# The huntingtin–HAP40 complex is a bidirectional cellular rheostat

**DOI:** 10.64898/2026.06.03.729778

**Authors:** Manuel Seefelder, Fabrice A C Klein, Enrico Calzia, Besnik Muqaku, Patrick Oeckl, Stefan Kochanek

## Abstract

Huntingtin-associated protein 40 (HAP40) is an obligate structural subunit of huntingtin (HTT) and is rapidly degraded when unbound, yet has been conserved across eukaryotes for over a billion years. Combining interactomics, quantitative respirometry, and transcriptomics, we show that the HTT-HAP40 complex functions as a bidirectional stoichiometric rheostat: unbuffered apo-HAP40 activates the Integrated Stress Response via ATF4 and DDIT3/CHOP, whereas unbuffered apo-HTT reciprocally drives cholesterol and fatty-acid biosynthesis through SREBF1/2. We identify the ER-mitochondria tether RMDN3 (PTPIP51) as a key HAP40 interactor, placing mitochondria-associated ER membranes (MAMs) at the rheostat’s convergence point, and demonstrate that HAP40 depletion specifically impairs respiratory complexes II/IV. Loss of rheostat balance reproduces transcriptional signatures of Huntington’s disease patient tissues, supporting a “dual failure” model in which collapse of stoichiometric buffering — rather than aggregation toxicity alone — drives pathogenesis. To our knowledge, this is the first obligate complex in which both unbound partners carry out distinct essential functions, defining stoichiometric buffering as a generalizable regulatory principle that couples complex assembly to metabolic and stress-response control across eukaryotes.

## Introduction

Huntington disease (HD) is a fatal inherited disorder characterised by a progressive loss of motor function, cognitive decline, and psychiatric symptoms (Burg *et al*, 2009). At its core, HD is a systemic pathology presenting with various non-neuronal symptoms including muscle atrophy, cardiac failure, and osteoporosis (Burg *et al*, 2009). The disease is caused by a CAG trinucleotide expansion in the huntingtin gene (HTT), resulting in a mutant protein with an expanded N-terminal polyglutamine (poluQ) tract. The 348 kDa HTT protein functions as a massive interactome hub, binding hundreds of partners (Saudou & Humbert, 2016; Harjes & Wanker, 2003; Haenig *et al*, 2020); yet, among this extensive network, Huntingtin-associated protein 40 (HAP40) occupies a unique niche, as it is to date the only known partner to form a stable, detergent-resistant complex with HTT involving all three domains of HTT (Guo *et al*, 2018; Harding *et al*, 2019).

Despite its centrality to HTT biology, HAP40’s physiological functions remain obscure (Seefelder *et al*, 2022) and existing reports are sparse; while some isolated studies link HAP40 to endosomal trafficking (Pal *et al*, 2006; Pal *et al*, 2008) or mitochondrial dynamics (Huang *et al*, 2017; Huang & Her, 2017), the molecular functions of HAP40 remain unknown. Despite the cytoplasmic instability of HTT-free HAP40 (apo-HAP40), numerous studies report that HAP40 can massively relocate to the nucleus upon overexpression or when HTT is downregulated (Peters & Ross, 2001; Milman & Woulfe, 2013). This suggests that apo-HAP40 may possess independent functions that are normally spatially restricted by HTT binding.

Unlike more transient effectors, HAP40 exhibits an obligate structural dependence on HTT. Cryo-EM structure and native mass spectrometry revealed that the non-covalent HAP40-HTT interface is exceptionally tight, resisting gas-phase dissociation even under conditions where the covalent HTT backbone begins to fragment (Guo *et al*, 2018; Harding *et al*, 2019; Harding *et al*, 2021; Alteen *et al*, 2023). Besides the extreme physical stability of the complex, there is also a strict biological hierarchy: HAP40 levels are linearly correlated with HTT, and most HAP40 is rapidly degraded upon HTT downregulation, whereas HTT stability is largely independent of HAP40 (Huang *et al*, 2021a; Harding *et al*, 2021; Xu *et al*, 2022). This co-evolved, high-affinity interaction, conserved from Drosophila melanogaster to humans (Seefelder *et al*, 2020), defines HAP40 not only as a binder, but as a subunit whose half-life depends on HTT binding. It also posits HAP40 as an archetype for obligate partners, i.e., proteins whose cellular stability is subordinate to their cognate binders (Huang *et al*, 2021a).

The existence of such extreme dependency raises a fundamental question: Why maintain a protein so unstable (HAP40) that it requires a massive energetic investment (HTT synthesis) to stabilise? We hypothesise here that this instability was selected as a potent regulatory feature. In this model, the HTT-HAP40 complex functions as a regulatory node, where HTT acts as a stoichiometric buffer (rheostat). Its role is to harness HAP40 for essential physiological tasks as a complex, while this functional sequestration simultaneously blocks alternate functions of apo-HAP40. Consequently, an imbalance between HAP40 and HTT, such as the progressive loss of HTT in HD, could trigger pathological effects both via the loss of the complex’s function and the dysregulation of apo-HAP40.

To test this hypothesis, we performed a comprehensive multi-omic characterisation (proteomics, transcriptomics) combined with functional studies in HAP40 knock-down cells, systematically distinguishing apo-HAP40 from the HTT-HAP40 complex. We show that apo-HAP40 and free HTT drive opposing transcriptional programmes for a specific set of modules — notably the integrated stress response (ISR) and SREBF-driven lipid biosynthesis — coupling these axes through a stoichiometric equilibrium. Disrupting this equilibrium produces pathological signatures that align with hallmarks of HD, supporting a model in which the HTT–HAP40 complex functions as a bidirectional rheostat whose failure contributes to neurodegeneration through both metabolic collapse and chronic stress signalling. More generally, our findings also underscore how tightly regulated obligate partner systems can act as rheostats that control potent biological activities and that can become catastrophic drivers of disease when their regulation fails.

## Results

To comprehensively map HAP40′s protein interactions, we devised a dual AP-MS strategy (Figure 1 A). We combined an *in vivo* pull-down using Strep-tagged HAP40 expressed in HEK293T cells with an *in vitro* pull-down using pre-purified GST-HAP40. Since the HTT-HAP40 complex cannot be reconstituted from its purified constituents *in vitro* (Guo *et al*, 2018), this dual design allowed us to systematically distinguish interactors of apo-HAP40 from those that may require the HTT-HAP40 complex. Both baits were robustly enriched over their matched controls (Figure 1 C), and the Bayesian integration of both assays defined a high-confidence interactome per experiment and a combined consensus set that recovers additional interactors whose evidence reached significance only when pooled across assays (Figure 1 B, D). Consistent with the rationale of the dual design, HTT was absent from the *in vitro* interactome but ranked as a top interactor in the *in vivo* pull-down (Figure 1 B, D). Stoichiometric analysis (iBAQ) revealed that only 12% (median; range 7.86%–16.15%) of overexpressed HAP40 assembled into a complex with HTT (Figure 1 E), establishing apo-HAP40 as the dominant species. Thus, even the *in vivo* dataset primarily reflects the apo-HAP40 interaction landscape. Functionally, the HAP40 interactome resolves into three dominant hubs — mitochondrial biology, nuclear gene-silencing machinery, and mitotic regulators (Figure 1 F) — each of which is dissected in the following sections.

**Figure 1:**
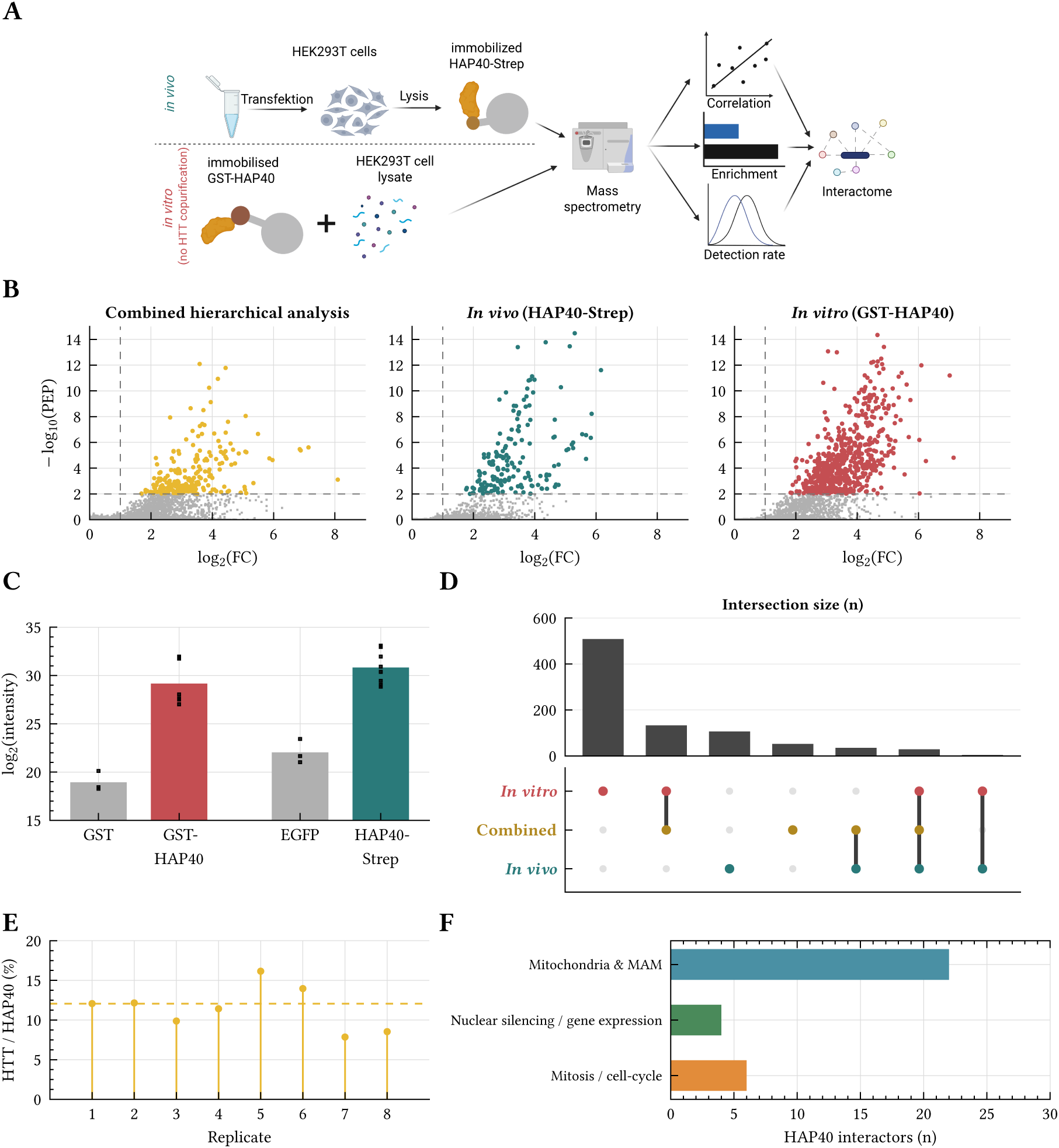
A dual AP-MS strategy based on strong stoichiometric dominance of apo-HAP40 reveals the core HAP40 interactome. **(A)** Experimental design. An *in vitro* pull-down using purified GST-HAP40 immobilised on GSH-sepharose (coral) and an *in vivo* pull-down of HAP40-TwinStrep overexpressed in HEK293T cells (teal) were processed through a common LC-MS/MS pipeline and integrated by the BayesInteractomics hierarchical Bayesian framework (PEP ≤ 0.01). GST-only and EGFP served as matched controls (*n*=3 replicate experiments per condition). **(B)** Volcano plots for the combined Bayesian model (gold), the *in vivo* (teal) and *in vitro* (coral) datasets. Dashed lines mark the hit thresholds. HTT is recovered only in the *in vivo* dataset, consistent with the known failure of the HTT-HAP40 complex to reconstitute from purified components (Guo *et al*, 2018). **(C)** Bait enrichment. log_2_-intensity of HAP40 (F8A1) in bait vs. matched-control pull-downs across all replicates; dots show individual replicates. **(D)** UpSet plot of high-confidence hits across the three analyses (*in vitro n*=674, *in vivo n*=174, combined *n*=249). Top bars show the size of each non-empty intersection; the dot matrix below indicates which sets contribute, with connected dots marking joint membership. HTT is captured exclusively by the *in vivo* assay. **(E)** Stoichiometric analysis. Per-replicate iBAQ-derived HTT:HAP40 molar ratio in the *in vivo* pull-down, ranging from 7.86% to 16.15% (median 12.07%), establishing apo-HAP40 as the dominant species. **(F)** Functional summary. Number of combined high-confidence interactors in three dominant hubs: mitochondria & mitochondria-associated ER membranes (MAM, blue), nuclear silencing / gene expression (green), and mitosis / cell-cycle regulators (orange).

### HAP40 regulates mitochondrial bioenergetics

Our proteomic analysis positions HAP40 as a central integrator of mitochondrial biology, revealing a deep and multifaceted engagement with the organelle’s homeostatic machinery that extends well beyond simple association. Gene Ontology (GO) analysis of the HAP40 interactome provided the first high-level evidence of this integration, showing a robust enrichment of mitochondrial proteins within the interactome (GO:0005739, FDR = 0.0051, 2.0-fold enriched), with a specific and striking over-representation of proteins localising to the mitochondrial outer membrane (GO:0005741, FDR = 3.85×10^−5^, 6.3-fold enriched). This mitochondrial signal extends across the broader mitochondrial membrane system (GO:0031966, FDR = 4.74×10^−5^, 3.3-fold enriched) and converges on bioenergetic substructures, including the inner mitochondrial membrane protein complex (GO:0098800, FDR = 0.0041, 5.4-fold enriched) and respiratory chain complex I (GO:0005747, FDR = 0.013, 9.5-fold enriched). Dissecting this enrichment revealed a network architecture organised around three core functional hubs: organelle dynamics, genetic maintenance, and bioenergetic structural assembly (Figure 2).

**Figure 2:**
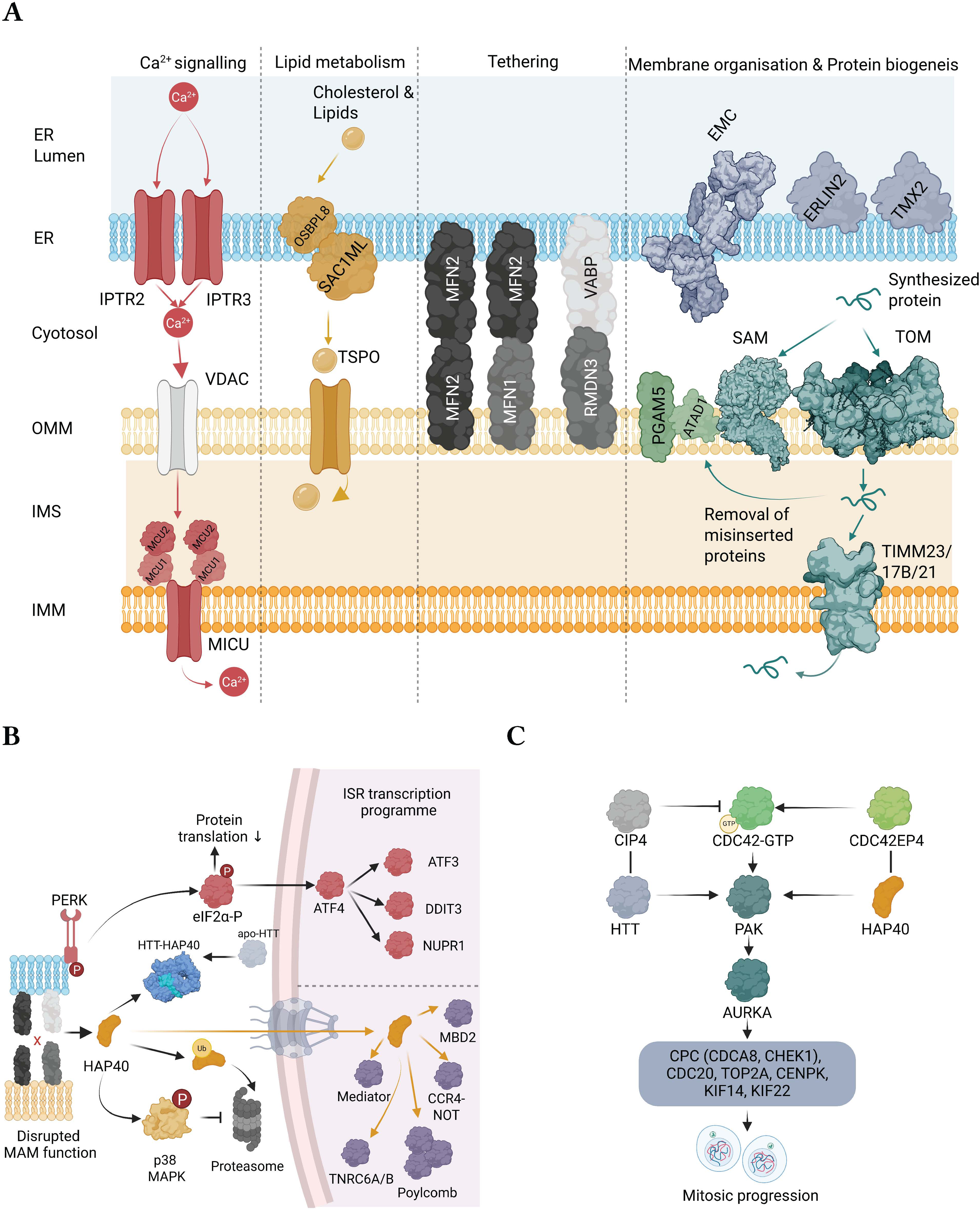
HAP40 as an organiser of the ER–mitochondria interface and nuclear stress coupling. **(A)** Schematic representation of HAP40 interactors at mitochondria-associated ER membranes (MAMs). HAP40-interacting proteins (coloured) are positioned at their established subcellular localisations. The ER membrane harbours the IP3 receptors ITPR2 and ITPR3, the lipid transfer proteins OSBPL8 and SACM1L, and the membrane organisation components EMC1, EMC9, ERLIN2, and TMX2. The MAM cleft is bridged by the RMDN3 (PTPIP51)–VAPB tethering complex and MFN2 homo-/heterodimers. The outer mitochondrial membrane hosts TSPO and PGAM5, while the inner membrane contains the calcium uniporter complex MCU–MICU2. Arrows indicate calcium (Ca^2+^) flux from ER to mitochondria and cholesterol transfer routes. Proteins in grey (VAPB and VDAC) were not identified as HAP40 interactors but are shown for structural context. **(B)** Model integrating MAM dysfunction with the nuclear stress response. Disruption of MAM contacts or MAM function liberates apo-HAP40, which is either cleared via ADRM1-mediated proteasomal turnover or imported into the nucleus; p38 MAPK activation biases the balance towards nuclear translocation (Huang *et al*, 2017; Huang *et al*, 2021c). Nuclear HAP40 engages the silencing apparatus (TNRC6A/B, CCR4-NOT, Polycomb, Mediator), while ATF4 drives induction of ATF3, DDIT3, and NUPR1 (Figure 4 C); a direct mechanistic link between these two arms remains unresolved. **(C)** Apo-HAP40 acts as a checkpoint effector in the CDC42–PAK–AURKA axis: binding to CDC42EP4 dampens CDC42 input to PAK1, phenocopying PAK1 knockdown and AURKA inhibition (Figure 4 D).

In the context of dynamics and transport, HAP40 appears to physically bridge the machinery governing mitochondrial shape and motility. We identified high-confidence interactions with both key fusion mediators MFN1 and MFN2, the fission-associated protein GDAP1, and the critical mitochondrial transport adaptor RHOT1 (Miro-1). The simultaneous association with MFN1, MFN2, and RHOT1 is functionally important; RHOT1 is known to form a complex with MFN1/2 to anchor kinesin and dynein motors, thereby facilitating axonal transport (Guo *et al*, 2005; Misko *et al*, 2010; Wang & Schwarz, 2009). Consistent with this, the interactome is significantly enriched for the Miro GTPase Cycle (Reactome HSA-9715370, FDR = 0.039, 35-fold enriched). This spatial integration extends to ER–mitochondria contact sites through the identification of the tethering protein RMDN3 (PTPIP51), which mediates lipid transfer and calcium signalling at mitochondria-associated ER membranes (Stoica *et al*, 2014). Given that mitochondrial motility is often antagonistic to the fission machinery—which requires organelle arrest for the recruitment of the fission executioner Drp1—HAP40’s association with both transport adaptors and fission regulators suggests it may act as a rheostat within this motility-dynamics coupling. This aligns with a previous report linking HAP40 overexpression to mitochondrial fragmentation via altered Drp1 phosphorylation (Huang *et al*, 2017).

Beyond morphology, the *in vivo* interactome highlights a direct role in mitochondrial genetics and protein synthesis, evidenced by the capture of TFAM, the master regulator of mtDNA replication and transcription (Kozhukhar & Alexeyev, 2023; Song *et al*, 2023), and several core components of the mitoribosome, including MTIF2, MRPL33, and MRPL42.

This regulation extends to the bioenergetic core, linking HAP40 to the assembly and regulation of the respiratory chain (Figure 2 A). We detected multiple Complex I subunits (NDUFA1, NDUFB6, NDUFB8, NDUFB11) and, crucially, the hypoxia-responsive assembly factors HIGD1A and its paralogue HIGD1C, as well as the CoA-synthesising enzyme COASY. Notably, we also identified the mitochondrial calcium uniporter MCU and its regulator MICU2, providing a potential physical link between HAP40 and mitochondrial calcium homeostasis—a key determinant of respiratory chain activity. These interactions were further complemented by the mitophagy-related phosphatase PGAM5 and the cellular prion protein PRNP, which has been reported to localise in part to the outer mitochondrial membrane and to modulate mitochondrial homeostasis (Faris *et al*, 2017). Notably, our *in vitro* assay using purified proteins confirmed that a subset of these associations — including those with respiratory chain components, mitoribosomal units, and dynamics regulators — occurs independently of HTT. This assay also captured association with the protein import machinery, including the outer membrane receptor TOMM20 and the inner membrane translocases TIMM17B and TIMM21.

Collectively, these data define HAP40 not merely as a passive passenger of the HTT-HAP40 complex, but as a regulator of mitochondrial homeostasis, proposing a molecular basis for HAP40′s role in mitochondrial dysfunction.

To orthogonally assess which of these associations reflect direct binary contacts versus engagement through higher-order assemblies, we subjected all high-confidence interactors (PEP ≤0.01 and below 50% disordered residues in the individual predicted structures) to pairwise AlphaFold3 (Abramson *et al*, 2024) docking and scored each complex with the adapted 4-metric C2Qscore (Genz *et al*, 2025) (Supplementary Figure 6). Confident direct interfaces emerged for the outer-membrane import receptor TOMM20 (C2Qscore 0.10, BF_dock_ 1.29), the redox enzyme CYB5R1 (C2Qscore 0.32, BF_dock_ 8.80), and Complex I subunit NDUFB6 (C2Qscore 0.16, BF_dock_ 2.17). In contrast, pairwise docking did not resolve confident interfaces for the mitofusins MFN1 and MFN2, the MAM tether RMDN3, the calcium uniporter subunits MCU and MICU2, the cholesterol import regulator TSPO, or the inner-membrane translocases TIMM17B, TIMM21 and TIMM23 (all C2Qscore ≤0 and BF_dock_ <1) — despite their robust capture in the AP-MS assays. This pattern is consistent with engagement of assembled holocomplexes rather than isolated subunits — the MFN homo- and heterodimers, the VAPB–PTPIP51 tether, the MCU–MICU2 calcium uniporter complex, the TSPO–VDAC cholesterol-import assembly, and the TIM translocase channel — and reflects the known limitation of pairwise AlphaFold3 prediction in the absence of membrane context and obligate partner assemblies. Within the silencing and transcriptional hub, only a subset of interactors yielded predicted direct interfaces: PCGF3 (Polycomb, C2Qscore 0.24, BF_dock_ 4.41), KAT2A (C2Qscore 0.14, BF_dock_ 1.74), and Mediator components MED10 and MED18 (C2Qscore 0.19 and 0.12, BF_dock_ 2.72 and 1.46). Other components of the same network — including CBX4 (Polycomb), MED29 (Mediator), DCP1B and MBD2 — fell into the assembly-mediated range (C2Qscore ≤0), consistent with the silencing machinery being predominantly engaged as preassembled complexes rather than through individual subunit contacts. The GW182 scaffold TNRC6A produced an elevated peak ipTM (0.61) but with substantial inter-model disagreement (ipTM *σ* = 0.11), yielding a modest C2Qscore (0.17) and BF_dock_ (2.33). Its paralogue TNRC6B, with comparable MS confidence, did not yield a positive C2Qscore (−0.01, BF_dock_ 0.48). We therefore interpret the structural evidence for a direct HAP40-GW182 interaction as tentative at best, noting that AlphaFold3 did not converge on a single interface hypothesis for either paralogue. Within the mitotic hub, AURKA produced a well-converged moderate-direct signal (C2Qscore 0.24, BF_dock_ 4.13, ipTM 0.52 ± 0.01), providing some additional structural support for the CDC42-PAK-AURKA axis proposed in the interactome section — although this finding alone does not constitute definitive evidence of a direct binary interaction and should be validated biochemically. PAK1, in contrast, fell into the assembly-mediated range (C2Qscore −0.07), consistent with the transcriptional-phenocopy model in which PAK1 need not contact HAP40 directly. CDC42EP4 could not be structurally evaluated because 54 % of its residues fell below the pLDDT threshold for ordered structure, triggering the disorder quality gate — a finding consistent with its nature as an intrinsically disordered CDC42 effector.

Superposing each AF3 HAP40–prey model onto the wild-type cryo-EM HTT–HAP40 complex (PDB 6EZ8 (Guo *et al*, 2018)) shows that 260 of 265 predictions place the prey on HAP40 surface that is otherwise engaged by HTT in the holo-complex; the remaining 5 (CLN5, TYSND1 and ATP6AP1 classified as accessible; CINP and NUDT16 as partially accessible) dock onto the disordered N-terminus 27–42 rather than onto any structured alternative patch (Supplementary Figure 7, Supplementary Figure 9). Because the HTT footprint is large (covering 124 of the 268 resolved HAP40 residues, 46 %) and contiguous, a randomly placed interface would overlap it appreciably by chance, so this binary classification alone is a weak test. We therefore compared each prediction against a size-matched null of random contiguous surface patches (Methods): the predicted prey interfaces overlapped the HTT footprint substantially more than chance (median overlap 0.75 versus 0.41 expected; 184 of 263 evaluable predictions exceeded the null at FDR < 0.05, and 189 of 263 against a simpler uniform null; Supplementary Figure 8, Supplementary Table 1), confirming that the interactome preferentially engages the surface HTT vacates rather than the HAP40 surface at large. The HTT footprint is itself essentially mutation-independent (Cohen’s *κ* ≥ 0.78 against 6EZ8 over the Q46 7DXJ (Huang *et al*, 2021b), Q128 7DXK (Huang *et al*, 2021b), and the higher-resolution wild-type 6X9O (Harding *et al*, 2021) reconstructions on shared residues). Because apo-HAP40 is the dominant species in the *in vivo* pull-down (median HTT:HAP40 iBAQ ratio 12.07%, Figure 1 E) and no HTT is captured at all in the GST-HAP40 *in vitro* pull-down (Figure 1 B, D), the geometric picture is internally consistent: the recovered prey partners engage HAP40 predominantly through the same surface that HTT vacates upon dissociation, and/or through the disordered N-terminus 1–41 and central loop 217–257 that are not visible in 6EZ8.

### HAP40 organises the ER–mitochondria interface

Strikingly, the HAP40 interactome extends beyond the mitochondrial matrix and inner membrane to encompass a broad interface with mitochondria-associated ER membranes (MAMs) (Figure 2). Beyond the tethering protein RMDN3 (PTPIP51), which mediates ER–mitochondria contacts via the VAPB–PTPIP51 complex (Stoica *et al*, 2014), and the dual MAM tether MFN2, we identified high-confidence interactions spanning the core functional axes of these contact sites. On the calcium signalling axis, HAP40 associates with the ER-resident IP3 receptors ITPR2 and ITPR3, which govern calcium release at MAMs, as well as the mitochondrial calcium uniporter MCU and its regulator MICU2, which mediate calcium uptake on the mitochondrial side. This positions HAP40 at both ends of the ER-to-mitochondria calcium transfer pathway. On the lipid transfer axis, we detected the oxysterol-binding protein OSBPL8, the fatty acyl-CoA reductase FAR1, and the alkylglycerone phosphate synthase AGPS — enzymes catalysing ether lipid synthesis, a process that requires enzymatic cooperation between ER and peroxisomes at membrane contact sites. This lipid transfer interface is complemented by the mitochondrial translocator protein TSPO, which mediates cholesterol import into mitochondria from the outer membrane. The MAM interface further includes the PI4P phosphatase SACM1L, which regulates phosphoinositide signalling at contact sites, the ER membrane complex components EMC1 and EMC9, the ER lipid raft-associated protein ERLIN2, the MAM-resident thioredoxin TMX2, and the mitophagy-associated phosphatase PGAM5. The convergence of tethering, calcium, lipid transfer, and quality control components within a single interactome is consistent with a role for HAP40 at the ER–mitochondria interface.

### HAP40 knock-down impairs mitochondrial bioenergetics

The structural association with respiratory assembly factors and dynamics machinery predicted that HAP40 is essential for mitochondrial bioenergetic capacity. To test this, we performed high-resolution respirometry on HAP40-depleted HEK293T cells (Figure 3 B-F). The analysis revealed a broad and severe bioenergetic deficit in HAP40-depleted cells.

**Figure 3:**
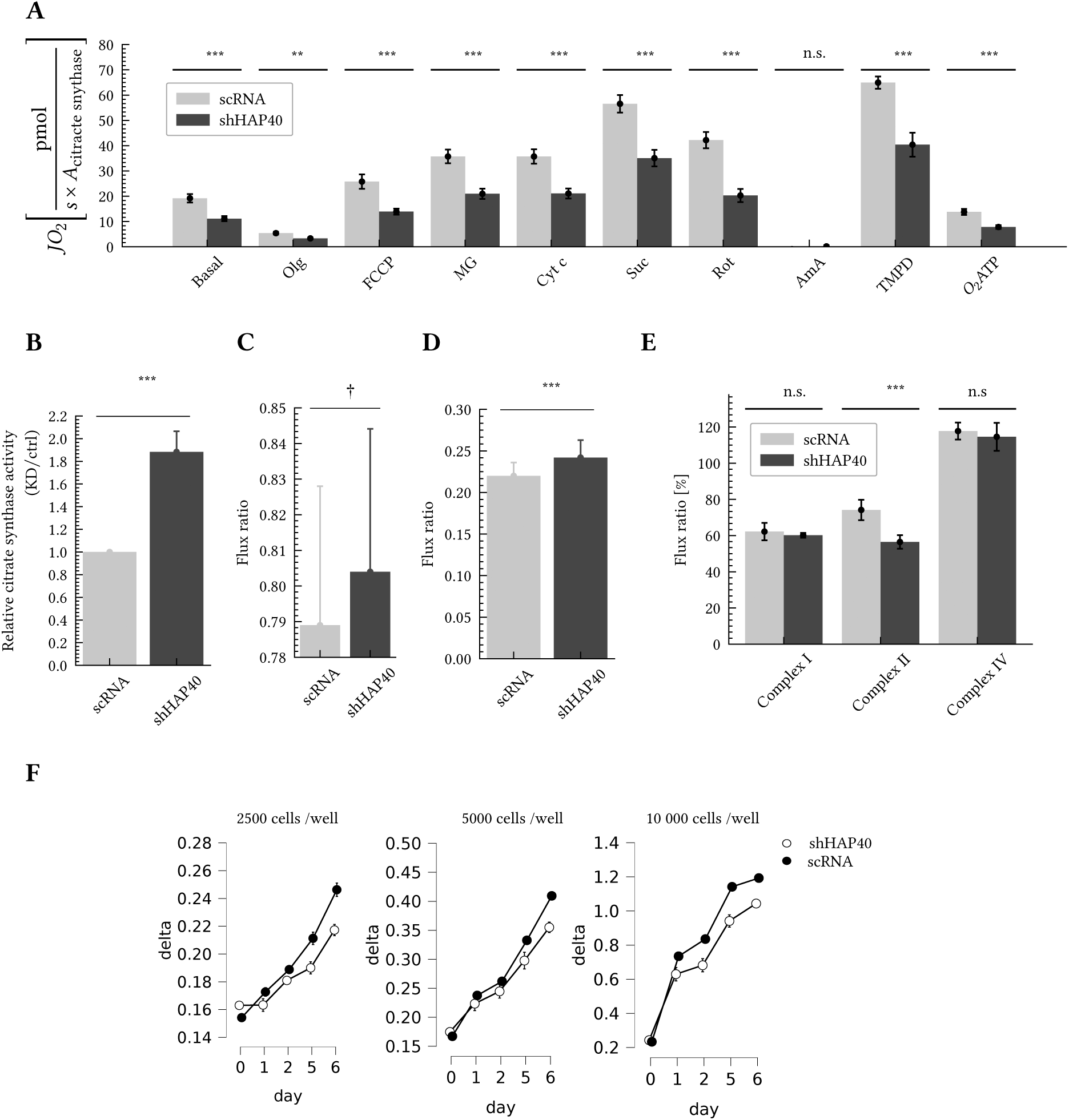
Functional assays in HEK293T cells with HAP40 knockdown or scramble shRNA. Error bars show 95% credibility intervals. (**A**) High-resolution respirometry. Olg: oligomycin, FCCP: carbonyl cyanide-p-trifluoromethoxyphenylhydrazone, MG: malate plus glutamate; Cyt c: cytochrome c; Suc: succinate, Rot: rotenone, Ama: antimycin A, TMPD: N,N,N′,N′-Tetramethyl-1,4-phenylenediamine, *O*_2_ATP: ATP-independent oxygen-flux **(B)** Relative citrate synthase activity in HAP40-KD versus control. **(C**) Fraction of maximal respiratory capacity utilised (basal / maximal) (N=7). **(D)** LEAK/U flux ratio. **(E)** Relative contribution of complex I, II, and IV (N=7) **(F)** Proliferation assay using WST-1 at three initial densities (N=3, 10 replicates; Bayesian ANOVA (JASP Team, 2025)). Statistical analysis with a Bayesian T-test. Bayes Factors (BF_10_) denoting evidence for difference between control (scRNA) and knock-down (shHAP40). † = Substantial evidence for null (BF_10_ 0.1 – 0.33) n.s. = Inconclusive evidence (BF_10_ 0.33 – 3), * = Substantial evidence for difference (BF_10_ 3 – 10), ** = Strong evidence for difference (BF_10_ 10 – 30), *** = Very strong/Decisive evidence (BF_10_ > 30)

We first observed elevated citrate synthase activity in HAP40-KD cells (Figure 3 B), indicating higher mitochondrial mass; all subsequent respirometry data were therefore normalised to citrate synthase activity. After this normalisation, HAP40-KD cells displayed a strong reduction in basal respiration (Figure 3 A). The reduced oxygen flux persisted after inhibiting ATP-synthase via oligomycin (LEAK state), and consequently ATP-linked respiration was significantly reduced (Figure 3 A). After uncoupling the electron transport system with FCCP, maximal respiratory capacity was also reduced in HAP40-KD cells (Figure 3 A). However, neither the R/U ratio (Figure 3 C) nor the LEAK/U ratio (Figure 3 D) were significantly altered, indicating that the coupling efficiency of the electron transport chain is preserved and that HAP40 depletion impairs total respiratory capacity rather than its utilisation.

Following permeabilisation with digitonin, the impairment was traced to specific respiratory chain components. Oxygen flux remained significantly impaired after the addition of the Complex I substrates malate and glutamate, and this defect persisted upon the addition of the Complex II substrate succinate and cytochrome c (Figure 3 B). To isolate Complex II-driven respiration, we inhibited Complex I with rotenone; the remaining oxygen flux was also reduced. In contrast, residual oxygen consumption after inhibition of Complex III with antimycin A showed no difference between groups, confirming comparable non-mitochondrial background levels. Direct assessment of Complex IV activity using TMPD and ascorbate confirmed a highly significant deficit (Figure 3 B). The consistent multi-complex respiratory failure confirms a fundamental bioenergetic defect resulting from HAP40 depletion.

To further dissect the respiratory chain defects, we calculated flux ratios normalised to the combined Complex I+II respiration (succinate state). The relative contributions of Complex I (cytochrome c / succinate) and Complex IV (TMPD / succinate) were not altered in HAP40-KD cells, whereas the relative contribution of Complex II (rotenone / succinate) was specifically reduced (Figure 3 E).

This functional defect aligns directly with the interactome findings showing HAP40′s association with the hypoxia-responsive assembly factors HIGD1A and HIGD1C, multiple Complex I subunits (NDUFA1, NDUFB6, NDUFB8, NDUFB11), and the mitochondrial calcium uniporter complex (MCU, MICU2).

### Apo-HAP40 modulates the AURKA-PAK1 axis

Our dual interactome analysis identified a robust, HTT-independent physical association between apo-HAP40 and the machinery controlling cell division, supported by significant enrichment for the centromeric region of chromosomes (GO:0000775, FDR = 4.74×10^−5^, 5.6-fold enriched), chromosome segregation (GO:0007059, FDR = 0.019, 4.4-fold enriched), sister-chromatid cohesion resolution (Reactome HSA-2500257, FDR = 0.020, 6.0-fold enriched), and metaphase plate congression (GO:0051310, FDR = 0.046, 8.7-fold enriched). While the *in vivo* analysis highlighted a striking enrichment for the Chromosome Passenger Complex (CPC), the *in vitro* screen using purified proteins revealed that this is driven by engagement of a dense network of mitotic regulators in the absence of HTT. The combined interactome captured the structural machinery of chromosome segregation, including CDCA8 (Borealin), a core CPC component identified in both screens, the centromere protein CENPK, Topoisomerase IIα (TOP2A), the mitotic kinesins KIF22 and KIF14 (mediating chromosome congression and cytokinesis, respectively), and the CDC42 effector CDC42EP4. Layered on this structural backbone, the *in vitro* screen further captured association of apo-HAP40 with the master mitotic regulator Aurora Kinase A (AURKA), the checkpoint kinase CHEK1, and CDC20, the key co-activator of the Anaphase-Promoting Complex/Cyclosome (APC/C) — interactions that occur in the absence of HTT and therefore represent bona fide apo-HAP40 binders.

To determine the functional necessity of these interactions, we assessed cell proliferation following HAP40 depletion. Consistent with diminished cell proliferation, HAP40 knockdown resulted in a significant, time-dependent reduction in cellular growth rates compared to controls (BF_10_=80 834.18; Figure 3 F). However, it is the transcriptomic data that illuminates the directionality of this regulation. We performed a comparative analysis of the transcriptional signature induced by HAP40 overexpression against public datasets of kinase inhibition. Strikingly, the gene expression profile of HAP40-overexpressing cells showed a significant overlap with the transcriptional signatures defined by AURKA inhibition (MLN8237 treatment (Althoff *et al*, 2014)) and the knockdown of its upstream activator, PAK1 (p21-activated kinase 1) (Villamar Cruz *et al*, 2016) (Figure 4 D). This transcriptional “phenocopying” strongly suggests that an excess of unbuffered apo-HAP40 acts as a functional inhibitor of the AURKA/PAK1 signalling axis (Figure 2 C). Integrating these physical and functional associations, we propose a network topology where the HAP40-HTT complex acts upstream of the CDC42-PAK-AURKA cascade (Figure 2 C). Notably, our *in vitro* pull-down also identified CDC42EP4 (CDC42 effector protein 4) as an apo-HAP40 interactor in the absence of HTT, providing a physical link between apo-HAP40 and the CDC42 signalling axis. Collectively, these data support a model where apo-HAP40 acts as a checkpoint effector: it physically engages AURKA and the mitotic machinery, and may interfere with the interaction of HTT with PAK1 (Luo *et al*, 2007) or the CDC42-inhibiting protein 4 (CIP4) (Holbert *et al*, 2003).

**Figure 4:**
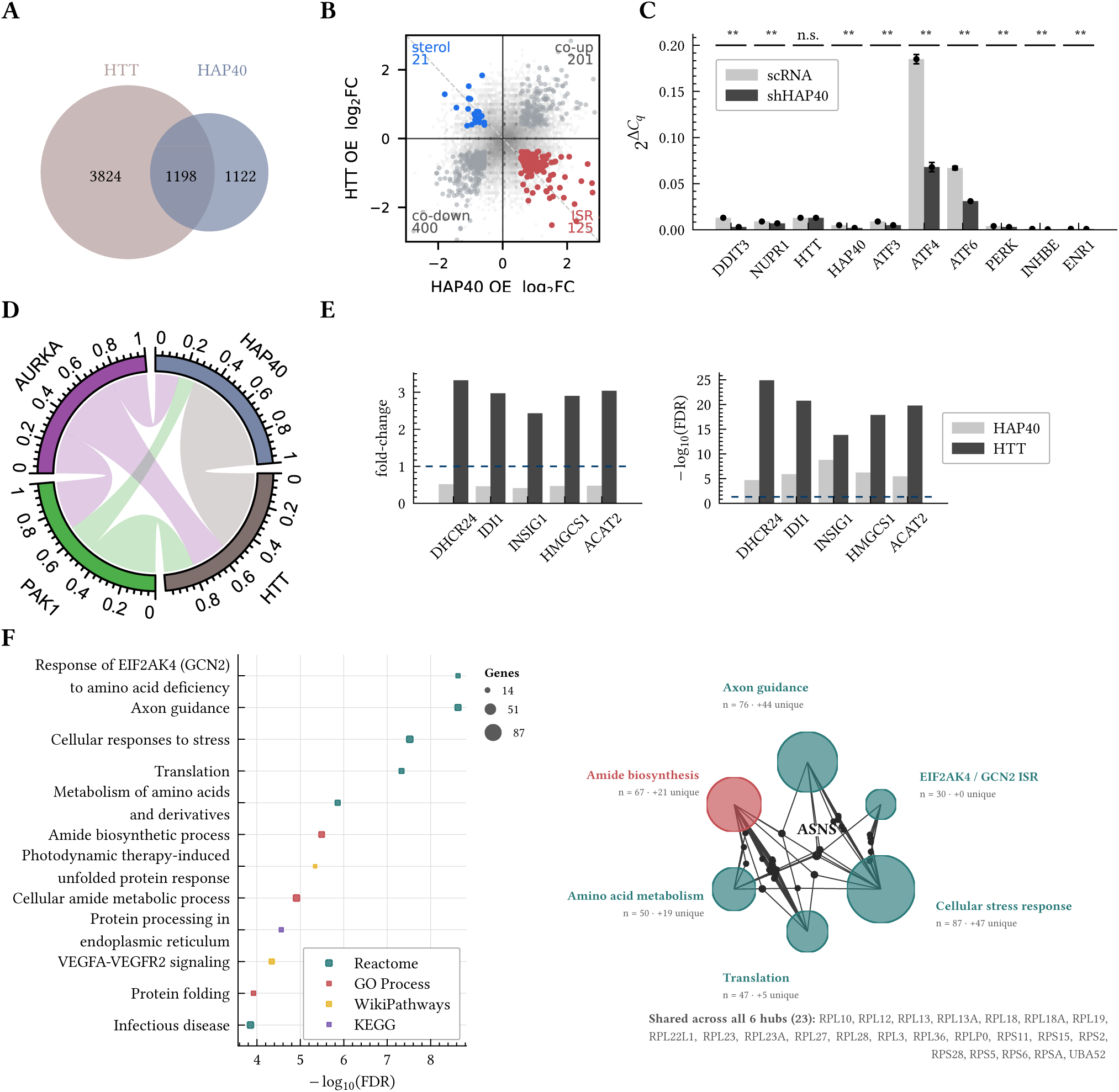
Reciprocal transcriptomic alterations upon HAP40 and HTT overexpression. **A**: Overlap of differentially expressed genes (DEGs) between HAP40 and HTT overexpression. **B:** Per-gene log_2_ fold changes upon HAP40 versus HTT overexpression: all common genes (grey density) and common differentially expressed genes (DEGs) (FDR < 0.05 in both, *n*=747) coloured by quadrant. The shared response is predominantly concordant (601 genes); the antagonistic subset is dominated by the ISR arm (HAP40-up / HTT-down, coral; 125 genes), with a smaller sterol arm (HAP40-down / HTT-up, blue; 21). Threshold-free RRHO map: Supplementary Figure 4. **C:** qRT-PCR of ISR regulators in HAP40-knockdown cells (*N*=3; mean ± SEM; Bayesian T-test). BF_10_: † (null, 0.1–0.33), n.s. (0.33–3), * (3–10), ** (10–30), *** (>30). **D:** Transcriptional overlap with PAK1 knockdown (Villamar Cruz *et al*, 2016) and AURKA inhibition (Althoff *et al*, 2014); ribbon width = shared gene fraction. **E:** Cholesterol biosynthesis genes with FDR≤0.05. **F:** STRING pathway enrichment of the 1198 reciprocally regulated differentially expressed genes (DEGs). Left: dotplot of the top 12 non-redundant terms (Reactome, GO Process, KEGG, WikiPathways; ≥80% containment, strength ≥0.2); dot size = gene count, colour = database. Right: cnetplot of the top 6 hubs (circle size = gene count); gene labels shown for nodes in ≥4 hubs; 23 ubiquitous genes shared across all 6 hubs are listed below.

### Apo-HAP40 binds silencing factors

Beyond the cell cycle, our analysis uncovered a high-confidence, HTT-independent association between apo-HAP40 and the core machinery of post-transcriptional gene silencing. The combined interactome identifies the GW182 paralogues TNRC6A and TNRC6B as top interactors, alongside the CCR4-NOT scaffold CNOT10 and the Polycomb chromobox protein CBX4. TNRC6 proteins function as the essential scaffolds for miRNA-mediated silencing and non-coding RNA-mediated transcriptional activation, bridging the Argonaute complex to downstream effectors (Hicks *et al*, 2017; Suzawa *et al*, 2017; Johnson *et al*, 2021). Consistent with this scaffolding role, the *in vitro* screen captured association of apo-HAP40 with the specific enzymatic machinery recruited by TNRC6, including the additional CCR4-NOT subunit CNOT11 (Hicks *et al*, 2017) and the mRNA decapping factors DCP1B and PATL1. This repressive network extended to the transcriptional level, evidenced by *in vitro* interactions with the Mediator complex (MED29, MED31) (Hicks *et al*, 2017) and chromatin regulators, including the additional Polycomb group proteins CBX6 and PCGF3 and the methyl-CpG binding protein MBD2. Crucially, the identification of TNRC6B as the top interactor in our combined analysis, alongside the preservation of these interactions in the *in vitro* assay, positions apo-HAP40 as an intrinsic, HTT-independent ligand of the nuclear silencing apparatus.

### HAP40 and HTT drive opposing programmes

Given the HTT-independent physical association of apo-HAP40 with nuclear silencing and stress-response machinery (TNRC6, CHEK1), we hypothesised that altering the HAP40:HTT ratio would drive a broad transcriptional reprogramming. To test this, we performed RNA-sequencing on HEK293 cells overexpressing HAP40, HTT, or a control vector. Both HAP40 and HTT induced substantial changes to the transcriptome, resulting in 2320 and 5022 differentially expressed genes (DEGs), respectively (FDR<0.05) (Figure 4 A). A statistically significant (*p*<2.2×10^−16^) overlap of 1198 genes was differentially expressed upon HAP40 and HTT overexpression (Figure 4 A).

Critically, a direct comparison of all common DEGs shows that the two programmes share a predominantly concordant overexpression footprint, yet contain a specific, mechanistically coherent antagonistic subset (Figure 4 B): genes driven up by HAP40 and reciprocally suppressed by HTT are dominated by the Integrated Stress Response (ISR) (ATF4, DDIT3, NUPR1, ASNS), whereas the reciprocal sterol-biosynthesis arm is smaller (Figure 4 E, Supplementary Figure 2). To determine if this functional antagonism extends beyond specific pathways and DEGs to the global transcriptome, we performed an Rank-Rank Hypergeometric Overlap (RRHO) analysis. Unlike threshold-based methods, RRHO compares the entire ranked list of transcriptional changes, allowing for a threshold-free assessment of global trends. Comparing the HAP40-induced versus HTT-induced transcriptomes revealed signal in both the diagonal (concordant) and off-diagonal (discordant) quadrants of the threshold-free RRHO map (Supplementary Figure 4). Alongside a broad concordant response — consistent with a shared overexpression-associated proteostatic footprint — a pronounced antagonistic subcomponent emerged in which genes strongly upregulated by HAP40 were reciprocally downregulated by HTT. This mirror-image subset tracks with the specific regulatory programmes dissected below (ATF4/DDIT3 arm of the ISR; SREBF-driven lipid biosynthesis; Supplementary Figure 2) and is mechanistically consistent with apo-HAP40′s physical engagement of the post-transcriptional silencing machinery (TNRC6A/B, CCR4-NOT, Mediator) and mitotic regulators identified in the interactome (Figure 1 F). To test this directly, we ran STRING functional-enrichment analysis on the concordant and the two discordant subsets (top-2000 genes per direction; Supplementary Figure 5). The concordant core was enriched for a non-specific mixture of pathways (cytoplasmic ribosomal proteins, axon guidance, Wnt signalling, generic cancer pathways and focal adhesion), with no single coherent mechanism emerging — compatible with a shared, non-directional response to protein overexpression in HEK293 cells rather than with specific rheostat biology. In stark contrast, the HAP40-dominant antagonistic arm was driven by ATF4/ISR biology (photodynamic-therapy-induced UPR, one-carbon metabolism, amino acid biosynthesis), and the HTT-dominant antagonistic arm converged on a single coherent module — sterol biosynthesis (Bloch/Kandutsch-Russell pathway). Together this supports the interpretation that the HAP40:HTT ratio acts as a bidirectional rheostat for two defined transcriptional modules — the ISR and lipid biosynthesis — rather than inverting the transcriptome as a whole.

To identify the upstream signalling nodes driving these reciprocal transcriptomic states, we performed transcription factor enrichment analysis using the TRRUST reference database (Supplementary Figure 2 A and Supplementary Table 7) and the CHEA3 database (Supplementary Figure 2 B and Supplementary Table 8). This analysis corroborated a striking “transcriptional mirroring” effect (Figure 4 B), where the HAP40-dominant and HTT-dominant states engage identical regulatory networks but drive them in diametrically opposing directions (Supplementary Figure 2). Specifically, the HAP40-induced transcriptome was dominated by the robust activation of key Integrated Stress Response (ISR) and Unfolded Protein Response (UPR) regulators. TRRUST analysis highlighted the master regulators ATF4 (*p*<10^−6^), ATF3, and XBP1 (Supplementary Figure 2 A and Supplementary Table 7), while ChEA3 independently validated this axis through the identification of DDIT3 (CHOP) (Supplementary Figure 2 B and Supplementary Table 8), the canonical pro-apoptotic effector of the ISR. This stress signature was paralleled by the enrichment of HSF1, the master regulator of the heat shock response, confirming that unbuffered apo-HAP40 acts as a potent proteotoxic stressor. Crucially, HTT overexpression produced the inverse signature, effectively suppressing these stress axes. Conversely, the metabolic failure observed upon HAP40 overexpression was mechanistically defined by the suppression of SREBF1 and SREBF2 (Supplementary Figure 2), the master transcription factors governing sterol and fatty acid biosynthesis. In the presence of excess HTT, however, SREBF1/2 targets were significantly upregulated, while they were downregulated upon HAP40 overexpression. Collectively, these data support a model in which the HTT-HAP40 complex is a bidirectional signalling rheostat where the ratio of apo-to-holo complex directly dictates the balance between anabolic growth (SREBF/E2F) and survival stress responses (ATF4/HSF1).

### Reciprocal ISR control by HAP40 and HTT

To understand the functional implications of these opposing transcriptional signatures, we performed Gene Set Enrichment Analysis (GSEA) and performed a network analysis of the resulting functional protein-protein interaction network formed by the DEGs (Figure 4 F). In HAP40-overexpressing cells, GSEA revealed a highly significant activation of the ISR and the eIF2α signalling pathway (Supplementary Table 6). This response appeared to be orchestrated by the robust upregulation of the master ISR transcription factor ATF4, and its partner ATF3. Consequently, we observed a broad induction of canonical ATF4 targets, including the pro-apoptotic factor DDIT3 (CHOP) and the feedback regulator PPP1R15A (GADD34). This stress signature extended to other key stress-responsive transcription factors, such as NUPR1 and JUND, as well as established stress markers like DDIT4, BBC3, and SESN2. In stark contrast, and consistent with a model of reciprocal antagonism, all of these key ISR drivers and stress marker genes were significantly downregulated upon HTT overexpression (Supplementary Table 6).

Crucially, this transcriptional reprogramming represents a specific activation of the ISR rather than a generic failure of cellular organelle homeostasis. We probed our dataset for markers of other organelle-specific stress responses and observed no significant enrichment of the Golgi stress response (GSR) pathways driven by TFE3 or CREB3 (Supplementary Table 5). Similarly, we detected no upregulation of the mitochondrial unfolded protein response (UPR_{mt}_) markers HSPD1 (Hsp60), HSPE1 (Hsp10), or DNAJA3 (Supplementary Table 6). This absence of broad organelle stress signalling confirms that unbuffered HAP40 does not induce non-specific toxicity, but rather selectively engages the cytosolic and ER-linked Integrated Stress Response.

To confirm that these changes were not artefacts of protein overexpression, we validated the expression of these ISR target genes in our earlier described (Huang *et al*, 2021a) stable HAP40 knock-down (HAP40-KD) cells (Figure 4 C) with robustly reduced HAP40 knock-down levels (Huang *et al*, 2021a). In direct opposition to the overexpression results, the loss of HAP40 led, as expected, to a significant downregulation of several stress markers, including ATF3, ATF4, ATF6, NUPR1, EIF2AK3 (PERK), INHBE, ERN1, and DDIT3 (Figure 4 C). This observed bidirectional regulation, where HAP40 overexpression activates the ISR while its depletion suppresses it, provides compelling evidence that a perturbation of HAP40′s functions significantly affects cellular homeostasis.

### Rheostat tunes cholesterol biosynthesis

Transcriptional downregulation of the mevalonate pathway is a well-established hallmark of HD, attributed to impaired SREBP1/2 activity (Sipione *et al*, 2002; Valenza *et al*, 2005; Valenza *et al*, 2007a; Di Pardo *et al*, 2019). Here, we demonstrate that this pathway is subject to bidirectional regulation by the HAP40-HTT axis. We observed a strong upregulation of cholesterol biosynthesis genes, including the rate limiting enzyme of the mevalonate pathway — 3-hydroxy-3-methylglutaryl-CoA synthase 1 (HMGCS1) — upon HTT overexpression, whereas the same genes were downregulated upon HAP40 overexpression (Figure 4 E). Notably, our interactome analysis did not identify any interaction of SREBP1/2 with either HAP40 or the HTT-HAP40 complex. Given the known physical interaction between HTT and SREBP1/2 (Di Pardo *et al*, 2019), the absence of an observable HAP40- / HTT-HAP40-SREBP interaction strongly suggests that the observed transcriptional changes are driven by the availability of apo-HTT.

We propose that HAP40 overexpression shifts the stoichiometric equilibrium, sequestering the endogenous apo-HTT pool into the stable HAP40-HTT complex. This sequestration limits the availability of apo-HTT to function as a co-regulator for SREBP1/2, thereby leading to the transcriptional downregulation of cholesterol biosynthesis genes.

### Rheostat failure mirrors HD pathology

To determine the pathological relevance of the transcriptional reprogramming driven by the HAP40-HTT rheostat, we compared our DEG against the Huntington’s Disease Signature Database (HDsigDB) and performed a detailed concordance analysis using RRHO. The HAP40/HTT-regulated gene set significantly overlaps with the core transcriptional dysregulation observed in HD patients and mouse models (Supplementary Figure 1). Specifically, we found striking convergence with signatures from neuronally-differentiated HD patient fibroblasts (GSE84013, FDR=3.2×10^−12^), HEK293 cells expressing mutant Huntingtin (GSE78928, FDR=3.6×10^−11^), human HD astrocytes (FDR=2.8×10^−10^), and HD Grade 3 caudate nucleus (GSE3790, Adjusted *P*=6.6×10^−10^). This overlap was driven by ISR effectors and amino acid metabolism genes (ATF3, DDIT4, ASNS, MTHFD2), stress-response regulators (NUPR1, HSPA1A/B), and cell-cycle regulators (CDKN1A). The conservation of this signature across patient-derived tissues, glial markers, and our rheostat model indicates that the identified defects represent a fundamental engagement of the HAP40-HTT machinery rather than cell-type specific artefacts.

To analyse whether the directionality of transcriptional changes in HD brain tissue (Labadorf *et al*, 2015) is concordant with HAP40 or HTT dominance, we performed RRHO analysis (Supplementary Figure 3). HAP40 overexpression showed pronounced global discordance with the HD brain signature, with the RRHO signal concentrated in the discordant quadrants (maximum −log_10_(*p*)≈26), yet revealed a statistically significant concordant subset (Supplementary Figure 3 A): genes downregulated in both HAP40 overexpression and HD comprised the cholesterol biosynthesis regulators SREBF2, HMGCR, and HMGCS1, as well as fatty acid metabolism enzymes (ACAT2, FASN, ACLY), while concordantly upregulated genes included ISR markers such as DDIT4. Conversely, RRHO analysis of HTT overexpression against the HD dataset (Supplementary Figure 3 B) produced a mixed directional signal of markedly lower magnitude (maximum −log_10_(*p*)≈11): a modest concordant component (co-downregulated genes) coexisted with comparably strong antagonistic subsets mapping to the rheostat’s metabolic arm. Cholesterol biosynthesis regulators (HMGCS1, HMGCR, ACAT2) were downregulated in HD but upregulated by HTT, while stress regulators (DDIT4, NUPR1) were upregulated in HD but suppressed by HTT. Together, these data support a model where HD pathology represents a “dual failure”: the loss of HTT-driven anabolic support and the concurrent destabilisation of HAP40-dependent bioenergetics.

## Discussion

### The HTT-HAP40 complex as a metabolic rheostat

HD has long been viewed through the lens of toxic gain-of-function (Saudou & Humbert, 2016), yet the physiological function of HTT and especially of its main partner, HAP40, itself has remained elusive, obscured by HTT’s large interactome (Seefelder *et al*, 2022). Our data fundamentally reframe HTT and HAP40 not merely as scaffolds, but as primary effectors of a bidirectional metabolic rheostat governed by the obligate nature of HAP40′s partnership with HTT. We propose that the exceptional stability of the HTT-HAP40 complex evolved to enforce a strict stoichiometric coupling between two mutually exclusive cellular programmes: a homeostatic programme — driven by a stoichiometric balance of apo-HTT, apo-HAP40 and the HTT-HAP40 complex — and a stress state, triggered by the accumulation of apo-HAP40 and a loss of apo-HTT.

In this model (Figure 5), HAP40 acts as a molecular “fuse” — a metastable protein that is continuously degraded unless buffered by HTT. This stoichiometric buffering ensures that under basal conditions the cell prioritises growth and anabolism; however, upon dissociation or stoichiometric imbalance, the release of apo-HAP40 rapidly interferes with metabolic processes and activates an ISR response.

**Figure 5:**
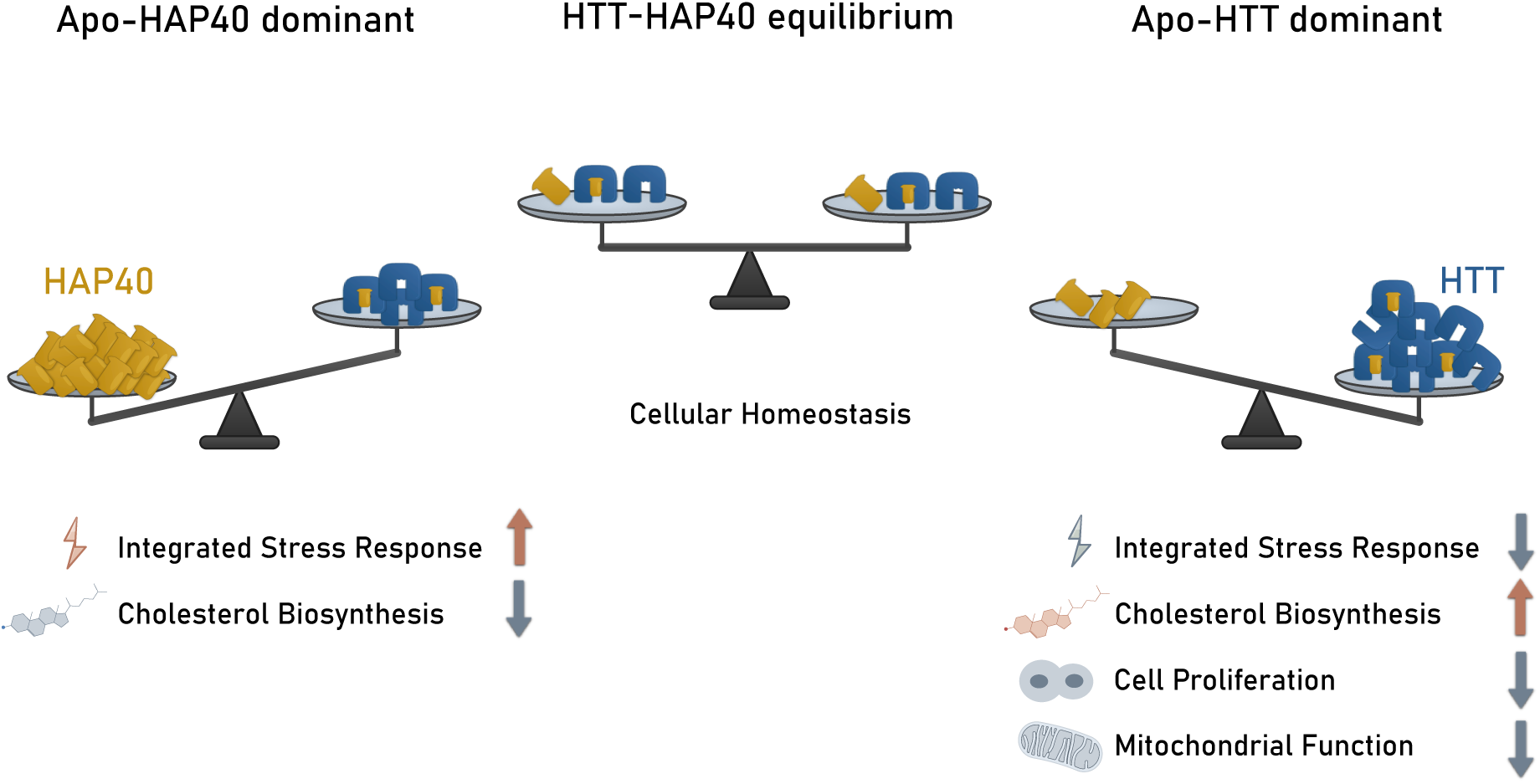
Model for HTT–HAP40 equilibrium-dependent regulation of cellular pathways. The stoichiometric balance between huntingtin (HTT, blue) and HAP40 (gold) governs downstream pathway activity, depicted as a balance scale across three states. When the HTT–HAP40 complex is intact (centre), cellular homeostasis is maintained. Excess free HAP40 (left) activates the integrated stress response (ISR) but suppresses cholesterol biosynthesis, whereas excess free HTT (right) leads to broad upregulation of ISR, cholesterol biosynthesis, cell proliferation and mitochondrial function. Upward orange arrows and warm-toned icons indicate pathway activation; downward grey arrows and cool-toned icons indicate suppression.

We utilised HEK293T cells as a robust human cell model to dissect the fundamental biochemical and stoichiometric properties of the HTT-HAP40 machinery. Given the evolutionary conservation of these proteins and their interaction from Drosophila melanogaster to humans (Seefelder *et al*, 2020) and the enrichment of the HAP40-HTT signature in HD patient-derived cells (Supplementary Figure 1), we posit that this rheostat represents a core cellular mechanism. In HD, the early degeneration of various brain structures, such as striatal neurons, may thus result from their disproportionate susceptibility to the failure of this bidirectional rheostat, likely driven by their exceptional bioenergetic demands (Joshi *et al*, 2025). In addition, it remains possible that the HTT-HAP40 complex may also play a role in the regulation of other neuron-specific cellular processes that have not been addressed in this study.

Our overexpression strategy was specifically designed to isolate the independent functional outputs of each unbuffered partner — a dissection that loss-of-function approaches alone cannot achieve, as HAP40 depletion necessarily follows HTT loss. Our iBAQ-based (Schwanhäusser *et al*, 2011) stoichiometric analysis confirms that even upon overexpression, only 12% (median; range 7.9–16.2%) of HAP40 assembles into the HTT-HAP40 complex (Figure 1 E), demonstrating that the system effectively interrogates the apo-HAP40 state. That this dissection captures physiologically relevant biology is supported by two independent lines of evidence: first, the qRT-PCR validation in HAP40 knockdown cells confirms the reciprocal regulation of ISR target genes (Figure 4 C); second, the transcriptional signature derived from our overexpression model significantly overlaps with datasets from HD patient tissues (Supplementary Figure 1 and Supplementary Figure 3).

### Cellular homeostasis relies on the synergistic action of apo-HTT and the HTT-HAP40 complex

We propose that the homeostatic state is not driven by a single species, but by the coordinated action of the holo-complex and the accessible apo-HTT and apo-HAP40 pools. Our data delineates a functional division of labour: HAP40 emerges as a key component of the mitochondrial homeostasis machinery, associating with the fusion and transport apparatus (MFN1, MFN2, RHOT1/Miro1) and required for full respiratory capacity (Figure 3), whereas the lipid biosynthetic arm is inferred to be driven by apo-HTT through SREBP1/2 — an interaction described previously (Di Pardo *et al*, 2019) that we did not detect for either apo-HAP40 or the holo-complex in our pull-downs, pointing to apo-HTT as the likely responsible species. Notably, the identification of the ER–mitochondria tethering protein RMDN3 (PTPIP51) as a HAP40 interactor suggests that this functional division may be physically coordinated at mitochondria-associated ER membranes (MAMs), where calcium and lipid transfer between the two organelles is regulated (Stoica *et al*, 2014). Physiological bioenergetics thus requires a precise equilibrium: sufficient holo-complex to stabilise the apo-HAP40 pool for mitochondrial maintenance, yet sufficient excess apo-HTT to drive cholesterol synthesis. Our findings are thus consistent with a dual failure model of HD: HAP40 depletion compromises mitochondrial integrity, while the loss of soluble HTT impairs cholesterol synthesis.

### The apo-HAP40 state drives stress adaptation

In contrast, the accumulation of apo-HAP40 functions as a stress sensor, initiating a multi-tiered stress response. We demonstrate that an excess of apo-HAP40 — mimicking a state where HTT levels are insufficient to buffer its obligate partner and where some HAP40 may consequently aberrantly relocate into the nucleus — triggers the ISR, characterised by the upregulation of ATF4, DDIT3 (CHOP), and PPP1R15A. Crucially, this is not merely a transcriptional event but a functional repurposing of the cell’s machinery. Building on earlier reports of a nuclear localisation of HAP40 (Peters & Ross, 2001; Milman & Woulfe, 2013; Seefelder *et al*, 2024), our interactome identifies apo-HAP40 as a binder of the nuclear silencing apparatus — including the GW182 scaffolds TNRC6A and TNRC6B, core subunits of the CCR4-NOT deadenylase complex (CNOT10, CNOT11) and components of the Mediator complex (MED29, MED31) (Figure 2 B) — concomitant with a broad downregulation of translational initiation and mRNA catabolic processes through the activation of the ISR. The identification of HAP40 at ER–mitochondria contact sites provides a spatial rationale for how organelle-level stress could be transduced to the nucleus: apo-HAP40 released from a disrupted MAM-associated pool would be readily available for nuclear import, linking loss of MAM integrity directly to the transcriptional stress response. This bidirectional rheostat is reinforced by the modulation of AURKA and CDKN1A. Together, these pathways constitute a coherent “Stress Adaptation” programme: when HTT is unavailable, apo-HAP40 coordinates the cessation of expensive anabolic processes and the activation of stress responses to counter the emerging deficits.

### Evolutionary implications

In the introduction we raised the question of why evolution would conserve a protein that is inherently toxic when unbuffered? Based on our data, we propose that the earlier observed intrinsic instability of apo-HAP40 (Huang *et al*, 2021a; Xu *et al*, 2022) serves a precise regulatory function, acting as a metabolic “fuse.” Under homeostatic conditions, HTT levels are sufficient to sequester HAP40 and tightly control levels of apo-HAP40, maintaining a homeostatic state. However, if HTT is depleted (e.g., during aggregation or increased turnover of mutant HTT) or functionally overwhelmed, apo-HAP40 accumulates. Because apo-HAP40 contains extensive disordered regions (Seefelder *et al*, 2020) and is rapidly degraded in the absence of HTT (Huang *et al*, 2021b; Xu *et al*, 2022; Harding *et al*, 2019), its signalling window is intrinsically transient. This design ensures that the “stress adaptation” state is self-limiting: the signal persists only as long as the imbalance remains, preventing a constitutively active stress response. The HAP40-HTT complex thus functions not just as a stabiliser, but as a sensor that couples the availability of its scaffold (HTT) to the metabolic decision to grow or arrest.

### Biological parallels

Unidirectional stoichiometric buffering systems involving obligate partners are well documented (Buchler & Louis, 2008). A prominent example is the Hsp90–HSF1 proteostasis rheostat, where Hsp90 sequesters HSF1 in an inert state and, upon proteostatic overload, releases it to trigger a transcriptional stress response (Barna *et al*, 2018; Guisbert & Morimoto, 2012). Our data suggest that HTT–HAP40 extends this principle to a bidirectional architecture: just as HTT buffers apo-HAP40 to restrain stress-related functions, HAP40 reciprocally buffers HTT to modulate lipid synthesis. The pathway decomposition of the RRHO (Supplementary Figure 5) supports this architecture at the module level: the antagonistic subsets map onto two mechanistically coherent axes — ATF4/ISR / one-carbon metabolism on the HAP40-dominant side and sterol biosynthesis on the HTT-dominant side — whereas the concordant core does not converge on any single coherent pathway, consistent with a non-specific response to protein overexpression rather than with rheostat biology. Thus, the HTT–HAP40 axis constitutes a bidirectional rheostat that couples two independent effector arms through a single obligate complex.

### Implications for HD

This model may offer a unified mechanistic explanation for the “double hit” observed in HD. Although the gain-of-function of FL-mHTT, mutant exon 1 fragments, aggregation, and somatic instability likely play central roles in disease aetiology, our results show that dysregulation of the HTT-HAP40 rheostat produces signatures that align strikingly with HD features. This underlines the significance of our findings, which add a further layer of understanding to the pathomechanisms underlying HD. First, the functional depletion of soluble HTT—via sequestration into aggregates or loss of wild-type allele function—impairs apo-HTT function. This creates a specific bottleneck in cholesterol synthesis, explaining the lipid deficits well-documented in HD models (Shankaran *et al*, 2017; Valenza *et al*, 2007a; Valenza *et al*, 2007b; Valenza & Cattaneo, 2005; Sipione *et al*, 2002; Valenza *et al*, 2015a; Valenza *et al*, 2015b). Second, the inability of mutant HTT to stabilise its obligate partner leads to the progressive degradation of HAP40 observed in patient tissues (Huang *et al*, 2021a; Xu *et al*, 2022; Harding *et al*, 2019). Our bioenergetic profiling demonstrates that this depletion is not benign; rather, the loss of HAP40 directly precipitates severe mitochondrial respiratory failure and Complex II/IV deficits (Figure 3). Moreover, the disruption of the HAP40–RMDN3 (PTPIP51) interaction at MAM contact sites may compound this failure by impairing ER-to-mitochondria calcium transfer and lipid exchange, thereby linking the bioenergetic and lipid biosynthetic arms of the rheostat at a single subcellular interface. This is consistent with recent evidence that the MAM-stabilising compound pridopidine rescues mitochondrial function in HD models via sigma-1 receptor activation (Naia *et al*, 2021) and is currently being evaluated in a phase 3 clinical trial for HD (Ko *et al*, 2026), underscoring the emerging pharmacological rationale for targeting organelle contact sites in neurodegeneration. This model also suggests that MAMs function as a subcellular sensor that couples organelle dysfunction to nuclear stress signalling: disruption of ER–mitochondria contacts — either through direct tether loss (RMDN3/MFN2) or impaired Ca^2+^/lipid transfer — may liberate apo-HAP40 from its MAM-associated pool and facilitate its relocation into the nucleus, where it engages the silencing machinery and triggers the ISR. Consistent with this, HAP40 depletion has been shown to activate p38 MAPK signalling (Huang *et al*, 2021c), a canonical upstream input into the ISR, while HAP40 overexpression induces mitochondrial defects via the proteasome-associated adaptor ADRM1 (Huang *et al*, 2017).

However, the stress signalling landscape in HD presents a complex layering of effects. Whereas pure HAP40 depletion suppresses the Integrated Stress Response (ISR) in stable knockdown cells (Figure 4 C), HD patient brains consistently exhibit a chronic, hyper-active ISR signature. We propose that in the disease state, the massive proteotoxic stress exerted by mutant HTT aggregates overrides the intrinsic ISR-suppressing effects of HAP40 loss, producing a composite “Dual Failure”: the metabolic collapse is driven by the breakdown of the HAP40-HTT rheostat (loss of mitochondrial and lipid maintenance), while the chronic stress signature is driven by the proteotoxicity of the mutant protein. Our HAP40 overexpression model is therefore best viewed as a proxy for this proteotoxic arm: by generating an unbuffered apo-HAP40 pool, it reproduces a high-stress environment analogous to that imposed by mutant HTT aggregates and demonstrates how an unchecked stress response may further exacerbate the metabolic blockade initiated by the rheostat failure.

Restoring the stoichiometric equilibrium of the HAP40-HTT complex, rather than targeting either protein in isolation, may thus offer a novel therapeutic strategy to simultaneously address the metabolic collapse and buffer the proteotoxic load.

## Methods

### Production and purification of GST-HAP40

Recombinant human HAP40, fused with an N-terminal Glutathione-S-transferase (GST) tag, and a GST-only control were expressed in E. coli. Bacterial pellets were lysed in a buffer containing 20 mM Tris-HCl pH 7.5, 150 mM NaCl, 10 mM EDTA, 5 mM MgCl_2_, 3 mM dithiothreitol (DTT), 1.7% (v/v) Triton X-100, 0.75 mg/mL lysozyme, and 17 µg/mL DNase I. The cleared lysate (10 000 × g) was incubated with Glutathione Sepharose 4B beads (Cat # 17-0756-01, Merck) for 2 hours at 4°C to allow binding.

After binding, the beads were washed three times with a total of 150 column volumes of 50 mM phosphate buffer (pH 7.0). The bead-bound proteins (GST-HAP40 or GST) were then covalently cross-linked to the glutathione resin using 1.5 mM disuccinimidyl suberate (DSS) in 50 mM phosphate buffer for 1 hour at room temperature. To remove any remaining non-covalently attached protein, the beads were subsequently washed with a stripping buffer of 0.2 M glycine-HCl (pH 2.6). The pH was immediately neutralised with 1 M Tris-HCl (pH 8.5), followed by a final wash with phosphate buffer. Successful covalent immobilisation of the bait proteins was confirmed by analysing a sample of the beads by SDS-PAGE and Coomassie Brilliant Blue staining.

### GST-HAP40 interactions assay

For each pull-down experiment, a pellet of approximately 3×10^8^ HEK293T cells was resuspended in ice-cold PBS supplemented with 0.5% (v/v) Tween-20 and 1x cOmplete™ Protease Inhibitor Cocktail (Roche, Cat# 11836170001). The cell suspension was lysed by sonication for five minutes on ice using a probe sonicator set to 70% amplitude (5 cycles). The resulting lysate was cleared by centrifugation at 10,000 x g for 10 minutes at 4°C.

To minimise non-specific binding of endogenous Glutathione-S-transferases, the cleared lysate was pre-cleared by sequentially incubating and loading it onto three GSH columns. Next, the lysate was applied to columns containing either covalently immobilised GST or GST-HAP40. The lysate was loaded onto the bait-bound beads and incubated for 30 minutes. After three washing steps, bound proteins were eluted with 0.1 M glycine (pH 2.6). Each fraction was immediately neutralised by the addition of 1 M Tris-HCl, pH 8.5.

### HAP40-Strep interactions assay

HEK293T cells were transiently transfected using polyethyleneimine (PEI) with expression plasmids encoding either human HAP40 with a C-terminal Twin-Strep-tag or EGFP as a negative control. Cells were harvested 48 hours post-transfection, and the protein lysate was prepared as described for the GST-HAP40 interaction assay. The cleared supernatant was loaded onto Strep-Tactin®XT 4Flow® resin (IBA, Cat # 2-5010-025).

Following incubation, the resin was washed three times with wash buffer (PBS supplemented with 0.1% (v/v) Tween-20). Bound protein complexes were eluted from the resin with an elution buffer consisting of PBS supplemented with 1.5 M MgCl_2_. The eluates were collected for mass spectrometry analysis.

### Mass spectrometry

Each sample was diluted to 500 µL with water and additionally 500 µL 1 M triethylammonium bicarbonate (TEAB) were added. Samples were spiked with 5 mM tris(2-carboxyethyl)phosphine hydrochloride (TCEP) and 10 mM 2-chloroacetamide (CAA), and proteins were reduced and alkylated by incubation for 30 min at 60 °C. Proteins were digested with 2 µg Trypsin/LysC (Promega) overnight at 37 °C. Digested peptides were purified using STAGE Tips (Affinisep SPE-Disks-Bio-DVB-47.20), dried overnight in a SpeedVac, and reconstituted in 12 µL 0.5% trifluoroacetic acid. Mass spectrometry analysis was performed with an UltiMate 3000 RSLCnano system and a Q Exactive mass spectrometer (both Thermo, Dreieich, Germany) by using the same equipment, solvent composition, and LC-MS settings as described previously for the analysis of tissue samples (Oeckl *et al*, 2019). Briefly, within each experiment (n=3) equal volumes of digested sample were injected onto a Thermo PepMap100 C18 trap column (75 µm × 2 cm, 3 µm, 100 A, C18) operating at 5 µL/min flow rate. Peptides were separated on a Thermo Acclaim PepMap analytical column (50 µm × 50 cm, 2 µm, 100 A, C18) by applying a step-gradient from 1-53% B over 160 min and a flow rate of 150 nL/min. MS data were acquired at a resolution of 70000 (MS) and 17500 (MS/MS) using a top15 DDA method. The data were searched against human database (downloaded from Uniprot on 12.01.2020) using MaxQuant software (Cox & Mann, 2008) (v. 1.5.2.8). Trypsin was set as the enzyme, allowing up to two missed cleavages. Carbamidomethylation was specified as a fixed modification, and protein N-terminal acetylation and methionine oxidation were specified as variable modifications. Protein intensities (iBAQ) reported by MaxQuant were used for statistical comparison between conditions.

### Statistical analysis

High-confidence protein interactions were identified from AP-MS data using our novel Julia-based framework, BayesInteractomics (https://github.com/ma-seefelder/BayesInteractomics). This pipeline integrates three evidence streams for each protein: enrichment (log2-fold change and p-value), bait-prey correlation across replicates, and a higher detection frequency in samples with the bait protein versus control samples using a Bayesian hierarchical model. A deep neural network (DNN) was used to generate a prior probability of interaction, which was then updated within a Bayesian two-component mixture model that combines the three evidence streams. The model uses copulas to learn the dependency structure between the evidence streams, and parameters were estimated using a Maximum a posteriori Expectation Maximisation (MAP-EM) algorithm. Final interactions were ranked based on the combined posterior error probability (PEP). Proteins with a PEP ≤0.01 were classified as interactors.

### Structural plausibility by AlphaFold3 docking

To orthogonally assess the structural plausibility of the identified HAP40 interactions, pairwise bait-prey complexes were predicted for all interactors passing the primary selection (PEP ≤0.01) using the AlphaFold Server (AlphaFold3) (Abramson *et al*, 2024). Predictions exceeding the 5000-token server limit per job or comprising more than 50% predicted disordered residues were excluded. For each pair, five predicted models were scored using an adapted C2Qscore (Genz *et al*, 2025), a linear combination of AlphaFold (AF) quality metrics calibrated for AF3 docking quality prediction.

The original C2Qscore is defined over five metrics: interface pLDDT, interface predicted aligned error (PAE), predicted TM-score, interface predicted TM-score (ipTM), and the Voronoi-based interface score VoroIF. In our implementation, VoroIF was omitted because it did not improve discrimination on the 1265-model AF3 benchmark — the 4-metric variant outperformed the full 5-metric score (AUC-ROC 0.929 vs. 0.919 AUC-PR 0.978 vs. 0.964) and also exceeded ipTM alone (AUC-ROC 0.924) and pDockQ2 (AUC-ROC 0.825); the logistic calibration was re-fitted accordingly. C2Qscore values were converted to a docking Bayes factor (BF_dock_) via the pre-fitted logistic calibration; independent quality gates were applied for fraction disordered (>0.5), mean pLDDT (<50), minimum interface PAE (>25 Å), and inter-model disagreement (ipTM standard deviation >0.15).

The docking Bayes factor served as a second Bayesian update to the MS-derived posterior, yielding a structurally refined combined posterior *P*_combined_=(odds_MS_⋅BF_dock_)/(1+odds_MS_⋅BF_dock_). We emphasise that a low C2Qscore or BF_dock_<1 does not constitute evidence against an interaction: because AlphaFold3 is trained predominantly on stable co-crystal structures deposited in the Protein Data Bank, its discriminatory power is systematically lower for transient, membrane-embedded, or assembly-mediated complexes — the very class of interactions that AP-MS is most sensitive to. Negative or low docking scores are therefore more consistent with an indirect or transient mode of association than with the absence of interaction. Both the MS-only and the structurally updated posteriors, together with the per-pair quality metrics and scoring tier used, are reported in the interactive BayesInteractomics report (Supplementary Table 2, Supplementary Table 3, Supplementary Table 4).

### Overlap of AF3 docking interfaces with the HTT–HAP40 interface

To assess whether the AlphaFold3-predicted prey-binding surface on HAP40 coincides with the surface engaged by HTT in the holo-complex, each predicted HAP40 chain was superposed onto the wild-type (Q17) cryo-EM HTT–HAP40 complex (PDB 6EZ8 (Guo *et al*, 2018)) using matchmaker in ChimeraX 1.11.1, and two residue sets on the superposed HAP40 were extracted: the 5 Å prey-contact set *C*_prey_ in the AF3 prediction, and the 5 Å HTT-contact set *C*_HTT_ in 6EZ8. To avoid crediting contacts placed in disordered regions not resolved in the cryo-EM reconstruction — the N-terminus 1–41, the central loop 217–257, and the short stretch 300–313 — the overlap fraction was computed on the 6EZ8-resolved portion *R*_6EZ8_ of HAP40 only: *f*_overlap_=|*C*_prey_∩*C*_HTT_|/|*C*_prey_∩*R*_6EZ8_|. Predictions were binned into three categories: non-accessible (*f*_overlap_≥0.30), partially accessible (0.10 – 0.30) and accessible (<0.10).

As a robustness check, the HTT interface was independently extracted from the higher-resolution wild-type structure of Harding *et al*. (PDB 6X9O (Harding *et al*, 2021)) and from the Q46 (PDB 7DXJ (Huang *et al*, 2021b)) and Q128 (PDB 7DXK (Huang *et al*, 2021b)) expanded-repeat cryo-EM structures; across the 227–268 HAP40 residues resolved in both 6EZ8 and each alternative, Jaccard similarities of the HTT-interface residue sets ranged from 0.78 to 0.90 and Cohen’s *κ* from 0.78 to 0.91, confirming that the interface definition is insensitive to the reference structure and to polyQ length. To ensure that the categorisation is not driven by arbitrary parameter choices, we performed a two-axis sensitivity analysis over the contact distance (4, 5, 6 Å) and the category thresholds (*f*_accessible_∈(0.05,0.10,0.15); *f*_non-accessible_∈(0.20,0.30,0.40)); the non-accessible fraction ranged from 87.5 % (4 Å, most permissive thresholds) to 98.9 % (6 Å, strictest thresholds), and the accessible count stayed ≤10 across all 27 cells, confirming that the main conclusion is parameter-insensitive.

To test whether the observed overlap exceeds chance given the large, contiguous HTT footprint, each prediction was additionally compared against a size-matched null. As real interfaces are spatially coherent, the primary null grew 4000 random connected surface patches of equal size per prey on the HAP40 C*α* contact graph (residues within 8 Å) and computed the empirical probability that a random patch overlapped the HTT footprint at least as much as observed (mean null overlap 0.41); p-values were Benjamini-Hochberg corrected across preys. A simpler uniform null, under which a prey is expected to overlap the 124 HTT-interface residues among the 268 resolved residues with fraction 0.46 (one-sided hypergeometric), gave concordant results. 184 of 263 preys (contiguity null) and 189 of 263 (uniform null) exceeded chance at FDR < 0.05.

### RNA-sequencing

Three HEK293 cell lines were used: HEK293TetOn, B1.21 (inducible overexpression of Huntingtin, HTT-Q17) (Huang *et al*, 2015), and B1.21HAP40 (constitutive expression of HAP40) (Guo *et al*, 2018). Cells were maintained in MEMα medium (Gibco) supplemented with 10% FBS (Gibco) at 37°C and 5% CO₂. HTT expression was induced with 1 µg/ml doxycycline for 48 hours. Protein expression was confirmed by Western Blot analysis using anti-HAP40 (SantaCruz Biotechnology #SC-69489) and anti-HTT (MAB2166, Millipore) antibodies.

Total RNA was extracted using the RNeasy Plus Mini Kit (Qiagen) following the manufacturer’s protocol. RNA integrity was verified on an Agilent Bioanalyzer 2100 system (RNA Nano 6000 Assay Kit). Library preparation and paired-end sequencing were conducted by Novogene Co., Ltd. using the NEBNext® Ultra™ RNA Library Prep Kit for Illumina® (NEB) with 1 µg total RNA per sample. Raw FASTQ reads were quality-filtered, aligned to the reference genome using HISAT2, and transcripts were assembled using the Cufflinks RABT method.

### Differential expression analysis

Differential expression analysis was performed using the DESeq2 R package. Genes with a Benjamini-Hochberg adjusted p-value (*p*_adj_)<0.05 were considered as DEG. To determine the statistical significance of the overlap between DEGs from HAP40 and HTT-Q17 overexpression, we used a hypergeometric test as implemented in the hypergea R package (Boenn, 2018). Genes with their respective log2-fold changes, found to be significant in both conditions, were visualised with a heatmap using the gplots R package (Warnes *et al*, 2024). Finally, to prioritise genes, we performed a robust rank aggregation analysis (Pihur *et al*, 2020; Kolde *et al*, 2012). Gene lists from each condition, sorted by both their absolute log_2_-fold-change and *p*_adj_, were aggregated to create a final ranked list of genes most affected by both HAP40 and HTT overexpression.

### Enrichment analysis

Gene Ontology (GO) term enrichment analysis for protein-protein interaction networks were performed using the tool in the STRING database (v11.0) with log_2_-fold-changes as input (Szklarczyk *et al*, 2022). Enrichment of transcription factors targets among HAP40 and HTT overexpressed genes were performed using EnrichR (Xie *et al*, 2021) and Chea3 (Keenan *et al*, 2019).

### RRHO analysis

To assess global transcriptional concordance and determine the extent of signal mirroring between datasets, RRHO analysis was performed using the RRHO2 R package (v. 1.0) (Cahill *et al*, 2018; Plaisier *et al*, 2010). Differential gene expression lists for HAP40 and HTT overexpression were processed to identify common gene identifiers. Genes were ranked based on a signed significance score, calculated as:

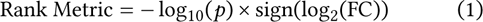

Raw p-values were used (rather than FDR-adjusted values) to preserve the full transcriptome and reduce tie density at the moderate-significance boundary. Duplicate gene identifiers were aggregated by calculating the mean of their ranking metric. The stratified overlap between the two ranked lists was computed using the RRHO2_initialize function with default parameters. The resulting hypergeometric *p*-values for each overlap threshold were log-transformed and visualised as a heatmap, where high signal intensity in the off-diagonal quadrants (top-left and bottom-right) indicates statistically significant discordant expression (transcriptional mirroring).

### Network analysis

Analysis of protein-protein interaction networks, retrieved from the STRING database (Szklarczyk *et al*, 2022) with a protein-protein interaction cut-off of 0.4, was conducted in Cytoscape (version 3.7.2) (Shannon *et al*, 2003) using genes identified by RRA and the top 50 hubs of the network were computed with the cytoHubba plug-in (Chin *et al*, 2014) using all eleven implemented scoring methods. Results of different scoring methods were aggregated by RRA (Kolde *et al*, 2012) to increase the robustness of the prediction. Hub proteins with an RRA score < 0.05 were considered further. Coexpression networks were retrieved with the GeneMania plug-in in Cytoscape (Shannon *et al*, 2003) and the top 50 hubs were computed as described before. Network communities, defined as a group of nodes that are more connected within themselves than with the rest of the network, represent functional modules and were computed with the community cluster algorithm (GLay) implemented in the Cytoscape plug-in clusterMaker2 (Shannon *et al*, 2003).

### Comparison with PAK1 and AUKRA transcriptome

Microarray data were downloaded from the NCBI GEO database with the accession number GSE71283 (Villamar Cruz *et al*, 2016) and GSE57810 (Althoff *et al*, 2014) and merged with their respective annotation information. Data normalisation was performed with full-quantile normalisation using the function normalise.quantiles.robust from the R-package preprocessCore (Bolstad, 2025) and transformed into Z-values. Z-ratios were computed as described (Cheadle *et al*, 2003) and genes with an absolute Z-ratio above 1.5 were considered as differentially expressed. For comparing murine with human data, the human orthologues of genes were retrieved using the R-package biomaRt (Durinck *et al*, 2009; Durinck *et al*, 2005).

### Quantitative reverse-transcription PCR (qRT-PCR)

shHAP40, shHTT cells and their control cell line were described previously (Huang *et al*, 2021a). RNA was isolated using the RNeasy Mini Plus kit (Qiagen) following manufacturer’s instructions and reversely transcribed into cDNA using the ThermoScript™ RT-PCR System with random hexamer primers. qRT-PCR was performed using KAPA SYBR® FAST qPCR Master Mix (2X) Kit. Forward (200 µM final concentration) and reverse primers (200 µM final concentration) were designed using the NCBI PrimerBlast tool and span exon-exon junctions for multi-exon genes to avoid detection of genomic DNA. Relative mRNA levels were normalised to 18S rRNA, hypoxanthine phosphoribosyltransferase 1 (HPRT1), and tubulin alpha 1a (TUBA1A). Statistical significance was determined by a Bayesian one-sided (*H*_*A*_:*μ*_KD_<*μ*_ctrl_), independent-samples Mann-Whitney U-Test based on computed 2_Δ*C*_*q* values using JASP (JASP Team, 2025).

### High-resolution respirometry

To remove the influence of antibiotics on mitochondrial respiration, control HEK293T cells expressing scramble shRNA and HAP40 knock-down cells (Huang *et al*, 2021a) were cultured in an antibiotic free medium four weeks prior to the experiment. Cellular oxygen uptake was quantified by high-resolution respirometry using the Oroboros Oxygraph-2K (Oroboros Instruments, Innsbruck, Austria). Cells were trypsinised, counted and resuspended in respiration medium. Respiration (JO_2_) was recorded as oxygen flux normalised to cell count (pmol*O*_2_/[*s*×10×10^6^cells]).A substrate-inhibitor-uncoupler-titration (SUIT) protocol was applied to intact cells. Basal respiration was recorded, followed by the addition of 1.25 µM Oligomycin to determine the LEAK-state. Maximum respiratory capacity (ETS) was then measured by titrating FCCP (1 mM stock, 1 nL/s via TIP-2K) to its optimum.Following this, cells were permeabilised (1.62 µM digitonin), and substrate-saturated respiration was measured with malate (2 mM), glutamate (10 mM), and succinate (10 mM). Finally, Complex I (0.5 µM rotenone) and Complex III (5 µM Antimycin A) were inhibited, and Complex IV activity was measured with ascorbate (2 mM) and TMPD (0.5 mM), correcting for auto-oxidation [49]. Data acquisition and analysis were performed with DatLab 7.0. Data of 7 independent experiments were analysed with JASP (JASP Team, 2025).

### Citrate Synthase Measurements

Control HEK293T cells expressing scramble shRNA and HAP40 knock-down cells (Huang *et al*, 2021a) underwent citrate synthase measurements as reported in the following protocol: http://wiki.oroboros.at/index.php/MiPNet17.04_CitrateSynthase. The measured values (Ultrospec 2100 pro, Amersham Biosciences) were normalised to the total cell number and to control cells. Data were analysed with a one-sided Bayesian Mann–Whitney U test in JASP (JASP Team, 2025).

### Cell proliferation assay

Cell proliferation was assessed using a WST-1 (Abcam) assay. Control (scramble shRNA) and HAP40 knock-down HEK293T cells were seeded in 96-well plates at initial densities of 2.500, 5.000, or 10.000 cells per well. The cells were cultured in antibiotic-free DMEM (Gibco) with 10% FBS, supplemented with 2.5% (v/v) WST-1 reagent.

Metabolic activity, correlating with cell proliferation, was quantified by measuring absorbance at 450 nm (signal) and 600 nm (background reference). Readings were taken at 1, 24, 48, 120, 144, and 192 hours post-seeding. The differential signal (ΔOD≔OD_450nm_−OD_600nm_) was used as the readout. A Bayesian ANOVA (JASP Team, 2025) was performed, with ΔOD as the dependent variable. The fixed effects included in the model were time (days), cell line, initial seeding density. The experiment number was included as a random effect to account for batch variability. The results of the WST-1 assay were independently confirmed by manual cell counting at the 192-hour endpoint using a Neubauer chamber.

### Declaration of generative AI and AI-assisted technologies in the manuscript preparation process

During the preparation of this work the authors used Claude (Anthropic) in order to assist with language editing (spelling, grammar, and phrasing). After using this tool, the authors reviewed and edited the content as needed and take full responsibility for the content of the publication.

## Data availability

All raw and processed data is available in the supplemental material.

## Code availability

The Julia package for the Bayesian analysis of the interactomes (BayesInteractomics, v1.1.6) used in this study is available on GitHub (https://github.com/ma-seefelder/BayesInteractomics) and archived on Zenodo (DOI: 10.5281/zenodo.19709177). During peer review, the Zenodo record is under restricted access; reviewers may obtain anonymous read-only access via the following preview link: zenodo.org/records/19709177 (anonymous preview). The record will be made openly available upon acceptance.

## Acknowledgements

This study was supported by the European Huntington Disease Network (EHDN) via their Lesley Jones Seed Fund Programme (project number: 1245). Further, M.S, F.K, and S.K. study was supported by the Deutsche Huntington Hilfe e.V. P.O. received research support from the ALS Association/ALS Finding A Cure (24-SGP-691, 23-PPG-674-2), Charcot Foundation (D.7090), Cure Alzheimeŕs Fund, DZNE Innovation-to-Application (I2A_call9_Oeckl), EU Horizon Europe (SYNAPSING), consulting fees from LifeArc and Fundamental Pharma, honoraria for lectures from GSK and travel support from Biogen. We thank Stephen Meier for excellent technical assistance. Figure 1 A and Figure 2 were created in BioRender. Seefelder, M. (2026) https://BioRender.com/6njc2pg.

## Author contributions

M.S., F.K., and S.K. conceptualised the study, with M.S. leading the design of the integrative omics strategy. M.S. and F.K. performed the GST-HAP40 and HAP40-Strep interaction assays. B.M. and P.O. conducted all mass spectrometry experiments and processed the raw data. E.C. and M.S. performed the mitochondrial respiration measurements. M.S. designed and carried out all bioinformatic and statistical analyses, including Bayesian interactome modelling, protein-protein interaction prediction, over-representation analysis, and candidate/pathway prioritisation. M.S. developed the BayesInteractomics Julia package used for interactome inference. M.S. performed RNA-sequencing of HAP40 and HTT overexpression cells and led all downstream transcriptomic analyses, including differential expression, pathway enrichment, and cross-omics integration. M.S. led the integration and interpretation of all datasets, produced all figures, and wrote the original draft. All authors reviewed, edited, and approved the final manuscript.

## Additional Information

**Correspondence and requests for materials** should be addressed to Manuel Seefelder.

## Supplementary Information

**Supplementary Figure 1:**
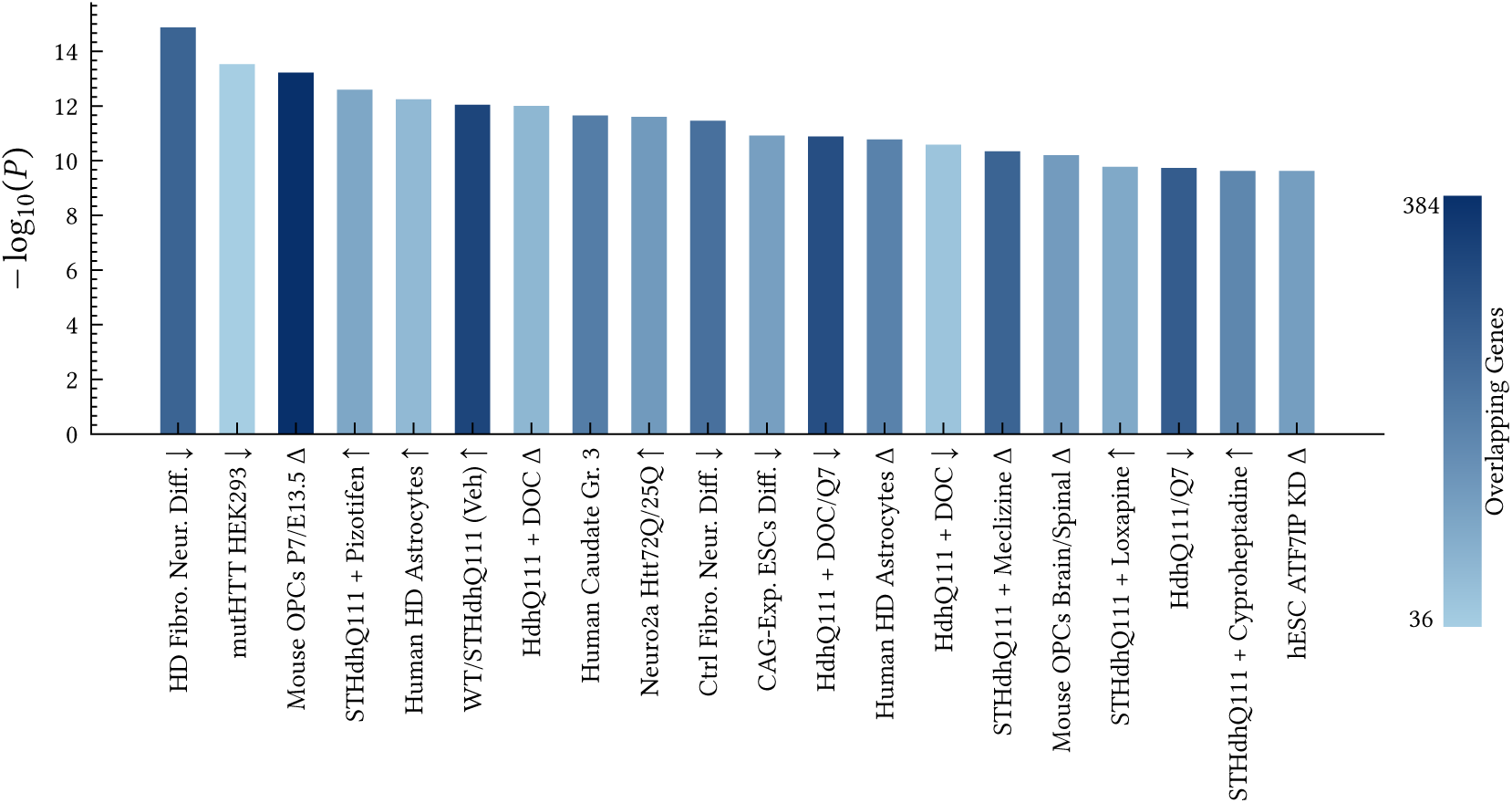
Enrichment of the HAP40/HTT-regulated gene set against the HDSigDB reference. The reciprocally regulated differentially expressed genes (DEgs) from HAP40 and HTT overexpression were queried against the Huntington’s Disease Signature Database (HDSigDB). Bar height encodes −log_10_(*p*) of the overlap; colour encodes the number of overlapping genes per signature. Arrows in the term labels indicate the direction of regulation in the reference dataset (↑ up, ↓ down, Δ changed).

**Supplementary Figure 2:**
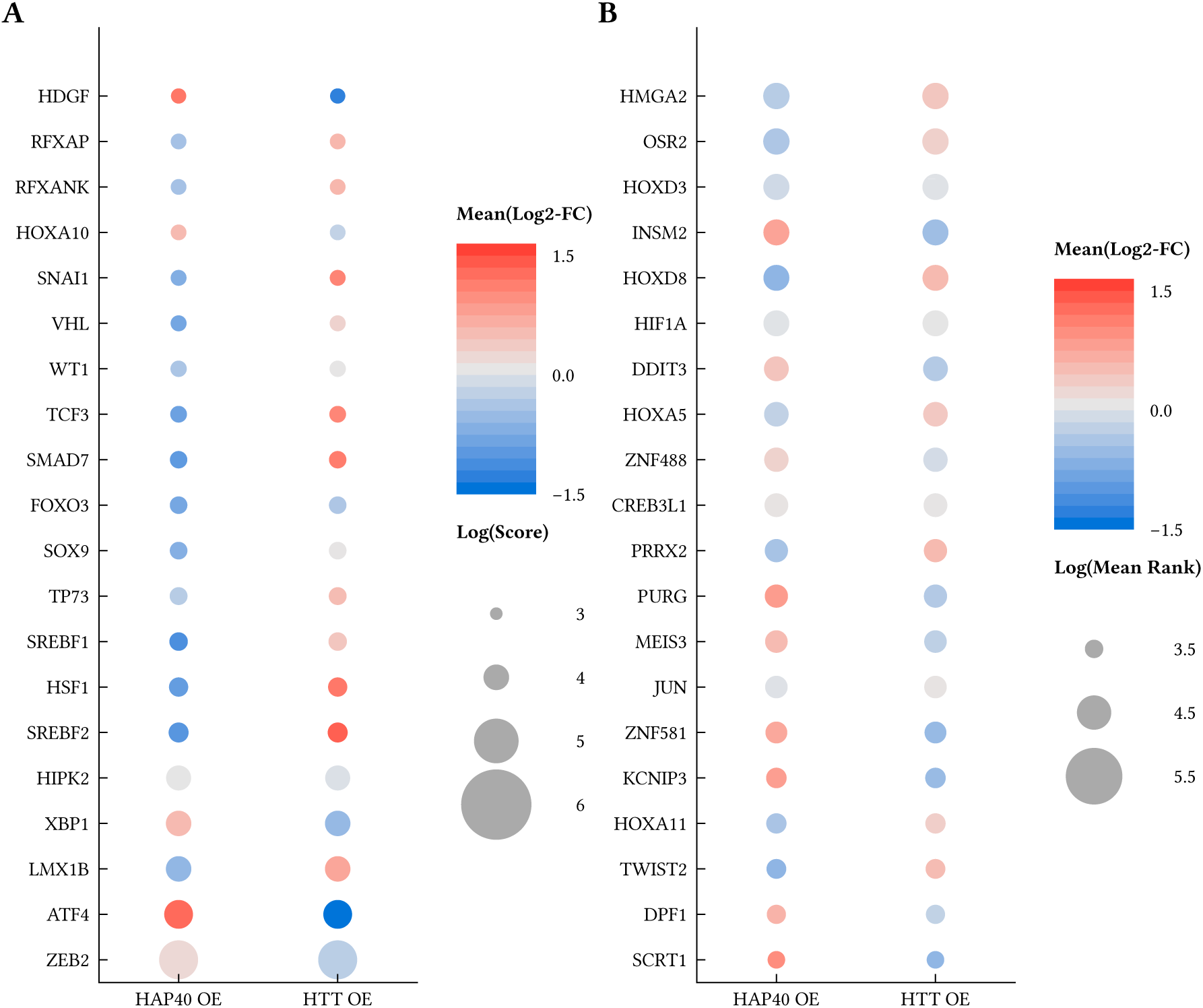
Transcription factor enrichment analysis of HAP40 and HTT overexpression. Enrichment of transcription factor target genes among differentially expressed genes upon HAP40 and HTT overexpression, queried against the TRRUST (A) and CHEA3 (B) databases. Bubble size represents the log(enrichment score) of target genes in the overlap; colour indicates enrichment significance mean log2-fold changes of the target genes.

**Supplementary Figure 3:**
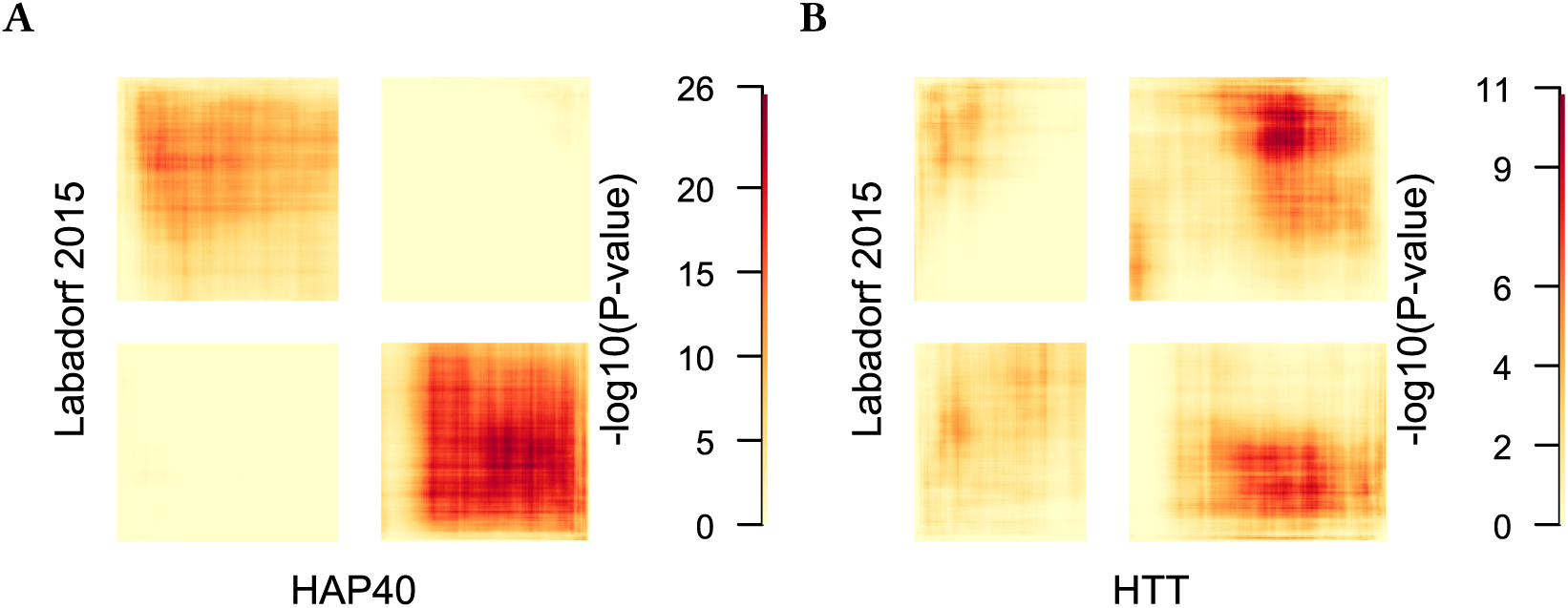
Directional concordance between the HAP40-HTT rheostat and HD brain transcriptomes. RRHO analysis comparing transcriptional changes upon HAP40 (A) or HTT (B) overexpression with changes in the cerebral cortex (Brodmann areas 4 and 9) of HD patients (Labadorf *et al*, 2015). Genes are ranked by signed significance [−log_10_(*p*)×sign(log_2_(fold change))]. Colour intensity represents −log_10_(hypergeometric p-values). Signal in the bottom-left and top-right quadrants indicates concordant regulation (co-up or co-down); signal in the top-left and bottom-right quadrants indicates discordant regulation.

**Supplementary Figure 4:**
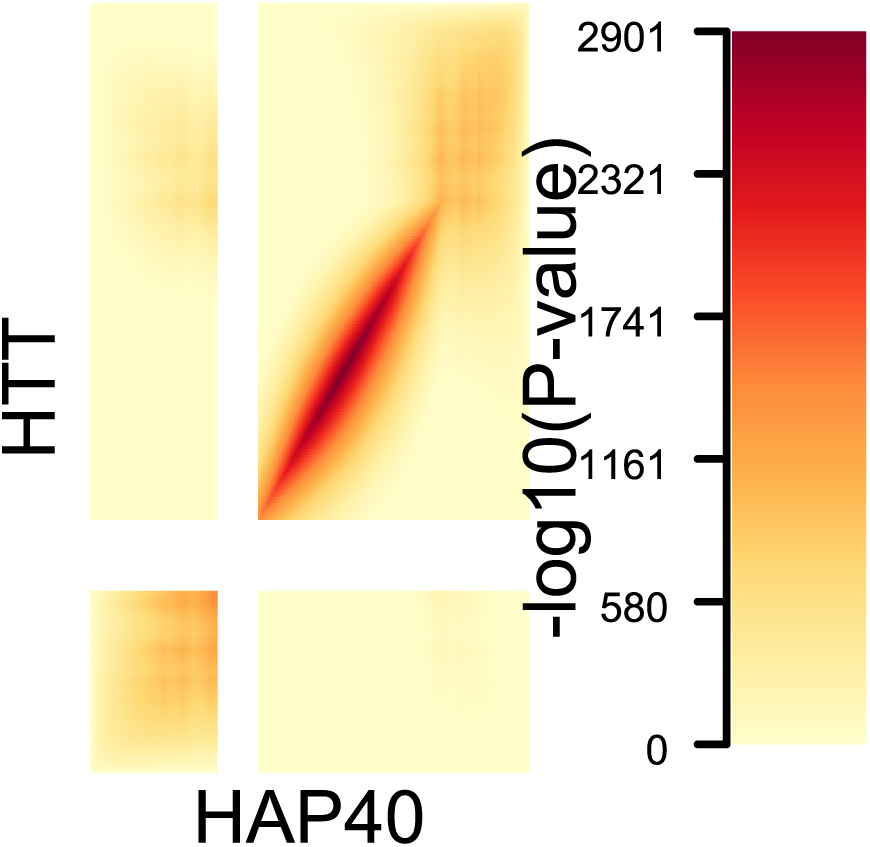
Threshold-free RRHO map of the HAP40- and HTT-induced transcriptomes. RRHO comparison of the full ranked transcriptomes (genes ranked by signed significance −log_10_(*p*)×sign(log_2_(fold change))). Colour intensity represents −log_10_(hypergeometric p-value). Signal in the bottom-left and top-right quadrants indicates concordant regulation (co-up / co-down); signal in the top-left and bottom-right quadrants indicates discordant (antagonistic) regulation. The map is dominated by the concordant co-down corner, consistent with a shared overexpression footprint; the antagonistic subcomponent is resolved at the gene level in Figure 4 B and by pathway in Supplementary Figure 5.

**Supplementary Figure 5:**
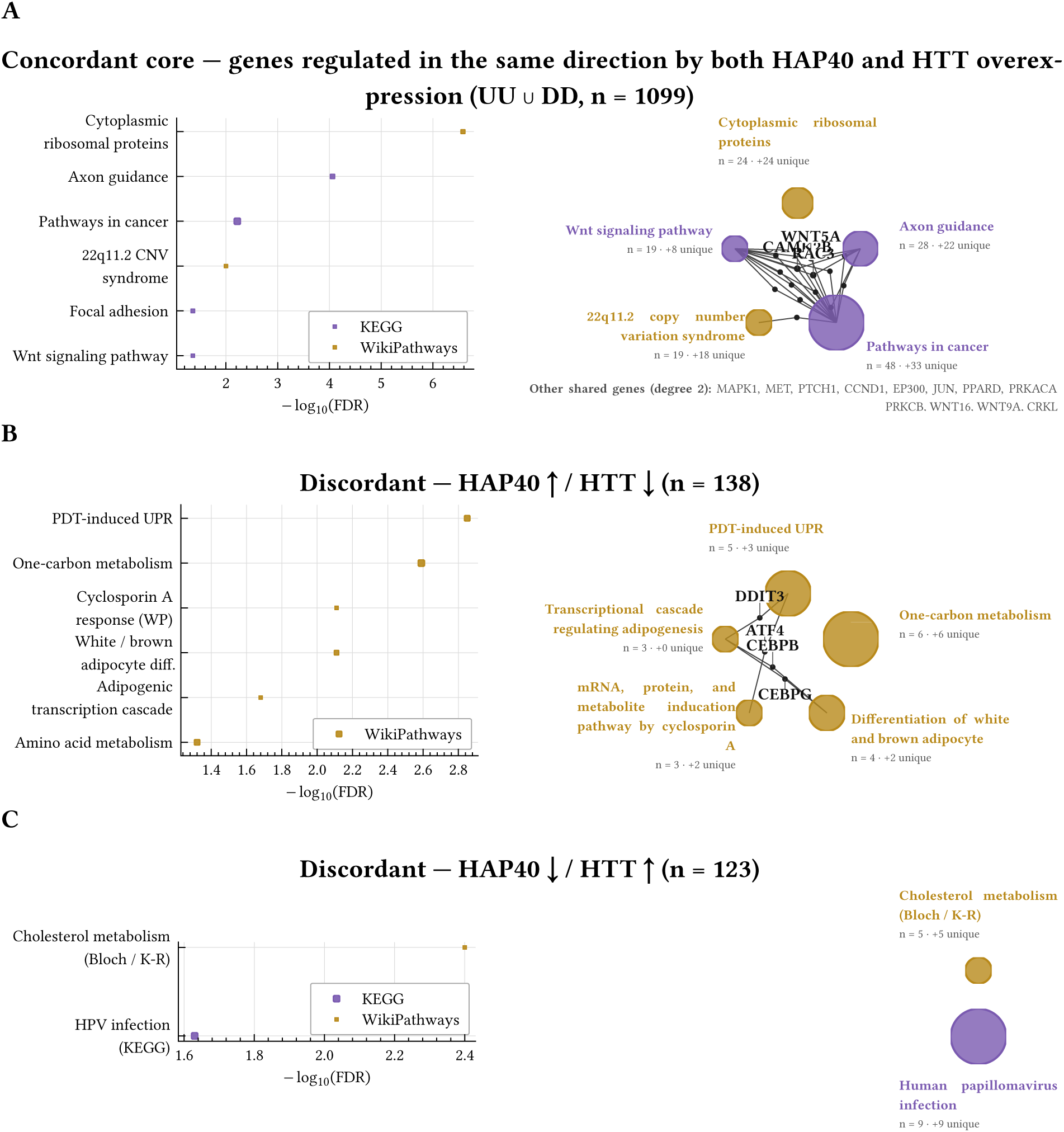
Pathway decomposition of the HAP40/HTT-overexpression RRHO. STRING functional-enrichment analysis of the three gene subsets defined by the RRHO (Supplementary Figure 4), using the top-2000 signed-rank positions per experiment. (A) Concordant core — genes regulated in the same direction by both overexpression conditions (union of up-up and down-down intersections, 1099 genes). Enrichment is dominated by generic overexpression-associated pathways (cytoplasmic ribosomal proteins, axon guidance, generic cancer/Wnt signalling), consistent with a shared proteostatic-stress footprint. (B) HAP40-dominant antagonistic arm (138 genes, upregulated by HAP40 and reciprocally downregulated by HTT). The signal is driven by the Integrated Stress Response (photodynamic-therapy-induced UPR, ATF4/DDIT3), one-carbon metabolism, and amino acid biosynthesis — the canonical ATF4/GCN2 downstream programme. (C) HTT-dominant antagonistic arm (123 genes, downregulated by HAP40 and reciprocally upregulated by HTT). The signal converges on sterol biosynthesis (Bloch/Kandutsch-Russell pathway). Dot colour encodes database; dot size encodes observed gene count. Cnetplot layout: hubs on the outer ring (size scaled by gene count; outer label = category colour), shared genes (membership ≥ 2 hubs) labelled within the ring. Rank metric: −log_10_(*p*)×sign(log_2_(FC)).

**Supplementary Figure 6:**
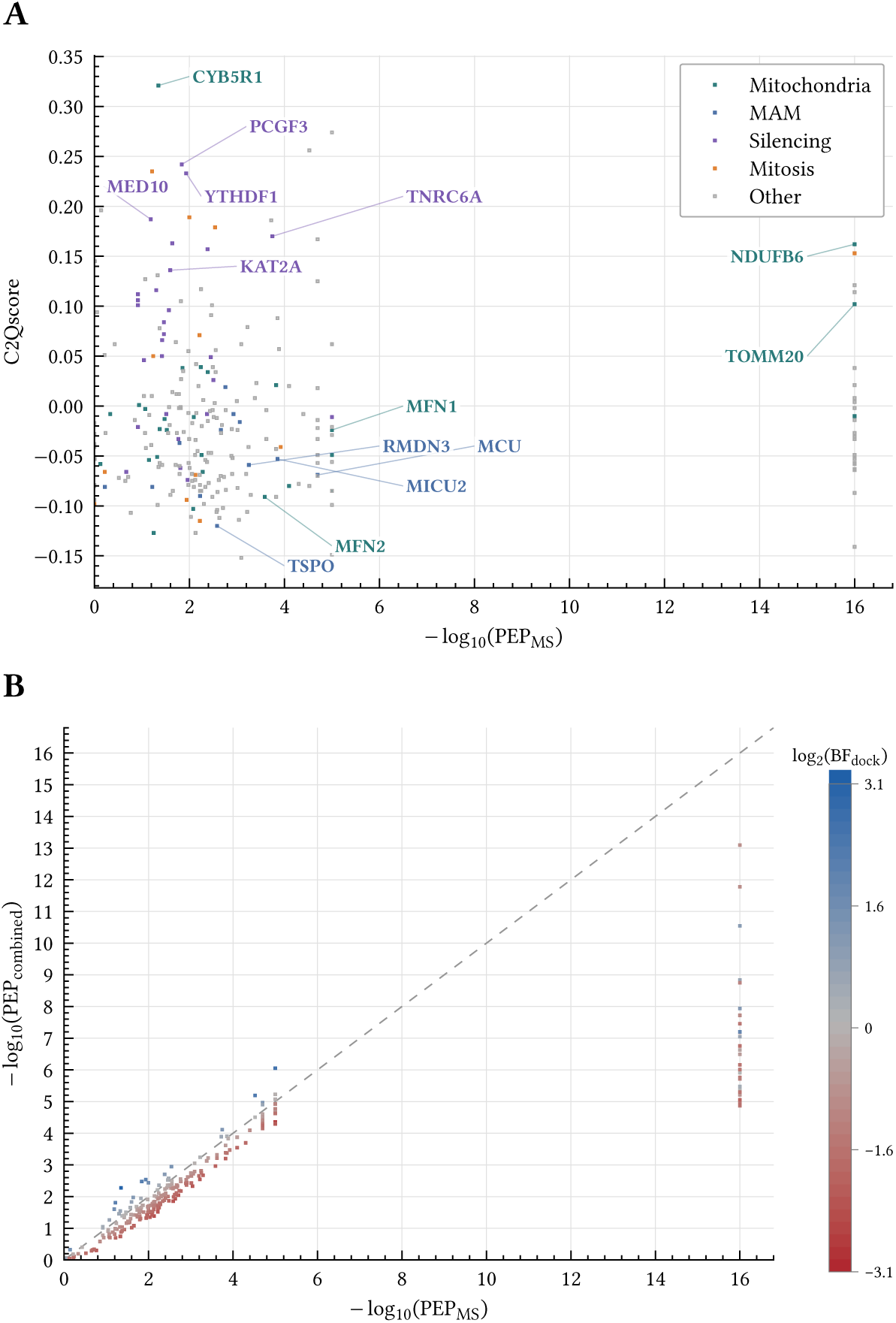
Structural plausibility of HAP40 interactions by pairwise AlphaFold3 docking. All high-confidence interactors (PEP ≤0.01) were submitted to the AlphaFold Server for pairwise bait-prey structure prediction, and each complex was scored with the adapted 4-metric C2Qscore (Genz *et al*, 2025) to derive a docking Bayes factor (BF_dock_). (A) C2Qscore versus the MS-derived −log_10_(PEP_MS_) of each interactor, coloured by functional hub. C2Qscore is a linear combination of interface pLDDT, interface PAE, pTM and ipTM calibrated on an AlphaFold3 benchmark; by construction, C2Qscore = 0 corresponds to a neutral docking Bayes factor (no update). Positive values indicate a structurally plausible pairwise interface; negative values indicate that a pairwise direct interface could not be resolved, consistent with association through higher-order assemblies, membrane context, or transient interfaces that AlphaFold3 is known to under-predict. (B) Posterior update: −log_10_(PEP_MS_) versus −log_10_(PEP_combined_) after the two-stage Bayesian update, with each point coloured by log_2_(BF_dock_). Points above the diagonal were boosted by docking evidence; points below were reduced. The MS posterior is effectively preserved for hits with PEP_MS_≪0.01, so the docking update chiefly reassigns confidence among borderline and unstructured candidates. Highlighted proteins (TNRC6A, MFN1, MFN2, RMDN3, MCU, MICU2, TSPO, TOMM20, CYB5R1, NDUFB6, PCGF3, KAT2A, MED10, YTHDF1) are discussed in the main text. Full per-pair structural metrics are available in the BayesInteractomics HTML report (Supplementary Table 4).

**Supplementary Figure 7:**
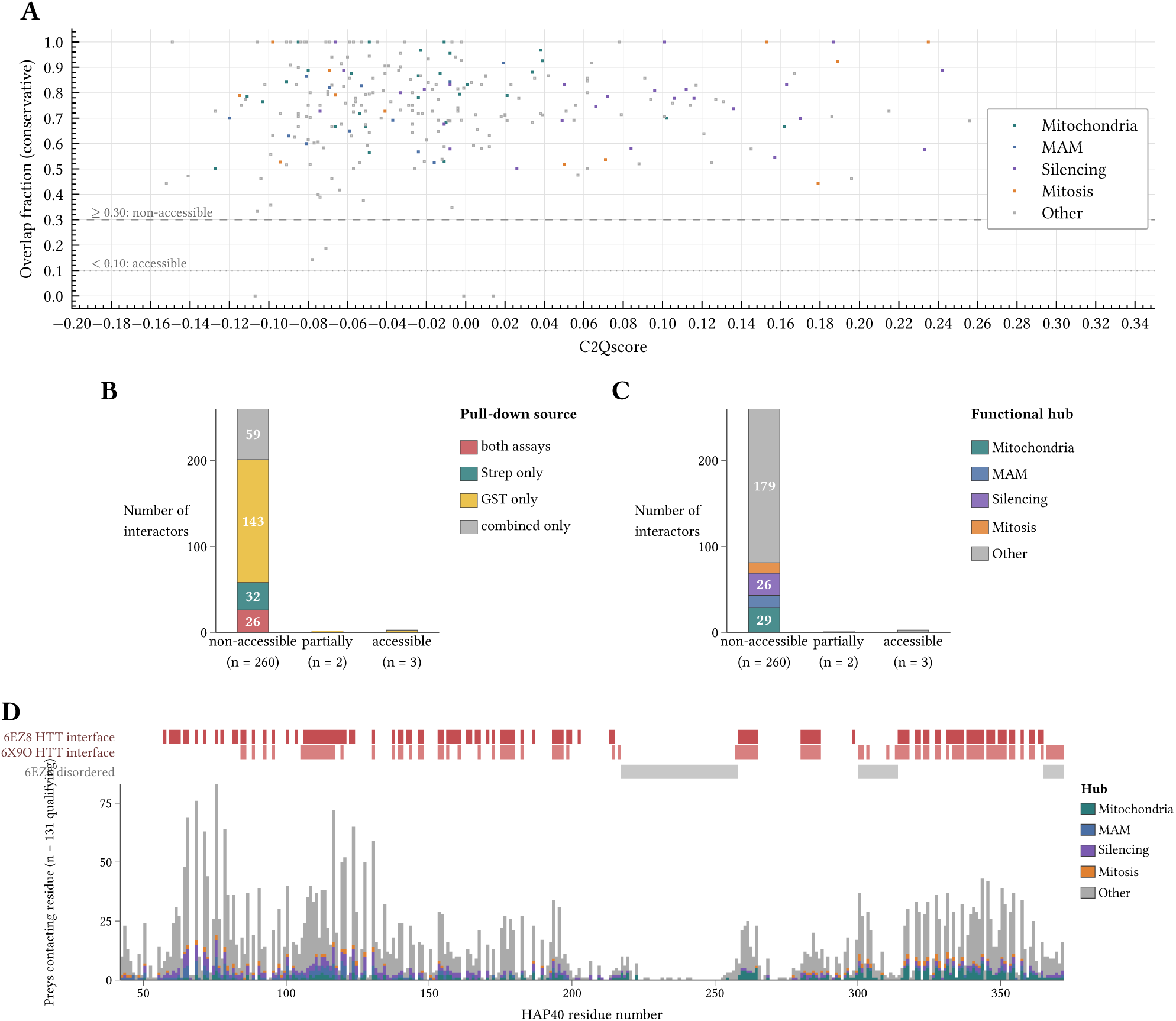
In silico comparison of AlphaFold3-predicted HAP40–prey interfaces with the experimental HTT–HAP40 interface. AF3 predictions were superposed onto the wild-type (Q17) cryo-EM HTT–HAP40 complex (PDB 6EZ8 (Guo *et al*, 2018)) and HAP40 residues within 5 Å of the prey were compared with HAP40 residues within 5 Å of HTT. The overlap fraction is computed on the 6EZ8-resolved portion of HAP40 only (residues 42–364), so disordered regions do not inflate the “accessible” category. Category cutoffs: non-accessible ≥0.30, partially accessible 0.10–0.30, accessible <0.10. Extracting the HTT interface independently from three alternative HTT–HAP40 cryo-EM structures gave concordant interface definitions (Jaccard/Cohen’s *κ* against 6EZ8): higher-resolution wild-type 6X9O (Harding *et al*, 2021) 0.7807/0.7788 Q46 expanded 7DXJ (Huang *et al*, 2021b) 0.9048/0.9095 Q128 expanded 7DXK (Huang *et al*, 2021b) 0.8333/0.8343. The HTT footprint on HAP40 is thus essentially mutation-independent, so the category assignments derived from 6EZ8 transfer to the expanded-repeat setting. (A) Conservative overlap fraction versus C2Qscore, coloured by functional hub. (B) Each category split by pull-down source. (C) Each category split by functional hub. (D) Per-residue view along HAP40 of the HTT interface in 6EZ8 and 6X9O (red strips), 6EZ8 disordered stretches (grey strip), and the stacked frequency of AF3 prey contacts per residue across 131 qualifying preys (PEP_combined_≤0.01), coloured by functional hub.

**Supplementary Figure 8:**
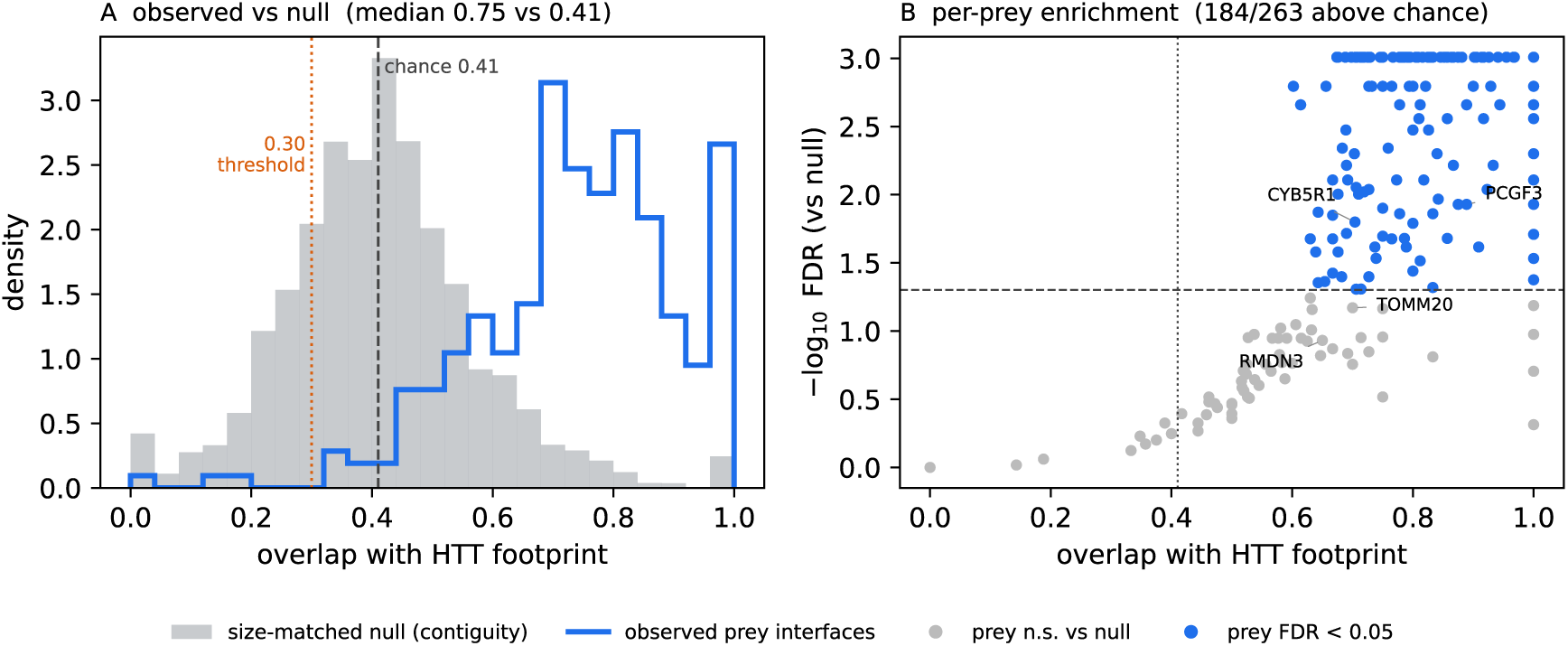
The HAP40–prey interfaces overlap the HTT footprint more than chance. Because the HTT footprint covers 46 % of the resolved HAP40 surface (124 of 268 residues) and is contiguous, the binary “non-accessible” classification (overlap ≥0.30; Supplementary Figure 7) is expected to be met by most randomly placed interfaces, so each AlphaFold3 prediction was compared against a size-matched null of random connected surface patches grown on the HAP40 C*α* contact graph (residues within 8 Å). (A) Distribution of the observed prey-interface overlap with the HTT footprint (blue) versus the size-matched contiguity null (grey). Dashed line, mean null overlap (0.41); dotted line, the 0.30 classification threshold. The observed overlap is shifted well above chance (median 0.75). (B) Per-prey enrichment: observed overlap versus the Benjamini-Hochberg–corrected significance of exceeding the null; 184 of 263 evaluable predictions exceed chance at FDR < 0.05 (blue). Dashed line, FDR = 0.05 dotted line, mean null overlap. A simpler uniform null (expected overlap 0.46) gives concordant results (189 of 263 preys above chance). Representative interactors are labelled; a minority (e.g. RMDN3, TOMM20) show overlap that is not individually distinguishable from chance, so the surface argument is made at the population level.

**Supplementary Figure 9:**
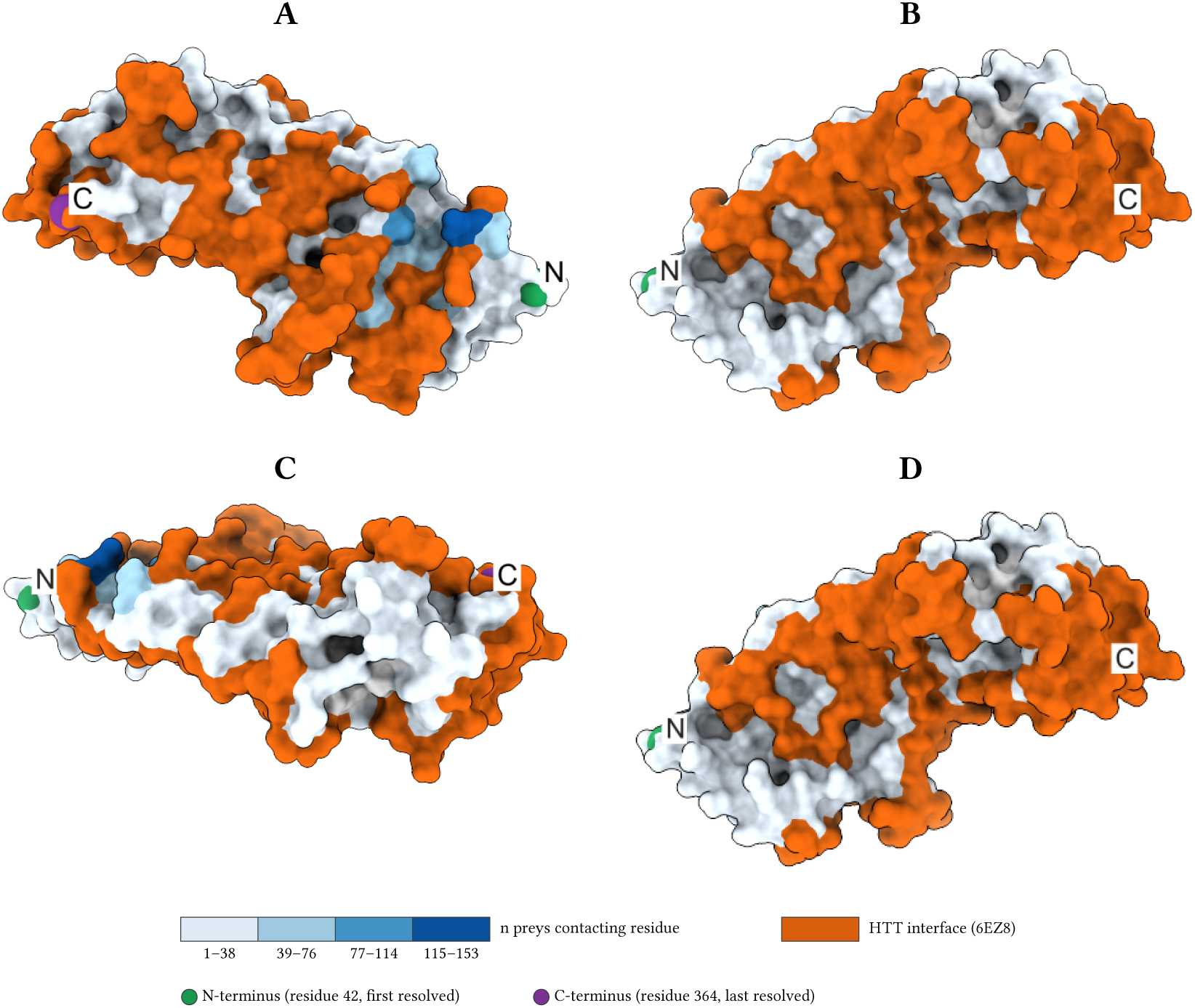
HAP40 molecular surface coloured by HTT interface and AF3 prey-contact frequency. HAP40 surface from the wild-type HTT–HAP40 complex (PDB 6EZ8 (Guo *et al*, 2018)); HTT is hidden to expose HAP40. Surface residues within 5 Å of HTT in 6EZ8 are coloured orange. All other residues are coloured by the number of AlphaFold3 HAP40–prey predictions in which they fall within 5 Å of a prey, binned into quartiles of a sequential blue palette. The N- and C-termini of the resolved construct (residues 42 and 364) are marked with green and purple spheres. Disordered residues (N-terminus 1–41, central loop) are absent from the surface because 6EZ8 does not resolve them. (A) Front view (HTT-binding face toward the viewer). (B) 180° rotation about the vertical axis. (C) 90° pitch up (top). (D) 90° pitch down (bottom). The near-complete orange coverage of the HAP40 surface explains why the vast majority of AF3 HAP40–prey predictions are classified as non-accessible in Supplementary Figure 7.

**Supplementary Figure 10:**
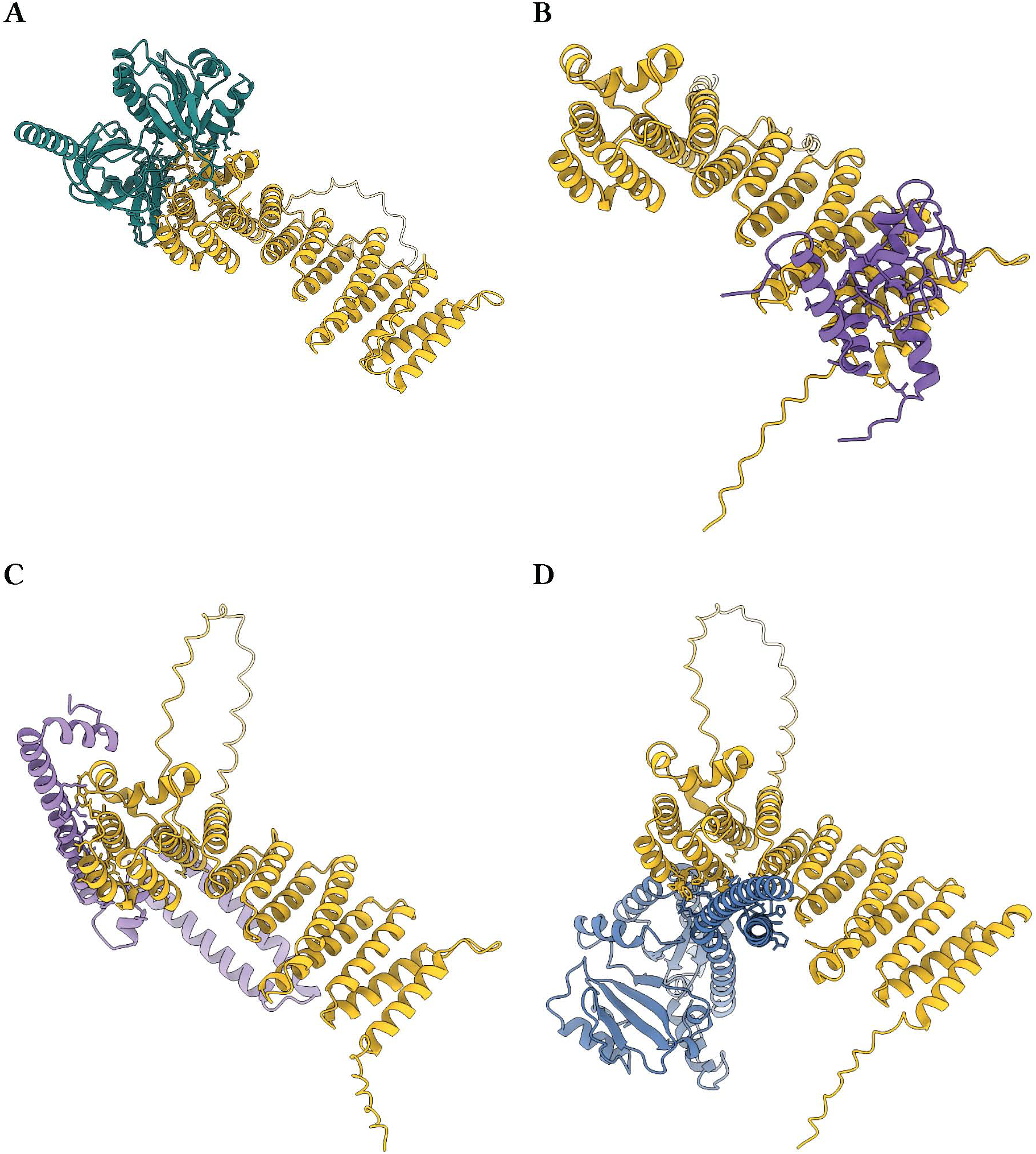
Representative AlphaFold3 HAP40–prey predictions. Best-ranked AF3 models for four representative preys, superposed on a common HAP40 orientation derived from the cryo-EM HTT–HAP40 complex (PDB 6EZ8). HAP40 is shown in gold; prey in the hub colour from Supplementary Figure 6. Interface residues within 5 Å of the partner chain are drawn as sticks. (A) CYB5R1 (C2Qscore 0.32, BF_dock_ 8.80, ipTM 0.59). (B) PCGF3 (C2Qscore 0.24, BF_dock_ 4.41, ipTM 0.47). (C) MED10 (C2Qscore 0.19, BF_dock_ 2.72, ipTM 0.44). (D) MFN2 (C2Qscore −0.09, BF_dock_ 0.24, ipTM 0.16), shown as a contrast example where AlphaFold3 does not resolve a confident pairwise interface, consistent with engagement through the obligate MFN homodimeric assembly.

**Supplementary Table 1:**
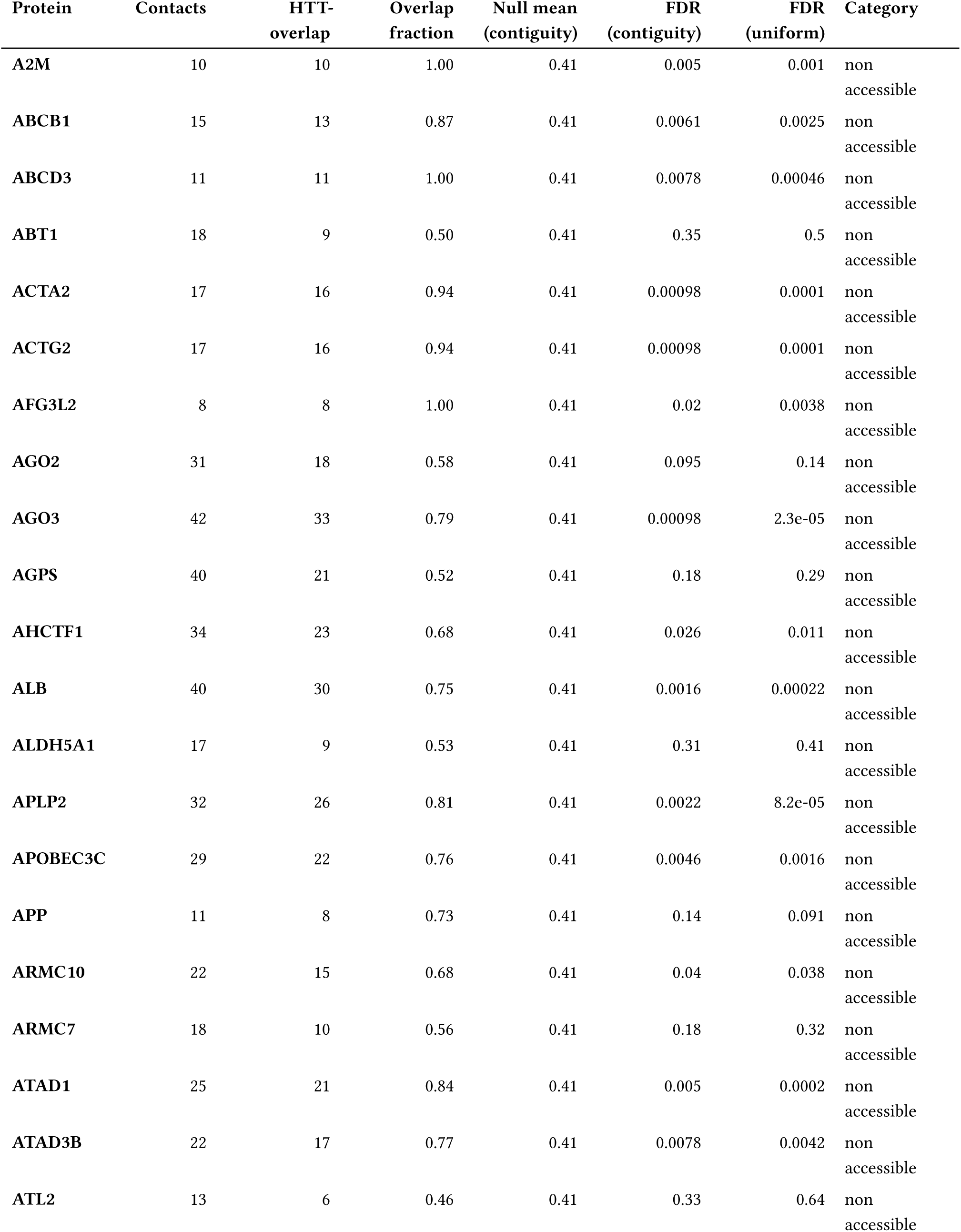
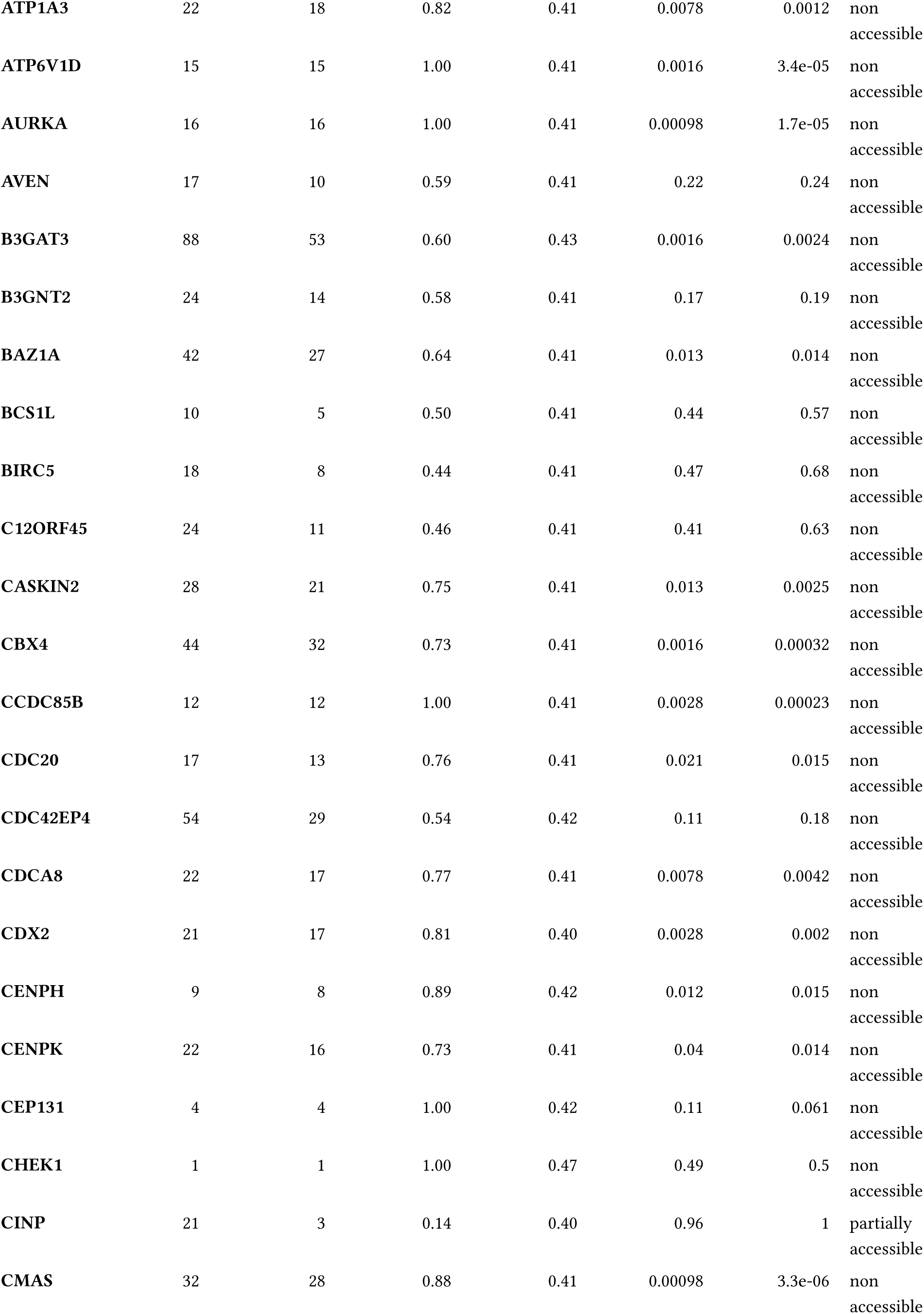
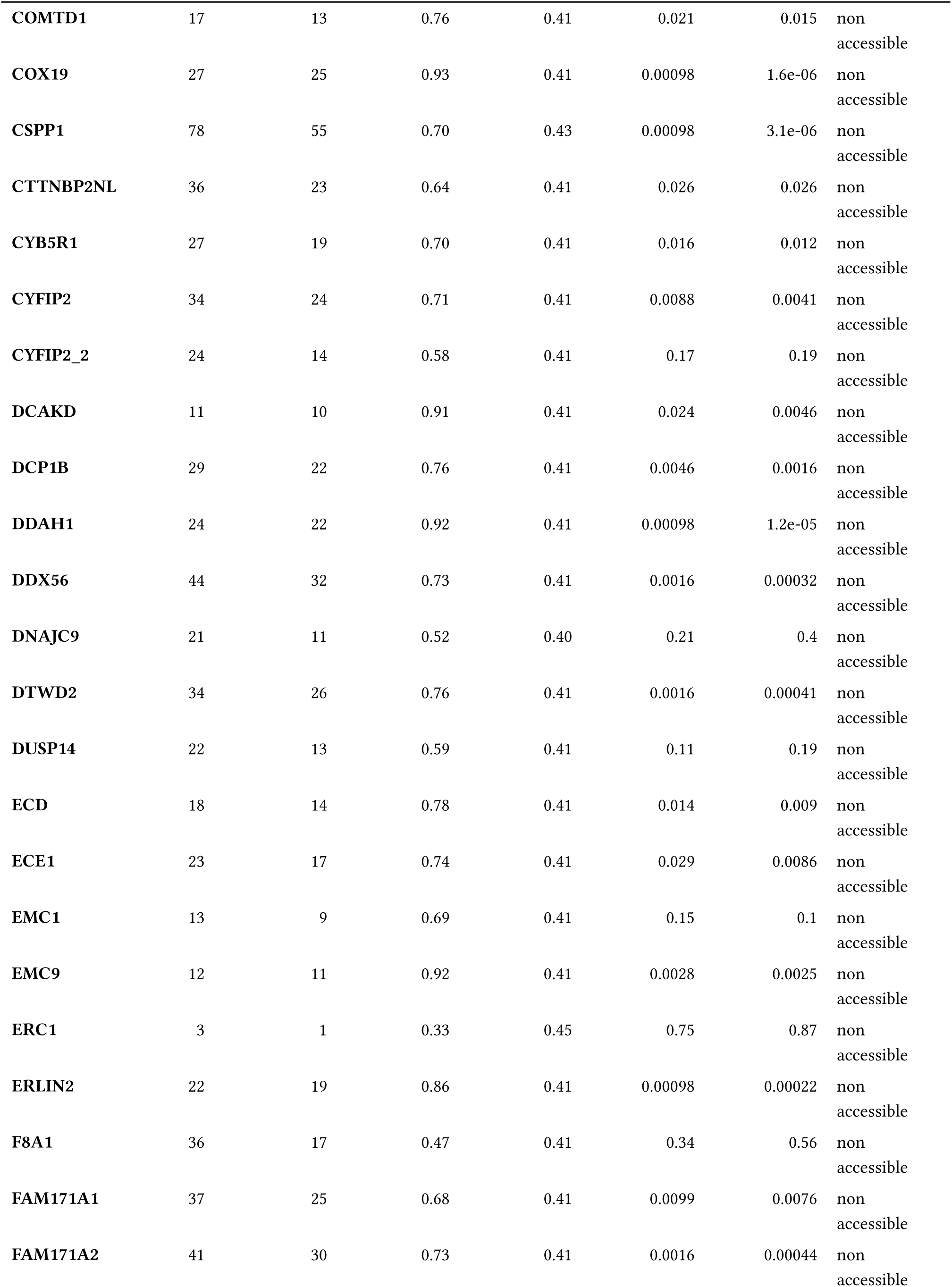
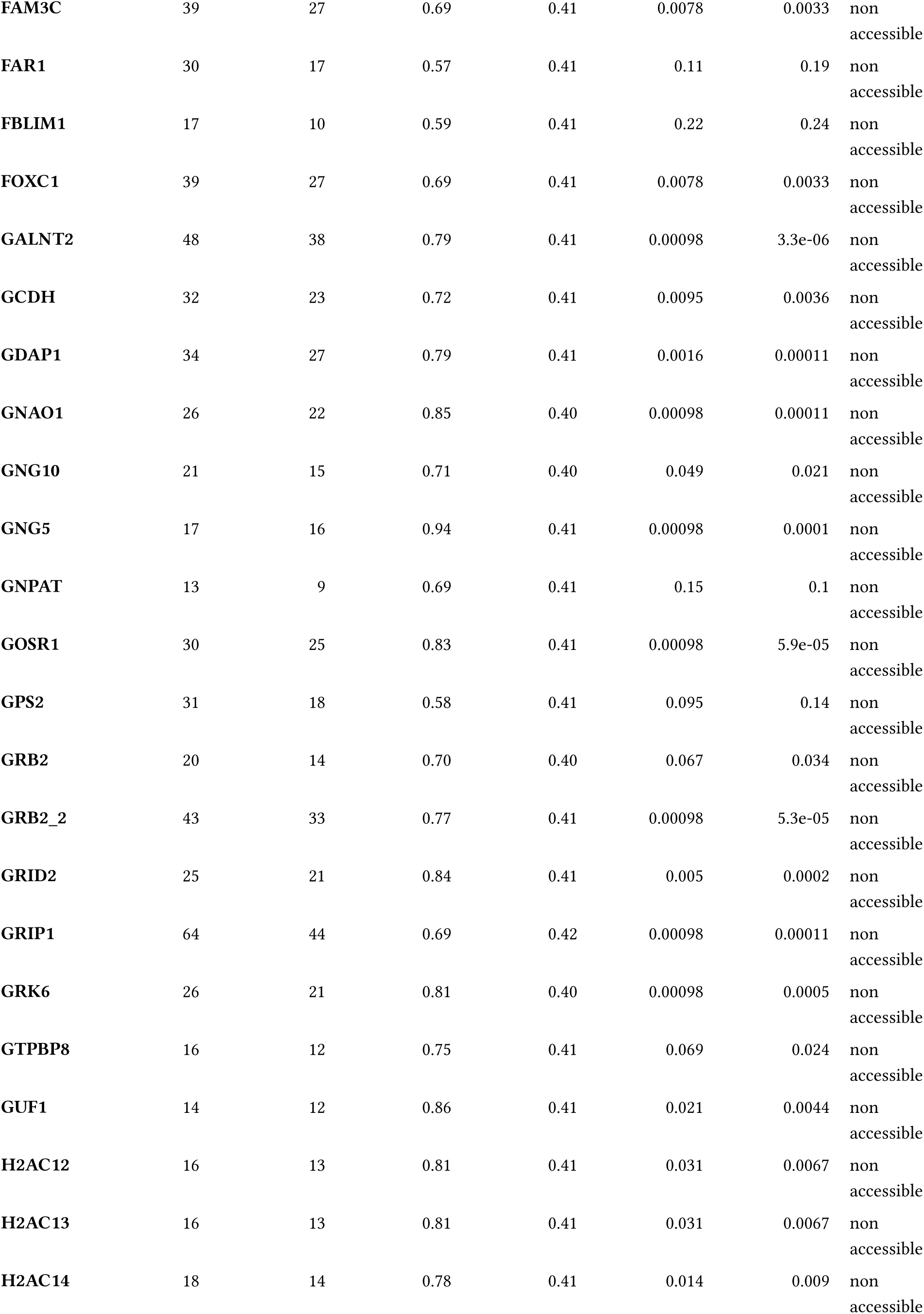
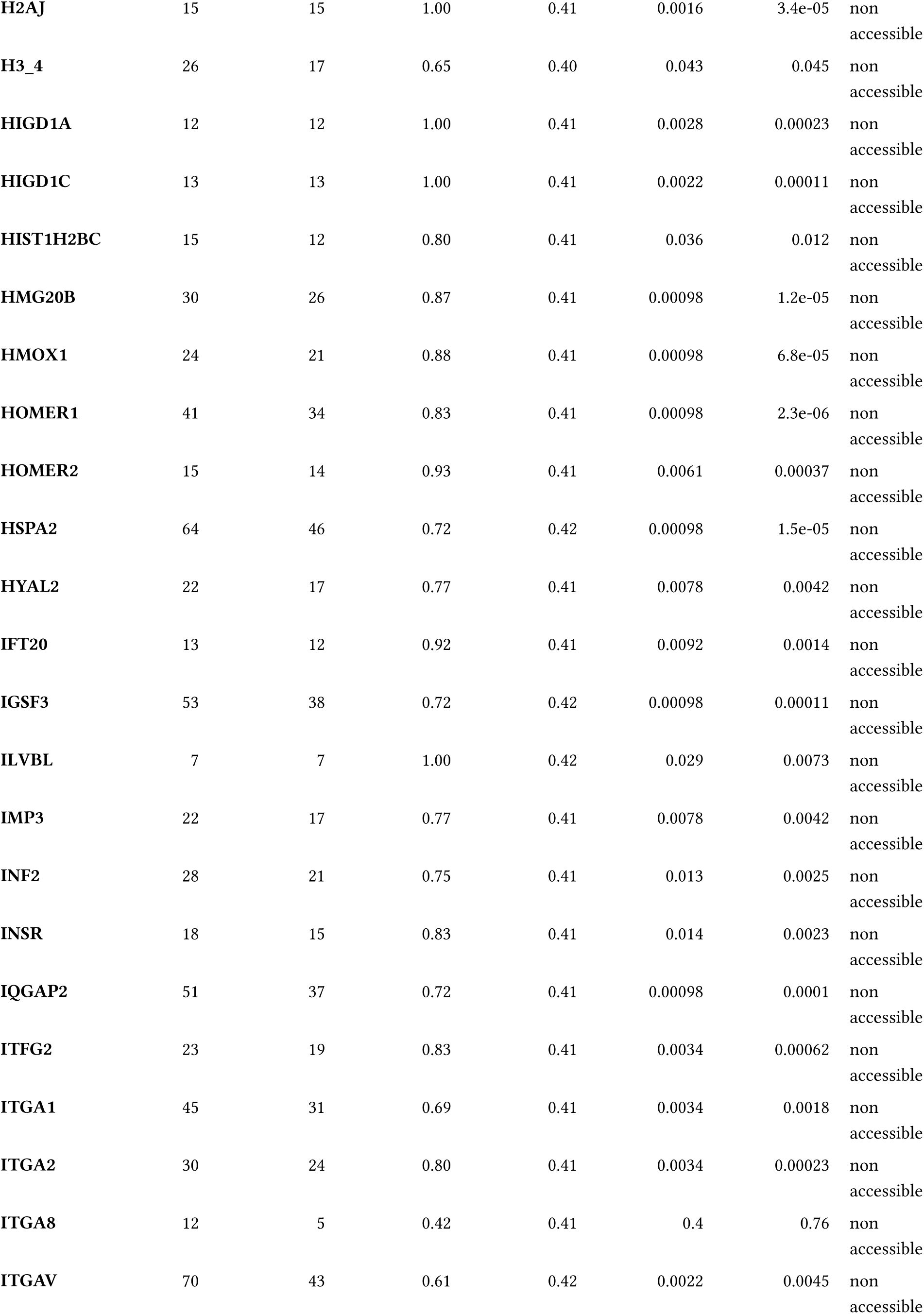
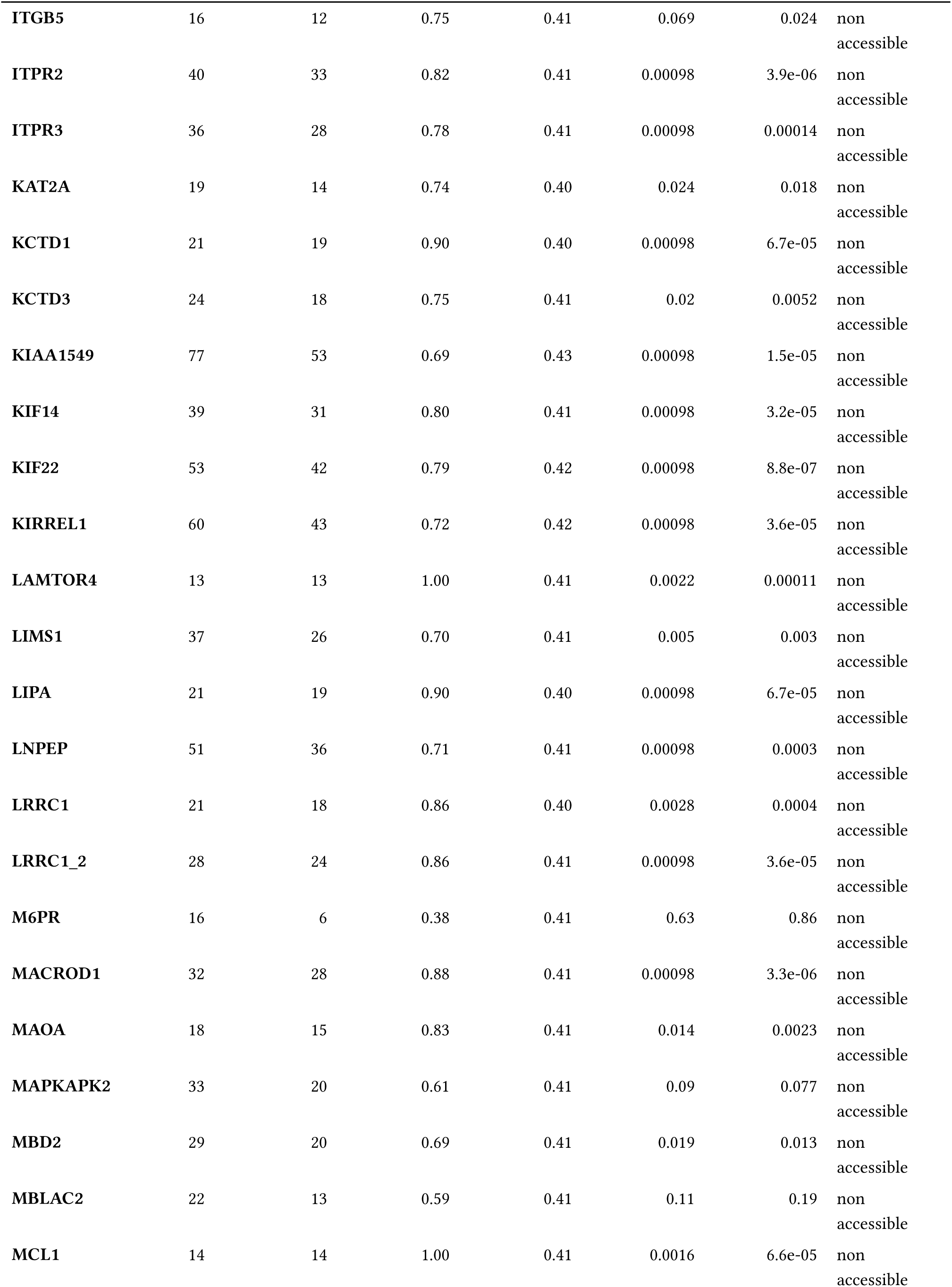
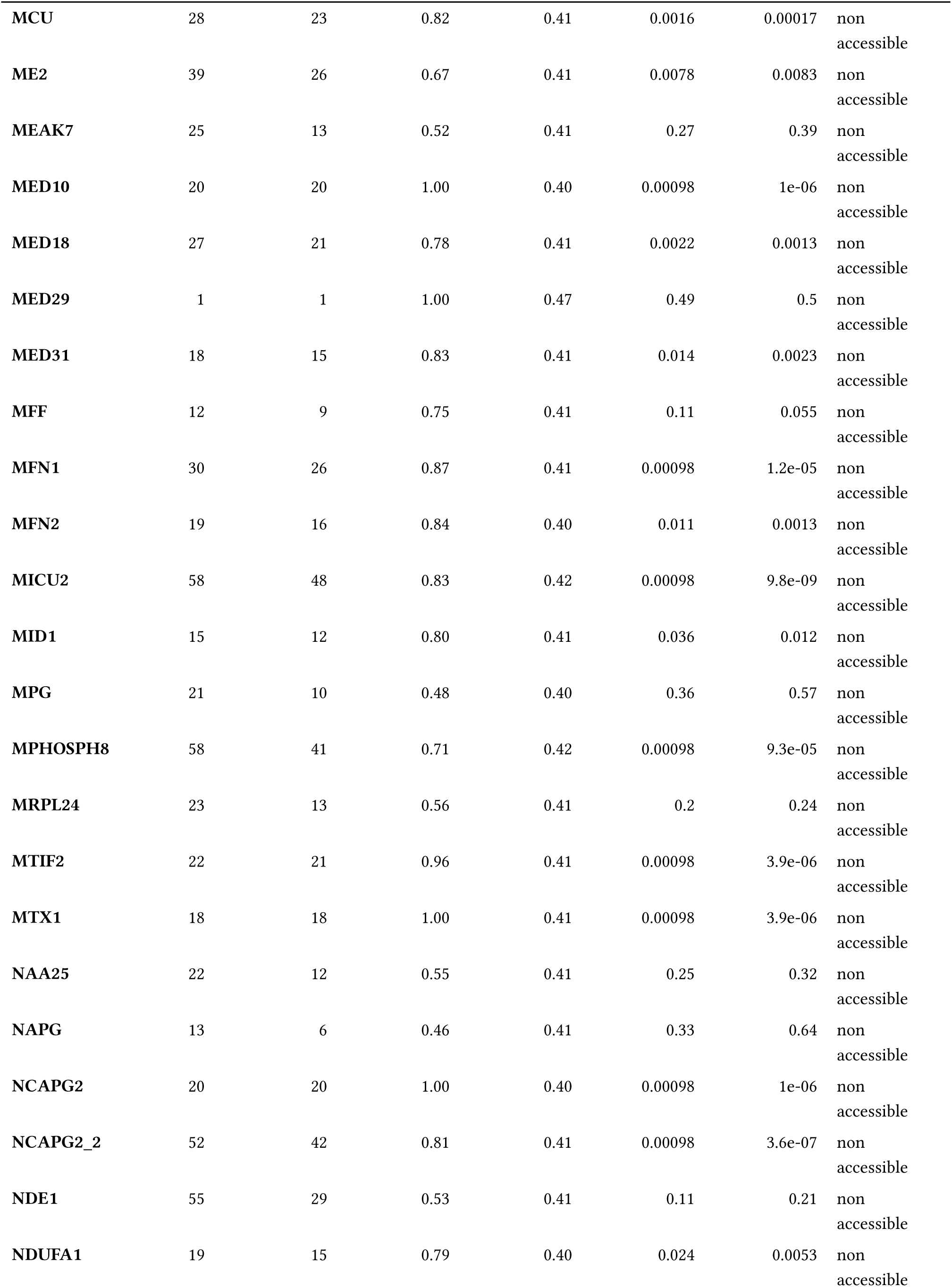
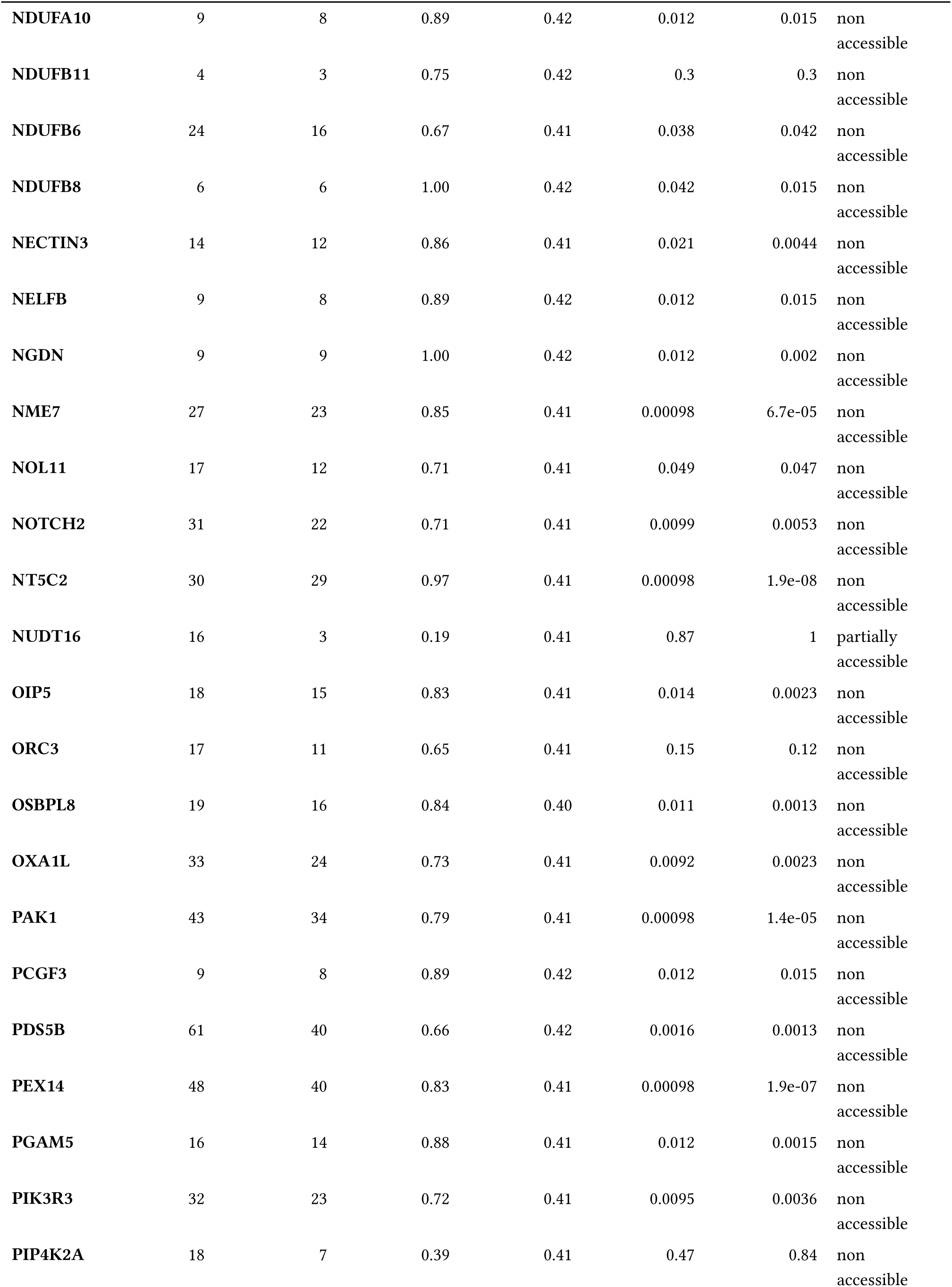
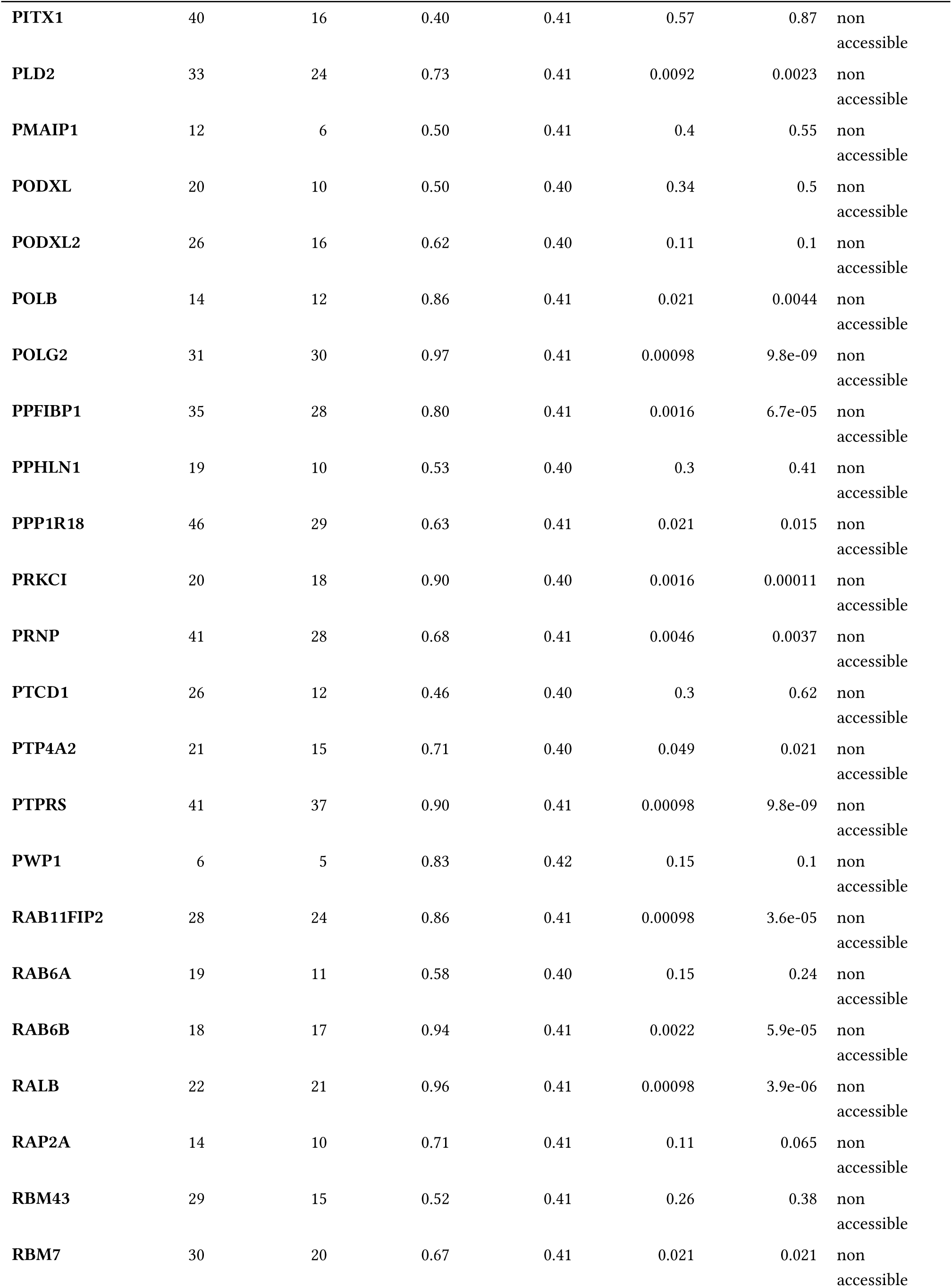
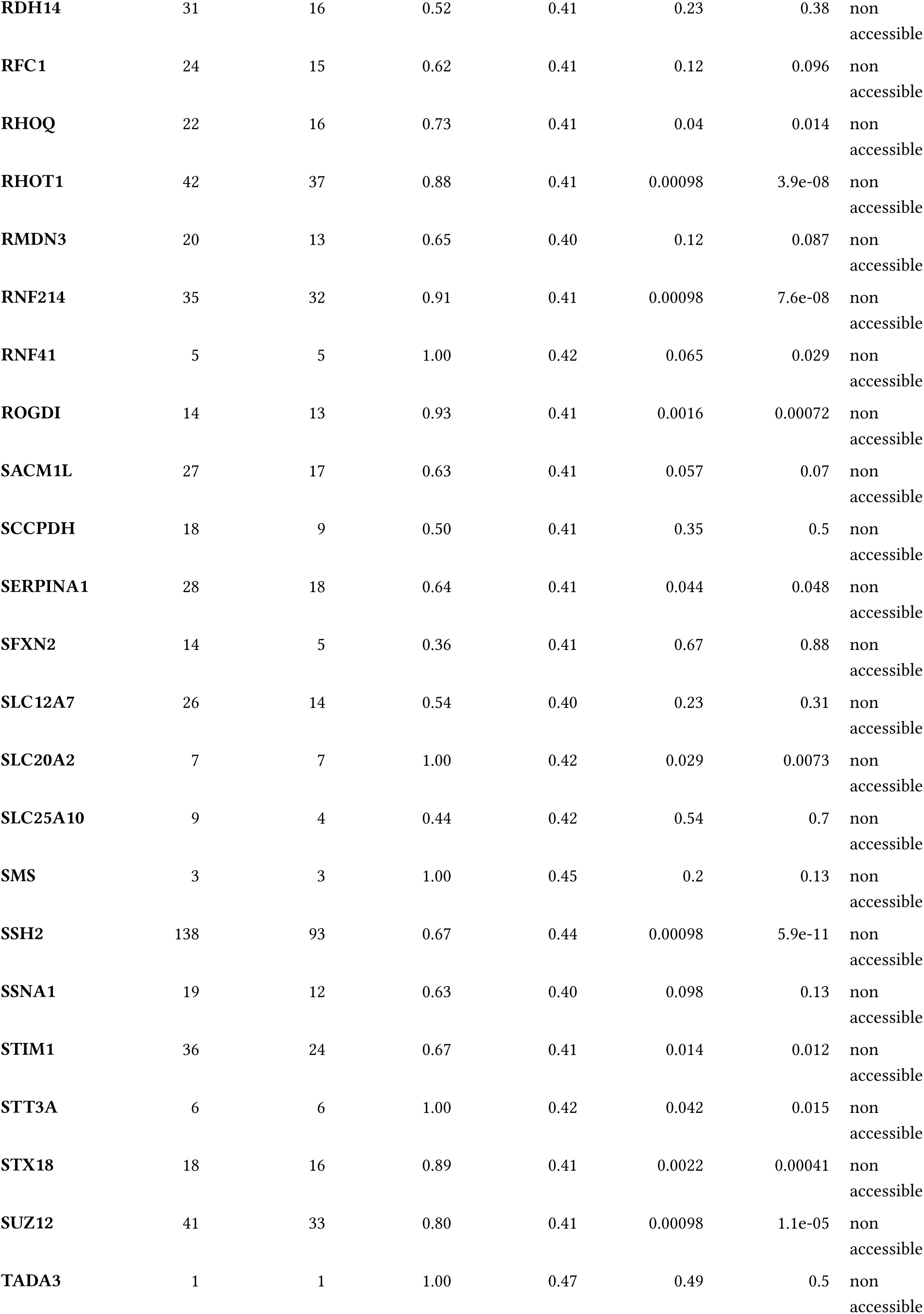
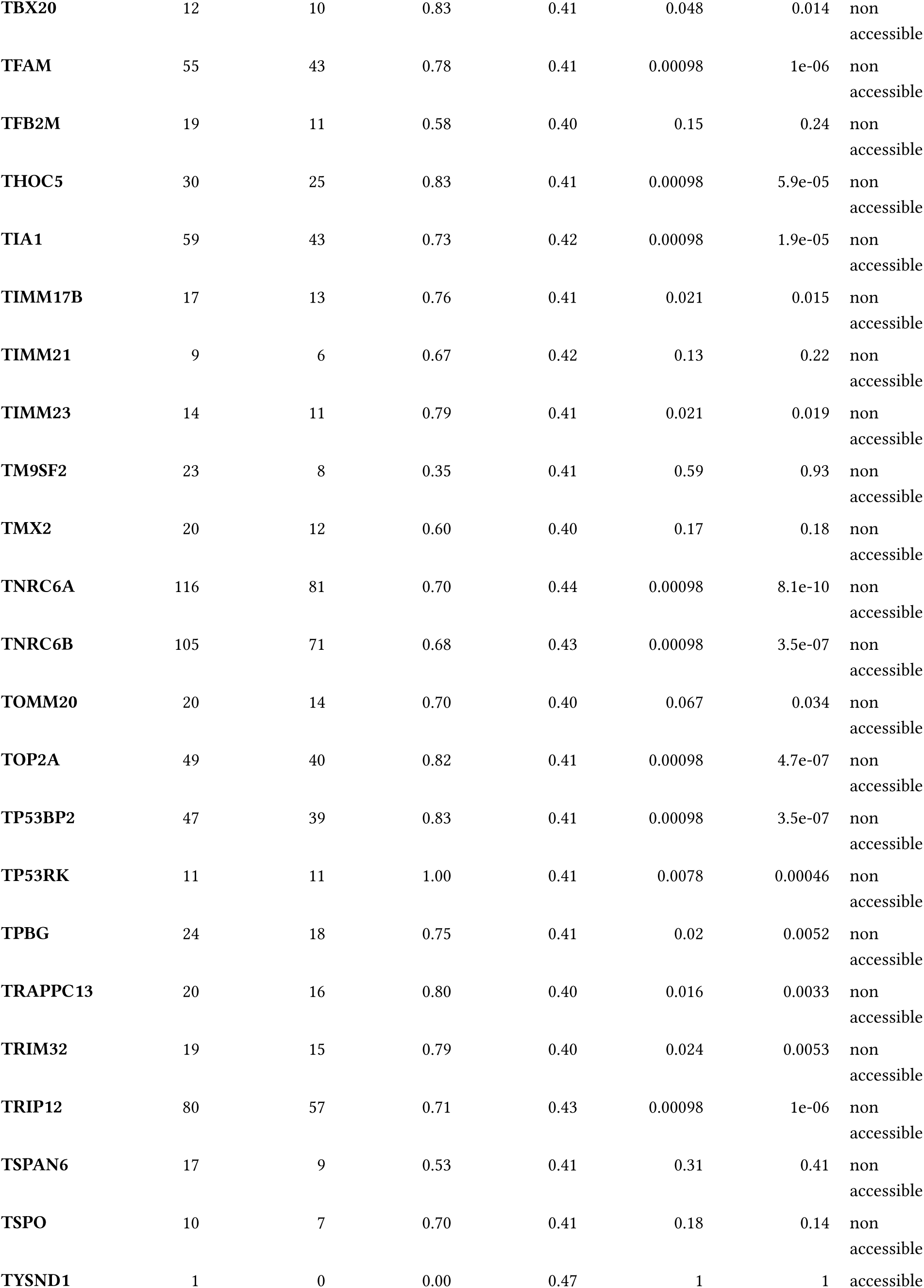
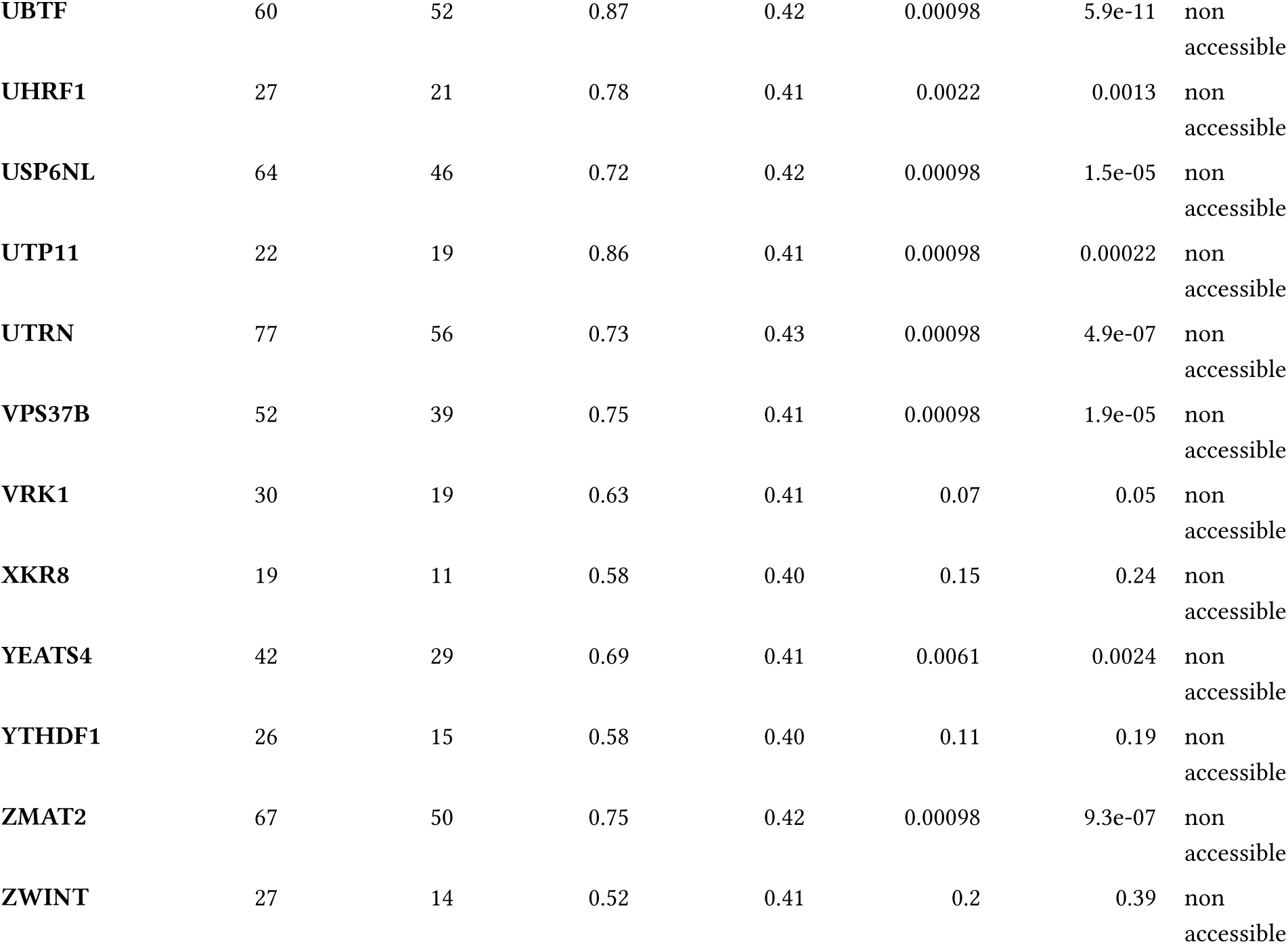
Per-prey surface-overlap null-model results. For every AlphaFold3 HAP40–prey prediction with contacts in the 6EZ8-resolved envelope (263 preys): the number of HAP40 contact residues, the number overlapping the HTT interface, the observed overlap fraction, the mean overlap expected under the size-matched contiguity null, and the Benjamini-Hochberg–corrected significance of exceeding the contiguity and uniform nulls. Categories follow Supplementary Figure 7 (non-accessible ≥0.30; partially accessible 0.10–0.30; accessible <0.10); see Supplementary Figure 8.

**Supplementary Table 2: *In vivo* HAP40 interactome.**

Interactive report from Bayesian analysis of the Strep-HAP40 affinity purification-mass spectrometry data.

Interactive BayesInteractomics report provided as a separate Supplementary Data file.

**Supplementary Table 3: *In vitro* HAP40 interactome.**

Interactive report from Bayesian analysis of the GST-HAP40 affinity purification-mass spectrometry data.

Interactive BayesInteractomics report provided as a separate Supplementary Data file.

**Supplementary Table 4: Combined HAP40 interactome.**

Interactive report from Bayesian analysis of the combined *in vitro* and *in vivo* HAP40 interaction assays.

Interactive BayesInteractomics report provided as a separate Supplementary Data file.

**Supplementary Table 5:** Changes in mRNA levels of genes involved in the Golgi stress response.

|  |  | HAP40 overexpression |  | HTT overexpression |  |
| --- | --- | --- | --- | --- | --- |
|  |  | log<sub>2</sub>(fold change) | FDR | log<sub>2</sub>(fold change) | FDR |
|  | **TFE3 pathway** |  |  |  |  |
|  | **GCP60 / ACBD3** | 0.233 | 0.595 | −0.052 | 0.833 |
|  | **GM130 / GOLGA2** | −0.207 | 0.677 | −0.529 | 0.002 |
|  | **Giantin / GOLGB1** | 0.284 | 0.500 | −0.110 | 0.745 |
|  | **SIAT4A / ST3GAL1** | 0.243 | 0.676 | −1.445 | 1.74·10<sup>−17</sup> |
|  | **SIAT10 / ST3GAL6** | 0.194 | 0.759 | −0.066 | 0.867 |
|  | **FUT1** | 1.461 | 2.74·10<sup>−08</sup> | −1.734 | 5.49·10<sup>−14</sup> |
|  | **B3GAT2** | 0.853 | 0.309 | 0.79 | 0.090 |
|  | **UAP1L1** | 0.903 | 5.51·10<sup>−4</sup> | −0.020 | 0.969 |
|  | **STX3A / STX3** | 0.496 | 0.082 | 0.229 | 0.437 |
|  | **WIPI49 / WIPI1** | 0.486 | 0.224 | −0.721 | 1.77·10<sup>−3</sup> |
|  | **RAB20** | 0.157 | 0.842 | 0.067 | 0.920 |
|  | **PCSK1** | −0.667 | 0.686 | −1.160 | 0.222 |
|  | **CREB3 pathway** |  |  |  |  |
|  | **ARF4** | −0.137 | 0.810 | −0.326 | 0.112 |
|  | **DR4 / TNFRSF10A** | 0.105 | 0.870 | 0.500 | 0.012 |
|  | **TRAPPC13** | 0.249 | 0.534 | 0.199 | 0.374 |
|  | **HSP47 pathway** |  |  |  |  |
|  | **HSP47 / SERPINH1** | −0.102 | 0.864 | −0.490 | 0.020 |

**Supplementary Table 6:**
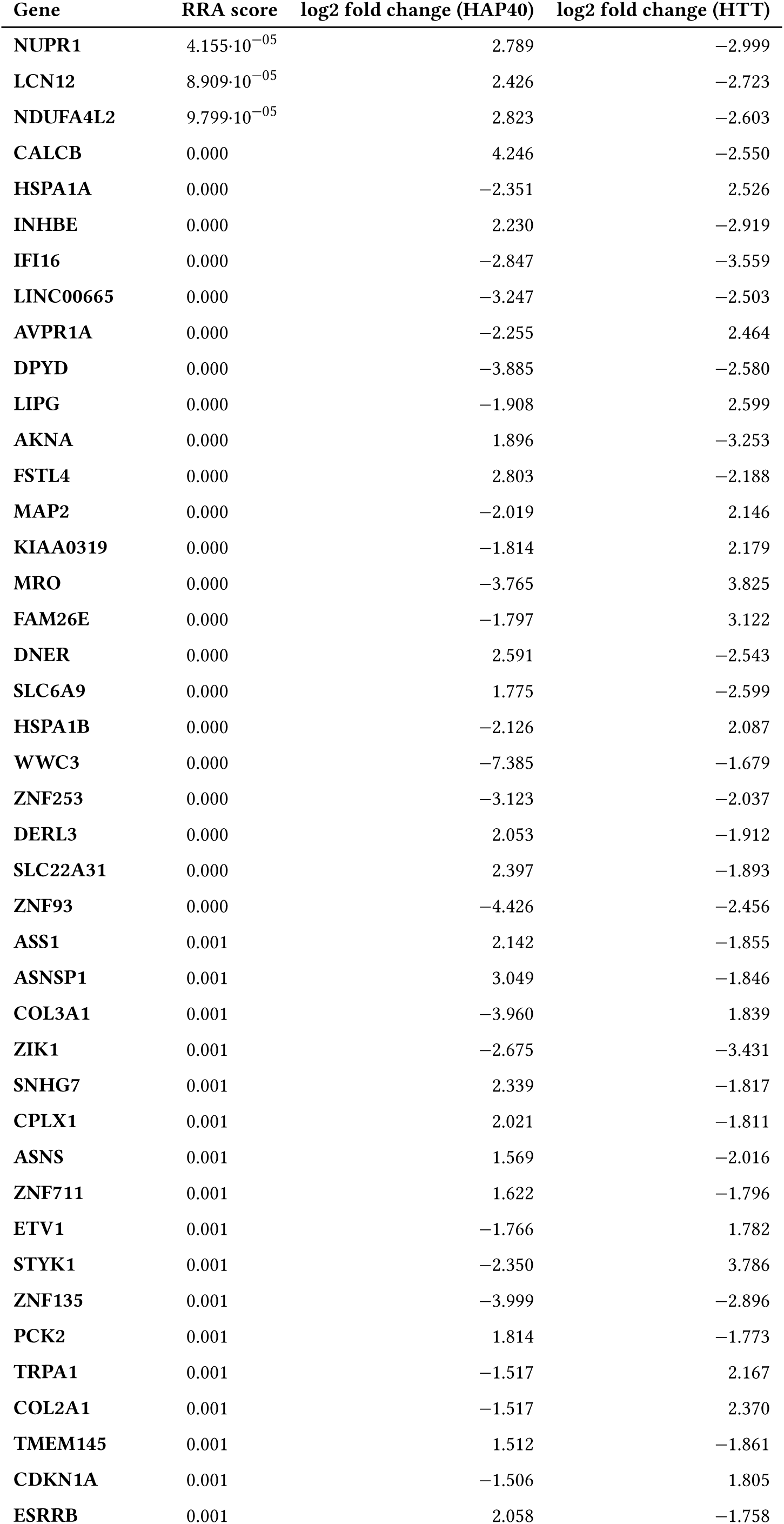
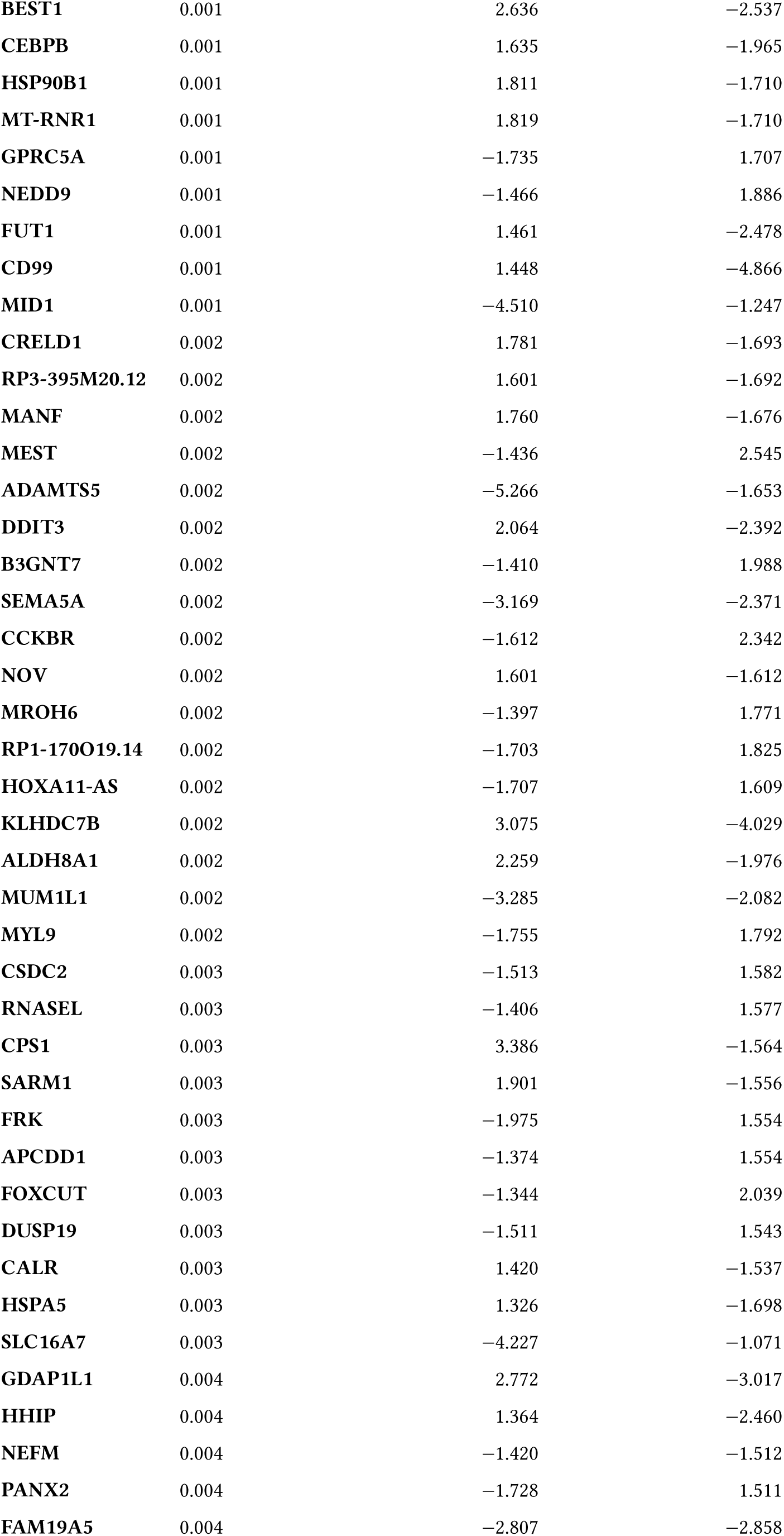
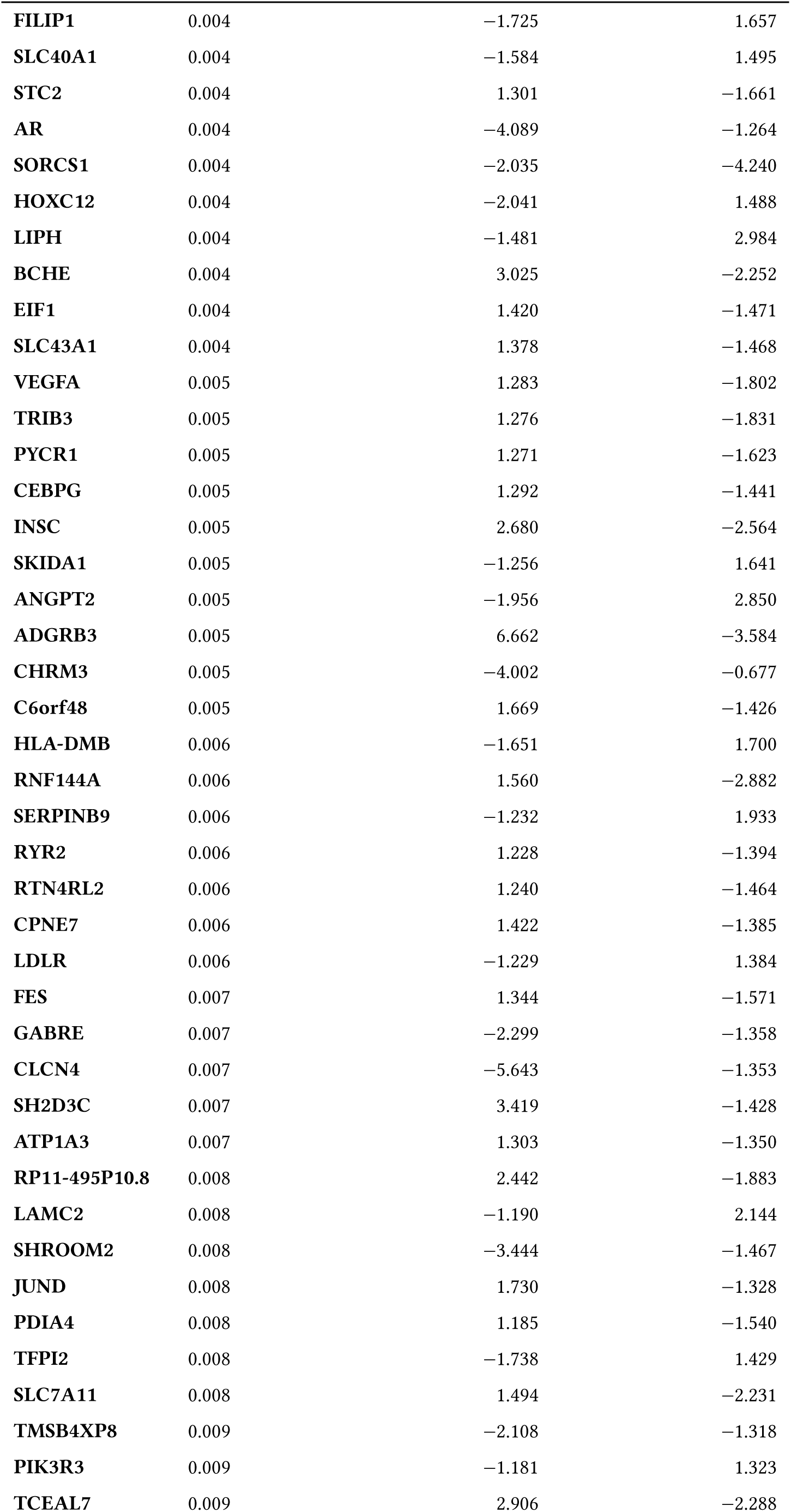
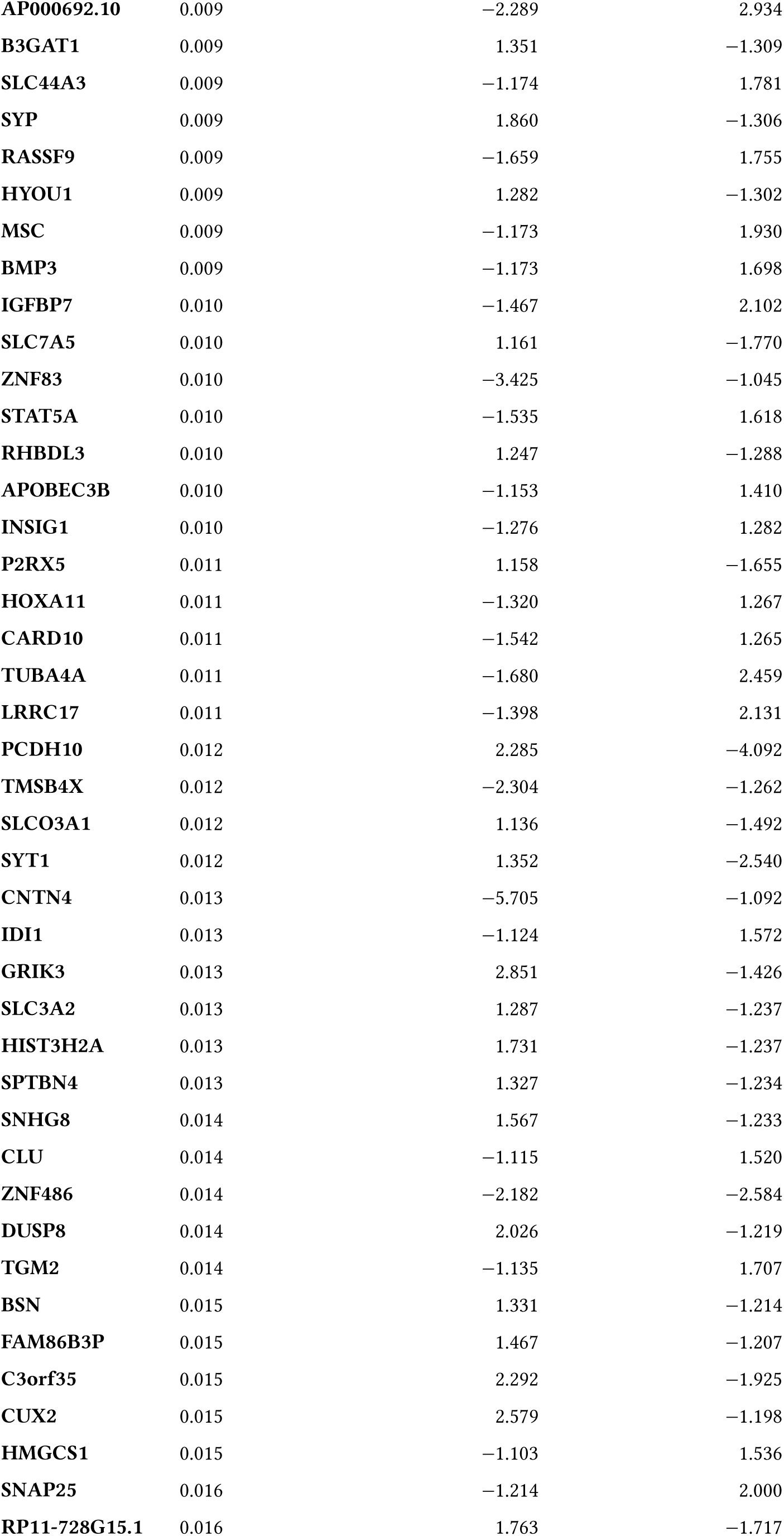
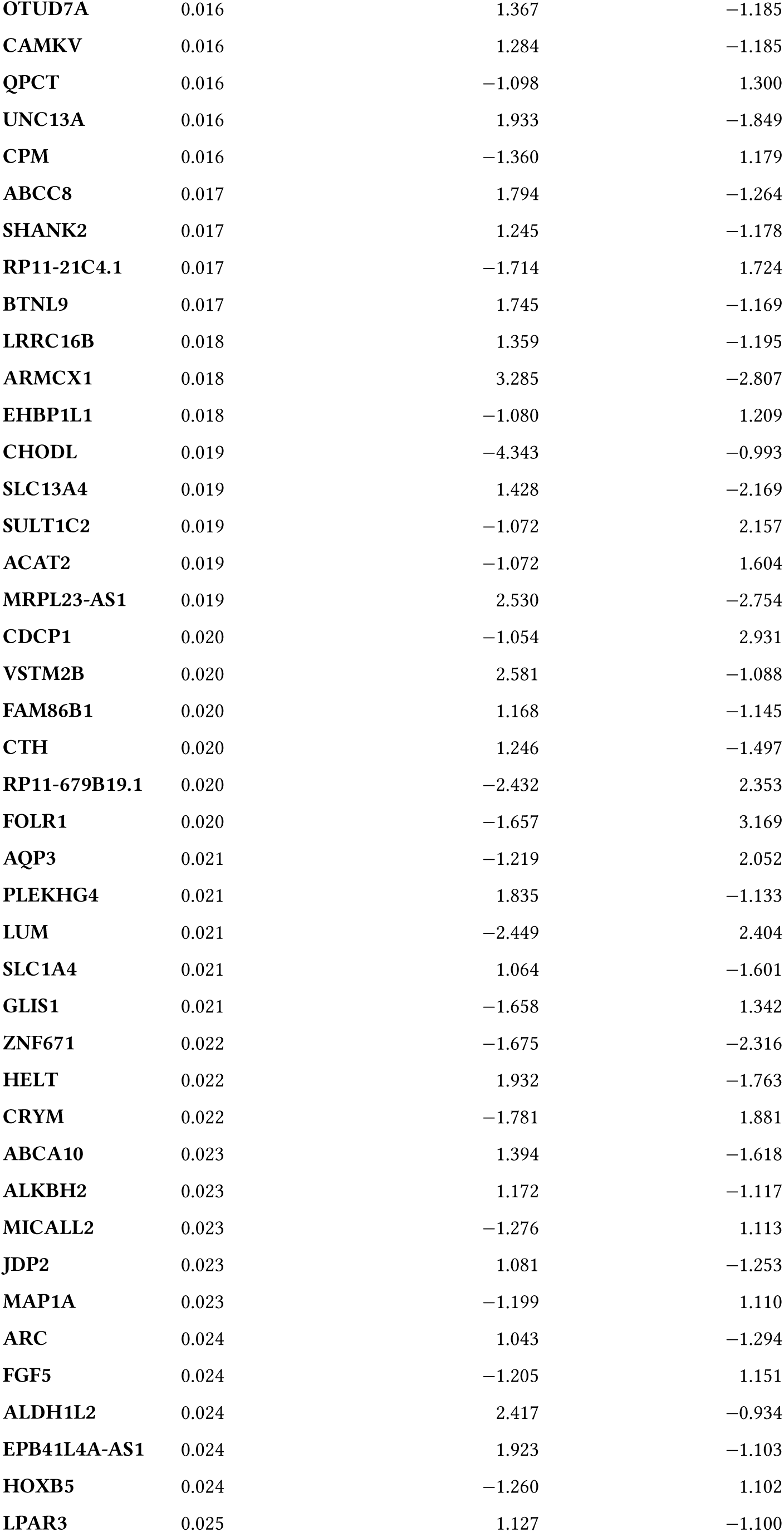
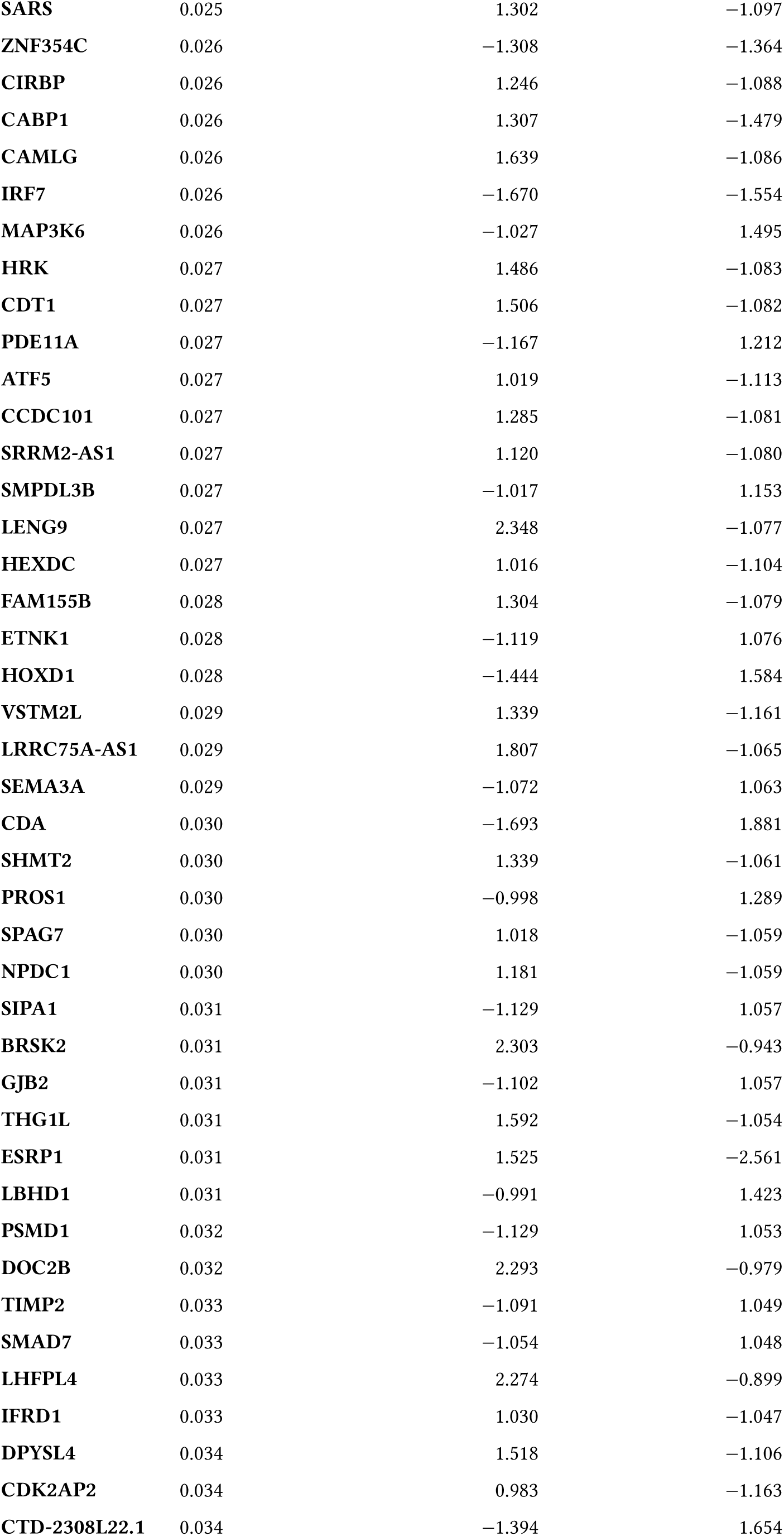
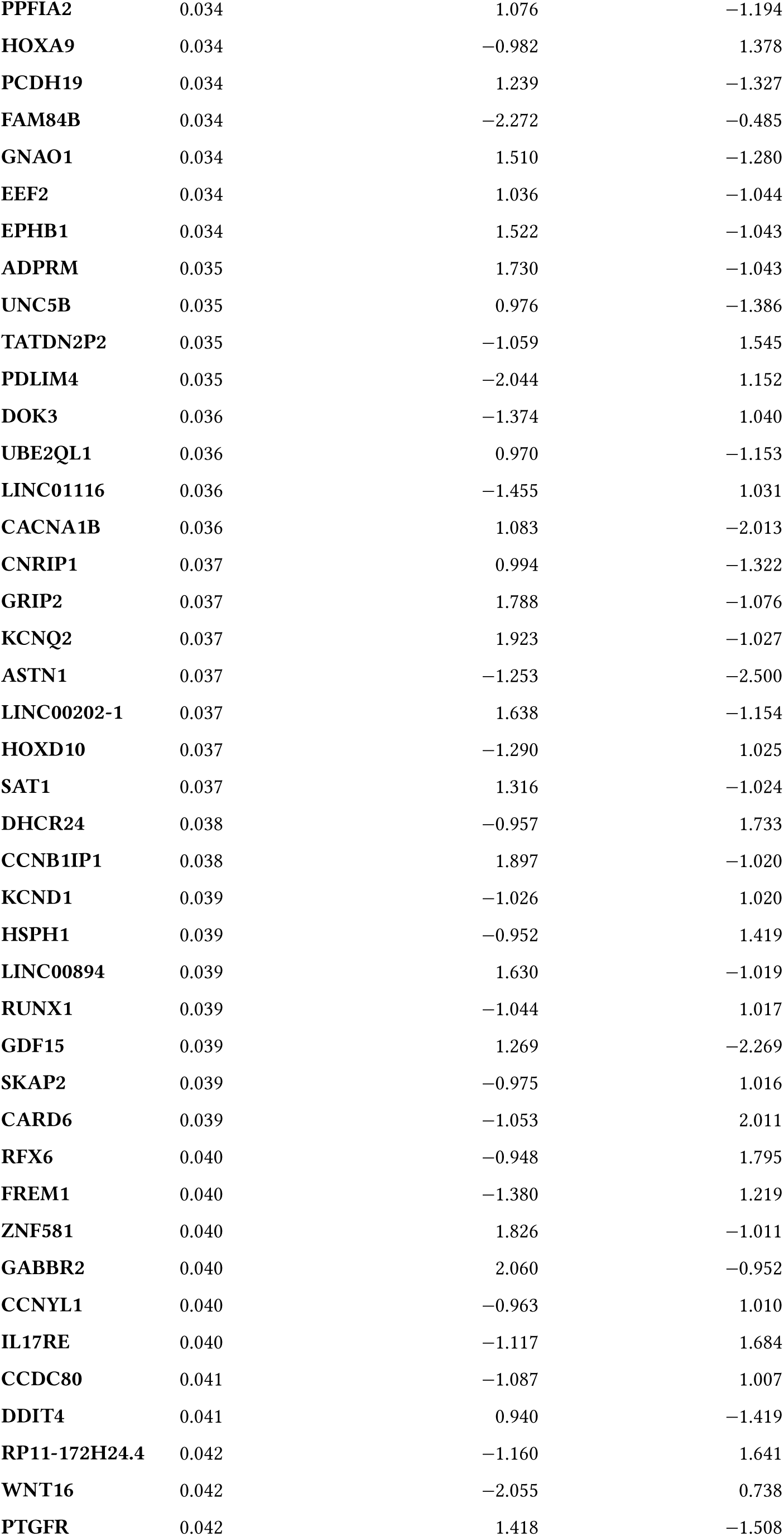
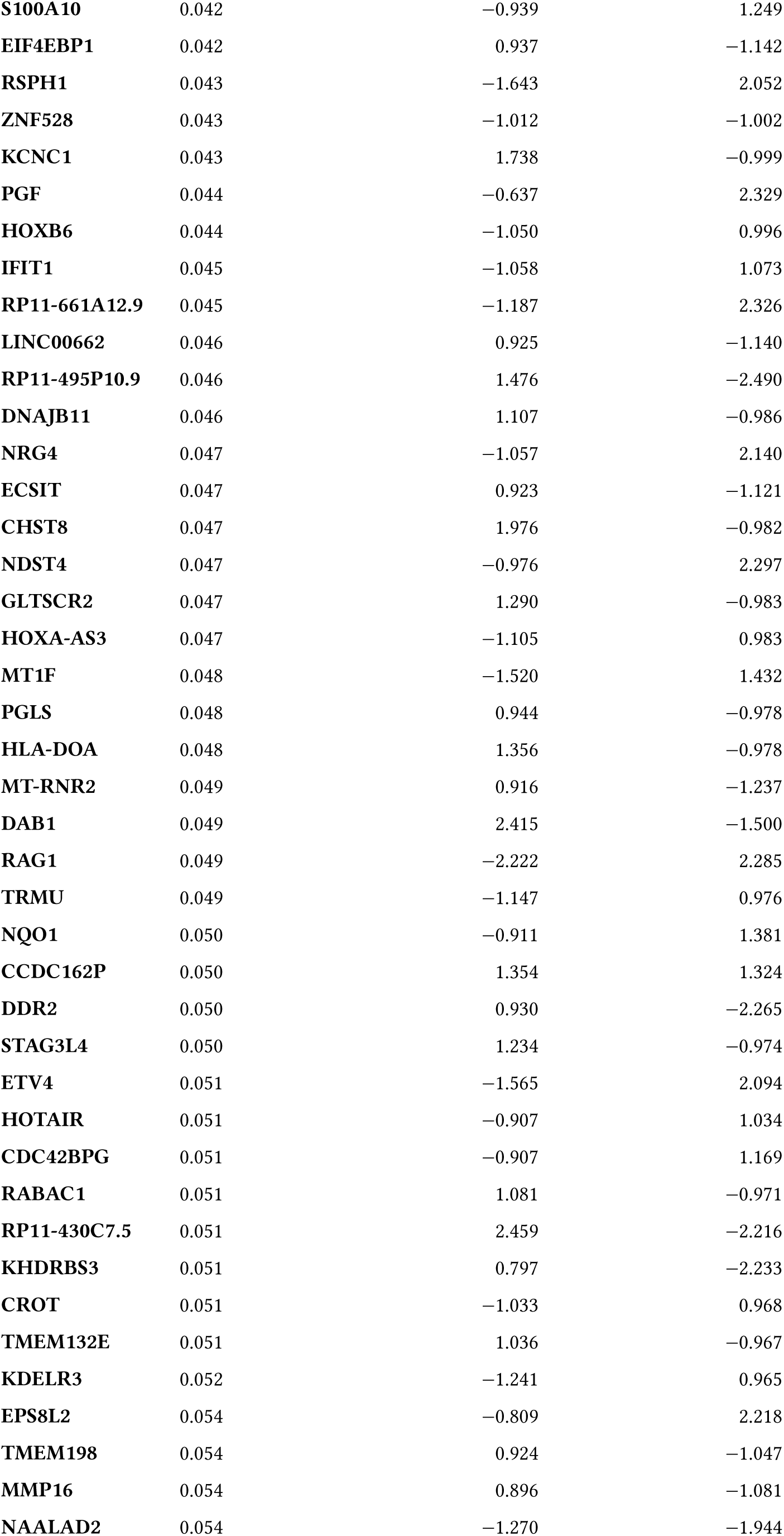
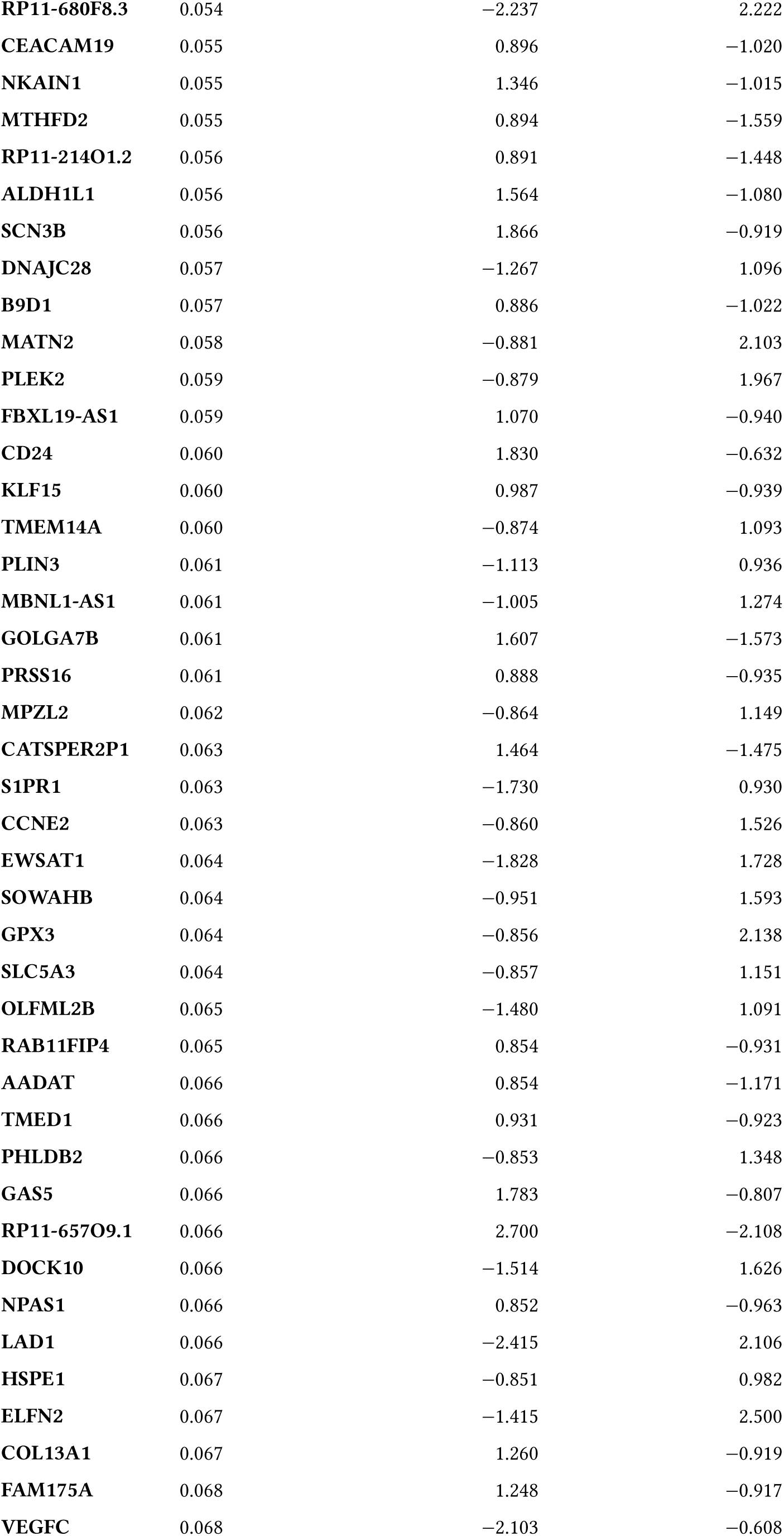
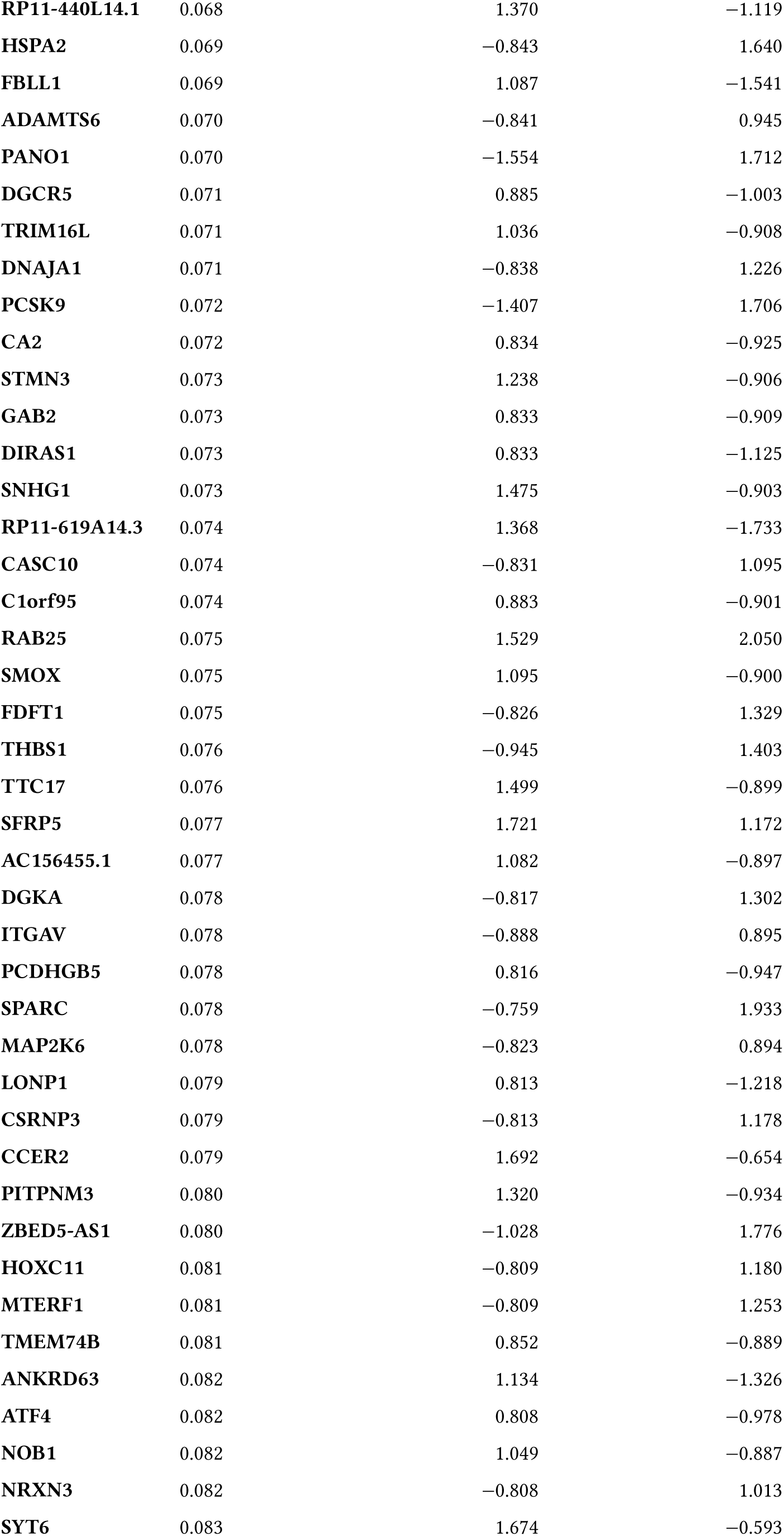
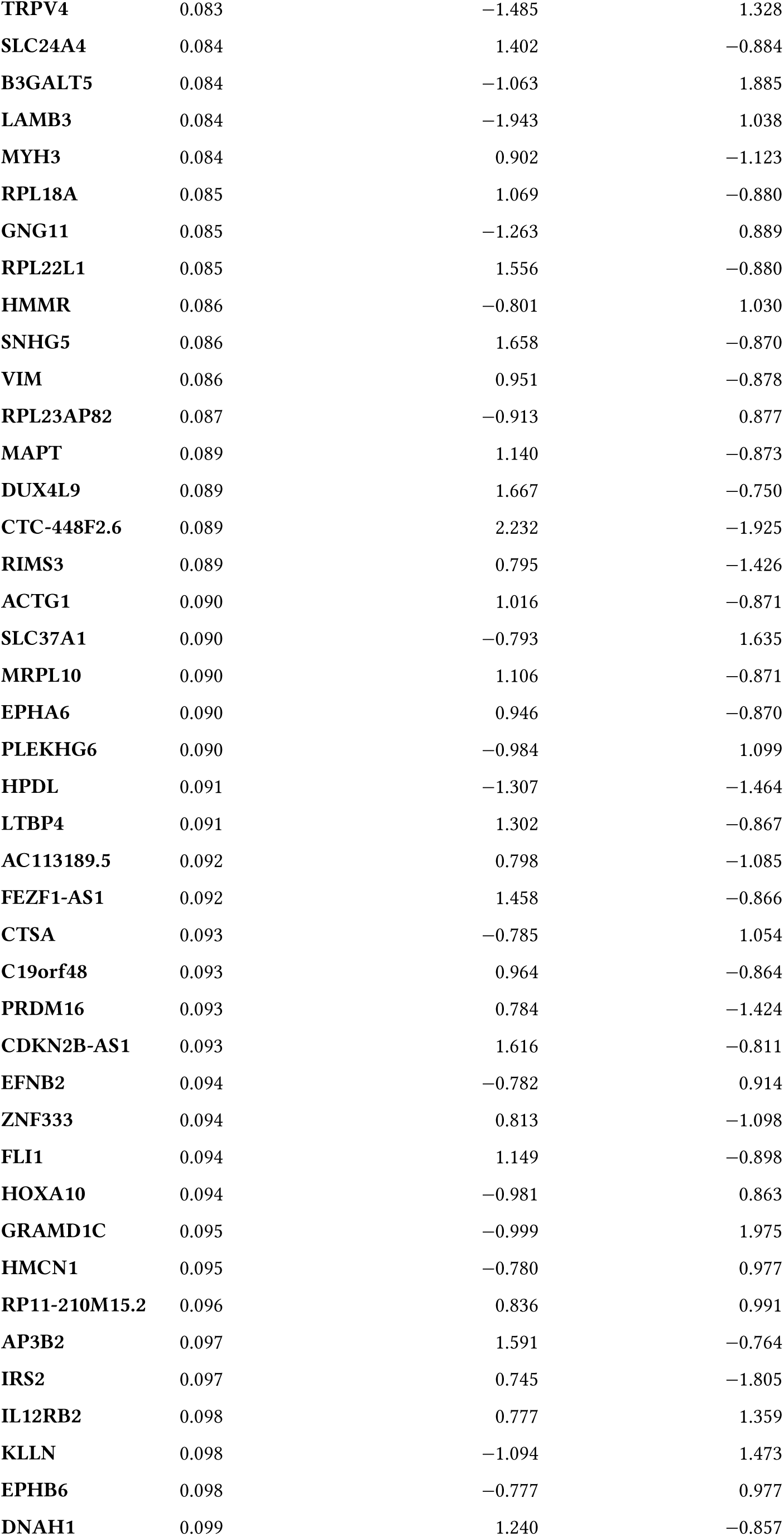
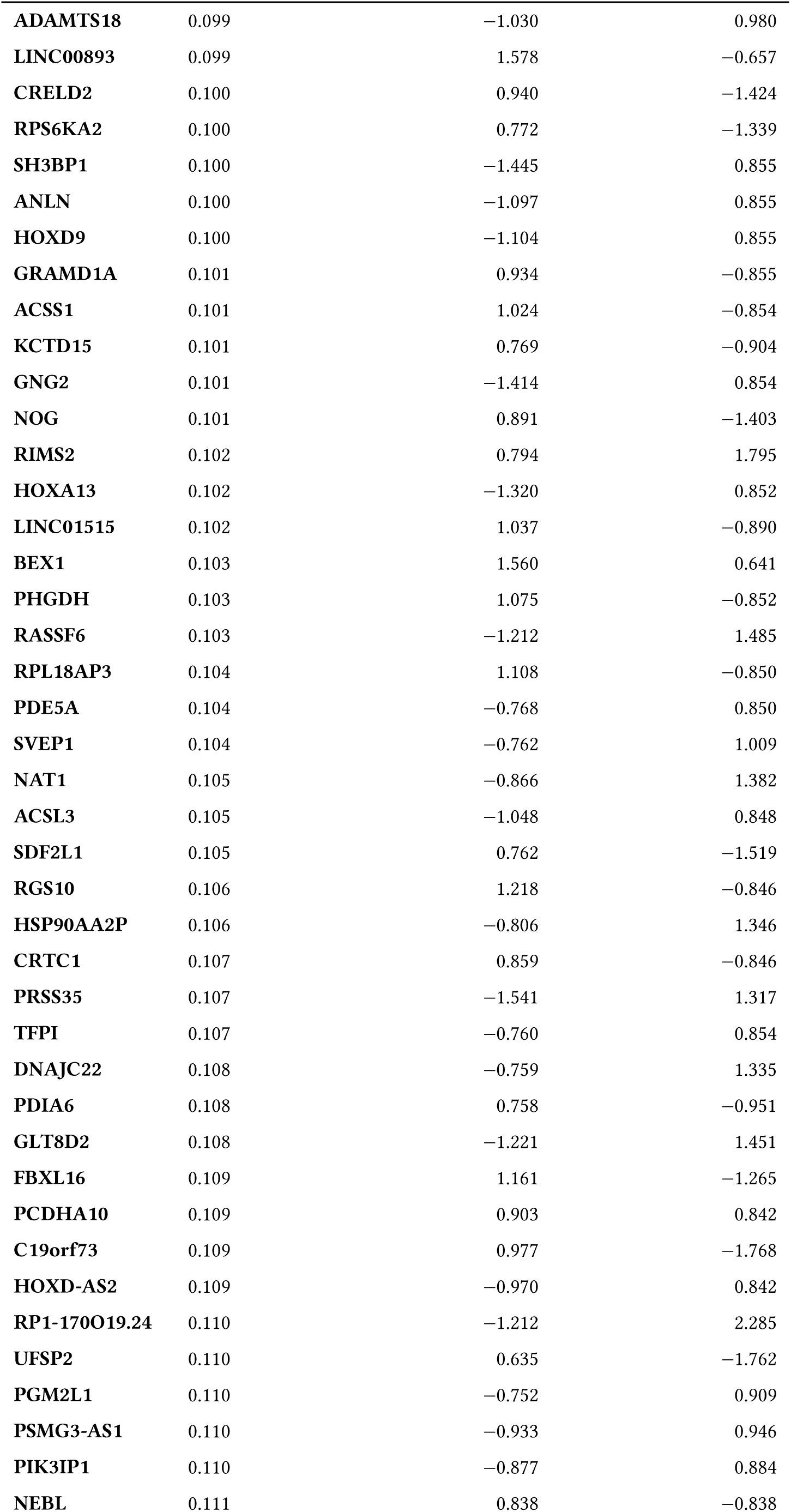
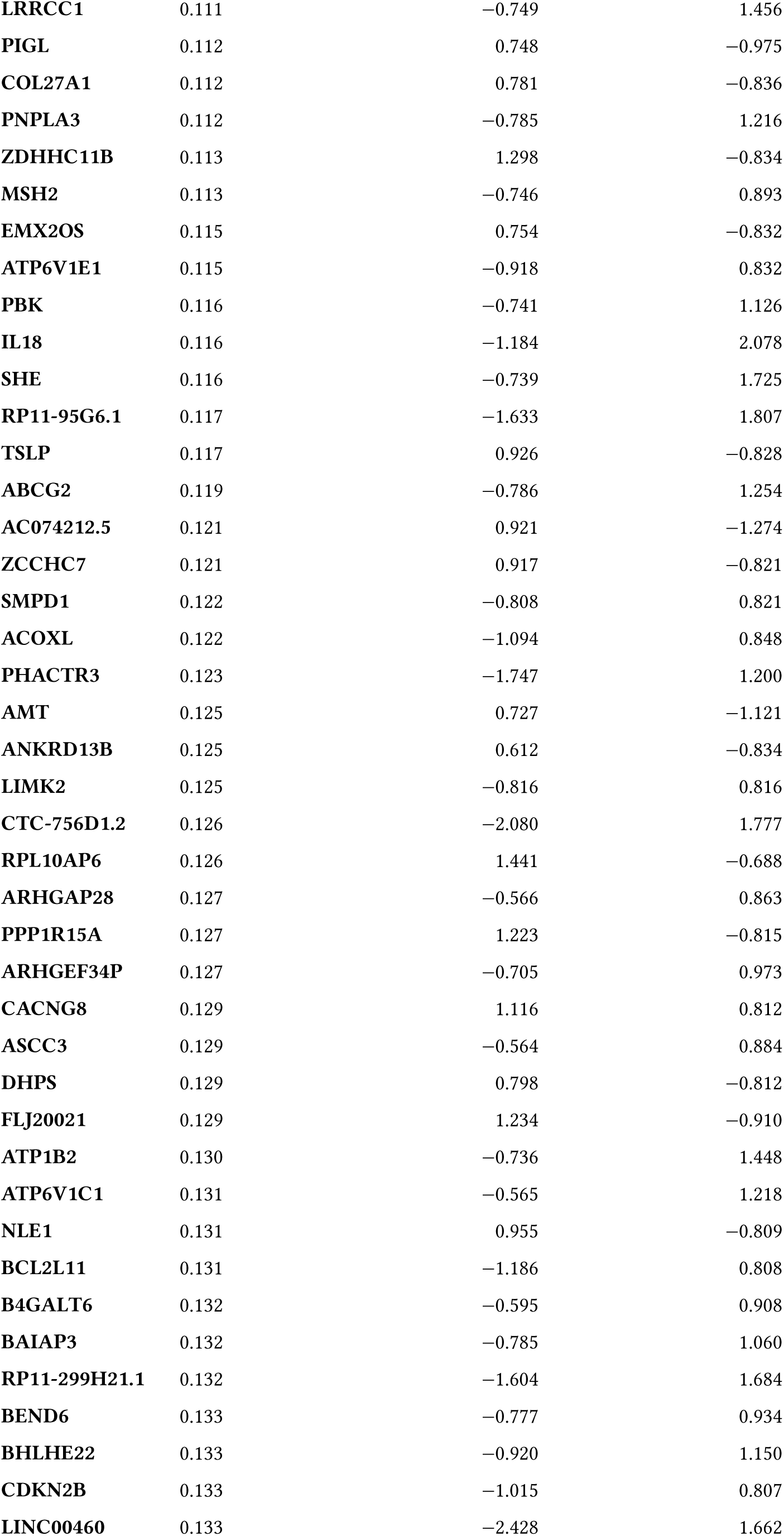
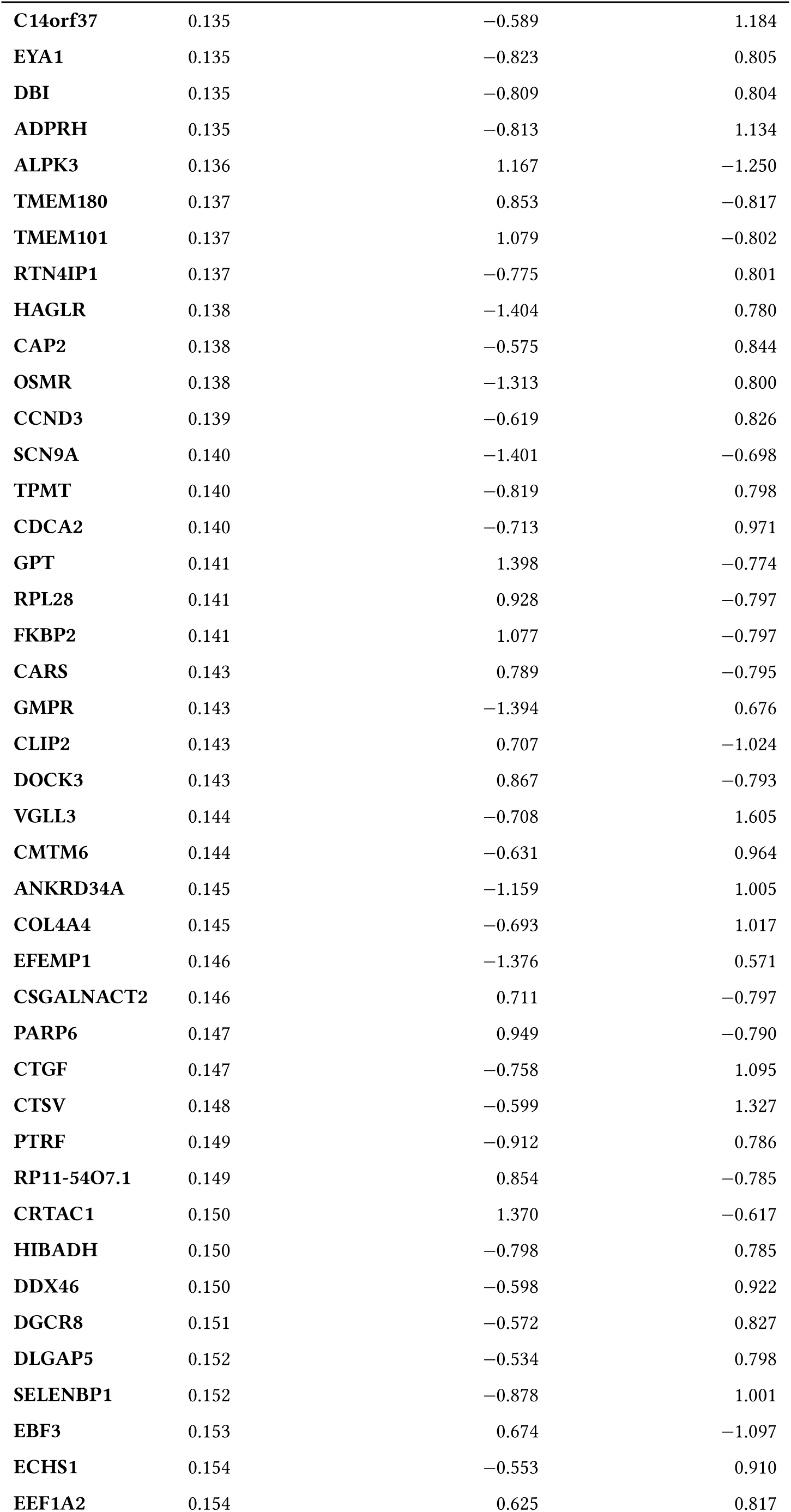
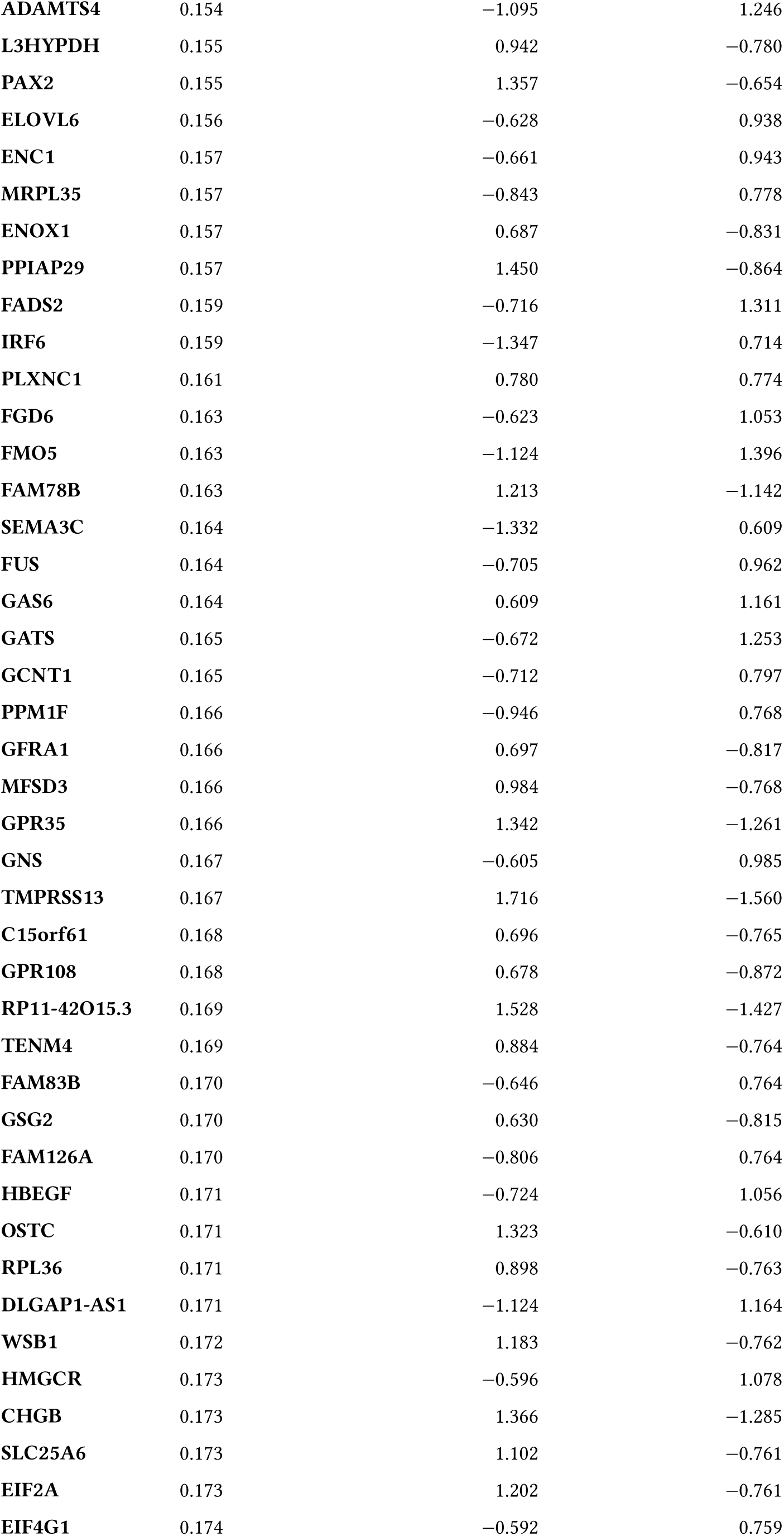
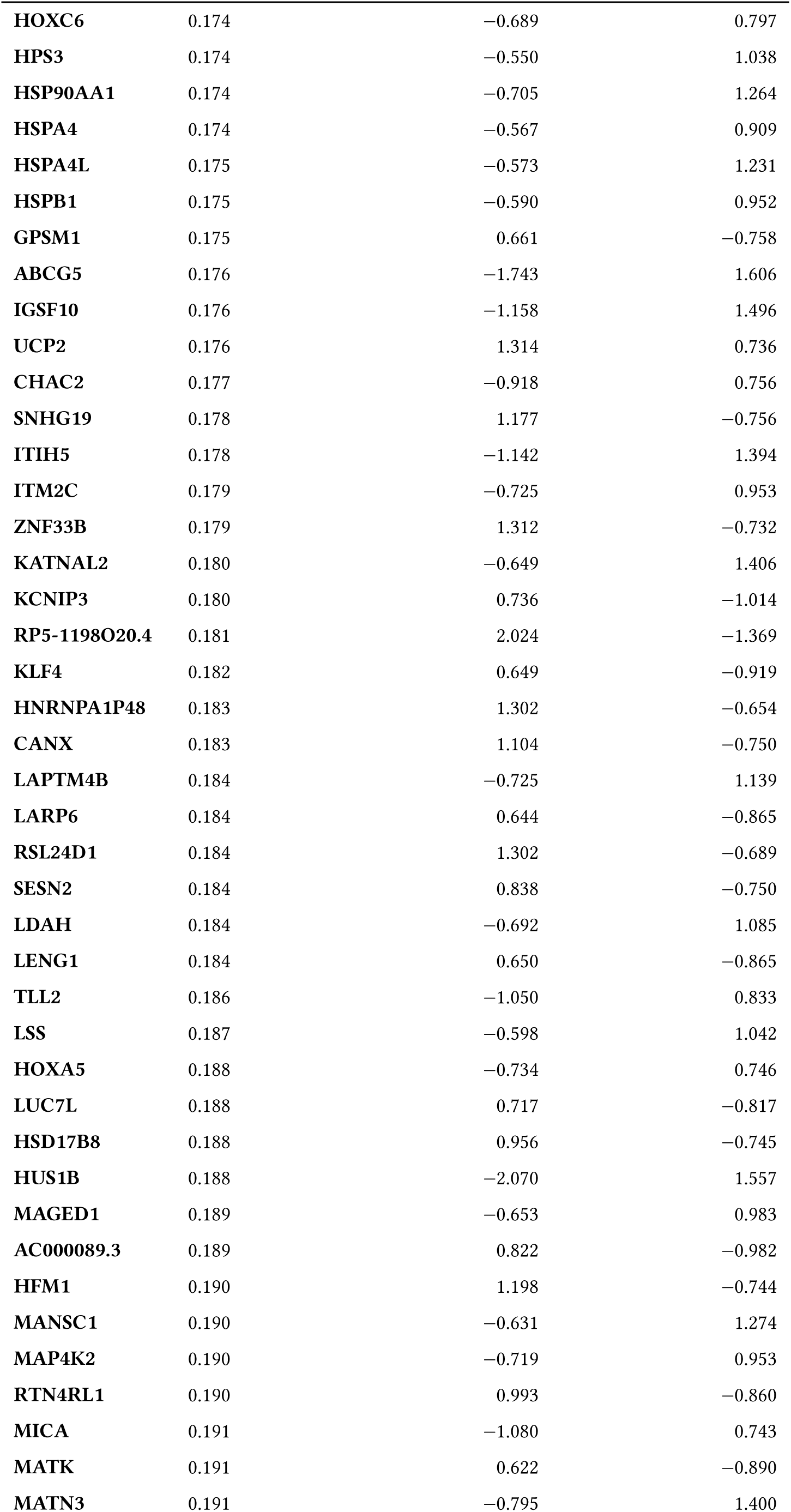
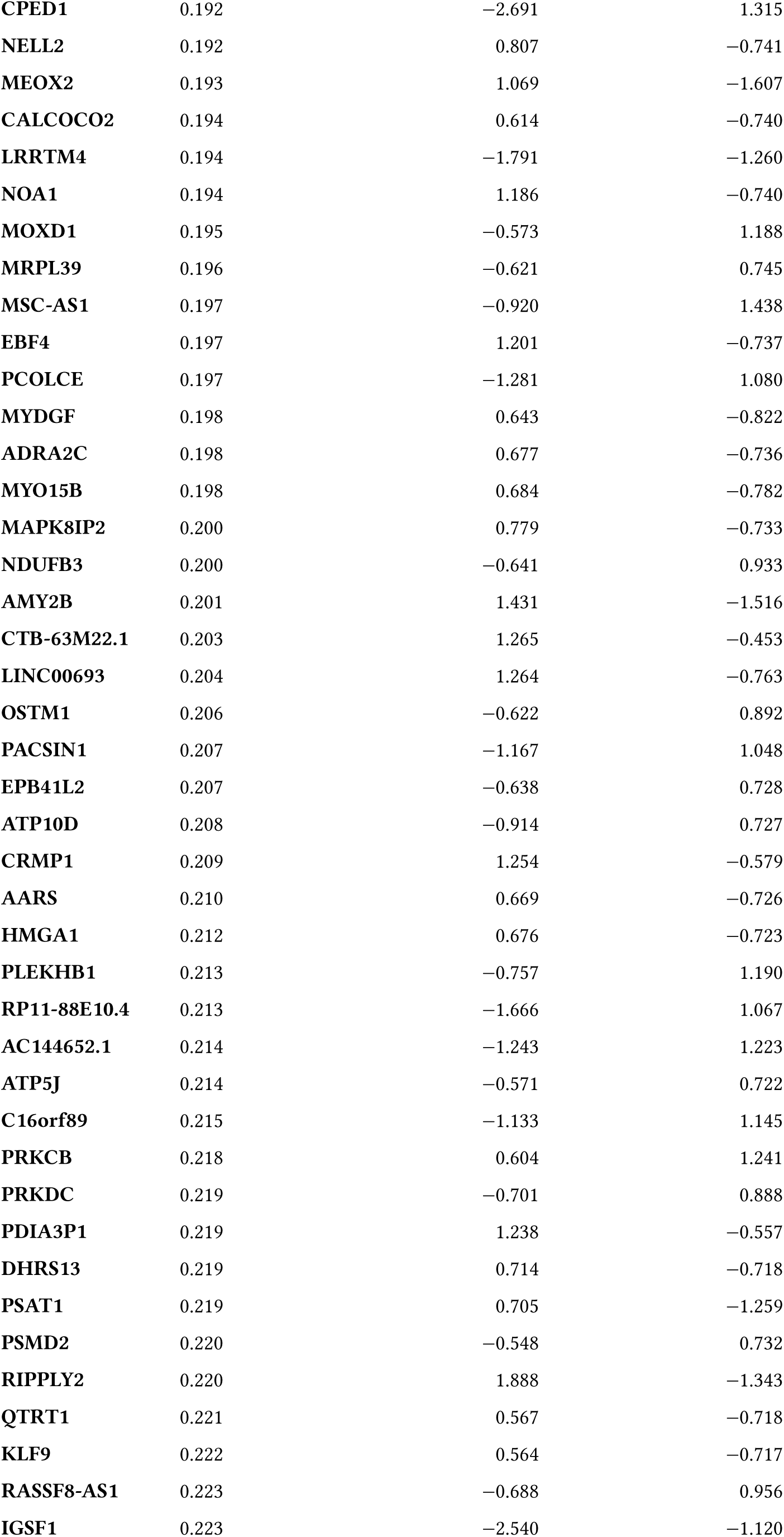
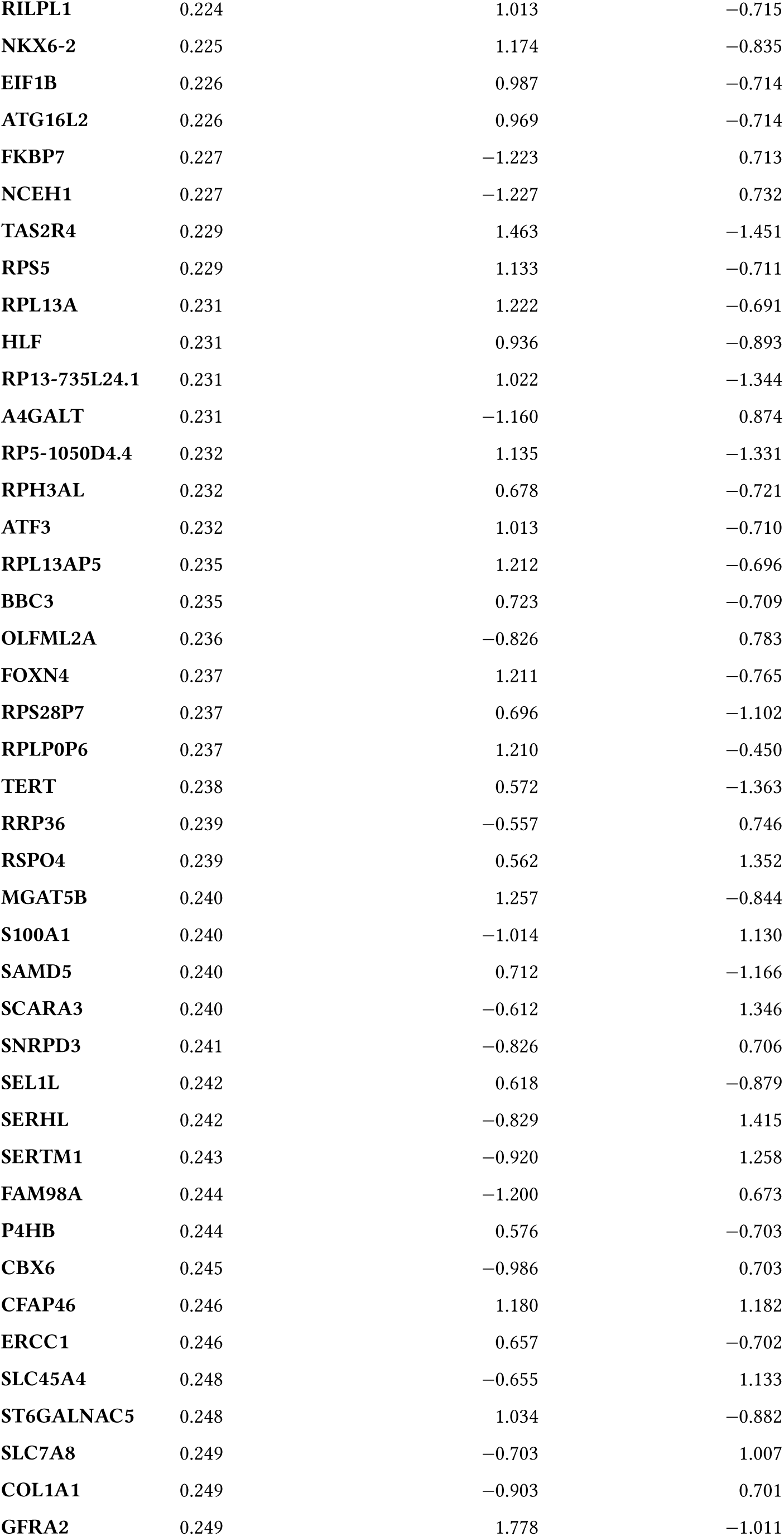
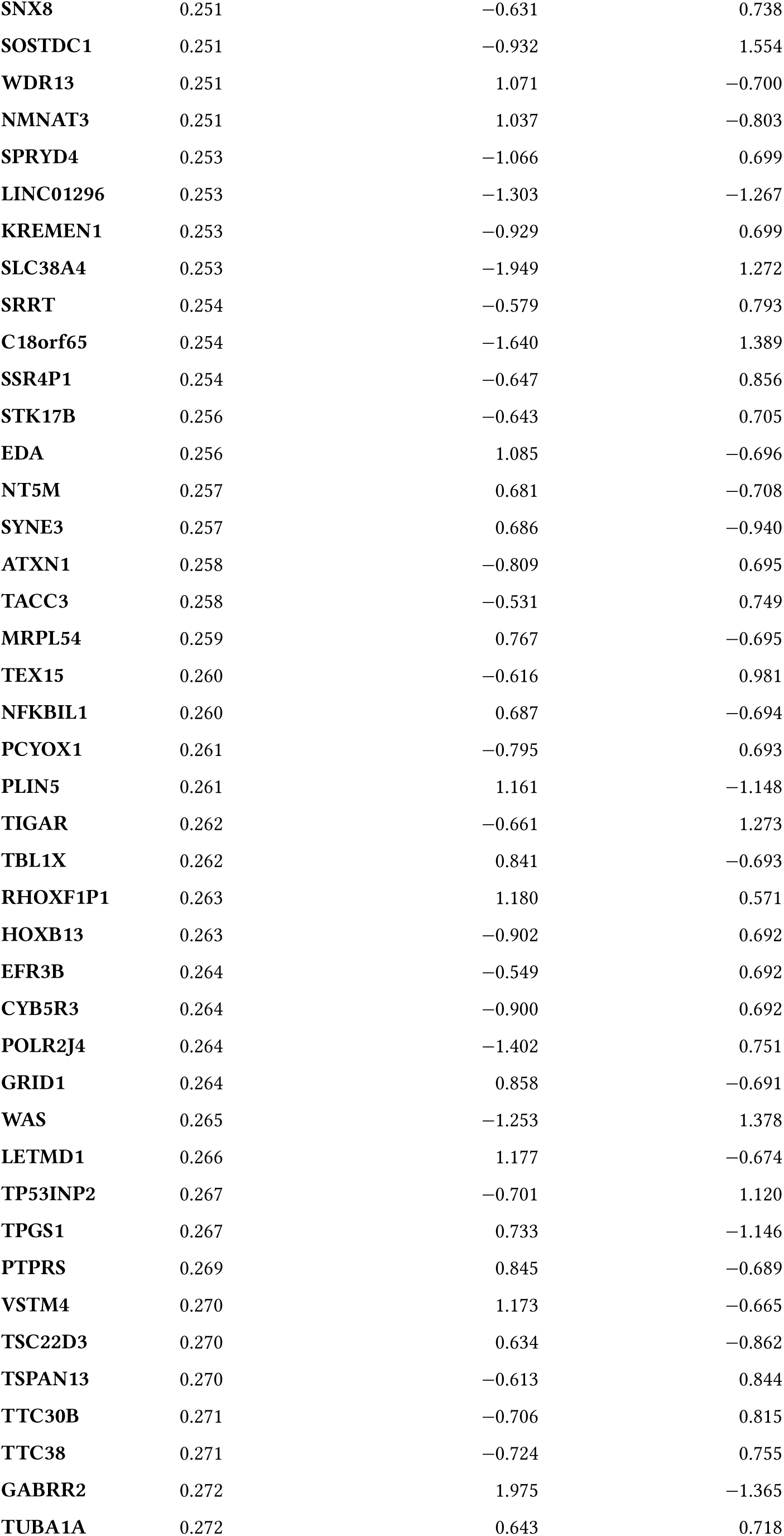
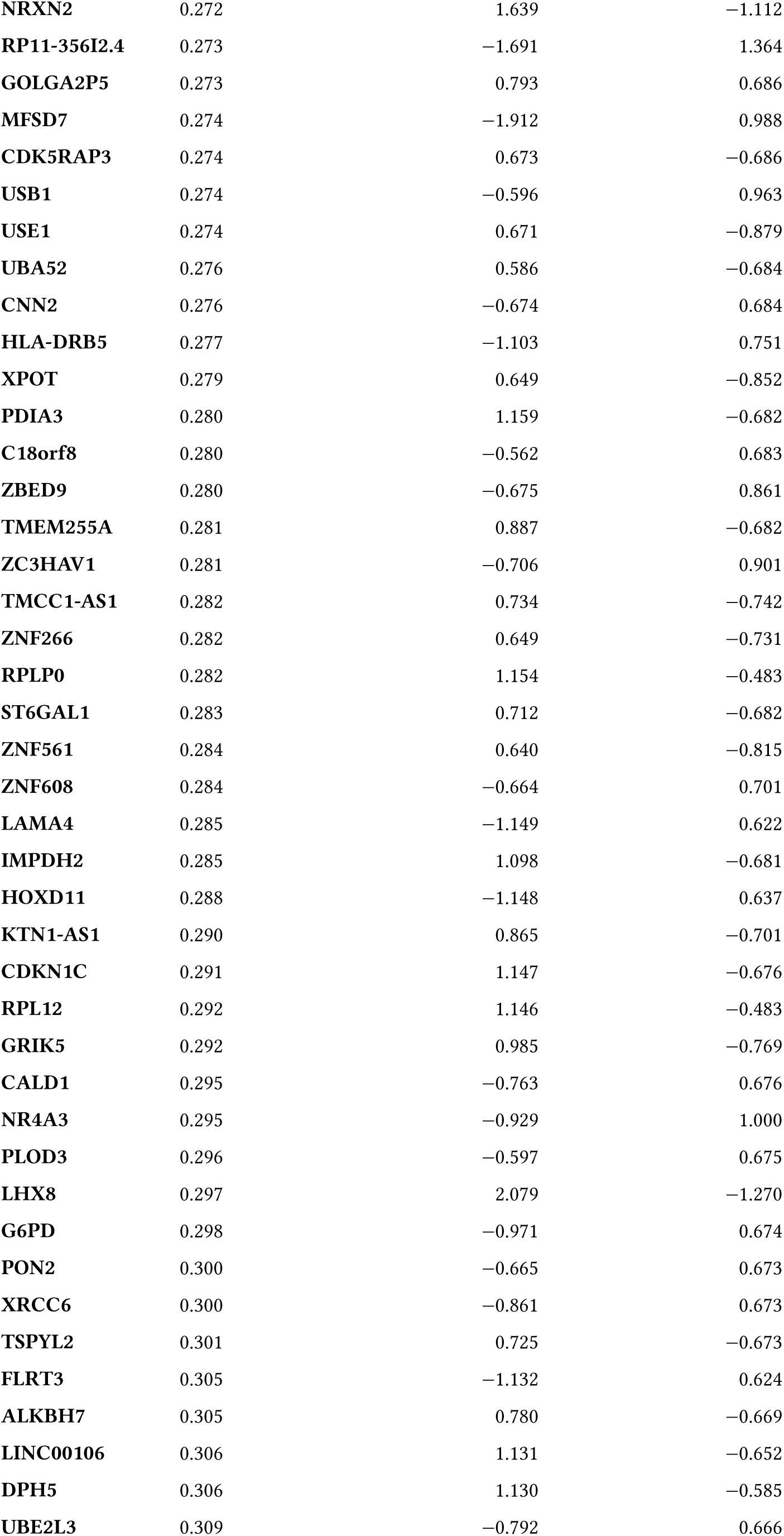
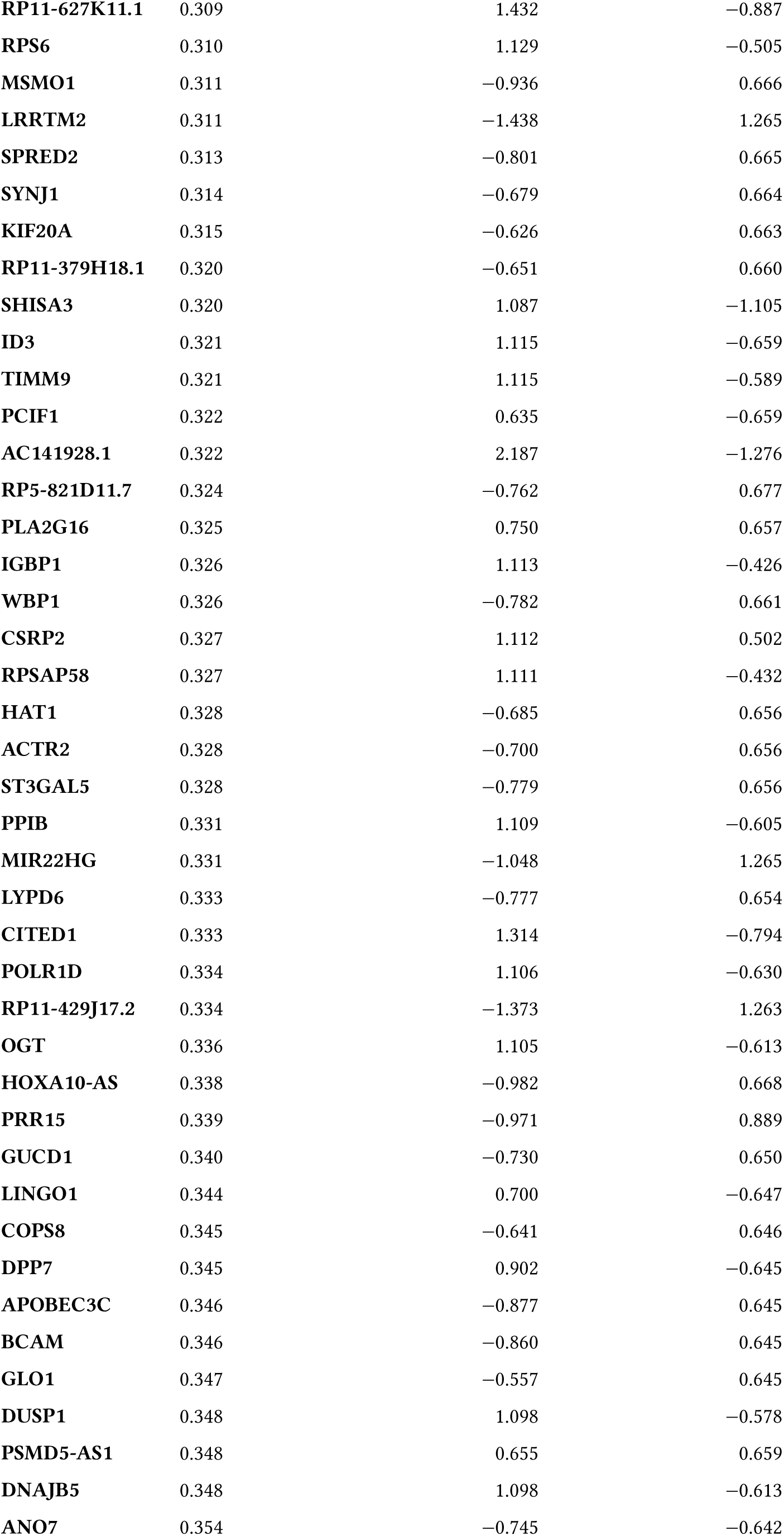
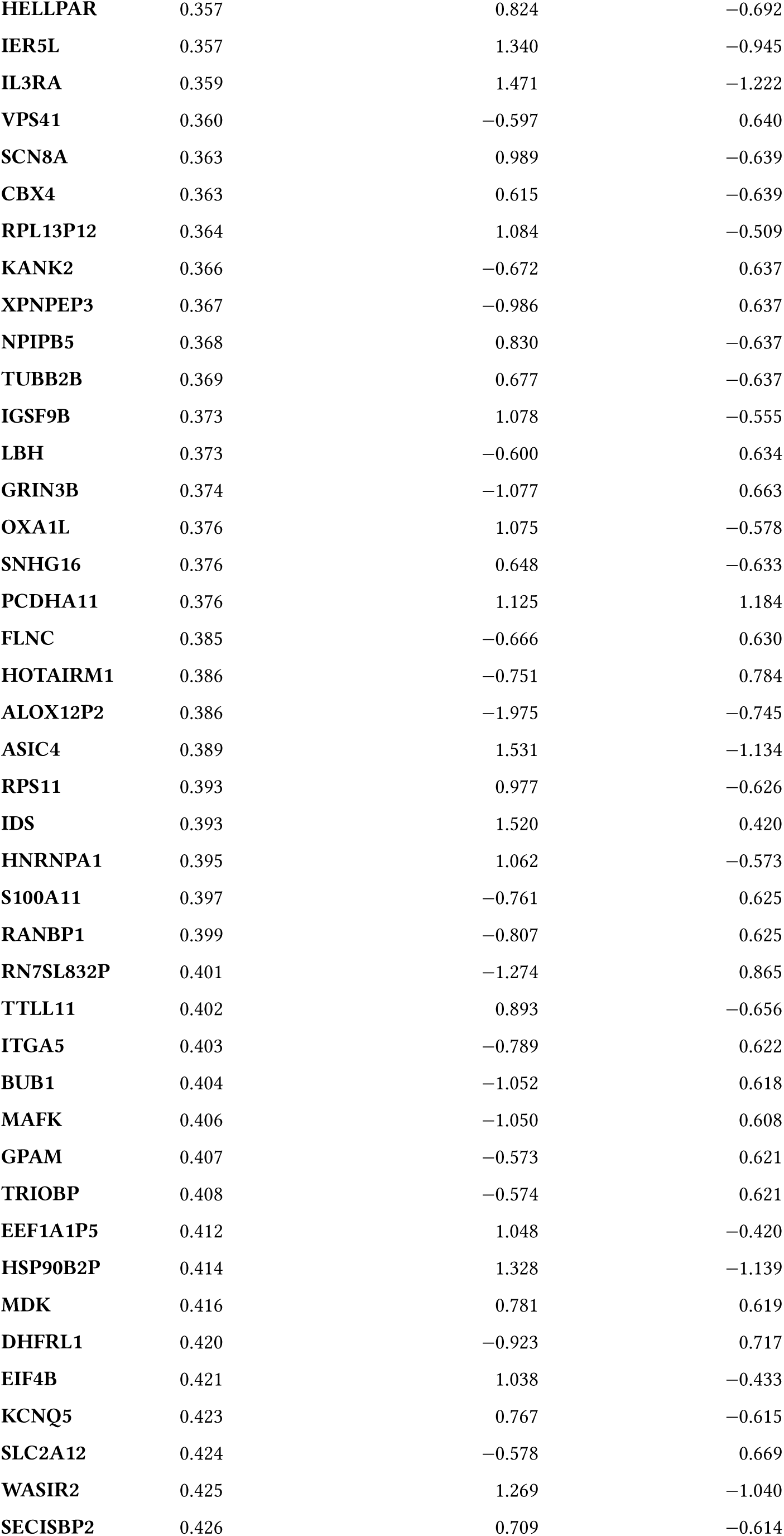
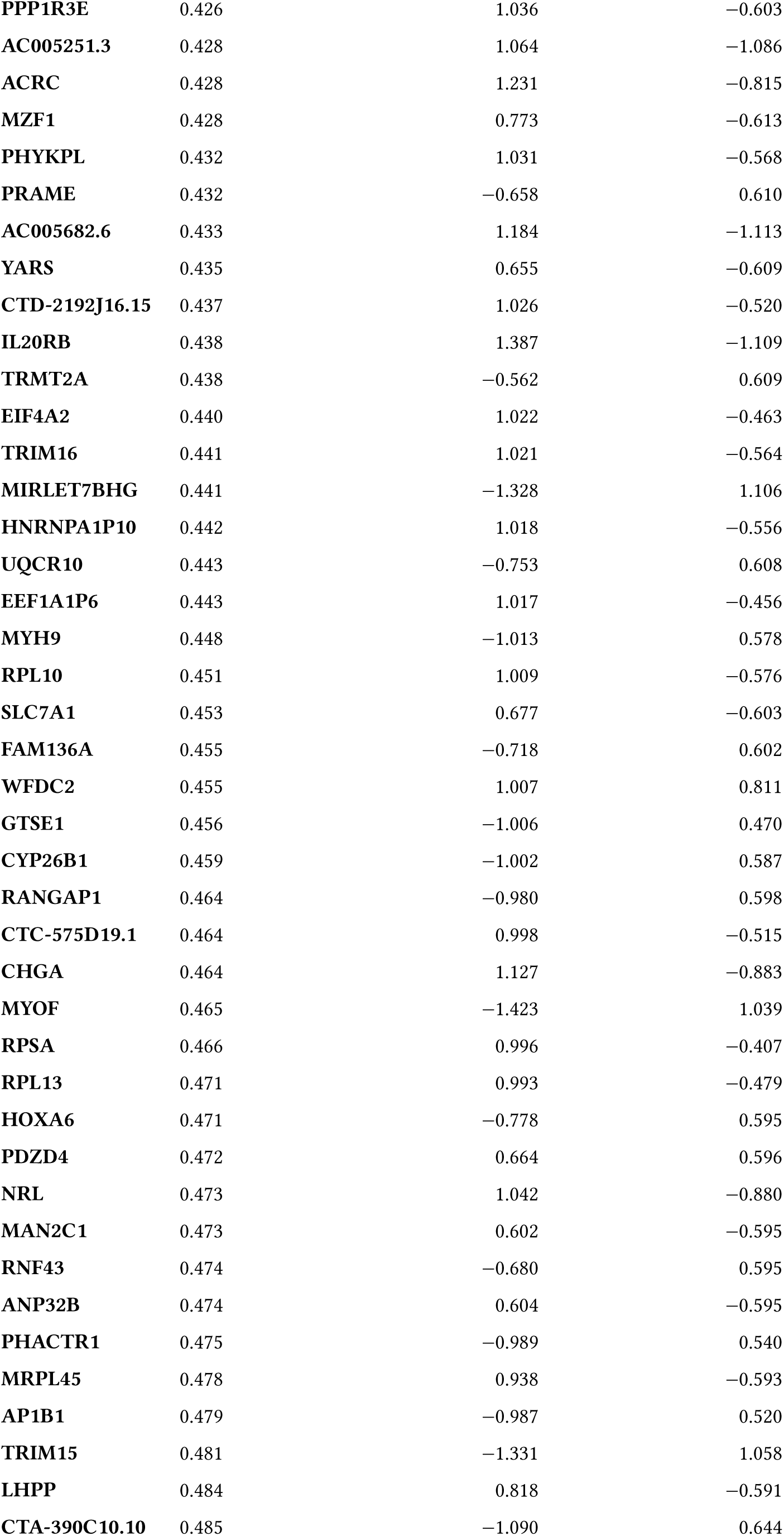
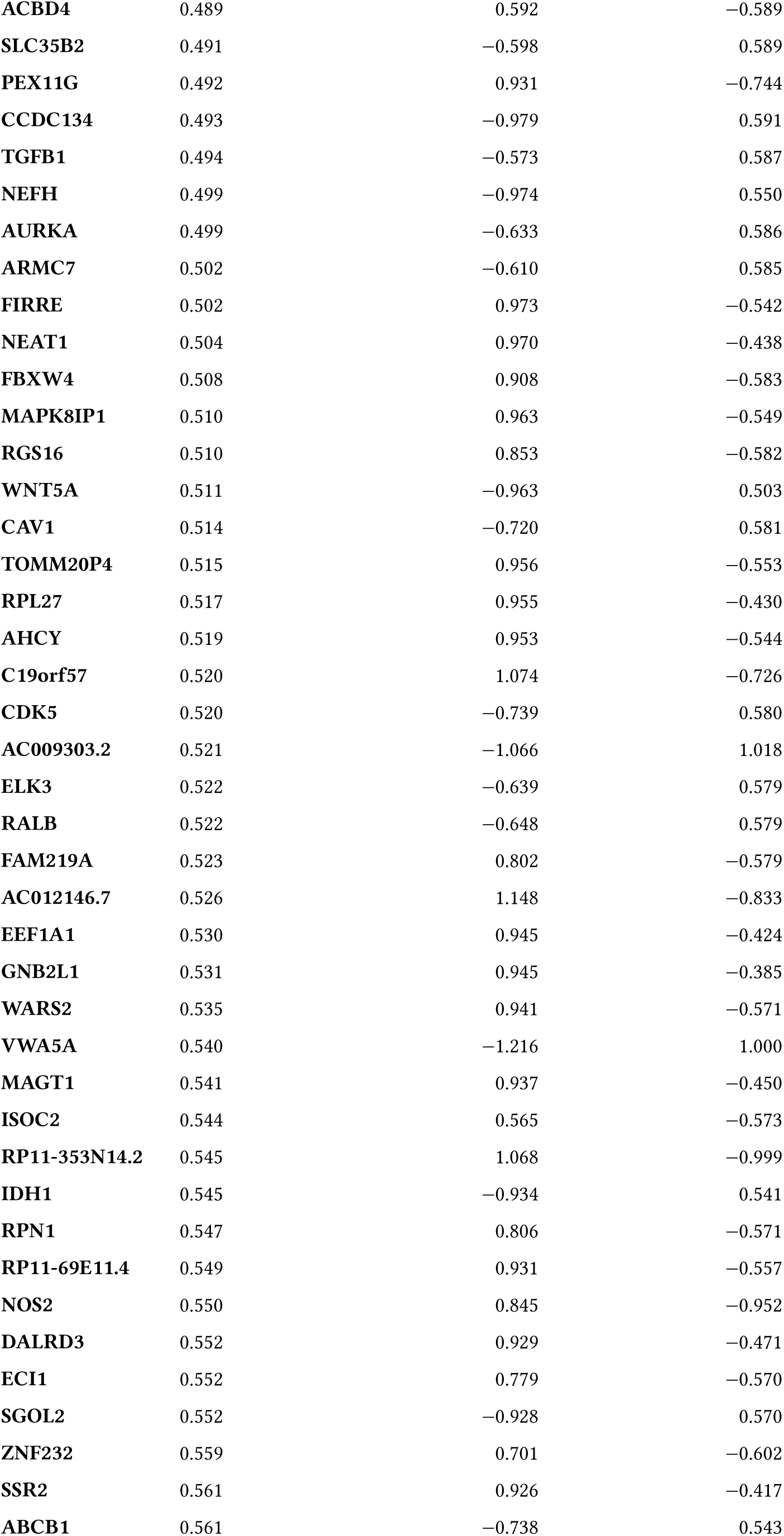
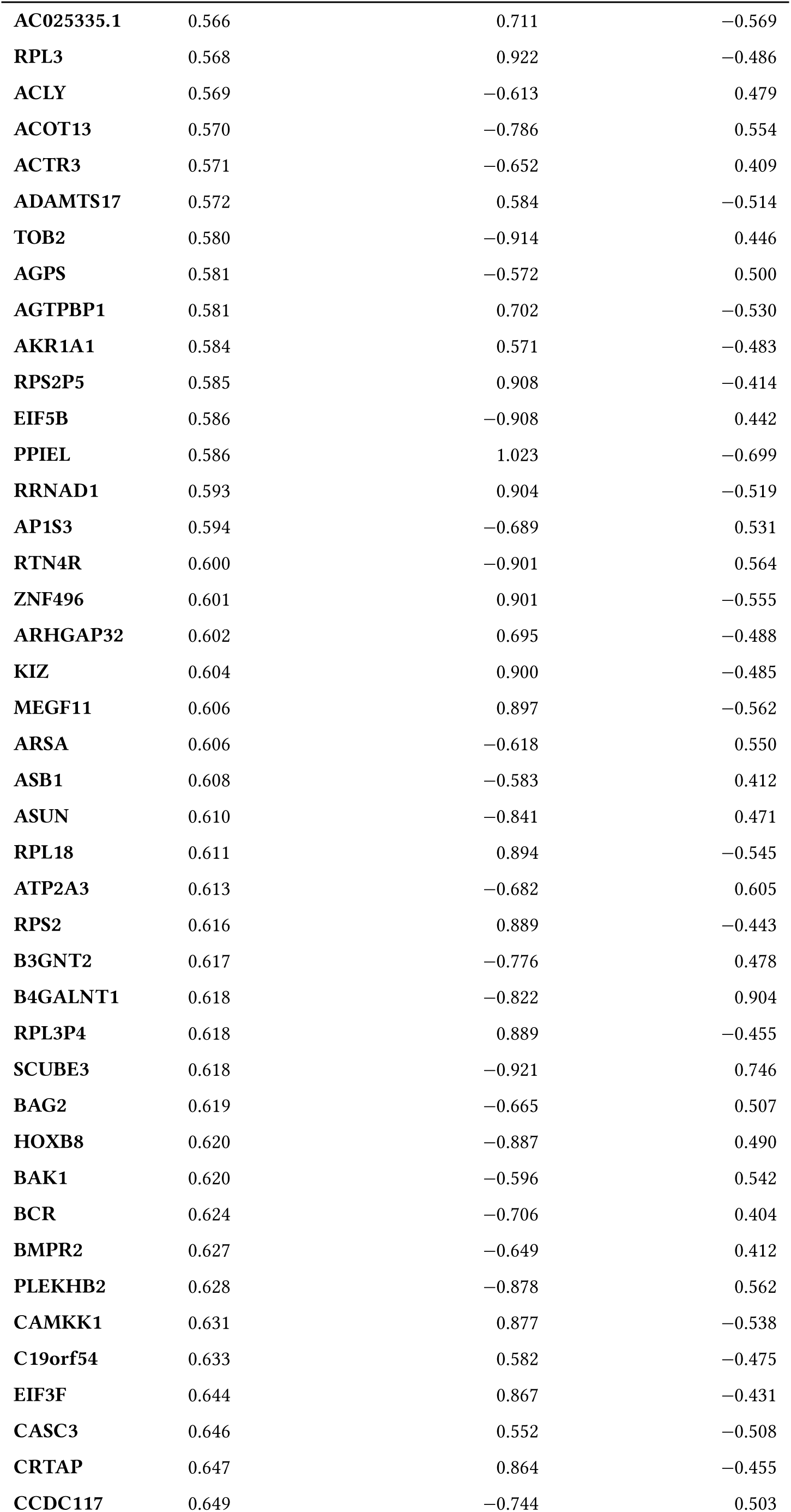
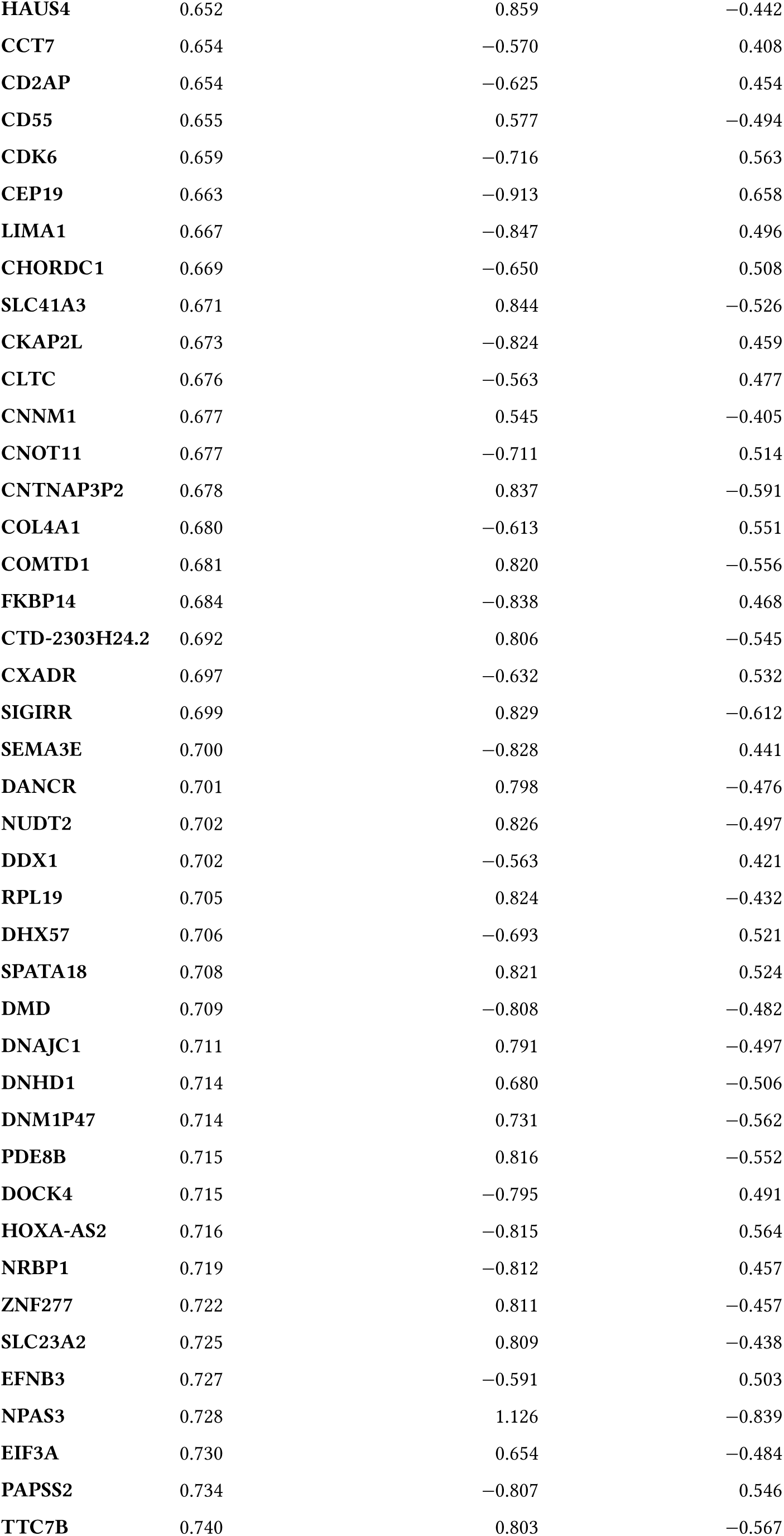
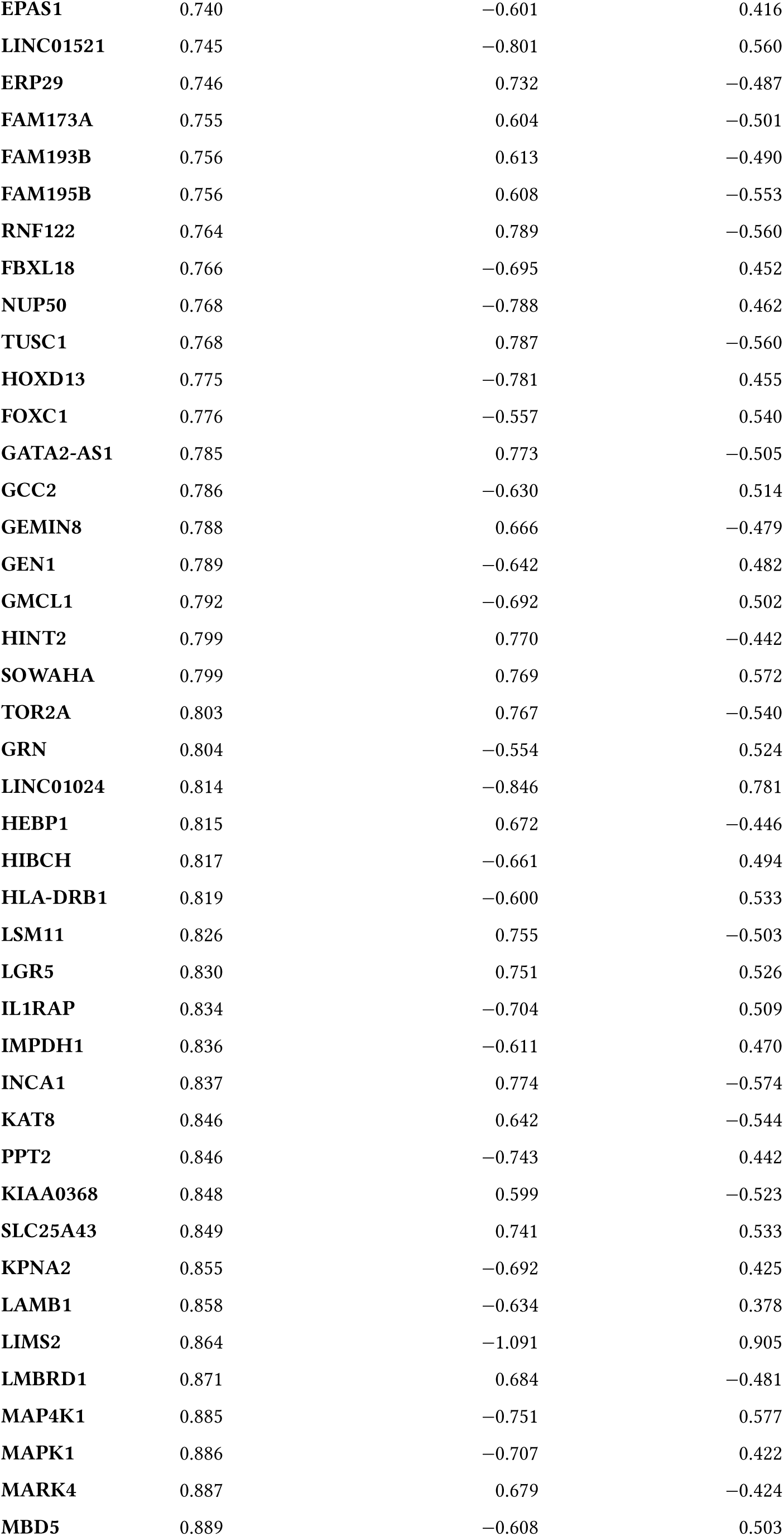
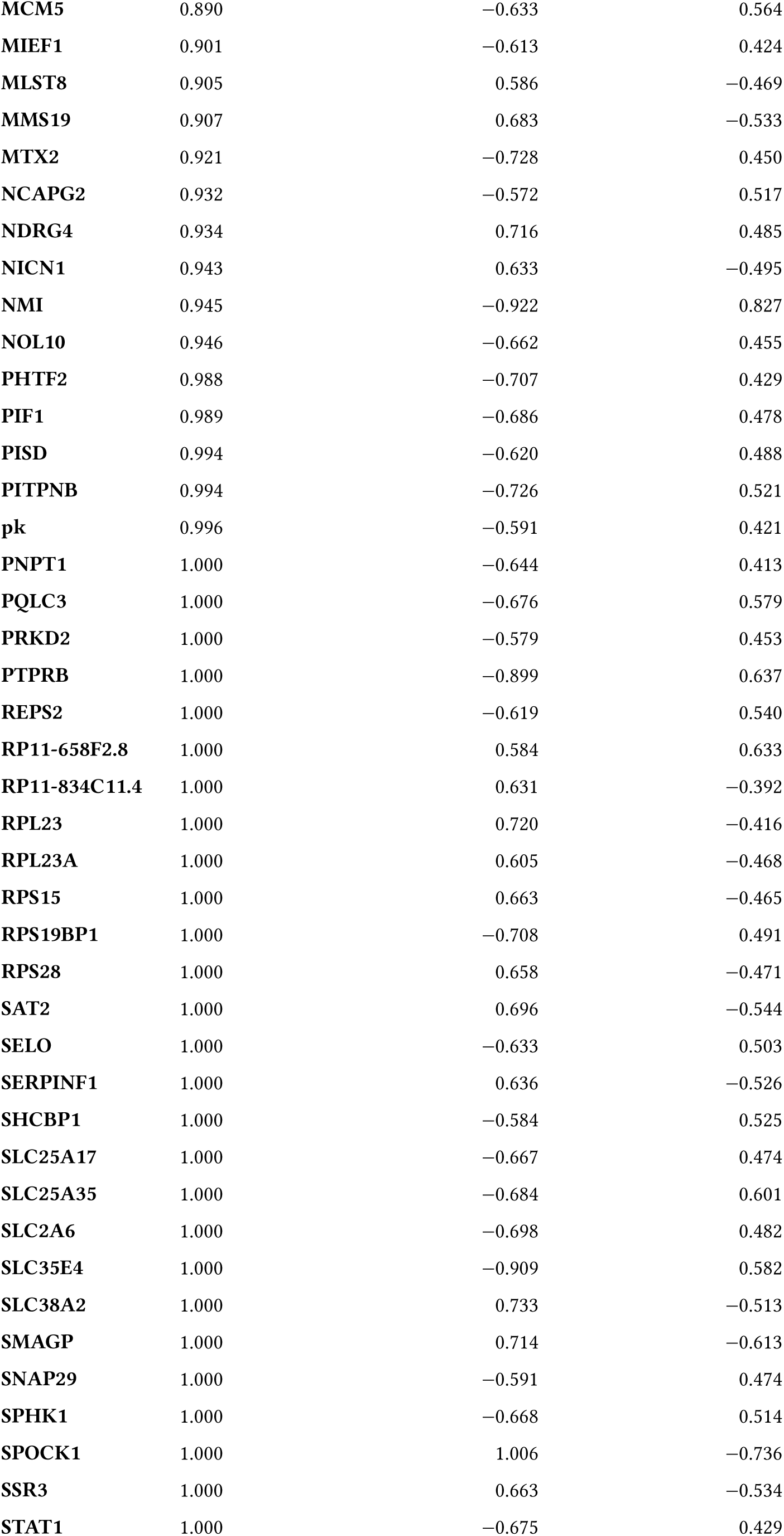
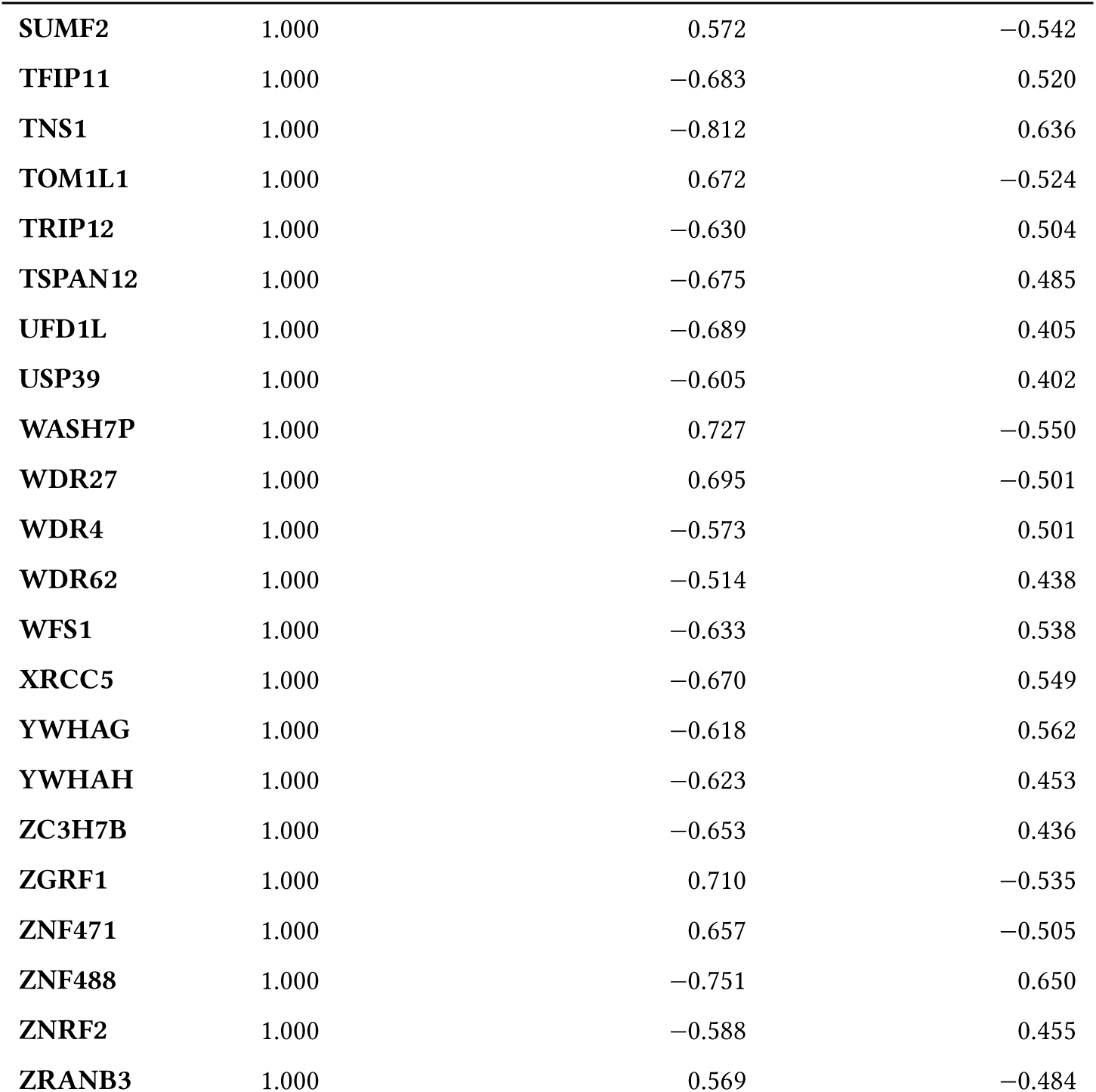
Changes in mRNA levels upon HAP40 and HTT overexpression.

**Supplementary Table 7:** Result of the transcription factor enrichment analysis against the TRRUST database.

| TF | Overlap | FDR | Odds Ratio | Combined Score | Mean log2-fold change<br>HAP40 | Mean log2-fold change<br>HTT |
| --- | --- | --- | --- | --- | --- | --- |
| ATF4 | 12/35 | 0.0 | 8.261 | 120.461 | 1.221 | -1.812 |
| XBP1 | 7/19 | 0.015 | 9.203 | 87.741 | 0.467 | -0.61 |
| ZEB2 | 4/6 | 0.023 | 31.491 | 272.547 | 0.15 | -0.333 |
| SP1 | 48/472 | 0.023 | 1.809 | 15.076 | -0.188 | 0.077 |
| HSF1 | 8/31 | 0.023 | 5.489 | 43.374 | -0.989 | 1.107 |
| E2F1 | 19/134 | 0.023 | 2.619 | 20.483 | -0.052 | -0.204 |
| WT1 | 11/57 | 0.025 | 3.779 | 28.788 | -0.448 | 0.006 |
| SREBF1 | 7/27 | 0.034 | 5.519 | 39.115 | -1.163 | 0.345 |
| SREBF2 | 6/20 | 0.034 | 6.755 | 47.731 | -1.078 | 1.297 |
| HIPK2 | 4/9 | 0.039 | 12.594 | 84.035 | -0.003 | -0.083 |
| SOX9 | 6/23 | 0.051 | 5.562 | 34.837 | -0.748 | 0.011 |
| STAT3 | 18/142 | 0.052 | 2.298 | 14.181 | -0.553 | 0.381 |
| FOXO3 | 5/18 | 0.068 | 6.057 | 34.432 | -0.857 | -0.432 |
| LMX1B | 3/6 | 0.068 | 15.731 | 87.923 | -0.644 | 0.674 |
| VHL | 5/20 | 0.092 | 5.249 | 27.249 | -0.875 | 0.208 |
| TP73 | 4/13 | 0.092 | 6.995 | 35.883 | -0.358 | 0.446 |
| TCF3 | 4/14 | 0.106 | 6.295 | 30.476 | -0.931 | 0.986 |
| CIITA | 6/31 | 0.106 | 3.781 | 17.713 | -0.596 | 0.523 |
| DNMT1 | 6/31 | 0.106 | 3.781 | 17.713 | 0.506 | -0.19 |
| SMAD3 | 6/31 | 0.106 | 3.781 | 17.713 | 0.004 | -0.099 |
| HDAC1 | 10/71 | 0.106 | 2.586 | 12.056 | -0.581 | -0.042 |
| HOXA10 | 4/15 | 0.106 | 5.723 | 26.203 | 0.467 | -0.274 |
| RFXANK | 4/15 | 0.106 | 5.723 | 26.203 | -0.5 | 0.502 |
| RFXAP | 4/15 | 0.106 | 5.723 | 26.203 | -0.5 | 0.502 |
| ATF2 | 6/32 | 0.106 | 3.635 | 16.461 | 0.456 | -0.214 |
| REST | 6/32 | 0.106 | 3.635 | 16.461 | 0.09 | -0.839 |
| FOXO1 | 5/24 | 0.117 | 4.143 | 18.152 | -1.248 | 0.153 |
| HDAC4 | 5/24 | 0.117 | 4.143 | 18.152 | 0.066 | -0.241 |
| ATF3 | 4/16 | 0.119 | 5.246 | 22.76 | 0.918 | -1.323 |
| SMAD7 | 3/9 | 0.122 | 7.864 | 33.736 | -1.032 | 1.067 |
| JUND | 6/34 | 0.125 | 3.375 | 14.293 | -0.692 | 0.873 |
| TCF4 | 5/25 | 0.125 | 3.936 | 16.56 | 0.58 | -0.506 |
| ESR1 | 10/76 | 0.125 | 2.39 | 10.043 | -0.38 | 0.03 |
| SMAD4 | 5/26 | 0.139 | 3.748 | 15.155 | -0.567 | 0.997 |
| SNAI1 | 3/10 | 0.139 | 6.741 | 26.814 | -0.768 | 0.997 |
| BRCA1 | 8/57 | 0.143 | 2.573 | 10.113 | -0.143 | -0.387 |
| RFX5 | 4/18 | 0.143 | 4.496 | 17.598 | -0.5 | 0.502 |
| SIRT1 | 7/48 | 0.164 | 2.689 | 10.111 | -1.019 | 0.046 |
| NPM1 | 3/11 | 0.168 | 5.898 | 21.847 | 0.145 | 0.068 |
| HIF1A | 10/83 | 0.173 | 2.16 | 7.861 | -0.353 | -0.268 |
| KLF6 | 4/20 | 0.181 | 3.933 | 13.962 | -0.347 | 0.393 |

|  | TF | Overlap | FDR | Odds Ratio | Combined Score | Mean log2-fold change<br>HAP40 | Mean log2-fold change<br>HTT |
| --- | --- | --- | --- | --- | --- | --- | --- |
| 3773 |  |  |  |  |  |  |  |
| 3774 |  |  |  |  |  |  |  |
| 3775 | EGR1 | 10/88 | 0.23 | 2.021 | 6.641 | -0.351 | -0.245 |
| 3776 | E2F3 | 3/13 | 0.23 | 4.718 | 15.301 | -0.286 | 0.197 |
| 3777 | ZEB1 | 3/13 | 0.23 | 4.718 | 15.301 | -1.971 | 0.471 |
| 3778 | ATM | 4/22 | 0.23 | 3.496 | 11.298 | 0.033 | 0.138 |
| 3779 | ETV4 | 4/22 | 0.23 | 3.496 | 11.298 | -0.047 | -0.092 |
| 3780 | HDGF | 2/6 | 0.249 | 7.859 | 24.239 | 1.118 | -1.34 |
| 3781 | WWTR1 | 2/6 | 0.249 | 7.859 | 24.239 | 0.486 | -0.737 |
| 3782 | ATF6 | 3/14 | 0.255 | 4.289 | 13.066 | 0.869 | -1.143 |
| 3783 | MYCN | 6/45 | 0.267 | 2.422 | 7.231 | -0.652 | 0.702 |
| 3784 | AR | 10/93 | 0.268 | 1.898 | 5.634 | -0.918 | -0.254 |
| 3785 | FOS | 7/57 | 0.272 | 2.204 | 6.48 | 0.099 | 0.036 |
| 3786 | FOSL2 | 3/15 | 0.282 | 3.931 | 11.273 | -0.281 | 0.385 |
| 3787 | JUNB | 3/15 | 0.282 | 3.931 | 11.273 | -1.178 | 1.569 |
| 3788 | PTTG1 | 3/15 | 0.282 | 3.931 | 11.273 | 0.436 | -0.471 |
| 3789 | CREBBP | 4/25 | 0.284 | 2.996 | 8.462 | -0.414 | 0.568 |
| 3790 | PGR | 4/25 | 0.284 | 2.996 | 8.462 | 0.585 | -0.733 |
| 3791 | MZF1 | 2/7 | 0.284 | 6.287 | 17.525 | -0.305 | -0.02 |
| 3792 | JUN | 14/149 | 0.289 | 1.635 | 4.511 | -0.123 | -0.304 |
| 3793 | PITX1 | 3/16 | 0.299 | 3.628 | 9.811 | -0.714 | -0.633 |
| 3794 | TP53 | 15/164 | 0.299 | 1.587 | 4.277 | -0.148 | -0.289 |
| 3795 | NR3C1 | 5/38 | 0.32 | 2.384 | 6.186 | -0.856 | 0.86 |
| 3796 | NFIC | 4/27 | 0.32 | 2.735 | 7.084 | -0.672 | 0.237 |
| 3797 | MYC | 10/100 | 0.32 | 1.75 | 4.502 | 0.191 | -0.21 |
| 3798 | EZH2 | 5/40 | 0.346 | 2.247 | 5.439 | -0.565 | 0.672 |
| 3799 | KLF4 | 5/40 | 0.346 | 2.247 | 5.439 | 0.641 | -0.731 |
| 3800 | DDIT3 | 3/18 | 0.346 | 3.144 | 7.594 | 1.288 | -1.654 |
| 3801 | YY1 | 9/91 | 0.346 | 1.728 | 4.077 | -0.347 | -0.063 |
| 3802 | DNMT3A | 2/9 | 0.346 | 4.49 | 10.451 | 0.107 | 1.489 |
| 3803 | FOSL1 | 2/9 | 0.346 | 4.49 | 10.451 | -1.002 | 1.208 |
| 3804 | SMARCA4 | 2/9 | 0.346 | 4.49 | 10.451 | -1.147 | 1.53 |
| 3805 | ZNF148 | 2/9 | 0.346 | 4.49 | 10.451 | 0.849 | -1.342 |
| 3806 | PPARG | 7/66 | 0.346 | 1.867 | 4.317 | -0.084 | 0.033 |
| 3807 | ETS1 | 8/79 | 0.346 | 1.774 | 4.085 | -0.674 | 0.764 |
| 3808 | ESR2 | 3/19 | 0.346 | 2.948 | 6.741 | 0.116 | -0.454 |
| 3809 | RUNX2 | 3/19 | 0.346 | 2.948 | 6.741 | -1.637 | -1.225 |
| 3810 | NFYA | 3/20 | 0.374 | 2.774 | 6.013 | 0.884 | -1.182 |
| 3811 | EPAS1 | 2/10 | 0.374 | 3.929 | 8.422 | 0.055 | 0.064 |
| 3812 | HOXB7 | 2/10 | 0.374 | 3.929 | 8.422 | 0.16 | -0.649 |
| 3813 | NKX3-1 | 2/10 | 0.374 | 3.929 | 8.422 | -1.403 | -1.533 |
| 3814 | TAL1 | 2/10 | 0.374 | 3.929 | 8.422 | -0.692 | 0.743 |
| 3815 | CDX2 | 4/32 | 0.384 | 2.246 | 4.723 | -0.667 | 0.758 |
| 3816 | ETS2 | 4/32 | 0.384 | 2.246 | 4.723 | -0.435 | 0.599 |
| 3817 |  |  |  |  |  |  |  |
| 3818 |  |  |  |  |  |  |  |
| 3819 | **HNF4A** | 5/45 | 0.401 | 1.966 | 4.009 | -1.392 | 1.622 |
| 3820 | **MEN1** | 2/11 | 0.401 | 3.492 | 6.921 | 1.151 | -1.346 |
| 3821 | **PITX2** | 2/11 | 0.401 | 3.492 | 6.921 | -0.851 | 0.523 |
| 3822 | **STAT2** | 2/11 | 0.401 | 3.492 | 6.921 | 0.085 | -0.262 |
| 3823 | **SP3** | 10/113 | 0.404 | 1.528 | 3.008 | -0.781 | 0.814 |
| 3824 | **CTNNB1** | 3/22 | 0.407 | 2.482 | 4.847 | -0.67 | -1.009 |
| 3825 | **USF2** | 5/47 | 0.418 | 1.872 | 3.569 | -0.512 | 0.324 |
| 3826 | **CEBPB** | 6/60 | 0.418 | 1.748 | 3.331 | 0.745 | -1.017 |
| 3827 | **APC** | 2/12 | 0.435 | 3.142 | 5.777 | 0.748 | -0.936 |
| 3828 | **OTX2** | 2/12 | 0.435 | 3.142 | 5.777 | 1.001 | -0.975 |
| 3829 | **FLI1** | 3/24 | 0.453 | 2.245 | 3.963 | 0.321 | -0.389 |
| 3830 | **HDAC3** | 3/24 | 0.453 | 2.245 | 3.963 | 0.833 | -0.844 |
| 3831 | **MTA1** | 3/24 | 0.453 | 2.245 | 3.963 | 0.451 | -0.929 |
| 3832 | **KAT2B** | 2/13 | 0.469 | 2.857 | 4.884 | -1.589 | 0.126 |
| 3833 | **PARP1** | 3/25 | 0.48 | 2.143 | 3.6 | 0.027 | 0.134 |
| 3834 | **RELA** | 22/301 | 0.499 | 1.242 | 2.031 | -0.448 | 0.02 |
| 3835 | **MITF** | 3/26 | 0.499 | 2.05 | 3.279 | 1.293 | -0.607 |
| 3836 | **NFKB1** | 22/303 | 0.499 | 1.233 | 1.964 | -0.545 | 0.215 |
| 3837 | **HDAC2** | 3/27 | 0.531 | 1.964 | 2.994 | 0.142 | -0.305 |
| 3838 | **RFX1** | 2/15 | 0.534 | 2.417 | 3.598 | -1.425 | 1.406 |
| 3839 | **TBP** | 2/15 | 0.534 | 2.417 | 3.598 | -1.106 | 1.384 |
| 3840 | **TP63** | 2/15 | 0.534 | 2.417 | 3.598 | -0.429 | 0.443 |
| 3841 | **VDR** | 4/42 | 0.566 | 1.654 | 2.345 | -0.537 | 0.446 |
| 3842 | **LEF1** | 3/29 | 0.575 | 1.813 | 2.511 | -0.626 | -0.948 |
| 3843 | **GATA1** | 5/57 | 0.581 | 1.511 | 2.066 | 0.226 | -0.075 |
| 3844 | **YBX1** | 3/30 | 0.581 | 1.746 | 2.307 | -1.84 | -0.098 |
| 3845 | **BCL6** | 2/17 | 0.581 | 2.094 | 2.734 | -1.261 | 1.306 |
| 3846 | **FOXM1** | 2/17 | 0.581 | 2.094 | 2.734 | -1.403 | -1.533 |
| 3847 | **RB1** | 3/31 | 0.581 | 1.683 | 2.123 | 0.614 | -0.796 |
| 3848 | **HDAC9** | 2/18 | 0.581 | 1.963 | 2.405 | 0.071 | 0.19 |
| 3849 | **FOXQ1** | 1/6 | 0.581 | 3.141 | 3.681 | 1.284 | -1.802 |
| 3850 | **GFI1** | 1/6 | 0.581 | 3.141 | 3.681 | -1.507 | 1.805 |
| 3851 | **IKBKB** | 1/6 | 0.581 | 3.141 | 3.681 | 0.846 | -0.952 |
| 3852 | **MBD1** | 1/6 | 0.581 | 3.141 | 3.681 | -0.739 | 0.543 |
| 3853 | **MEF2D** | 1/6 | 0.581 | 3.141 | 3.681 | 1.013 | -0.71 |
| 3854 | **NFKB2** | 1/6 | 0.581 | 3.141 | 3.681 | -0.717 | 0.564 |
| 3855 | **PIAS1** | 1/6 | 0.581 | 3.141 | 3.681 | -0.676 | 0.429 |
| 3856 | **SFPQ** | 1/6 | 0.581 | 3.141 | 3.681 | 0.97 | -0.439 |
| 3857 | **SOX6** | 1/6 | 0.581 | 3.141 | 3.681 | -1.507 | 1.805 |
| 3858 | **SPI1** | 5/62 | 0.581 | 1.378 | 1.599 | -0.13 | 0.282 |
| 3859 | **GATA2** | 2/19 | 0.581 | 1.848 | 2.126 | -3.023 | 0.793 |
| 3860 | **NFKBIA** | 2/19 | 0.581 | 1.848 | 2.126 | 0.09 | -0.139 |
| 3861 |  |  |  |  |  |  |  |
| 3862 |  |  |  |  |  |  |  |
| 3863 | HOXA1 | 1/7 | 0.591 | 2.617 | 2.739 | -0.602 | 0.416 |
| 3864 | HSF2 | 1/7 | 0.591 | 2.617 | 2.739 | -0.568 | 0.91 |
| 3865 | ILF3 | 1/7 | 0.591 | 2.617 | 2.739 | -0.601 | 0.534 |
| 3866 | ING1 | 1/7 | 0.591 | 2.617 | 2.739 | -1.507 | 1.805 |
| 3867 | KLF10 | 1/7 | 0.591 | 2.617 | 2.739 | -1.507 | 1.805 |
| 3868 | MSX2 | 1/7 | 0.591 | 2.617 | 2.739 | -0.787 | 1.254 |
| 3869 | MTF1 | 1/7 | 0.591 | 2.617 | 2.739 | -0.637 | 2.33 |
| 3870 | NCOR1 | 1/7 | 0.591 | 2.617 | 2.739 | -4.09 | -1.265 |
| 3871 | NFATC2 | 1/7 | 0.591 | 2.617 | 2.739 | 0.573 | -1.364 |
| 3872 | SALL4 | 1/7 | 0.591 | 2.617 | 2.739 | -0.983 | 1.378 |
| 3873 | TFAP4 | 1/7 | 0.591 | 2.617 | 2.739 | -1.507 | 1.805 |
| 3874 | PAX5 | 2/21 | 0.601 | 1.653 | 1.684 | -0.467 | 0.221 |
| 3875 | STAT6 | 3/36 | 0.601 | 1.428 | 1.432 | -0.16 | 0.184 |
| 3876 | MYB | 3/37 | 0.601 | 1.386 | 1.328 | -0.461 | 0.489 |
| 3877 | ATF5 | 1/8 | 0.601 | 2.243 | 2.112 | -0.619 | 0.826 |
| 3878 | CUX1 | 1/8 | 0.601 | 2.243 | 2.112 | -0.963 | 0.503 |
| 3879 | ERCC2 | 1/8 | 0.601 | 2.243 | 2.112 | -1.507 | 1.805 |
| 3880 | FOXF2 | 1/8 | 0.601 | 2.243 | 2.112 | -1.52 | 1.433 |
| 3881 | HES1 | 1/8 | 0.601 | 2.243 | 2.112 | -2.02 | 2.146 |
| 3882 | IRF9 | 1/8 | 0.601 | 2.243 | 2.112 | -1.507 | 1.805 |
| 3883 | KLF8 | 1/8 | 0.601 | 2.243 | 2.112 | 0.952 | -0.878 |
| 3884 | MEF2C | 1/8 | 0.601 | 2.243 | 2.112 | 1.284 | -1.802 |
| 3885 | MYBL2 | 1/8 | 0.601 | 2.243 | 2.112 | -0.903 | 0.701 |
| 3886 | NR0B2 | 1/8 | 0.601 | 2.243 | 2.112 | 1.815 | -1.774 |
| 3887 | RBMX | 1/8 | 0.601 | 2.243 | 2.112 | 1.73 | -1.328 |
| 3888 | ZFP36 | 1/8 | 0.601 | 2.243 | 2.112 | 1.284 | -1.802 |
| 3889 | E2F4 | 2/23 | 0.612 | 1.496 | 1.353 | -1.394 | -1.066 |
| 3890 | NANOG | 2/23 | 0.612 | 1.496 | 1.353 | -0.676 | 0.471 |
| 3891 | TFAP2A | 5/71 | 0.612 | 1.19 | 1.026 | -0.224 | 0.214 |
| 3892 | ELF1 | 1/9 | 0.612 | 1.963 | 1.672 | 1.15 | -0.898 |
| 3893 | EWSR1 | 1/9 | 0.612 | 1.963 | 1.672 | 0.573 | -1.364 |
| 3894 | GATA6 | 1/9 | 0.612 | 1.963 | 1.672 | -0.721 | 0.582 |
| 3895 | HIC1 | 1/9 | 0.612 | 1.963 | 1.672 | 1.284 | -1.802 |
| 3896 | POU3F2 | 1/9 | 0.612 | 1.963 | 1.672 | -0.768 | 0.85 |
| 3897 | PPARGC1A | 1/9 | 0.612 | 1.963 | 1.672 | 1.815 | -1.774 |
| 3898 | SP2 | 1/9 | 0.612 | 1.963 | 1.672 | -0.807 | 0.546 |
| 3899 | RUNX1 | 3/40 | 0.614 | 1.273 | 1.067 | 0.059 | -0.31 |
| 3900 | EP300 | 4/56 | 0.614 | 1.208 | 1.008 | -0.494 | 0.338 |
| 3901 | CREM | 2/25 | 0.616 | 1.365 | 1.101 | 0.064 | 0.048 |
| 3902 | SRF | 2/25 | 0.616 | 1.365 | 1.101 | -1.282 | 0.655 |
| 3903 | CREB1 | 6/90 | 0.616 | 1.122 | 0.884 | -1.215 | 0.482 |
| 3904 | ABL1 | 1/10 | 0.616 | 1.744 | 1.351 | -1.507 | 1.805 |

| 3905 | TF | Overlap | FDR | Odds Ratio | Combined Score | Mean log2-fold change | Mean log2-fold change |
| --- | --- | --- | --- | --- | --- | --- | --- |
| 3906 |  |  |  |  |  | HAP40 | HTT |
| 3907 | LMO2 | 1/10 | 0.616 | 1.744 | 1.351 | -1.957 | 2.851 |
| 3908 | MBD2 | 1/10 | 0.616 | 1.744 | 1.351 | -0.746 | 0.893 |
| 3909 | PPARD | 1/10 | 0.616 | 1.744 | 1.351 | 1.317 | -1.025 |
| 3910 | TFAP2C | 1/10 | 0.616 | 1.744 | 1.351 | -1.507 | 1.805 |
| 3911 | TFDP1 | 1/10 | 0.616 | 1.744 | 1.351 | -0.721 | 0.582 |
| 3912 | ZNF382 | 1/10 | 0.616 | 1.744 | 1.351 | -0.717 | 0.564 |
| 3913 | HNF1A | 2/26 | 0.617 | 1.308 | 0.996 | 0.105 | -0.099 |
| 3914 | NR1I2 | 2/26 | 0.617 | 1.308 | 0.996 | -0.842 | 0.973 |
| 3915 | FOXP3 | 1/11 | 0.622 | 1.57 | 1.11 | -1.507 | 1.805 |
| 3916 | KLF2 | 1/11 | 0.622 | 1.57 | 1.11 | -1.507 | 1.805 |
| 3917 | MECP2 | 1/11 | 0.622 | 1.57 | 1.11 | -0.739 | 0.543 |
| 3918 | PAX6 | 1/11 | 0.622 | 1.57 | 1.11 | -0.79 | 0.623 |
| 3919 | PROX1 | 1/11 | 0.622 | 1.57 | 1.11 | 1.147 | -0.676 |
| 3920 | SP4 | 1/11 | 0.622 | 1.57 | 1.11 | 2.142 | -1.855 |
| 3921 | CREB5 | 1/12 | 0.646 | 1.427 | 0.923 | -0.676 | 0.429 |
| 3922 | IRF7 | 1/12 | 0.646 | 1.427 | 0.923 | 0.809 | -0.979 |
| 3923 | NR5A2 | 1/12 | 0.646 | 1.427 | 0.923 | -1.743 | 1.607 |
| 3924 | TCF7L2 | 1/12 | 0.646 | 1.427 | 0.923 | -0.739 | 0.543 |
| 3925 | CTCF | 2/30 | 0.657 | 1.121 | 0.683 | 0.648 | -1.037 |
| 3926 | RUNX3 | 2/30 | 0.657 | 1.121 | 0.683 | -1.276 | 1.412 |
| 3927 | USF1 | 4/65 | 0.657 | 1.029 | 0.613 | -0.604 | -0.224 |
| 3928 | HDAC7 | 1/13 | 0.657 | 1.308 | 0.777 | -1.507 | 1.805 |
| 3929 | KLF5 | 1/13 | 0.657 | 1.308 | 0.777 | -1.015 | 0.807 |
| 3930 | NFYC | 1/13 | 0.657 | 1.308 | 0.777 | -0.739 | 0.543 |
| 3931 | STAT5A | 1/13 | 0.657 | 1.308 | 0.777 | -1.015 | 0.807 |
| 3932 | STAT1 | 5/84 | 0.671 | 0.993 | 0.557 | -0.795 | 0.363 |
| 3933 | NF1 | 1/14 | 0.671 | 1.207 | 0.66 | -0.746 | 0.893 |
| 3934 | NFYB | 1/14 | 0.671 | 1.207 | 0.66 | -0.739 | 0.543 |
| 3935 | SNAI2 | 1/14 | 0.671 | 1.207 | 0.66 | -0.633 | 0.532 |
| 3936 | CEBPA | 3/51 | 0.68 | 0.981 | 0.507 | 0.834 | -0.481 |
| 3937 | ING4 | 1/15 | 0.68 | 1.121 | 0.565 | -1.507 | 1.805 |
| 3938 | IRF3 | 1/15 | 0.68 | 1.121 | 0.565 | -1.67 | -1.554 |
| 3939 | NR4A1 | 1/15 | 0.68 | 1.121 | 0.565 | -1.092 | 1.049 |
| 3940 | NRF1 | 1/15 | 0.68 | 1.121 | 0.565 | -0.911 | 1.381 |
| 3941 | SATB1 | 1/15 | 0.68 | 1.121 | 0.565 | -0.76 | 1.934 |
| 3942 | NFE2L2 | 1/16 | 0.702 | 1.046 | 0.487 | -0.911 | 1.381 |
| 3943 | TWIST1 | 2/35 | 0.702 | 0.951 | 0.442 | 0.281 | -0.923 |
| 3944 | CEBPD | 1/17 | 0.714 | 0.981 | 0.422 | 2.065 | -2.392 |
| 3945 | GATA3 | 2/38 | 0.735 | 0.872 | 0.345 | 0.11 | 0.95 |
| 3946 | POU2F1 | 2/38 | 0.735 | 0.872 | 0.345 | -1.689 | -0.973 |
| 3947 | PML | 1/19 | 0.751 | 0.872 | 0.322 | -1.507 | 1.805 |
| 3948 | ARNT | 1/20 | 0.756 | 0.826 | 0.283 | 1.284 | -1.802 |

| 3949 | TF | Overlap | FDR | Odds Ratio | Combined Score | Mean log2-fold change | Mean log2-fold change |
| --- | --- | --- | --- | --- | --- | --- | --- |
| 3950 |  |  |  |  |  | HAP40 | HTT |
| 3951 | NR1H4 | 1/20 | 0.756 | 0.826 | 0.283 | 1.284 | -1.802 |
| 3952 | FOXA1 | 1/21 | 0.773 | 0.785 | 0.25 | -2.351 | 2.527 |
| 3953 | AHR | 1/22 | 0.774 | 0.747 | 0.222 | -0.787 | 1.254 |
| 3954 | NR5A1 | 1/22 | 0.774 | 0.747 | 0.222 | 1.601 | -1.613 |
| 3955 | REL | 1/22 | 0.774 | 0.747 | 0.222 | -1.507 | 1.805 |
| 3956 | SOX2 | 1/22 | 0.774 | 0.747 | 0.222 | 0.952 | -0.878 |
| 3957 | POU5F1 | 1/24 | 0.799 | 0.682 | 0.176 | -1.507 | 1.805 |
| 3958 | TWIST2 | 1/25 | 0.809 | 0.654 | 0.157 | 1.269 | -2.27 |
| 3959 | RARA | 1/26 | 0.818 | 0.627 | 0.14 | -1.507 | 1.805 |
| 3960 | ERG | 1/27 | 0.828 | 0.603 | 0.126 | 0.952 | -0.878 |
| 3961 | PPARA | 1/39 | 0.915 | 0.413 | 0.039 | -1.535 | 1.619 |
| 3962 | IRF1 | 1/51 | 0.96 | 0.313 | 0.014 | -1.507 | 1.805 |

**Supplementary Table 8:**
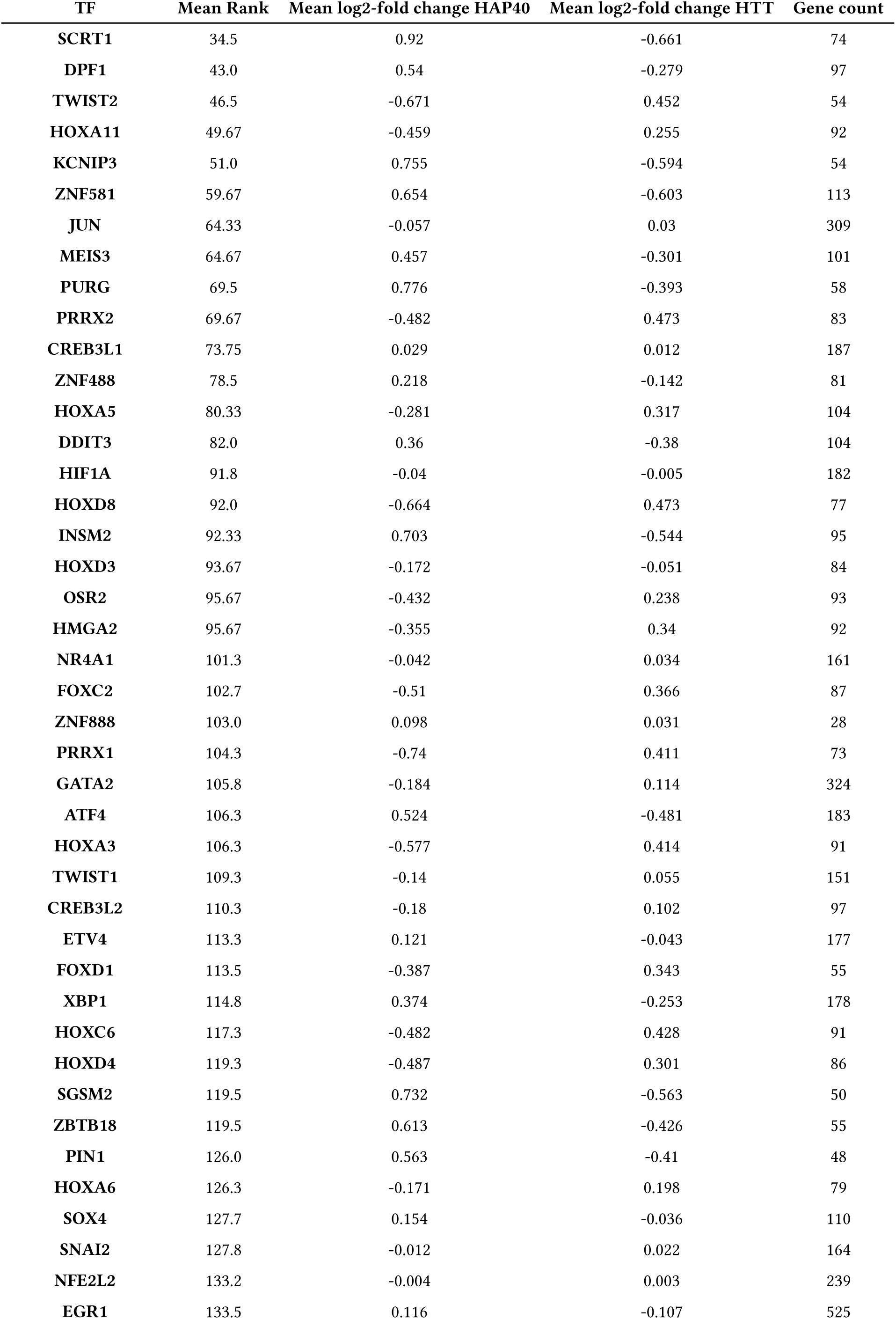
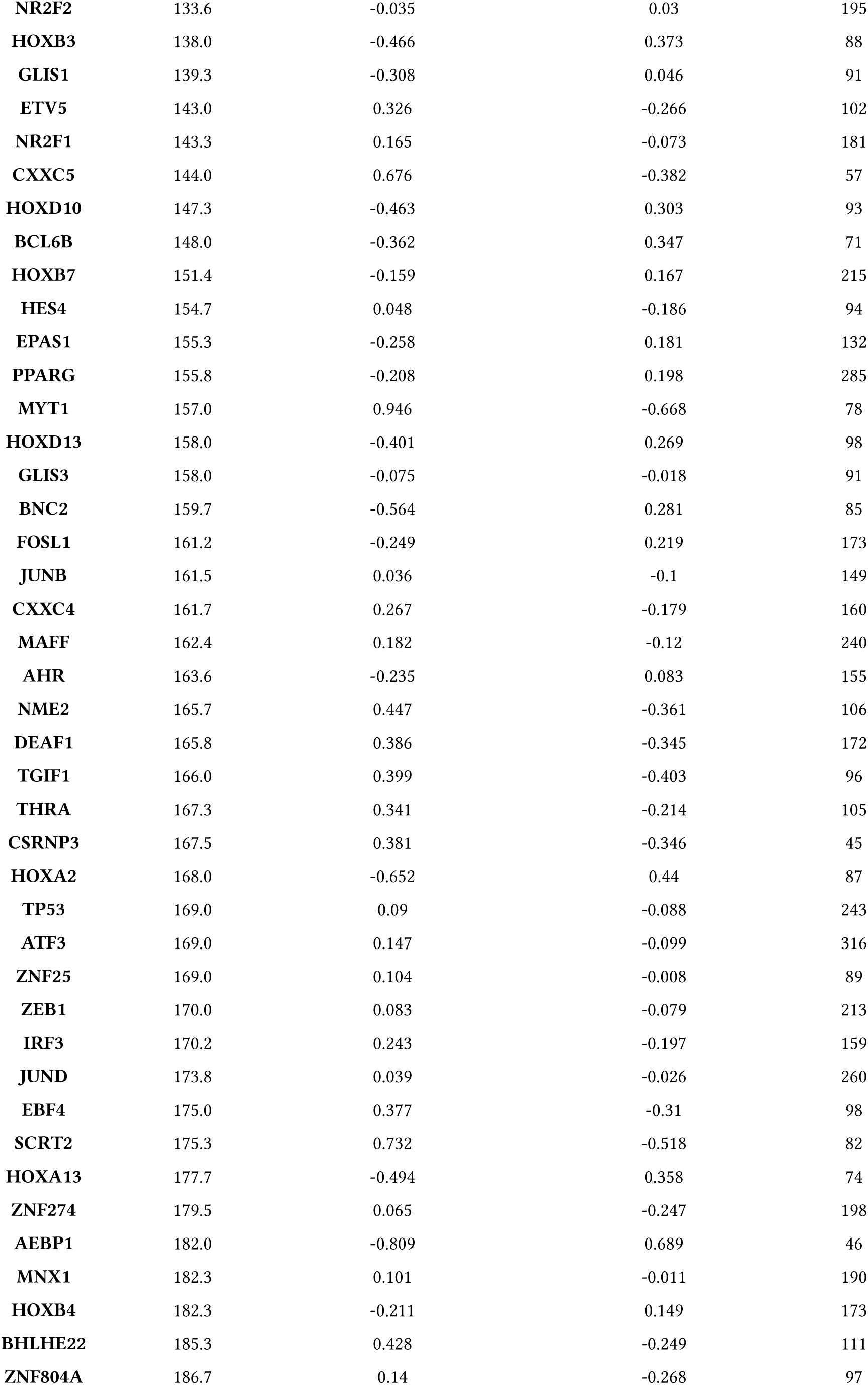
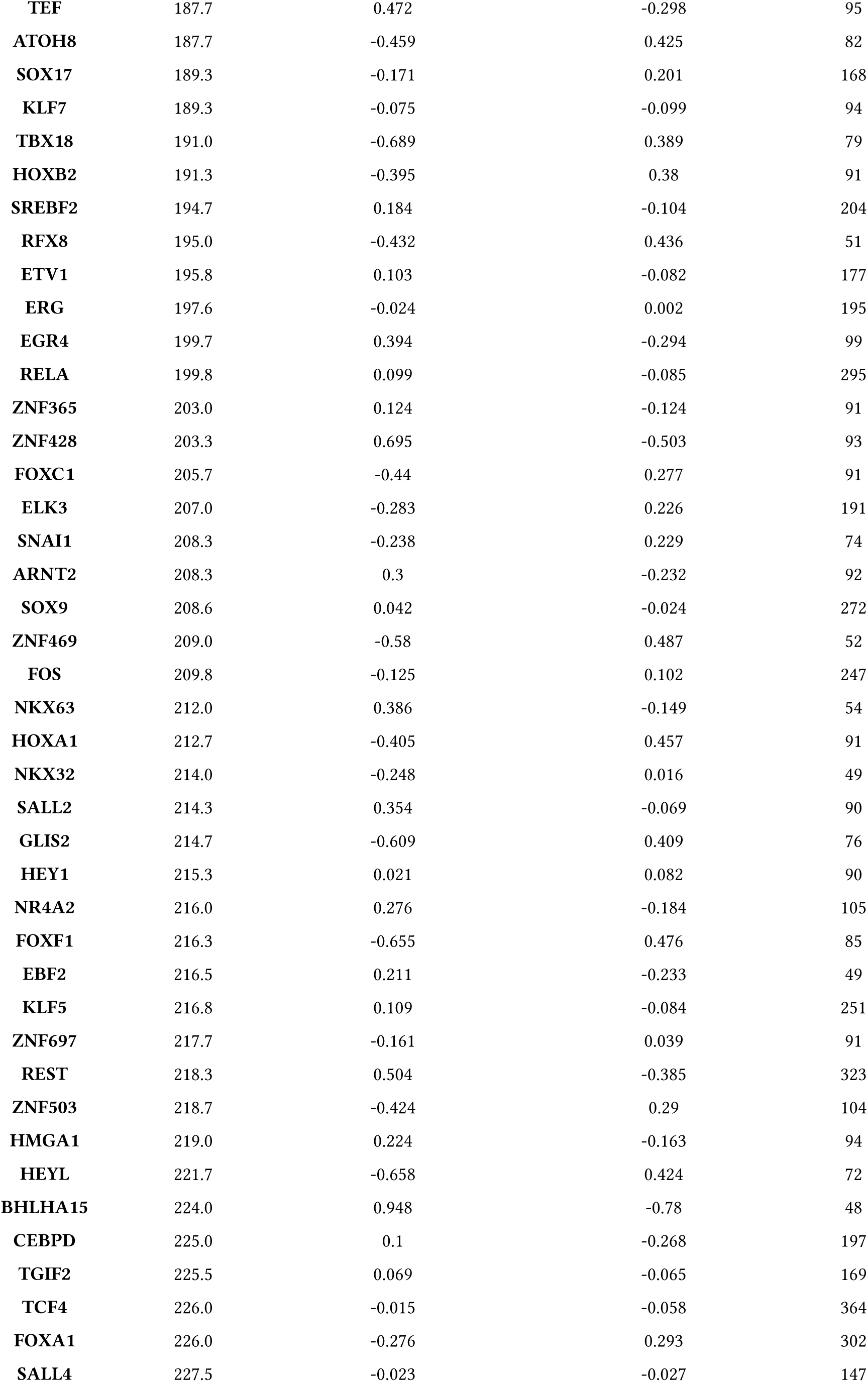
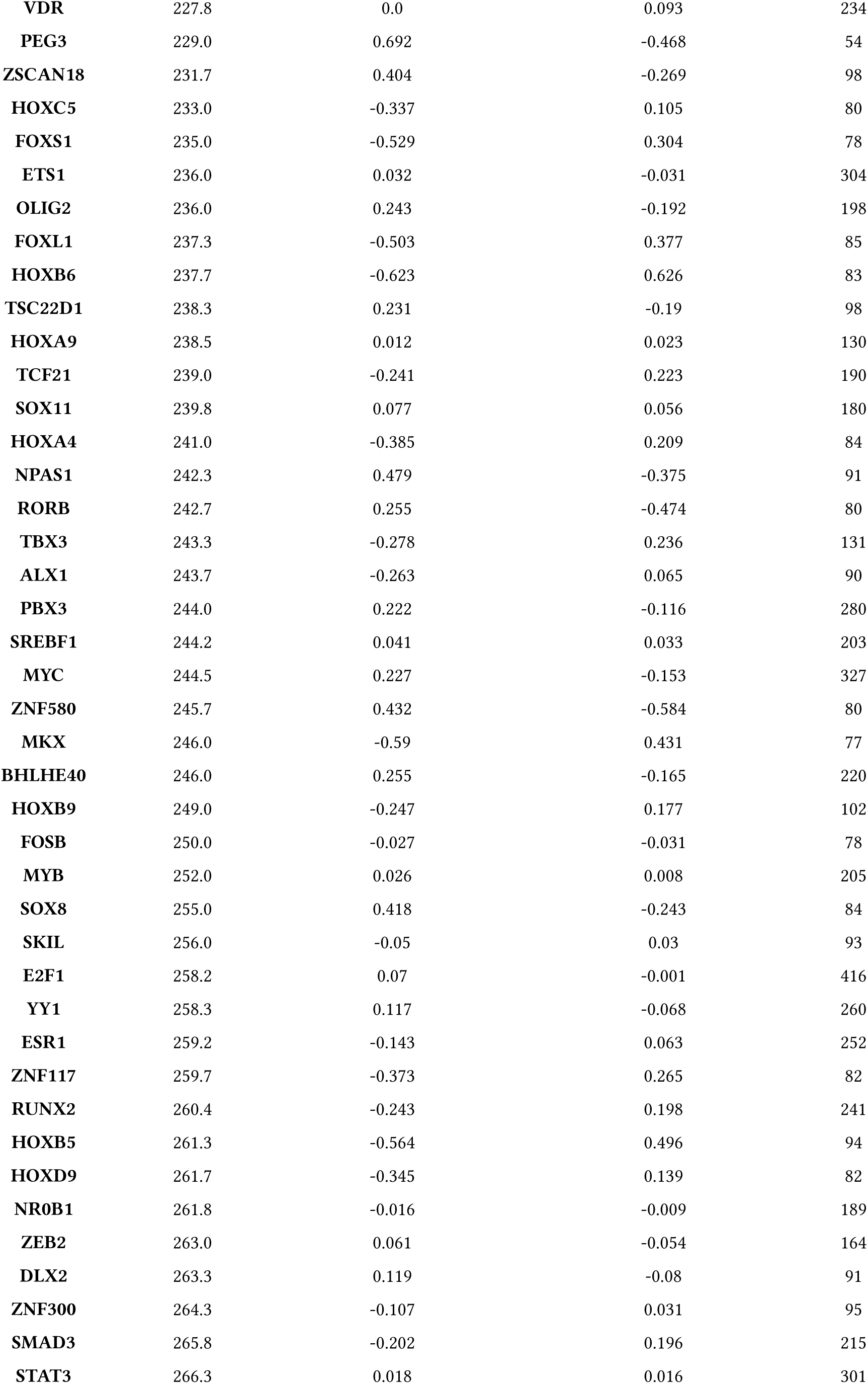
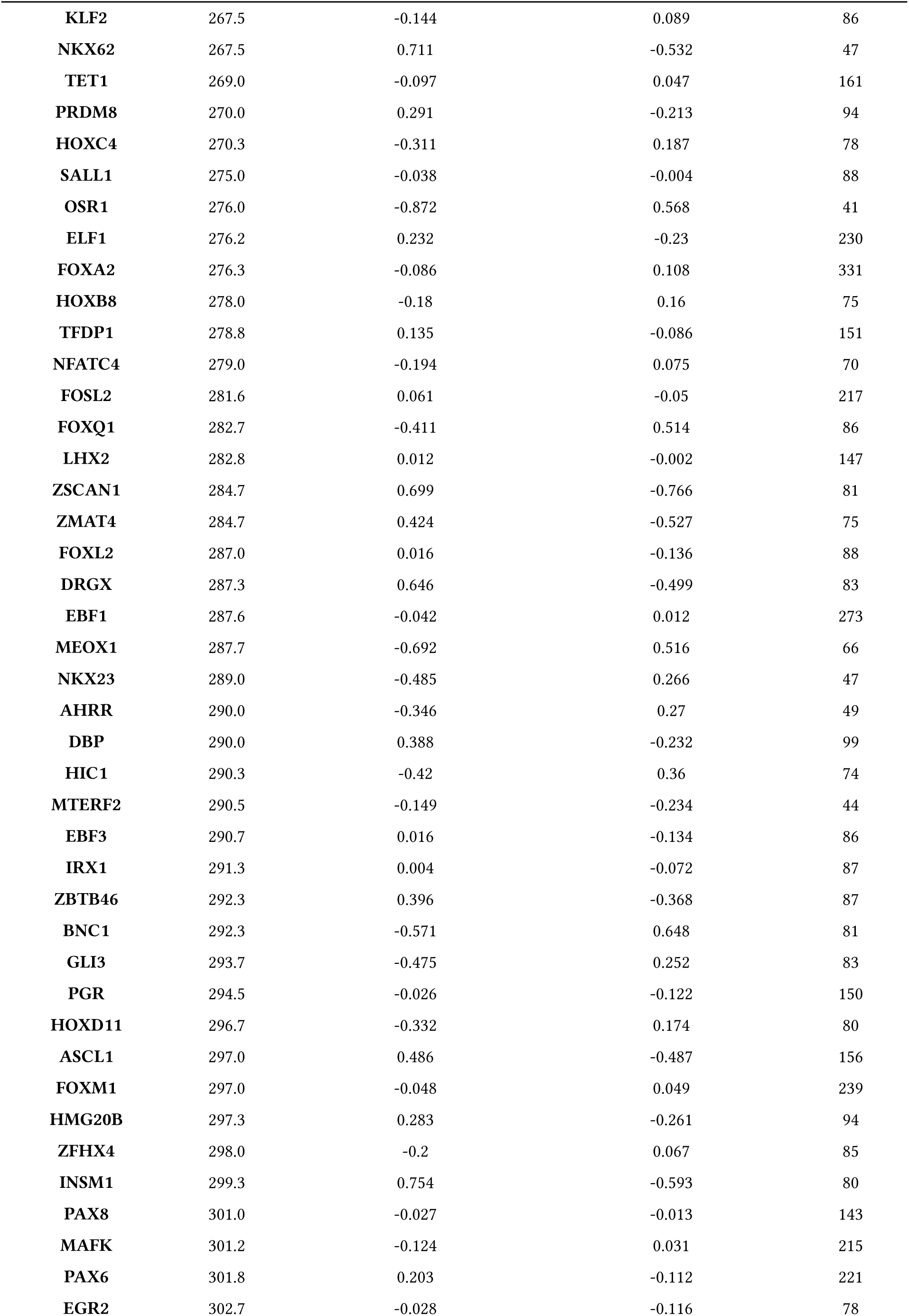
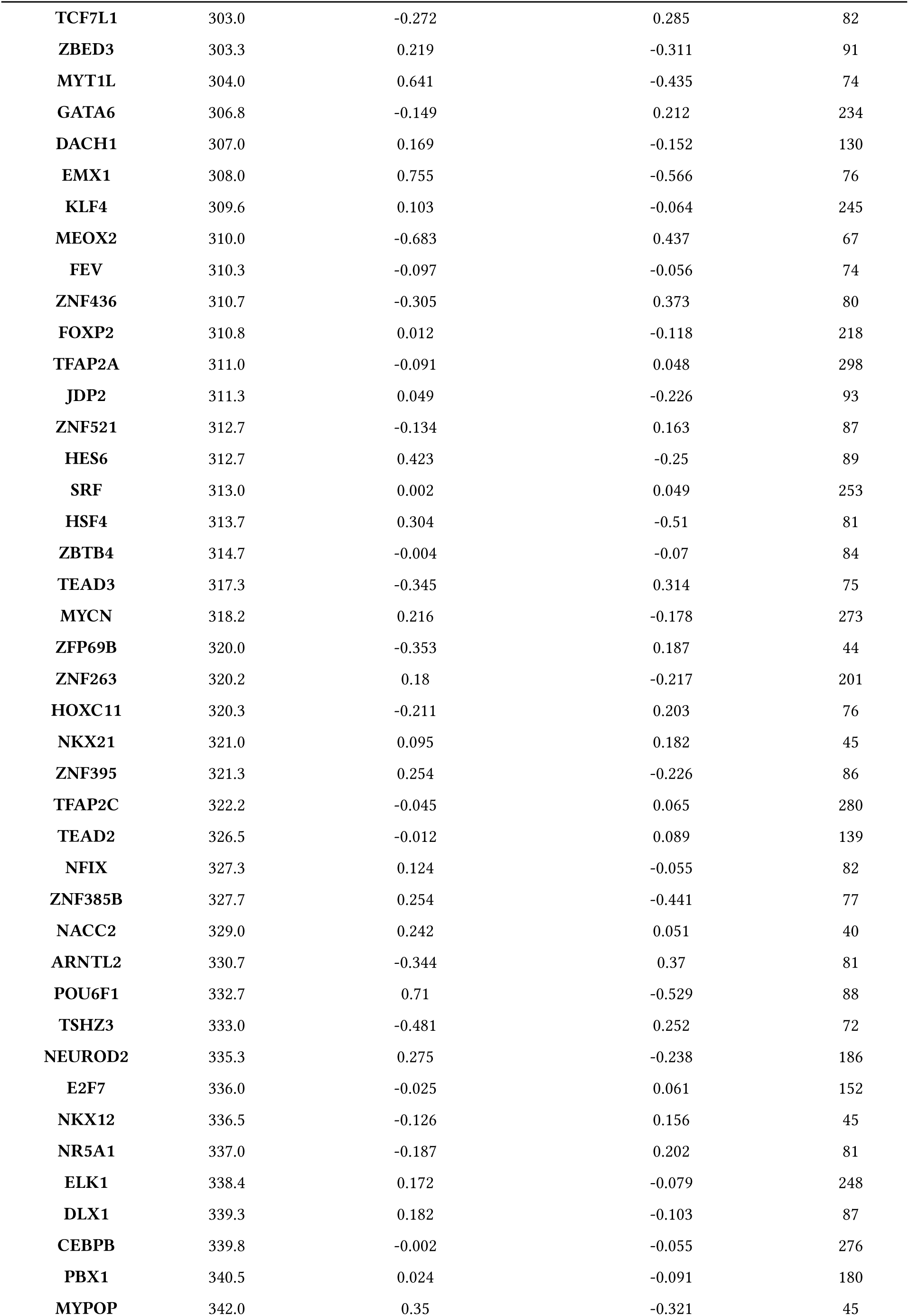
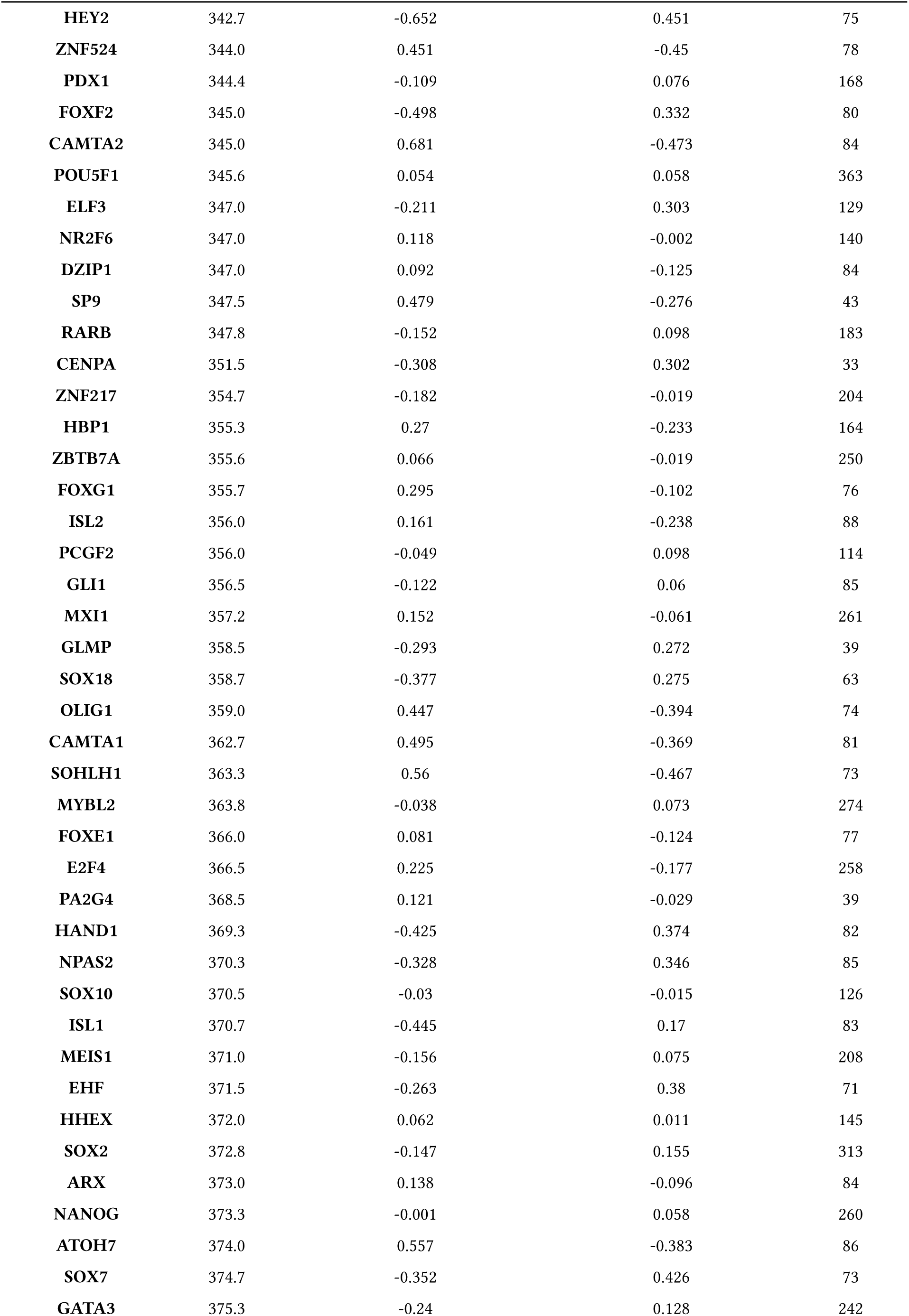
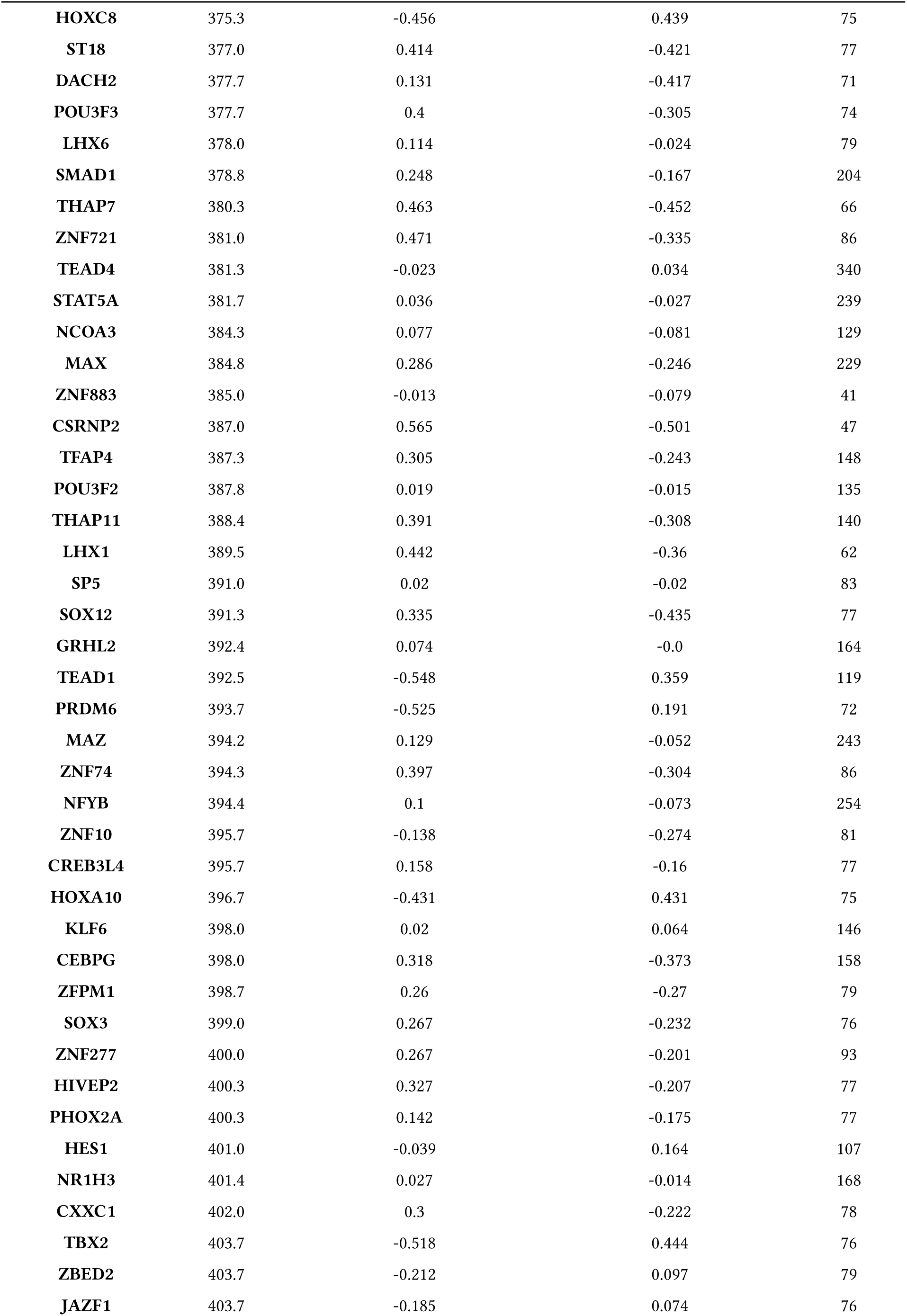
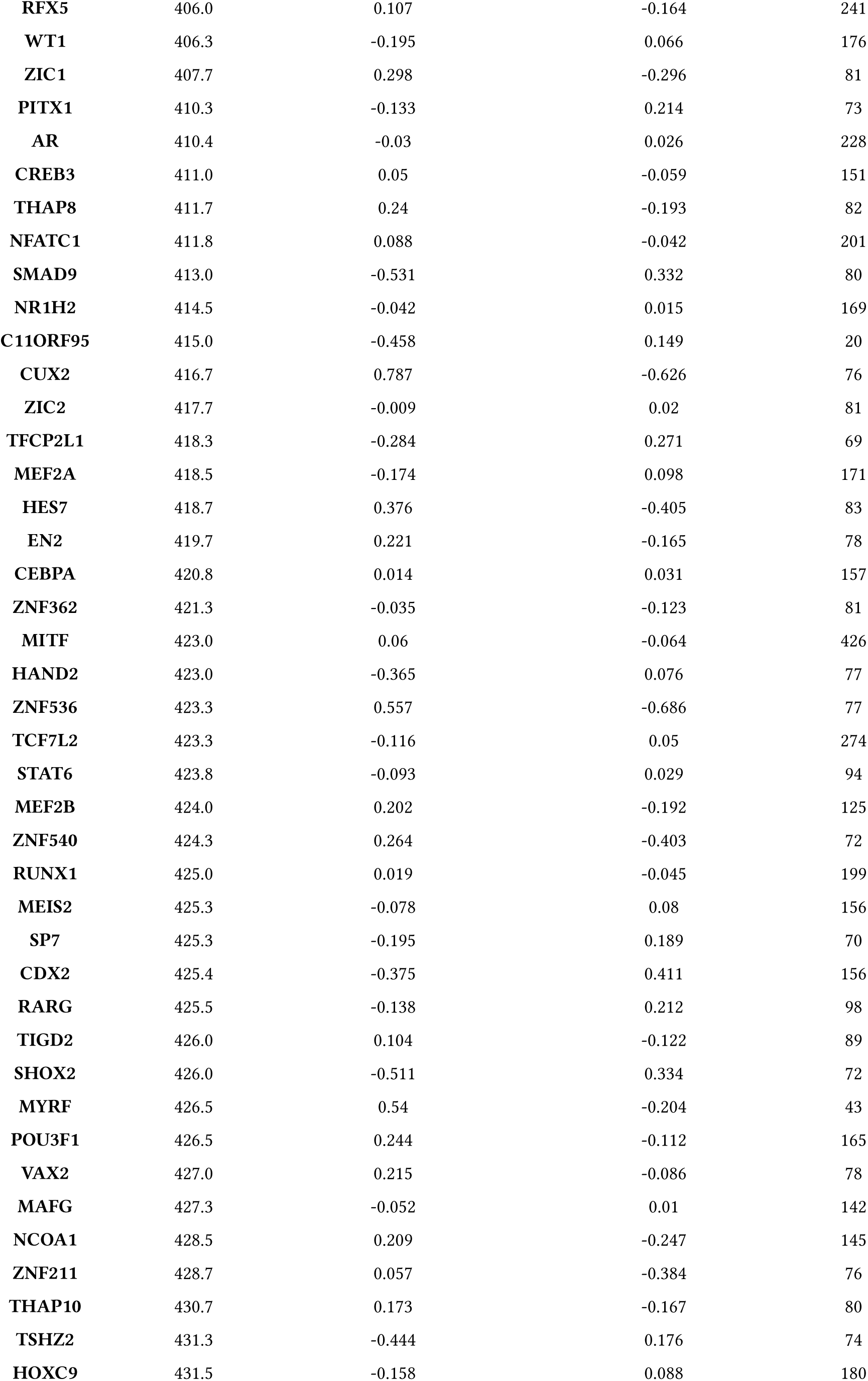
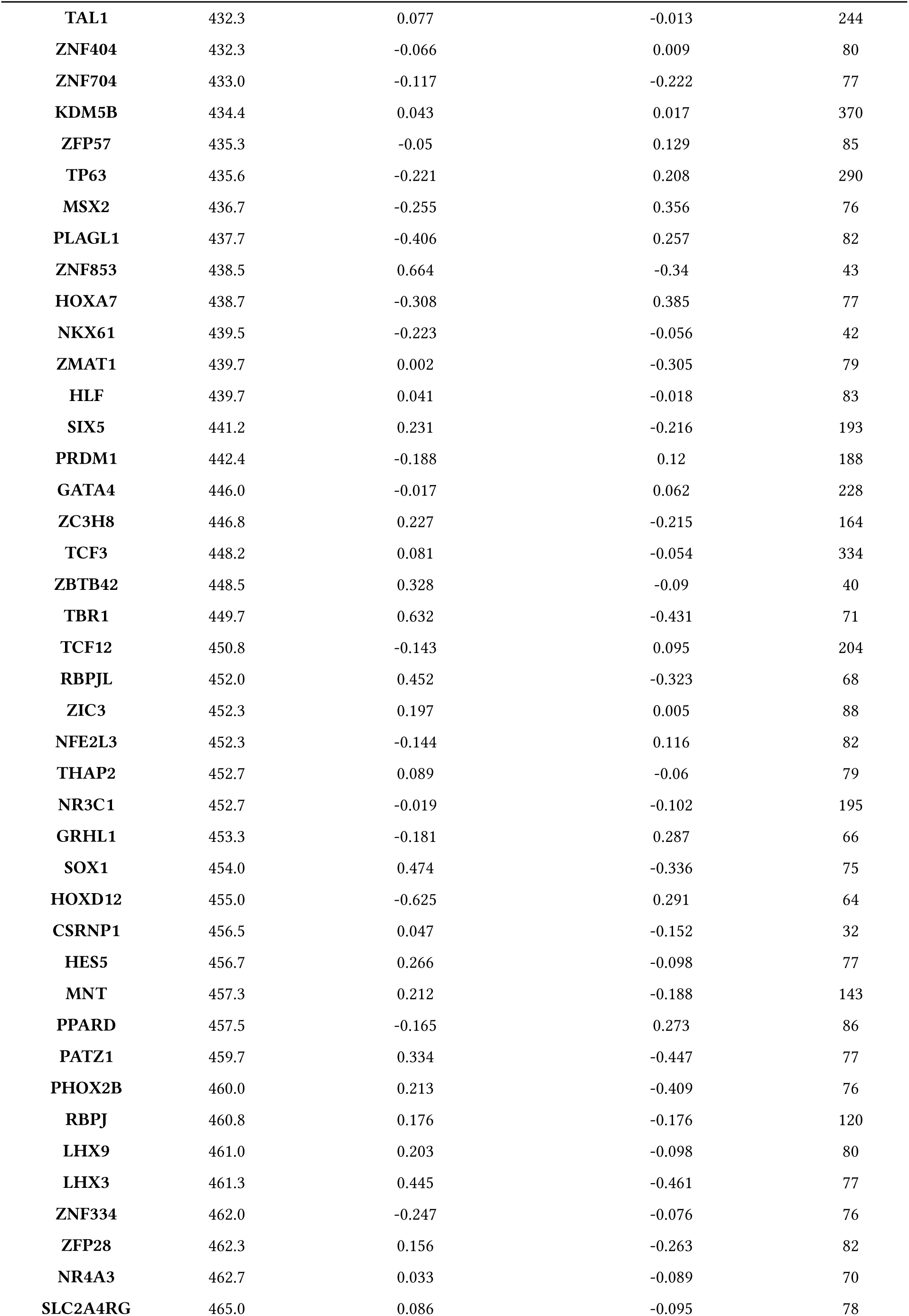
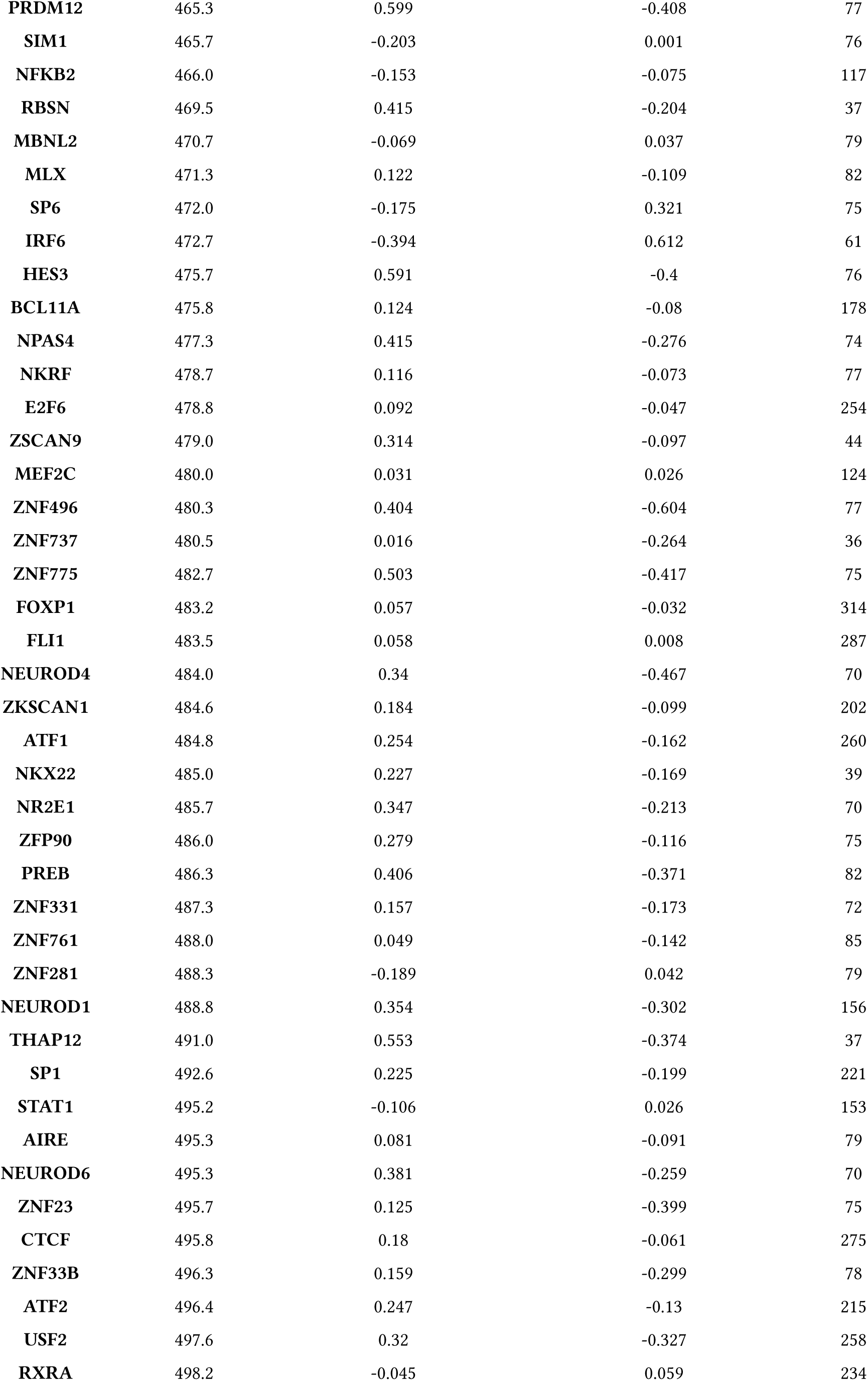
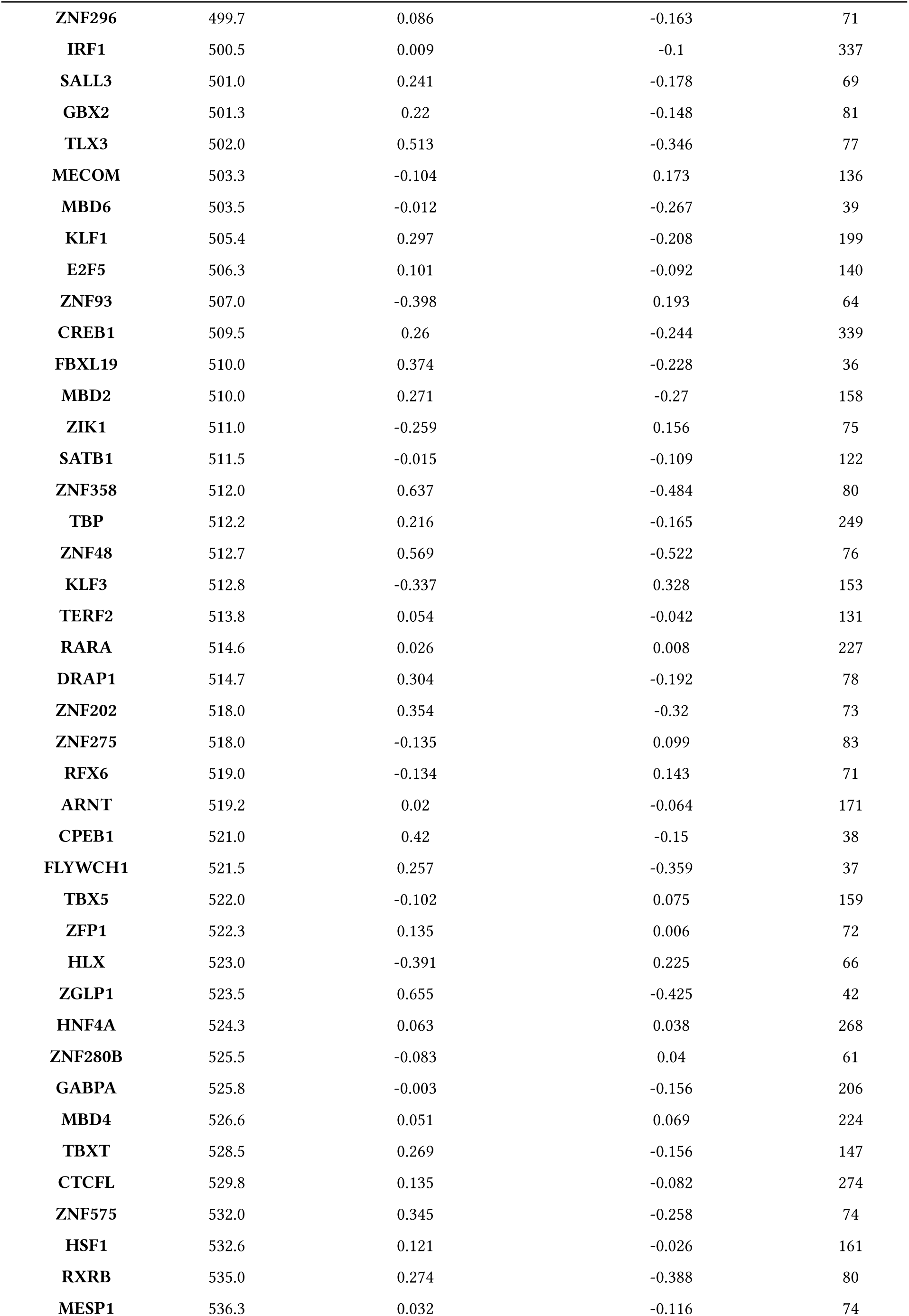
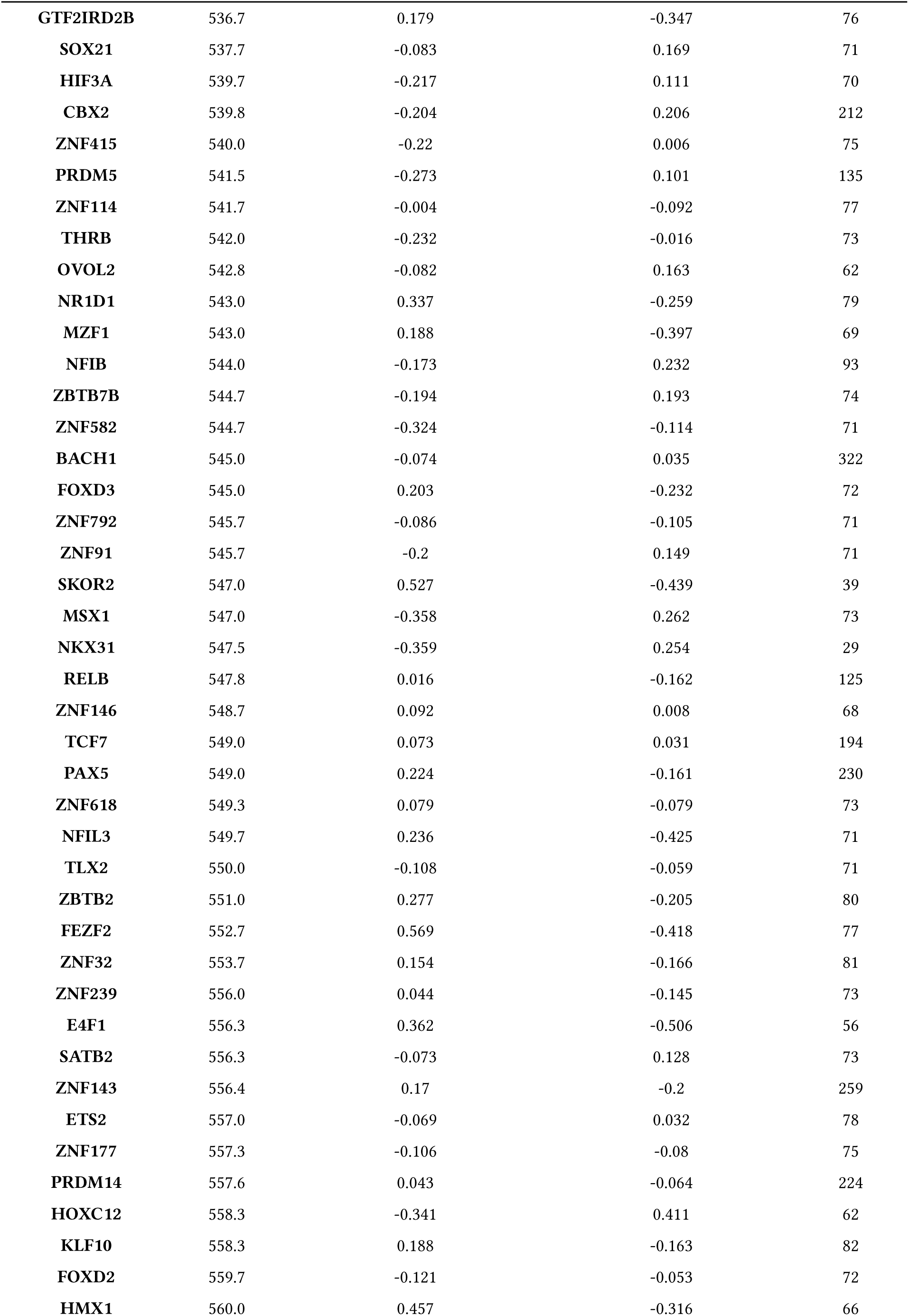
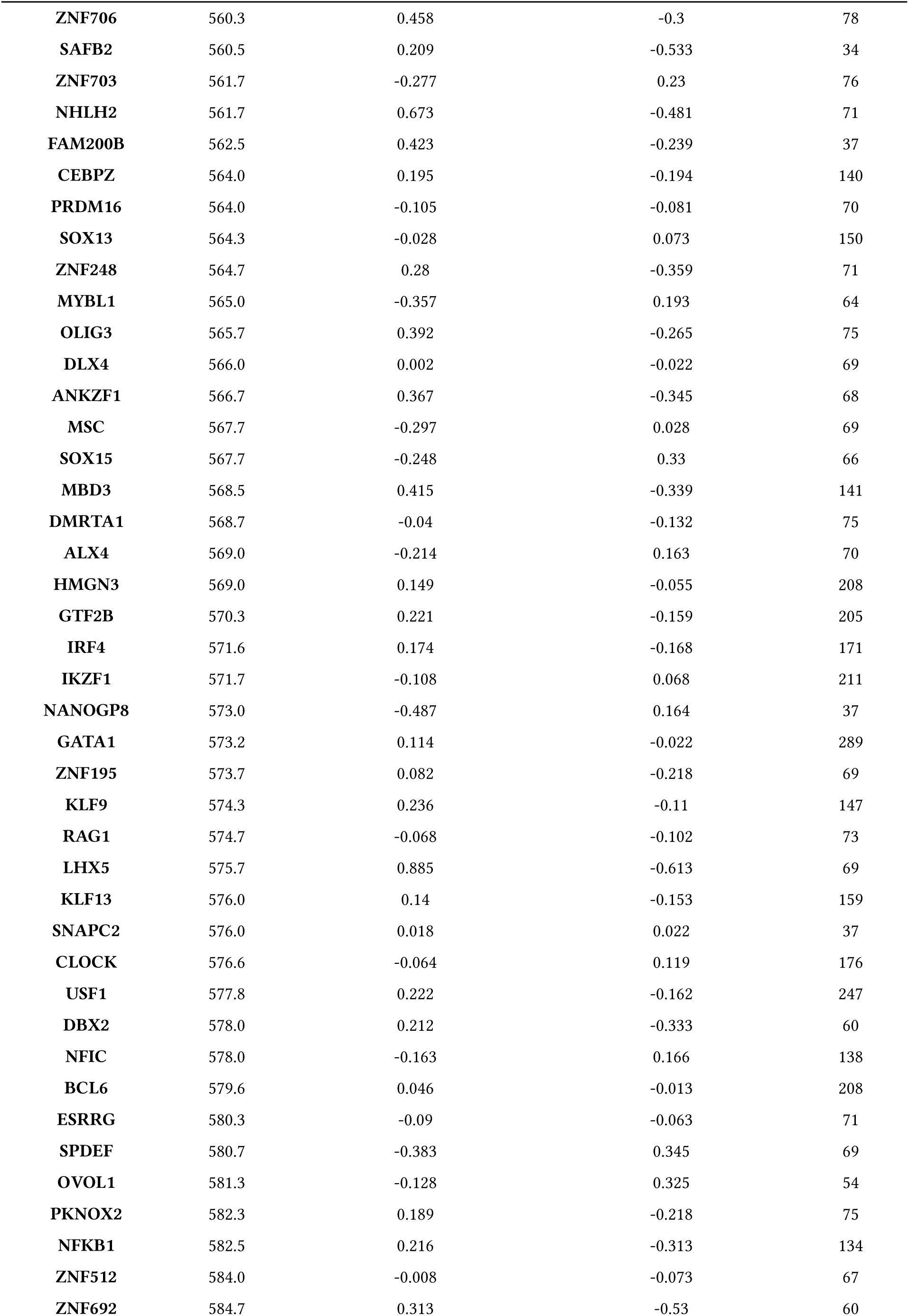
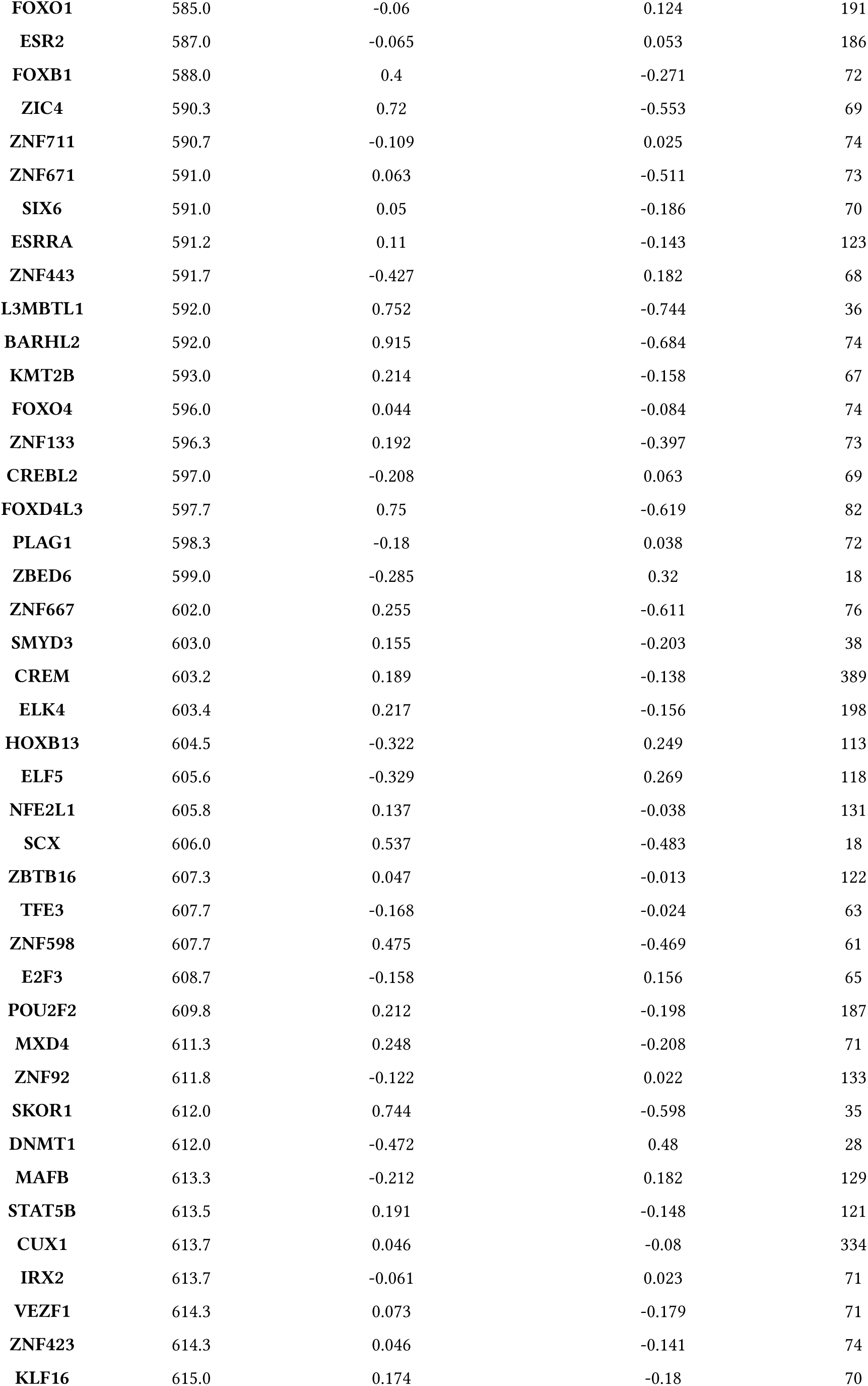
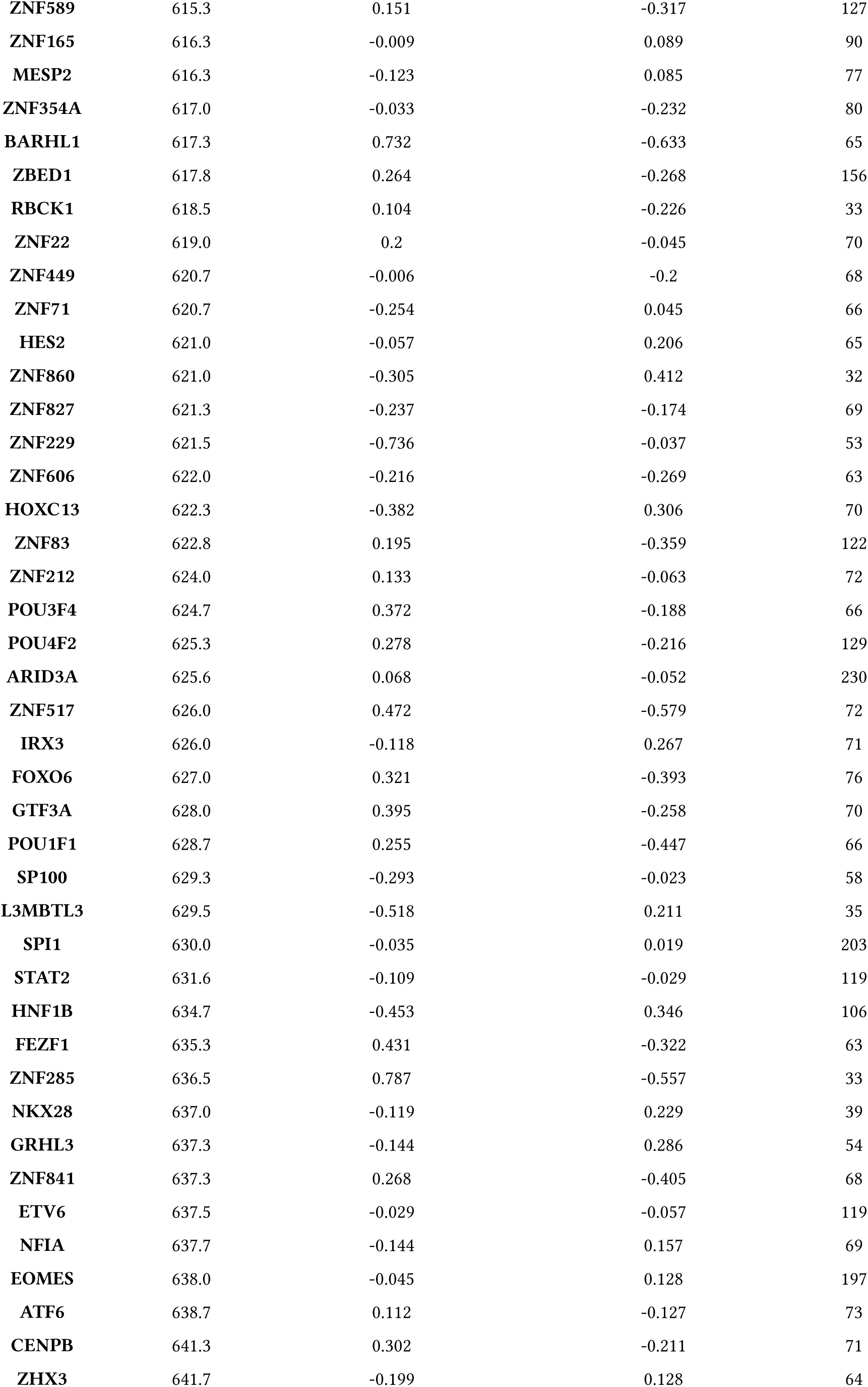
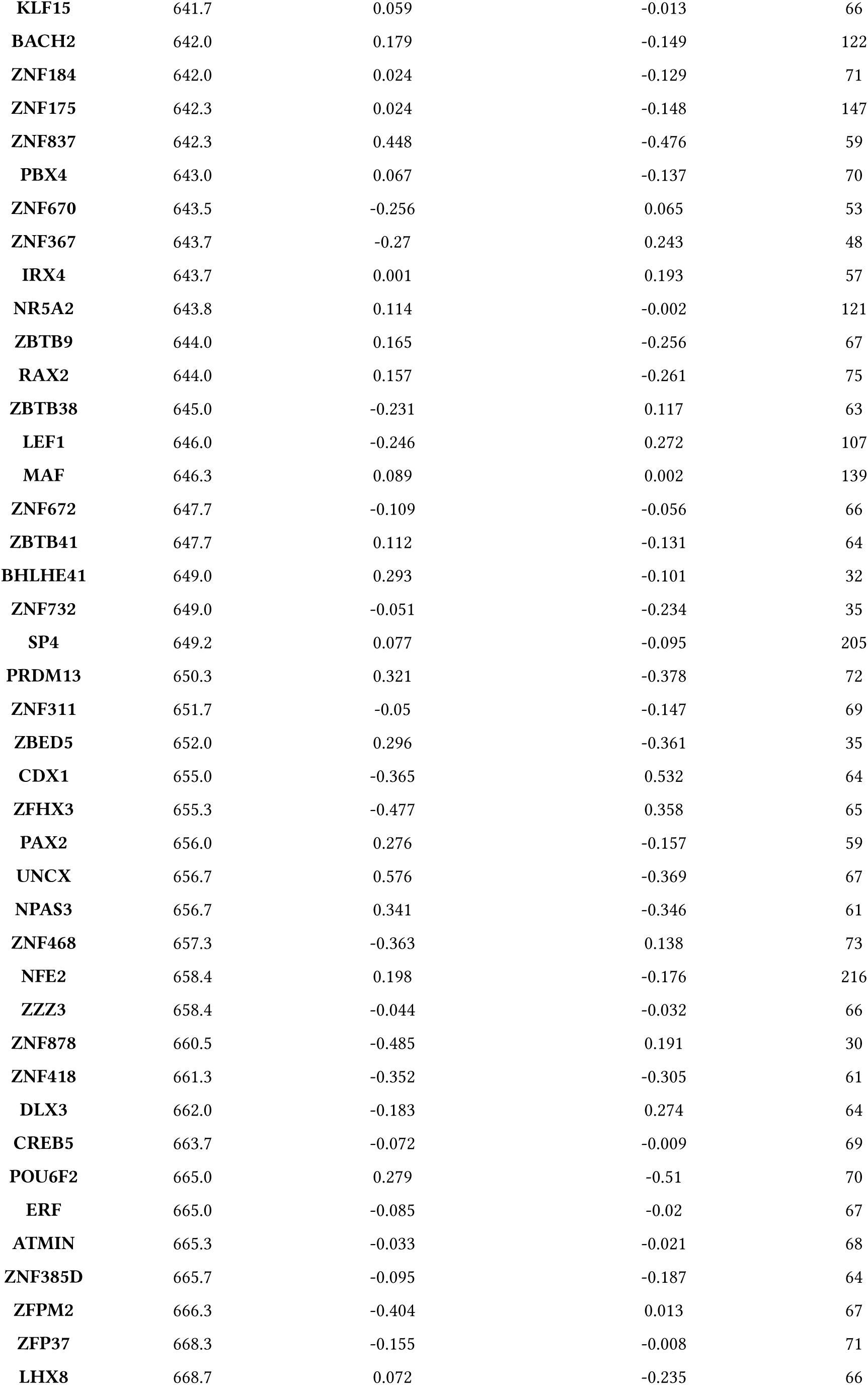
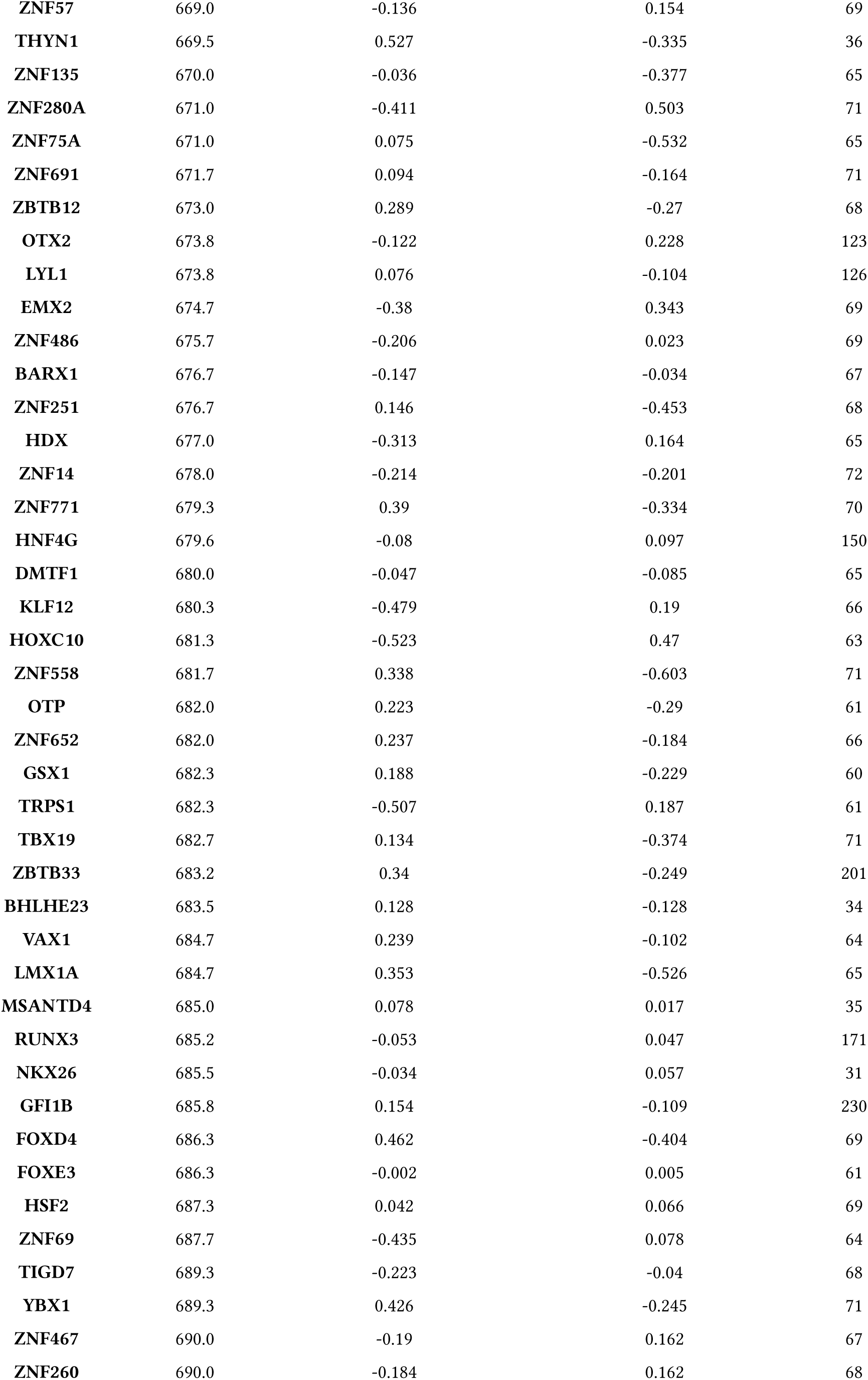
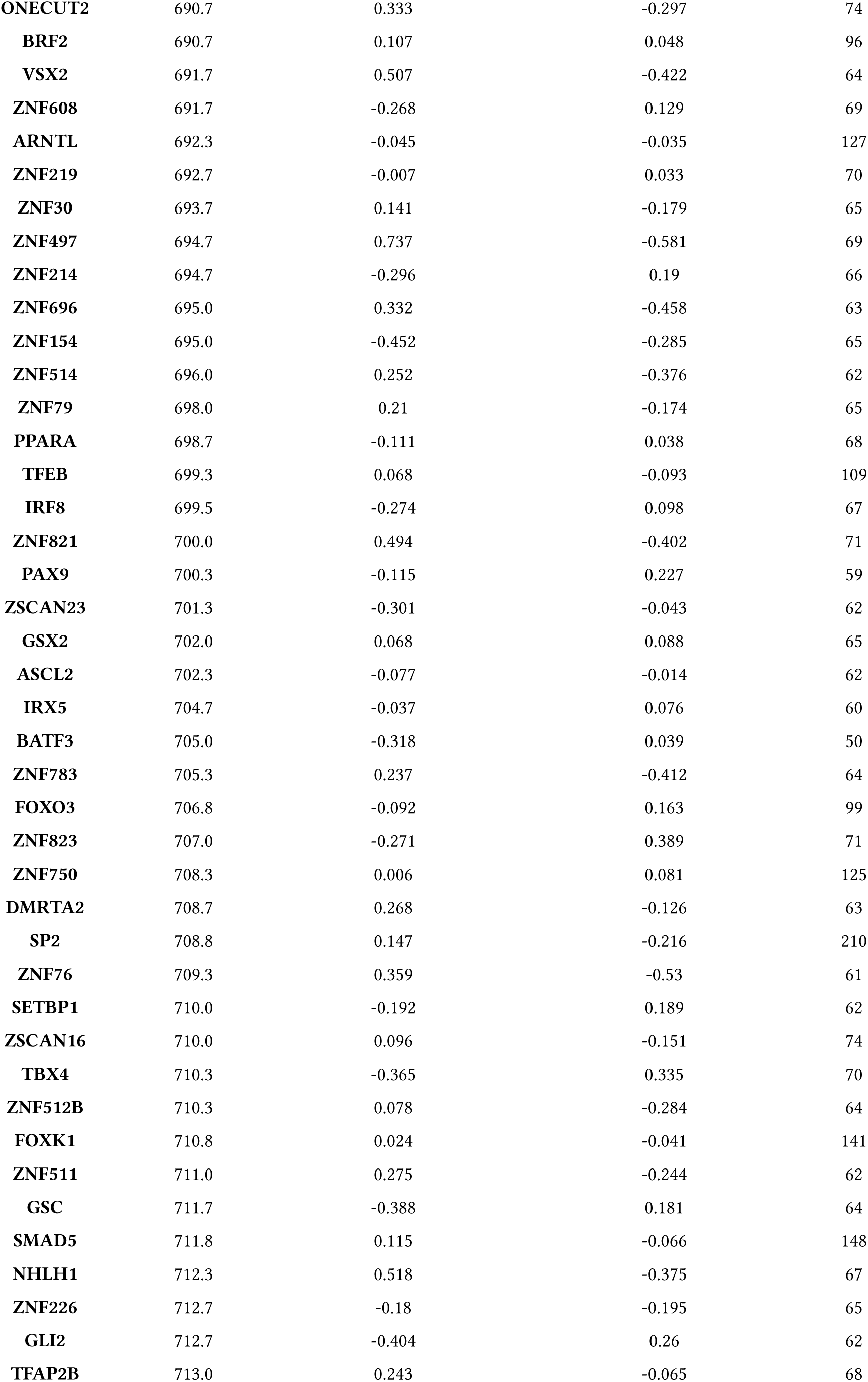
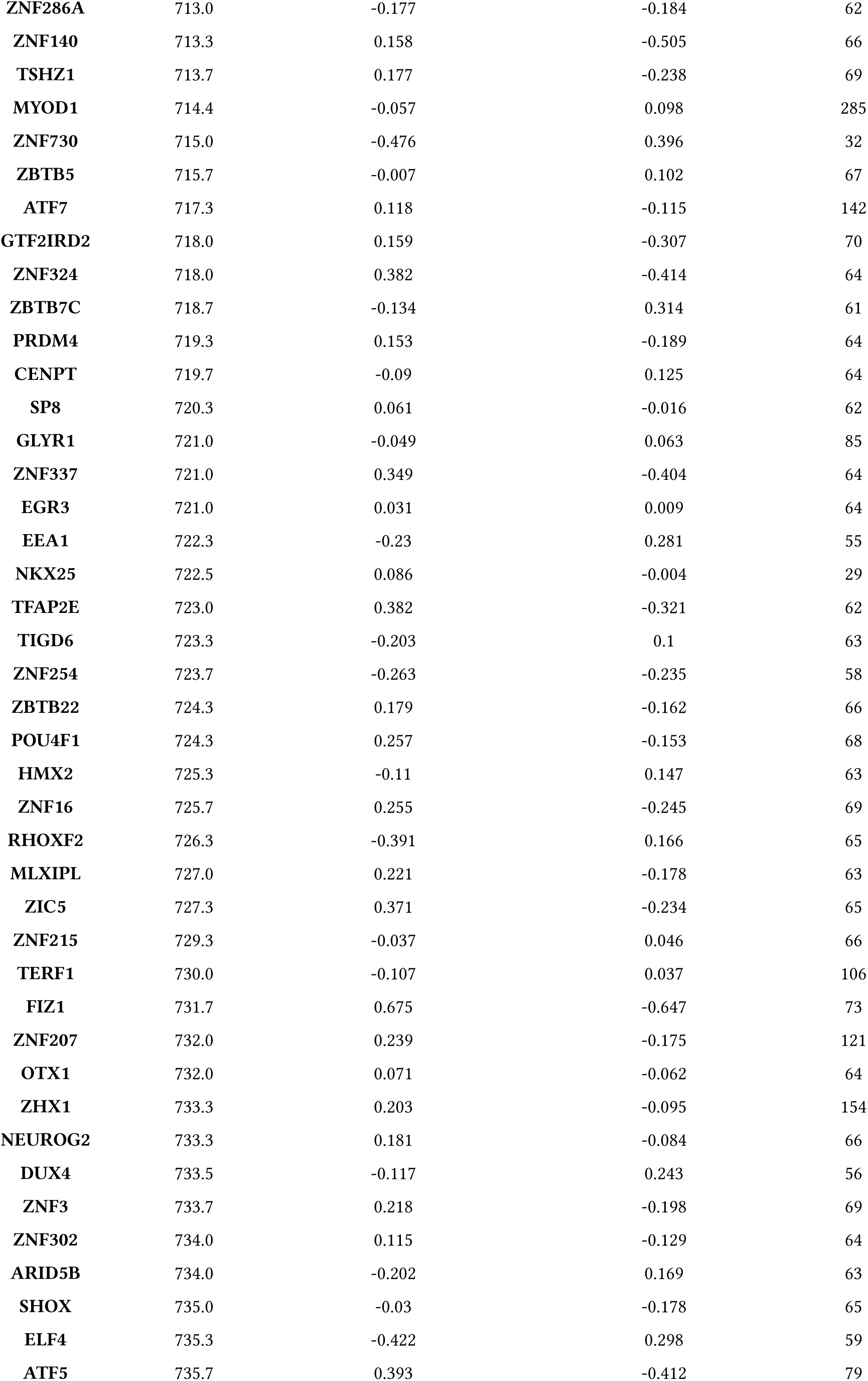
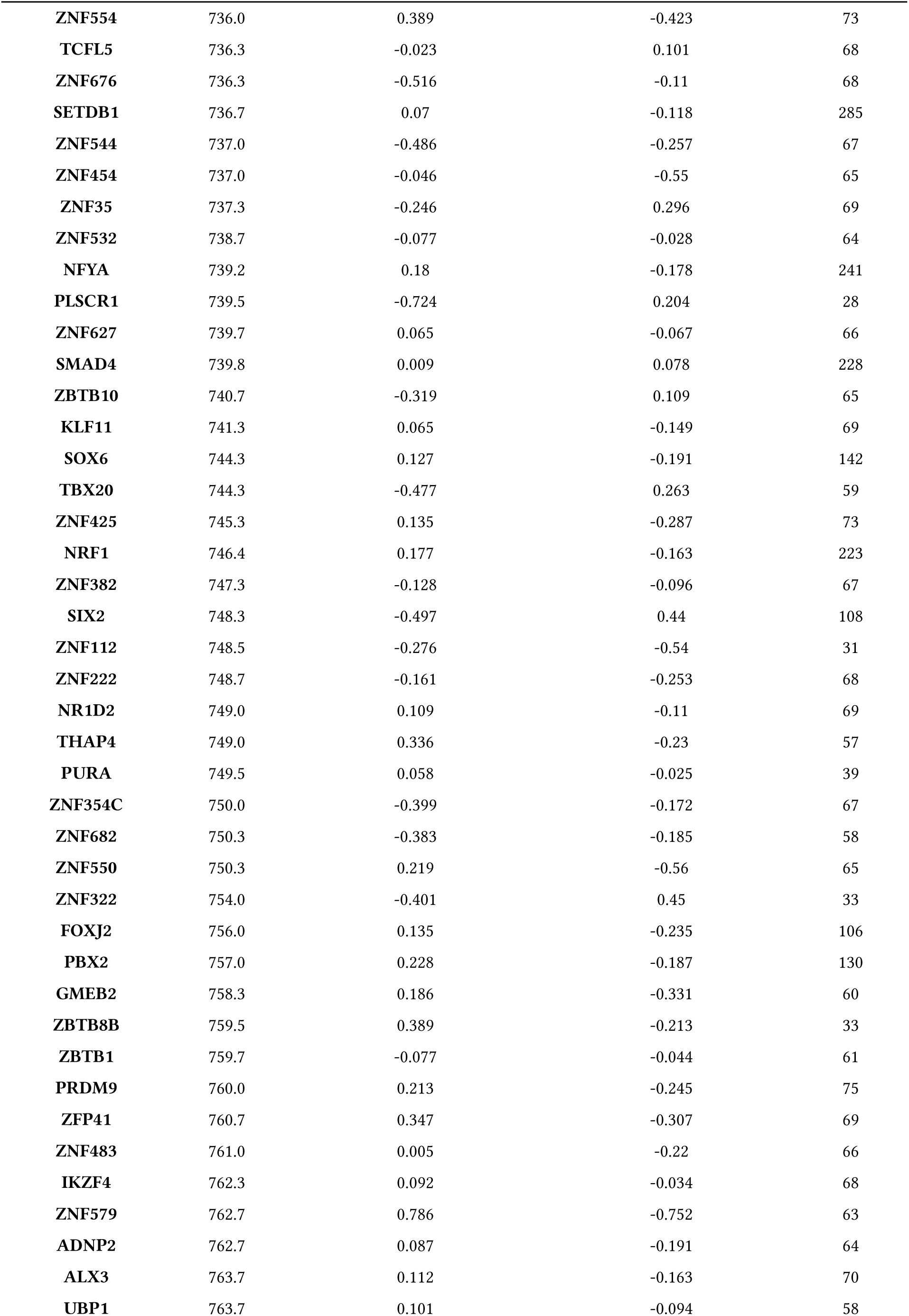
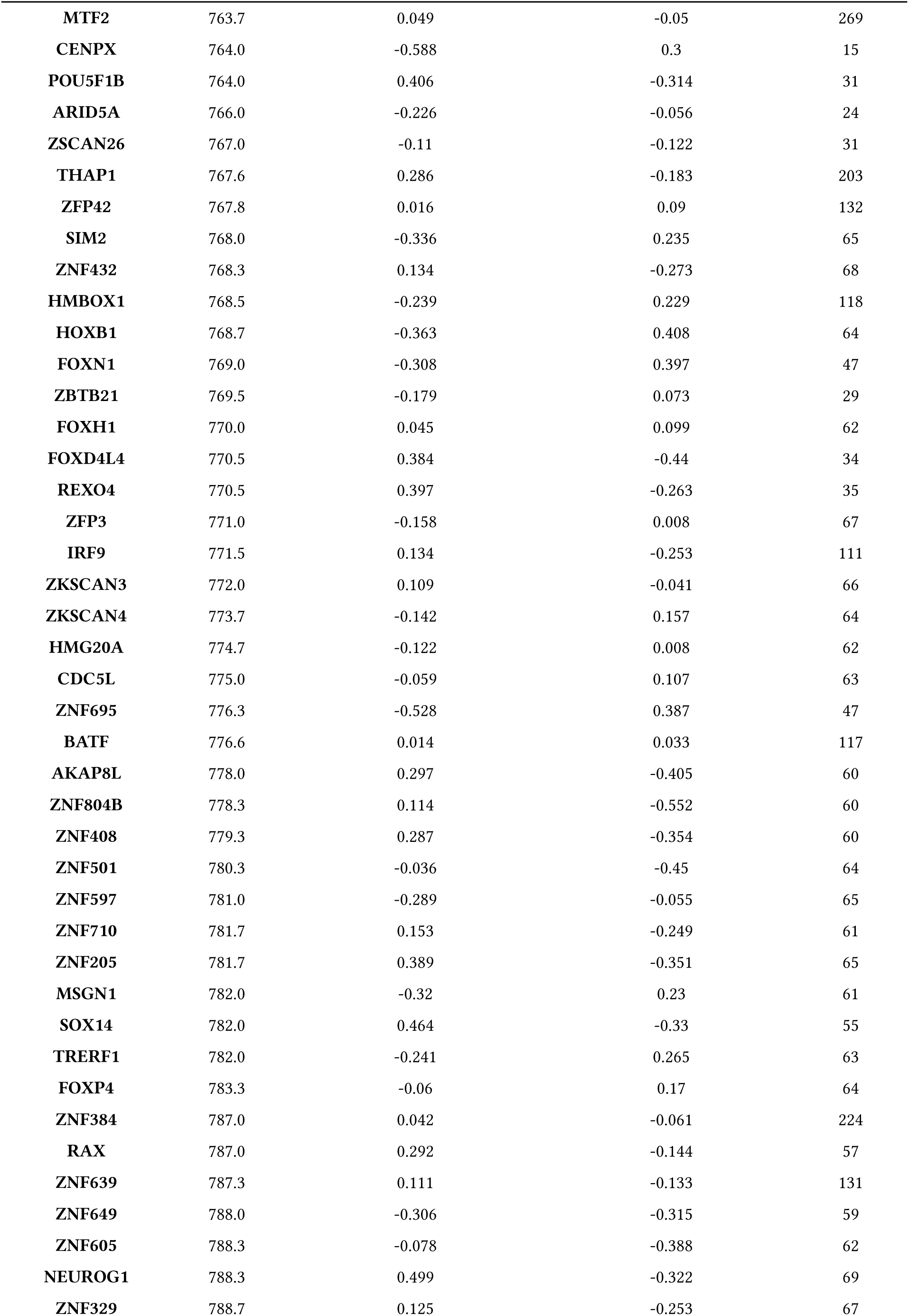
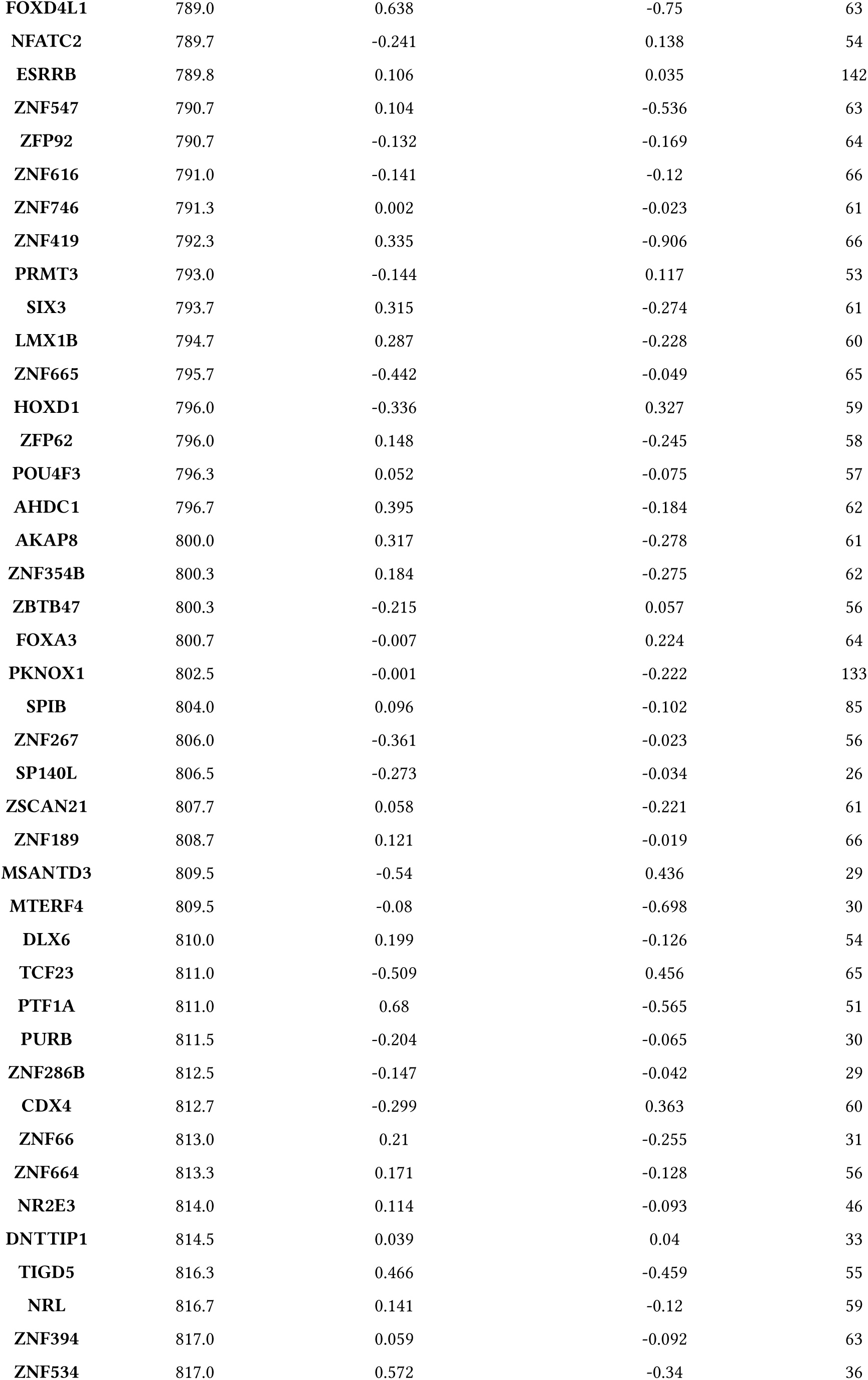
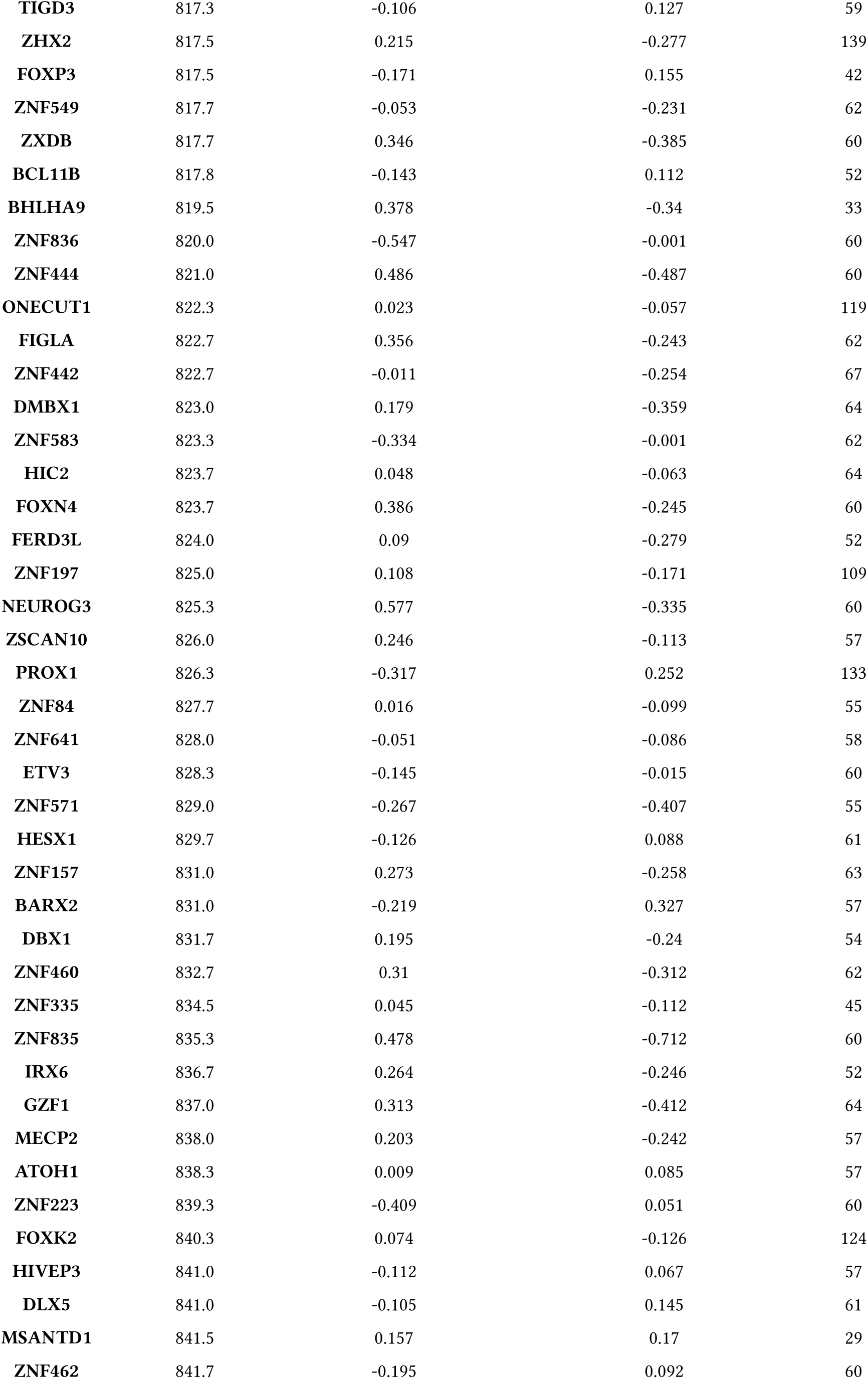
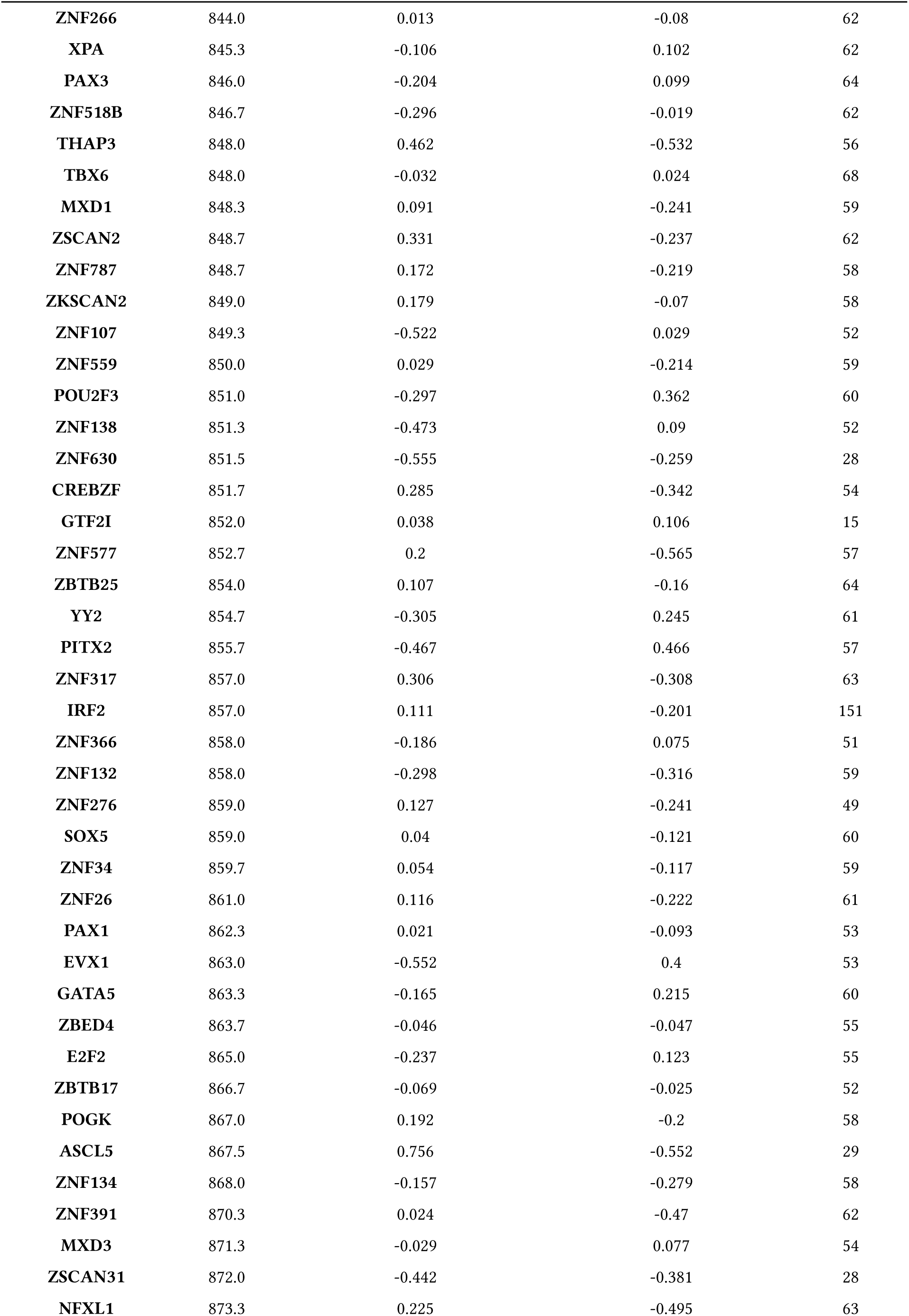
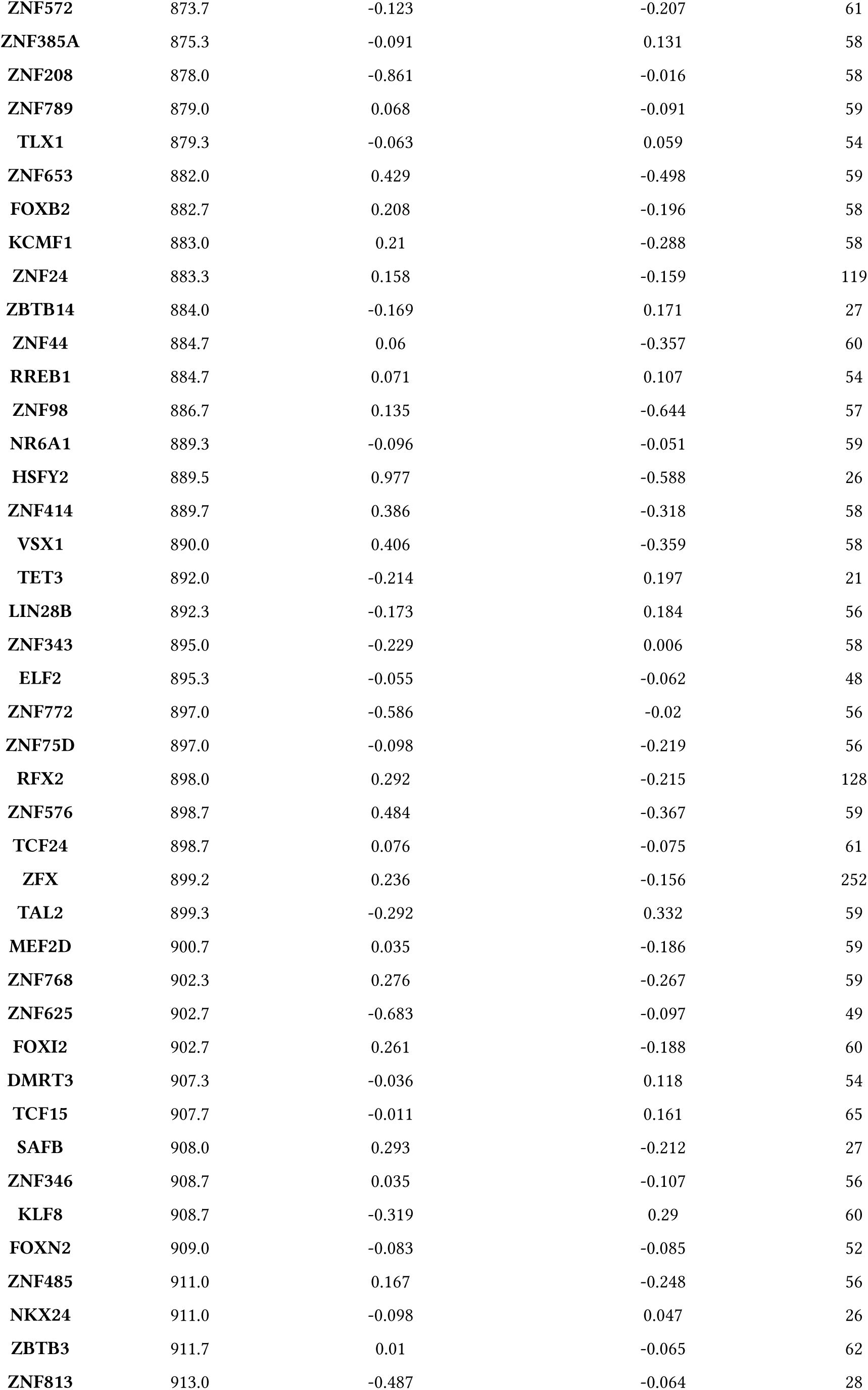
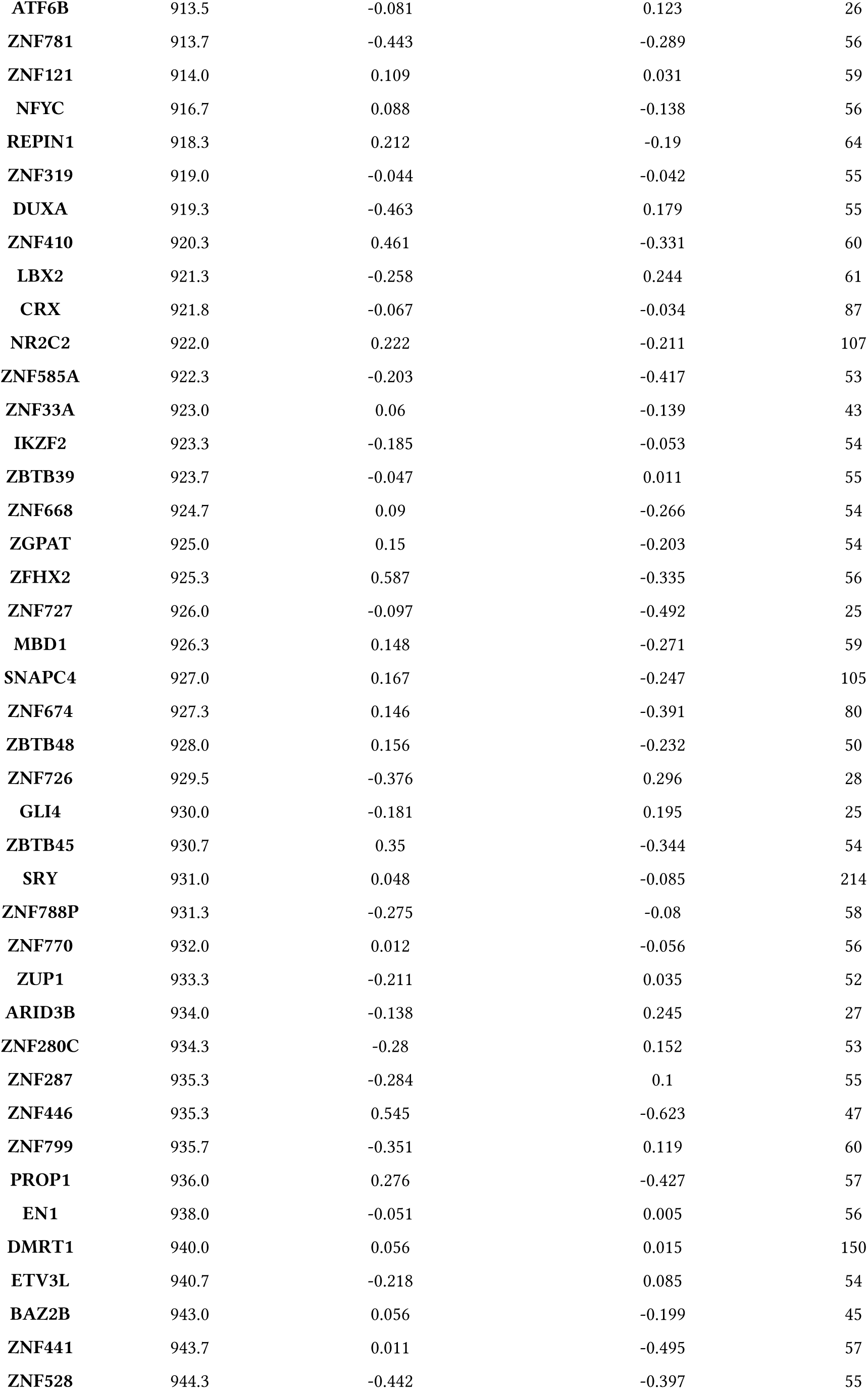
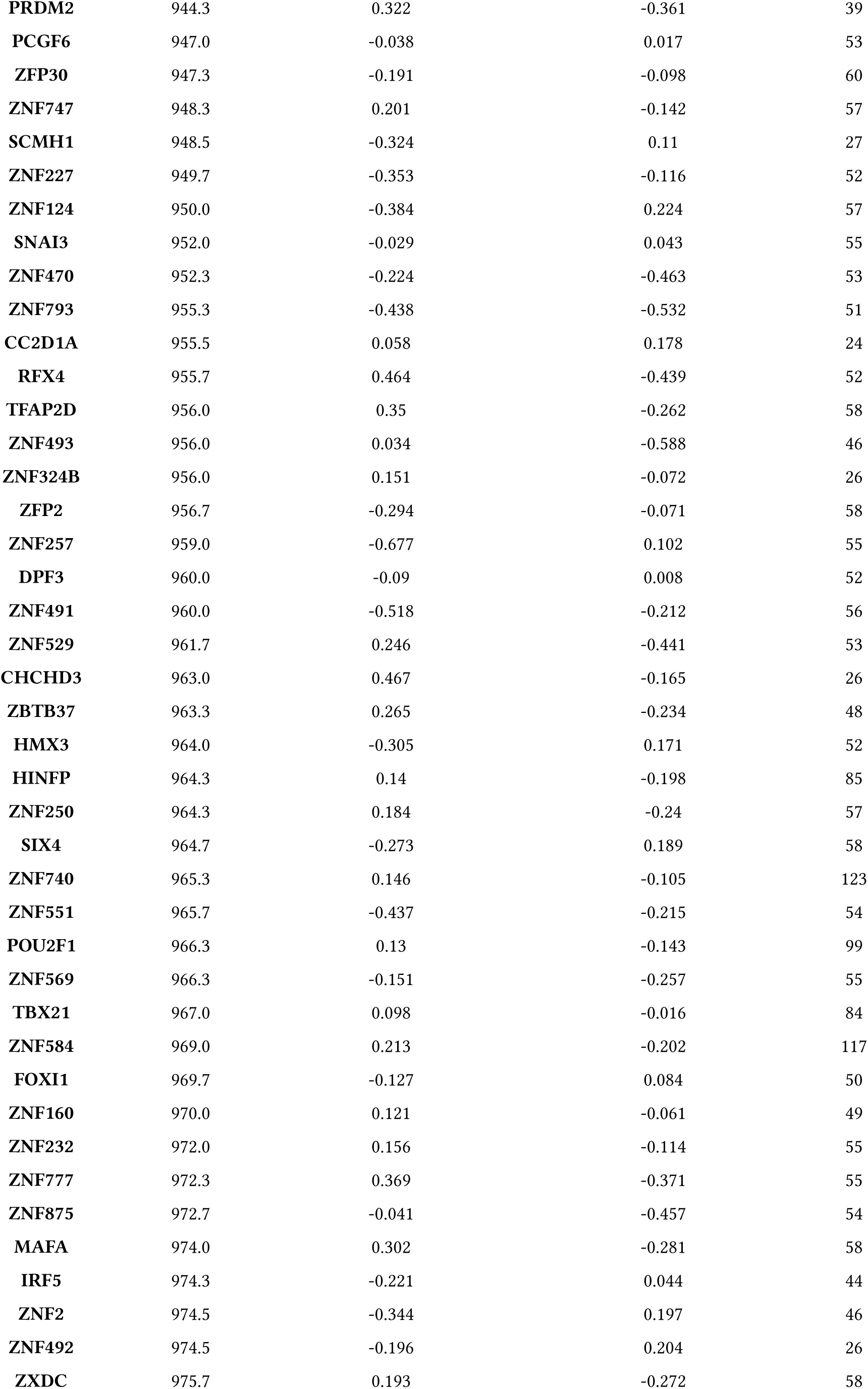
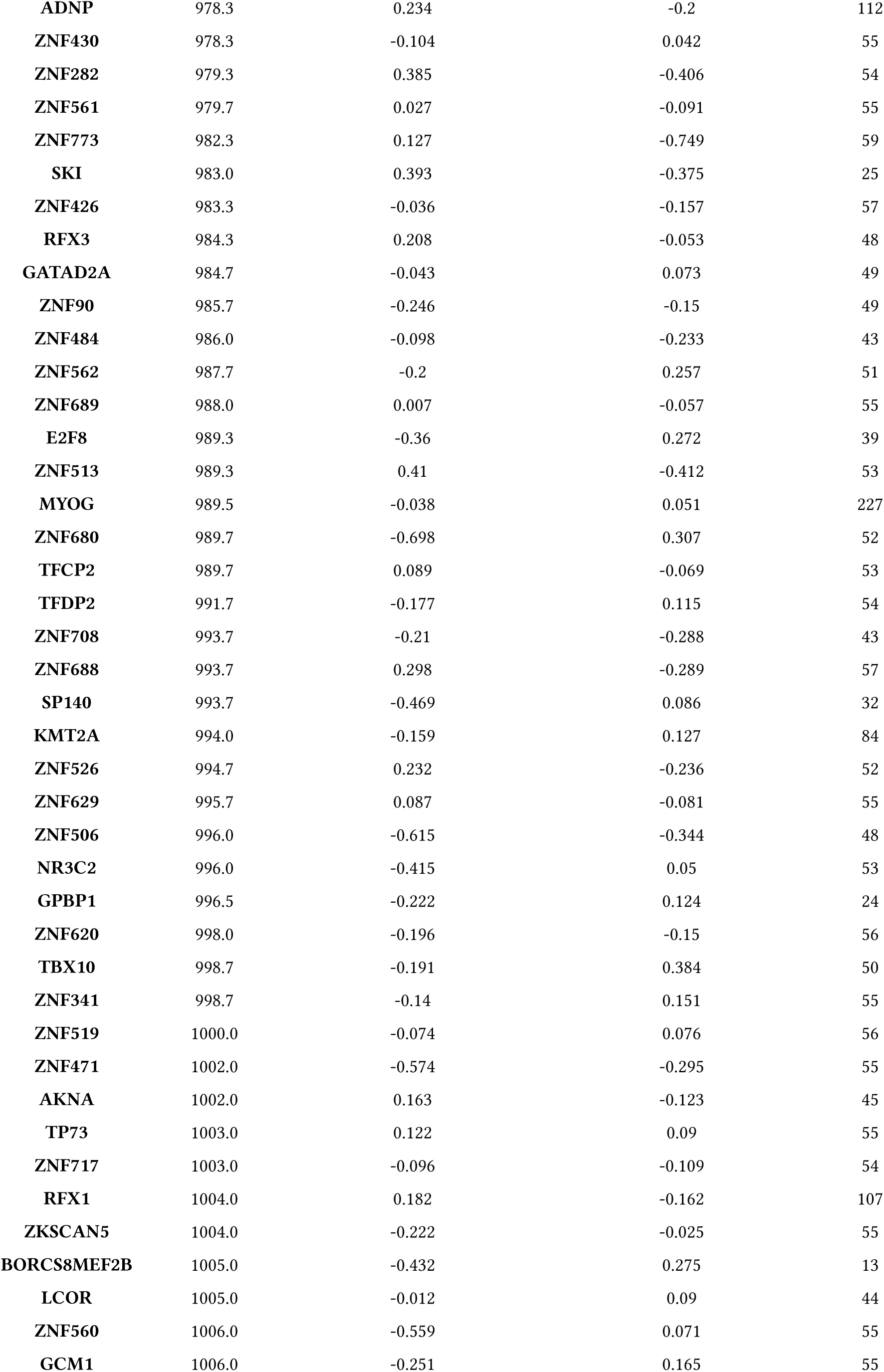
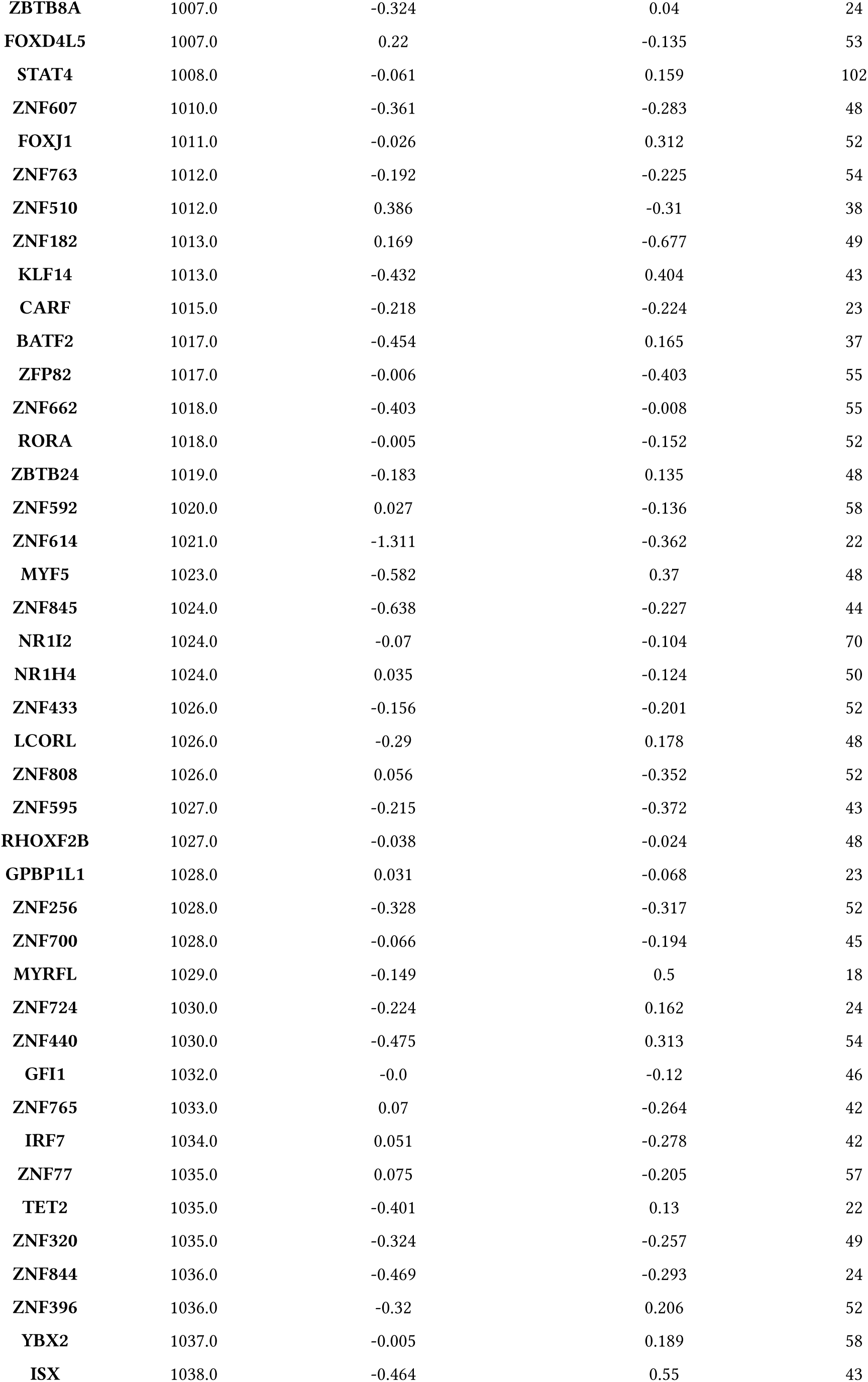
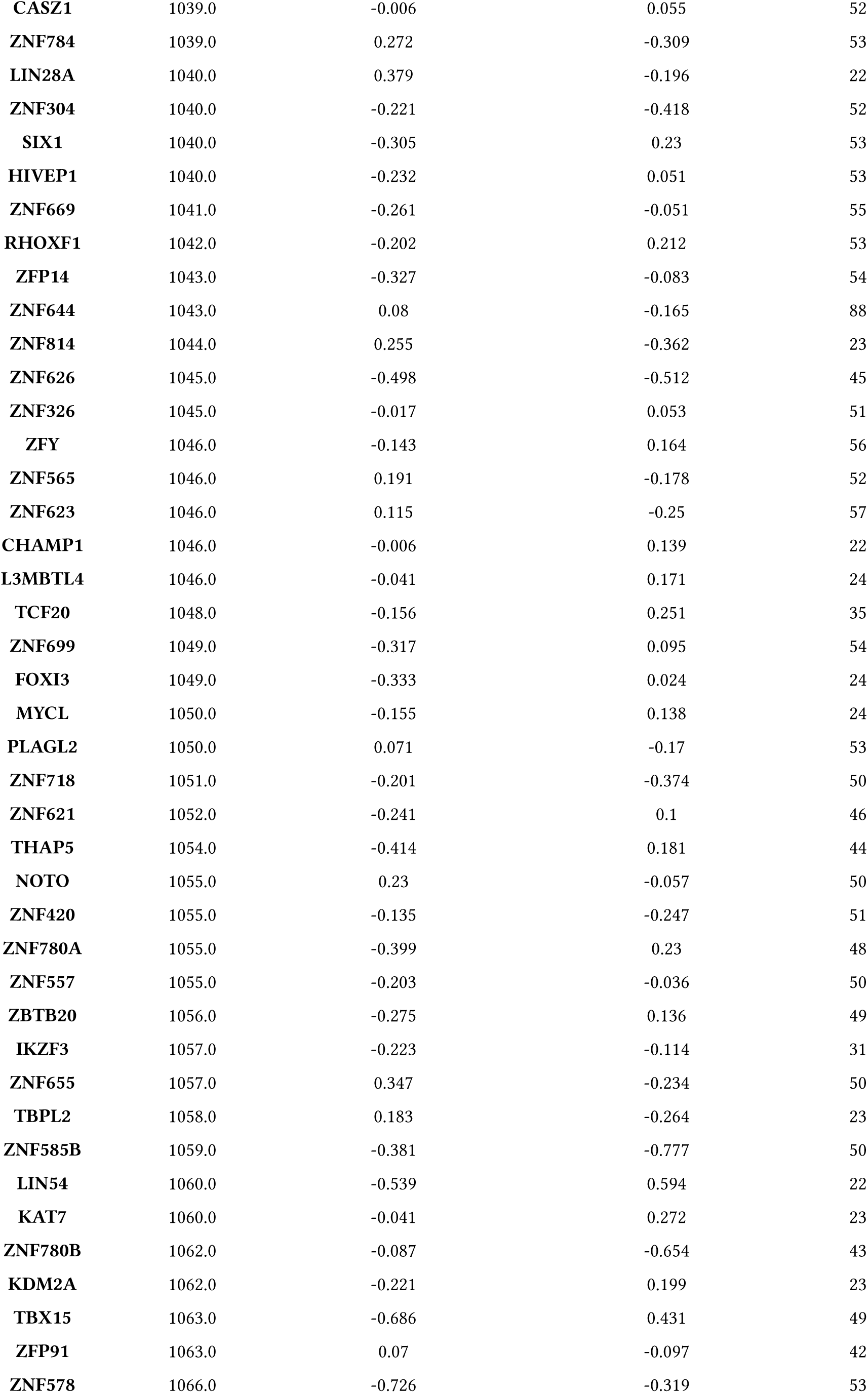
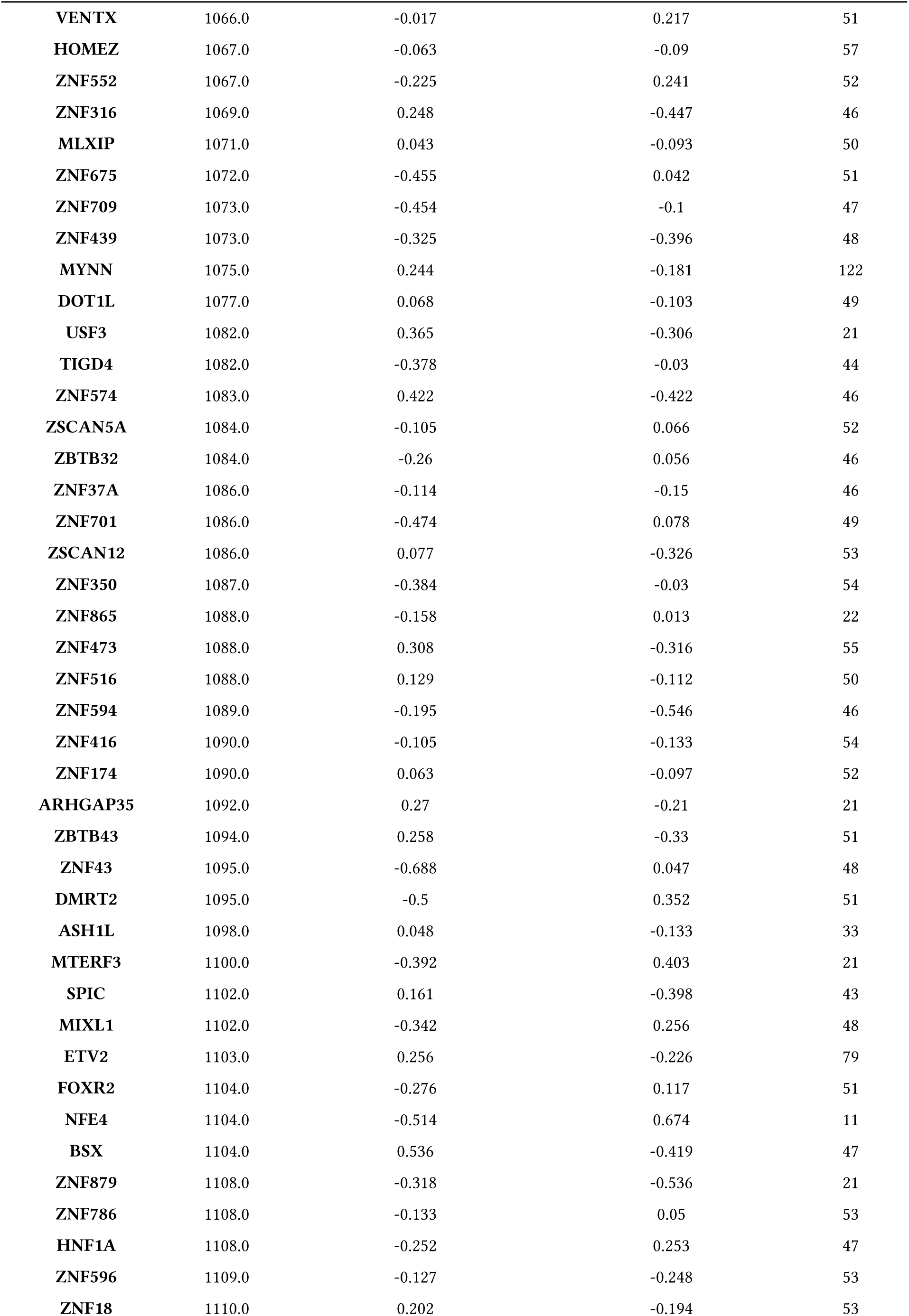
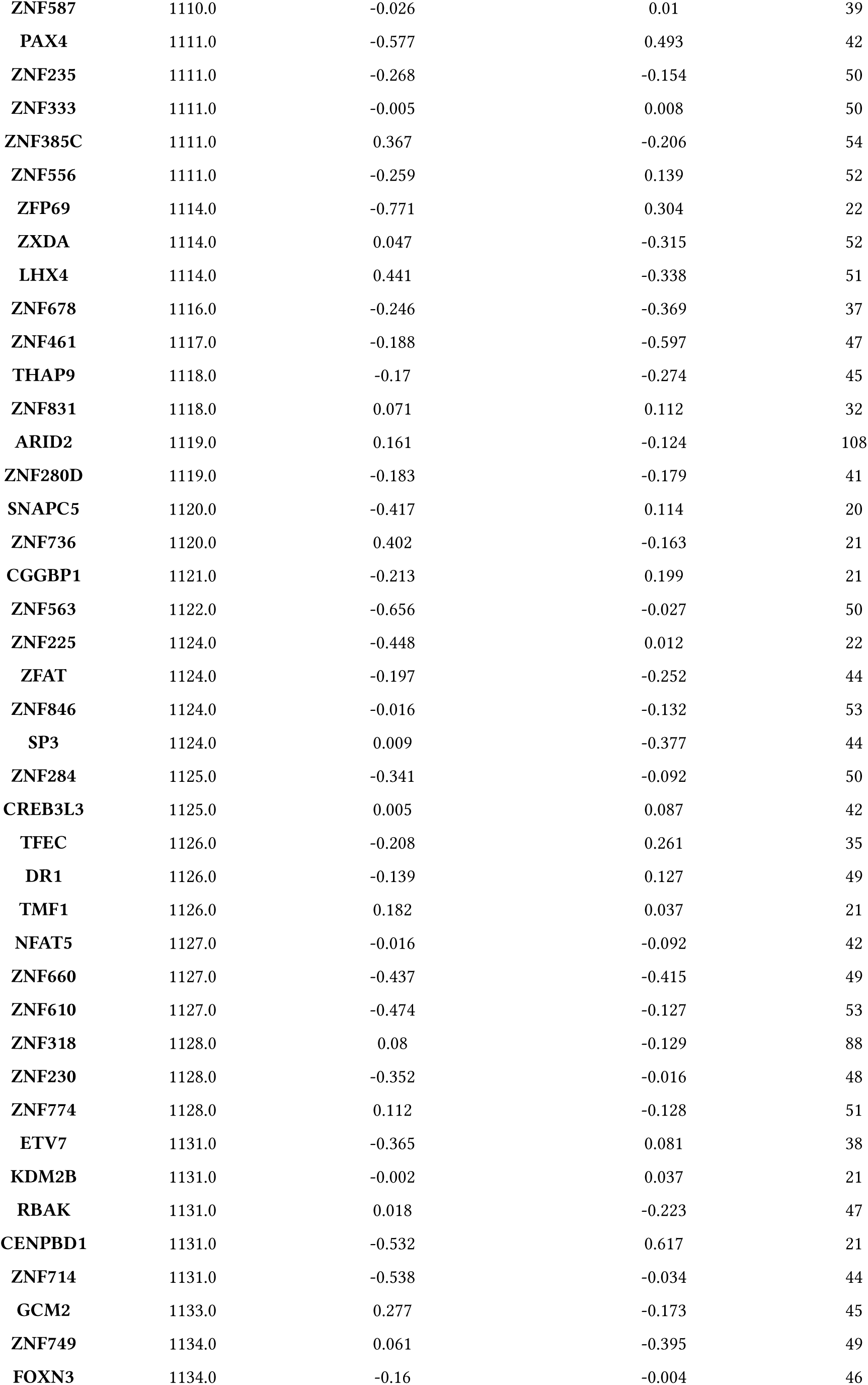
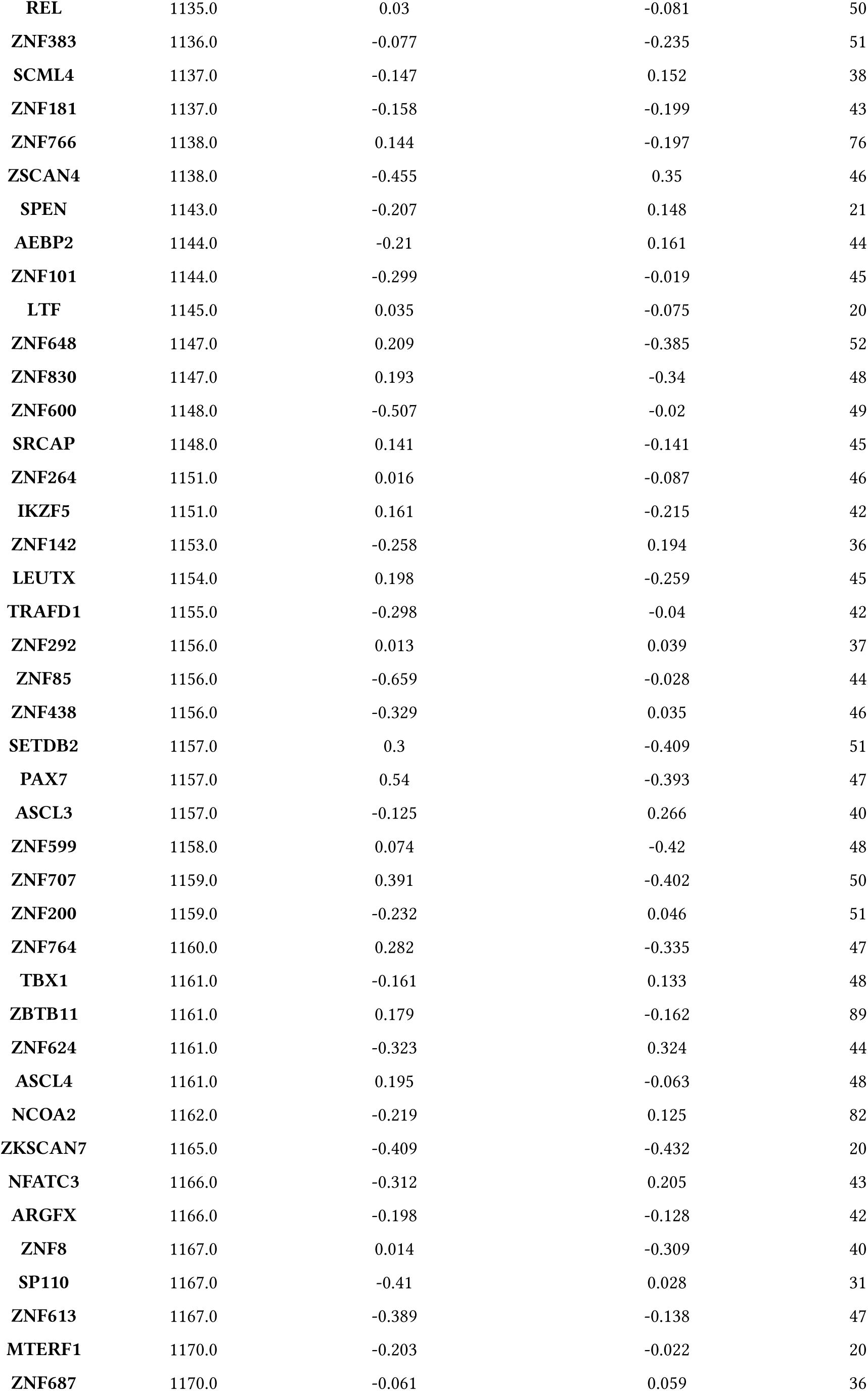
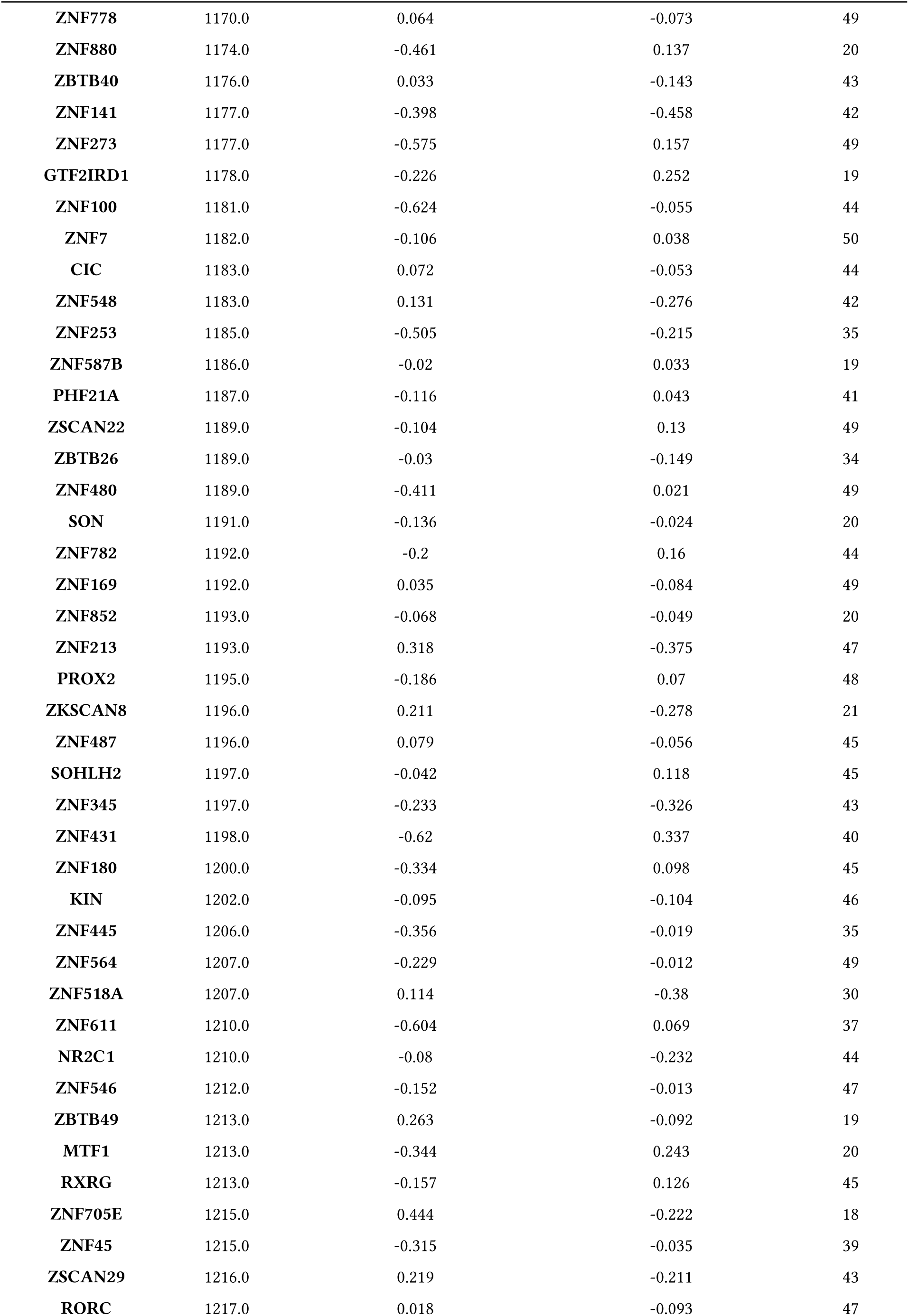
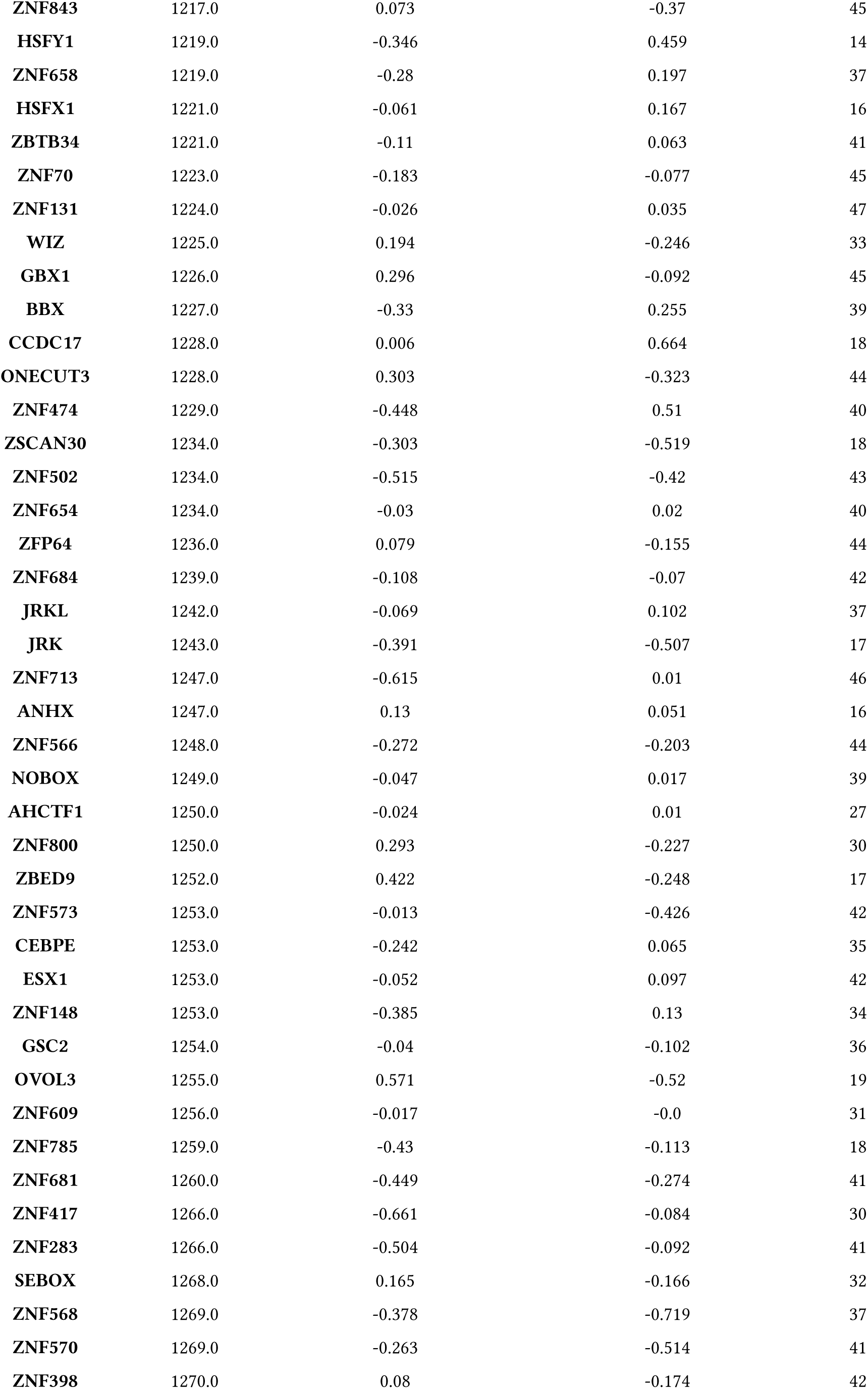
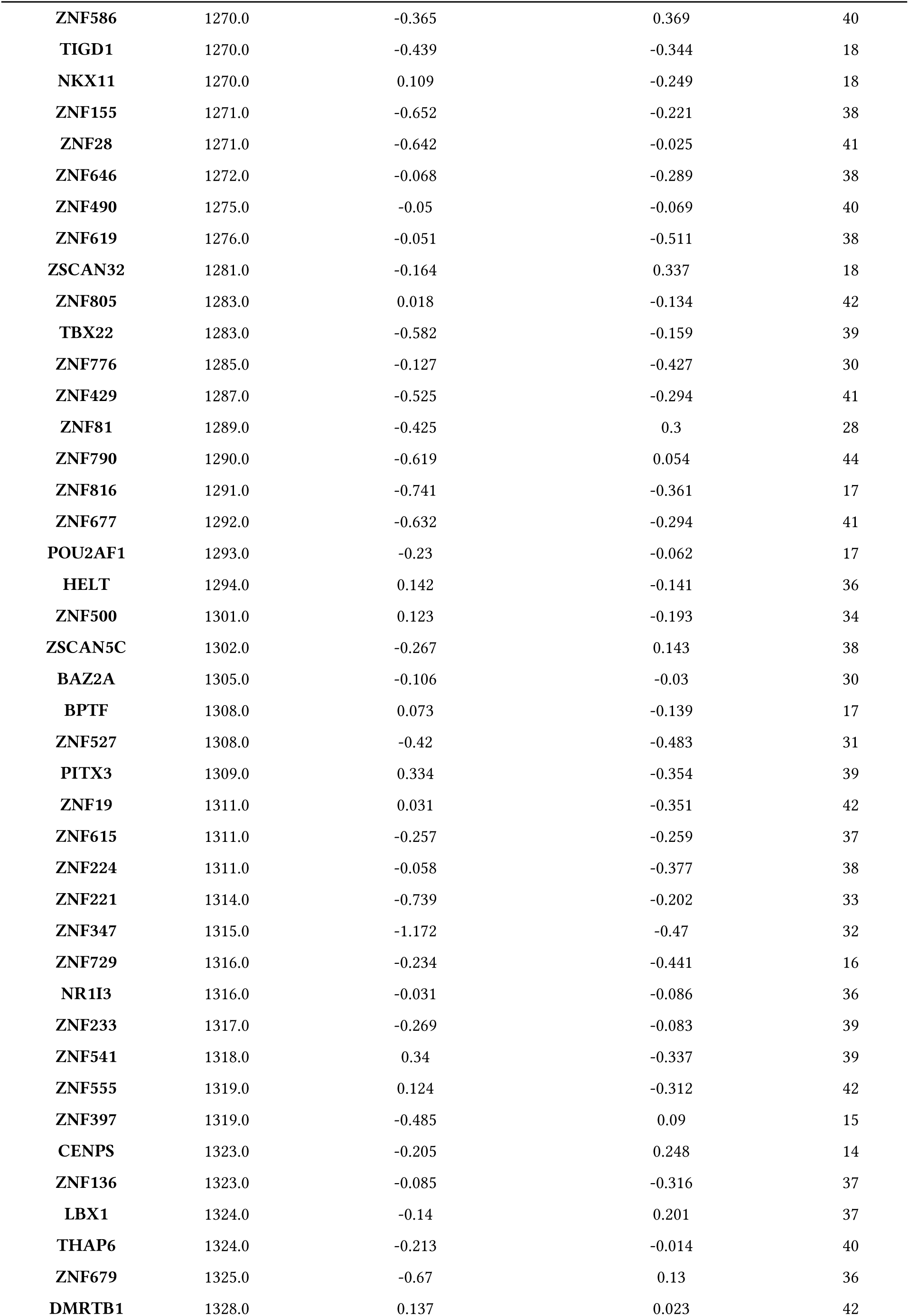
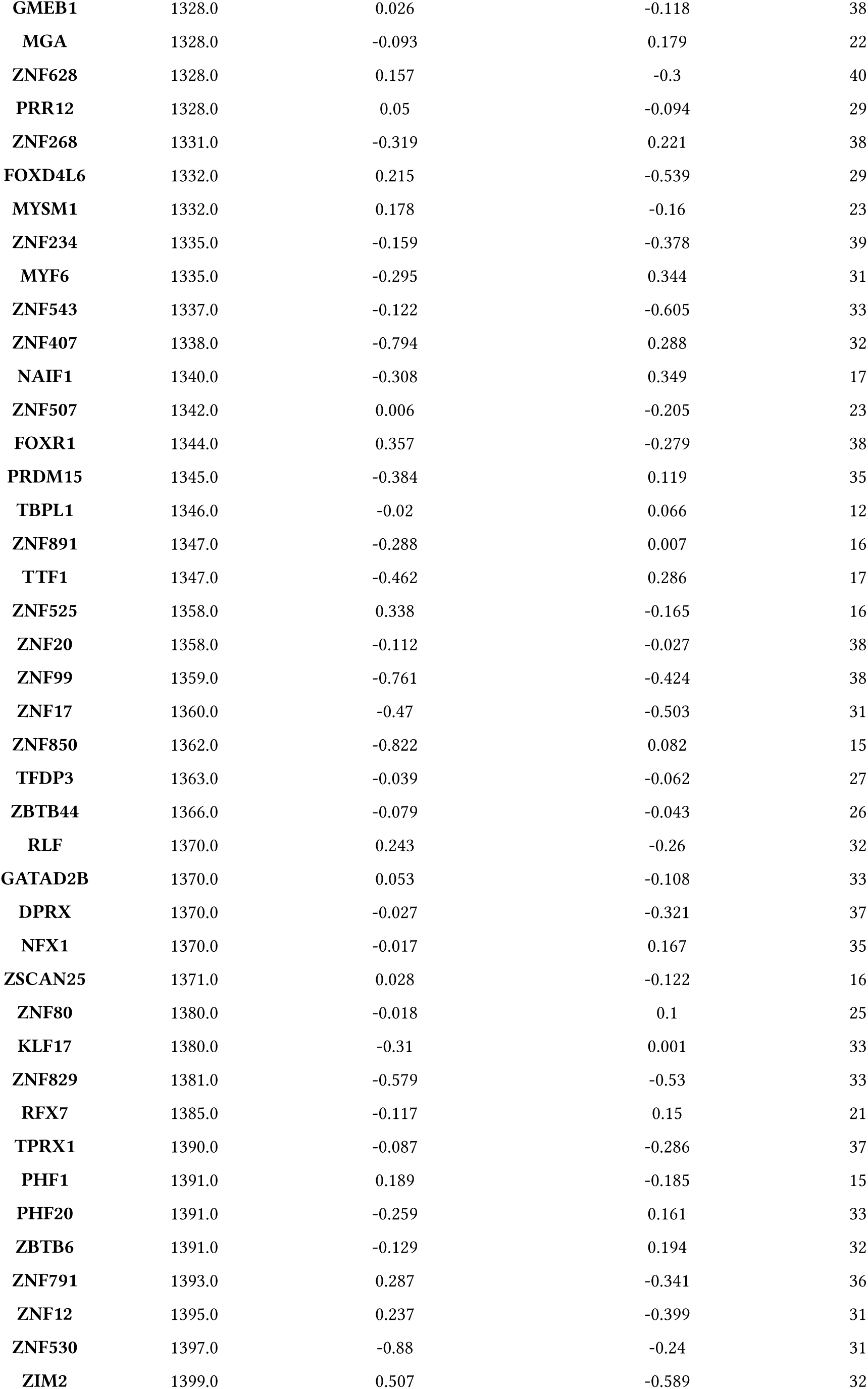
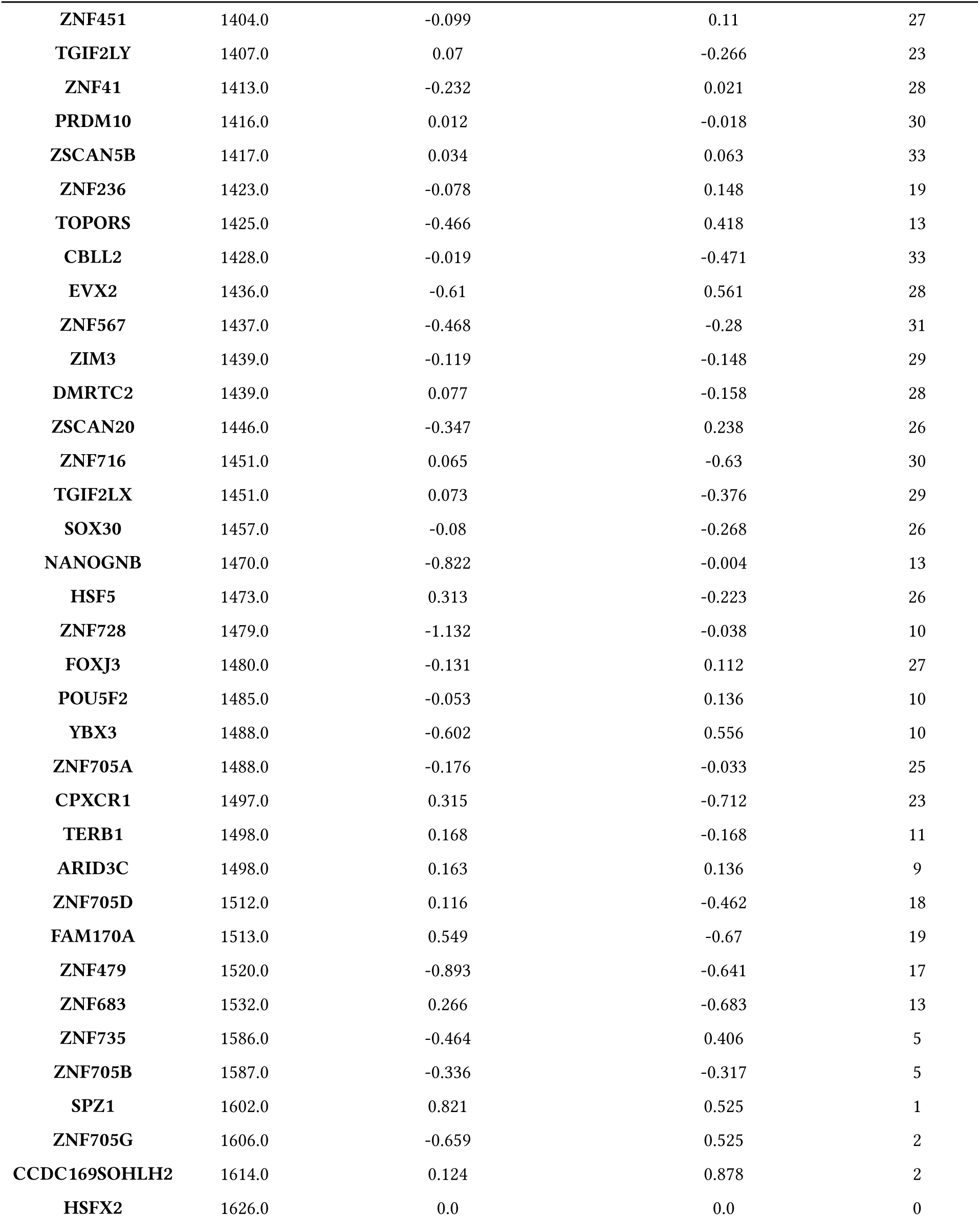
Result of the transcription factor enrichment analysis against the CHEA3 database.

