## Supplementary Table 3 for "The huntingtin–HAP40 complex is a bidirectional cellular rheostat": GST-HAP40.html

BayesInteractomics — Interactome Report

BayesInteractomics

H0 (no interaction) | H1 (interaction) | Agnostic (ambiguous)

**Selected:**

× Clear

- Results
- Evidence
- Calibration
- Sensitivity
- Mixture Model
- Structural Evidence
- Differential
- Data Quality
- Methods

**Posteriors are uncalibrated.** Run with `run_simulation=true` for calibrated posteriors and FDR.

Volcano Plot
Reset Selection

Click a point to highlight it across all plots and the table.
What is this?

This plot shows how strongly each protein interacts with your bait versus how confident the analysis is in that interaction. Proteins in the upper-right corner are both highly enriched and statistically supported as genuine interactors.

The x-axis shows log2 fold change (how much more abundant in bait vs control). The y-axis shows the combined Bayes factor on a log10 scale. Horizontal dashed lines mark conventional evidence thresholds (BF = 3, 10, 30). Click any point to highlight that protein across all tabs.

Rank-Rank Plot
Reset Selection

X: log₁₀(BF) · Y: log₂FC · Colour: log₁₀(BF correlation)
What is this?

This plot helps identify proteins that rank consistently as strong interactors across multiple types of evidence. Proteins in the upper-right corner show both high enrichment and strong correlation with the bait.

The x-axis shows the Bayes factor for enrichment (log10 scale), the y-axis shows log2 fold change, and colour intensity reflects the correlation Bayes factor. This multi-dimensional view helps distinguish true interactors from contaminants that may score high on only one metric.

Results Table
What is this?

This table lists every protein detected in your experiment with its interaction scores. Use the filters above to focus on high-confidence interactors or search for specific proteins of interest.

Key columns: posterior probability (overall confidence, 0-100%), PEP (per-protein false positive probability), BFDR (estimated FDR if you called everything at or above this protein significant), and individual Bayes factor values for enrichment, correlation, and detection. Click a row to highlight that protein in all plots.

⬇ Export CSV
Reset

Show sub-model BFs

Filters:

BF ≥

P(int|data) ≥

0.8
0.95

BFDR ≤

0.05
0.01

PEP ≤

0.05
0.01

Diag.

All
✔ OK
⚠ Warning
✘ Fail

Sens. ≤

Component:

H0

Agnostic

H1

Stability:

Robust

Sensitive

Fragile

Disagree

All
Disagree
Agree

✕ Clear filters

Diagnostic Flag Legend

✔ **OK** — No issues detected  
⚠ **Warning** — Low data (< 4 observations) or moderate residuals  
✘ **Fail** — Residual outlier (|mean residual| > 2)  
— No diagnostic data for this protein

| Protein | Bayes Factor | P(interaction\|data) | PEP | BFDR | log₂FC | BF Enrich. | BF Correl. | BF Detect. | Evidence | Diag | Sens. Range | Component | P(H0\|data) | P(agn\|data) | P(H1\|data) | Disagree | Stability | BF 3c-EM | BF Copula | Pareto k |
| --- | --- | --- | --- | --- | --- | --- | --- | --- | --- | --- | --- | --- | --- | --- | --- | --- | --- | --- | --- | --- |

0 proteins excluded (not detected in any sample)

| Protein ID | Name |
| --- | --- |

FDR vs Posterior Threshold
What is this?

This plot shows how many false positives you can expect at different confidence cutoffs. It helps you choose a threshold that balances discovering real interactions against accepting some false positives.

The x-axis is the posterior probability threshold; the y-axis is the estimated FDR. The curve is derived from simulation-based calibration. A typical target is FDR below 5% (dashed line).

Sensitivity vs Posterior Threshold
What is this?

This plot shows what fraction of true interactors you would recover at each confidence threshold. Stricter thresholds miss more real interactions but produce fewer false positives.

The x-axis is the posterior probability cutoff. The y-axis is sensitivity (true positive rate). The curve is estimated from synthetic data generated to match your experiment's characteristics.

ROC Curves
What is this?

This plot evaluates the overall ability of the analysis to distinguish true interactors from background proteins. A curve hugging the top-left corner indicates excellent discrimination.

The ROC curve plots sensitivity (true positive rate) against 1 minus specificity (false positive rate) at all possible thresholds. The area under the curve (AUC) summarizes performance: 1.0 is perfect, 0.5 is random.

Reliability Diagram
What is this?

This plot checks whether the reported confidence values are trustworthy. If a protein is assigned 80% probability, it should truly be an interactor about 80% of the time.

Points on the diagonal line indicate well-calibrated probabilities. Points above the diagonal mean the analysis is underconfident; below means overconfident. Bars show how many proteins fall in each confidence bin.

Threshold Recommendations
What is this?

This table suggests optimal confidence thresholds for different analysis goals, such as maximizing the number of discoveries or minimizing false positives.

Each row shows a threshold strategy with its estimated FDR, sensitivity, specificity, and the number of proteins that would be called significant. Choose based on whether you prioritize completeness or precision.

| pi\_H1 | Effect Scale | FDR≤1% | FDR≤5% | FDR≤10% | Sensitivity@5%FDR |
| --- | --- | --- | --- | --- | --- |

Reliability Diagram: Raw vs Calibrated
What is this?

This plot compares confidence values before and after calibration. Calibrated probabilities should align more closely with the diagonal, meaning they better reflect actual interaction rates.

Blue points show raw posterior probability values; orange points show calibrated values. If calibration improved the estimates, orange points will be closer to the diagonal than blue.

Calibration Function
What is this?

This plot shows the mathematical transformation applied to adjust raw confidence values into calibrated probabilities.

The x-axis is the raw posterior probability; the y-axis is the calibrated value. The calibration function (Platt scaling) is a logistic regression fitted on simulation ground truth. Points near the diagonal indicate minimal adjustment was needed.

Cross-Validation ECE
What is this?

This table reports how well the calibration holds up when tested on data not used to fit the calibration model, ensuring the correction generalises beyond the training data.

ECE (Expected Calibration Error) measures the average gap between predicted probabilities and observed frequencies. Lower ECE means better-calibrated posteriors. Values are from cross-validation folds.

| Fold | Posterior ECE | FDR ECE |
| --- | --- | --- |

FDR Accuracy: Declared vs Actual
What is this?

This plot checks whether the declared false discovery rates match the actual error rates observed in simulations. Points near the diagonal mean the FDR estimates are accurate.

The x-axis is the declared FDR (what you are told); the y-axis is the actual FDR (measured from simulation ground truth). A well-calibrated analysis produces points along the diagonal.

Reading the Sensitivity Labels

|  |  |
| --- | --- |
| LC | Latent Class model — the 3-component EM that separates null, agnostic, and interactor proteins. The α vector sets the Dirichlet prior on mixing weights. |
| α | Dirichlet concentration parameters [α_H0_, α_Ag_, α_H1_]. Larger values impose a stronger prior belief about the proportion of each component. |
| E[π_1_] | Expected prior probability of a protein being an interactor (H1 component). Derived from the Beta(α,β) prior on the copula EM mixing weight. E.g. E[π_1_]=0.125 means 12.5% of proteins are expected to be true interactors a priori. |
| BMA | Bayesian Model Averaging — combines Copula and LC posteriors using LOO stacking weights. Each grid point re-estimates the stacking weights from scratch. |

Sensitivity Rank Correlation
What is this?

This plot summarises how similar the protein rankings are across all tested prior specifications. High correlations mean the ordering of interactors is stable regardless of prior choice.

Each point is the Spearman rank correlation between the default analysis and one alternative prior sensitivity specification. Correlations above 0.95 indicate excellent robustness.

Pairwise Spearman Rank Correlation
What is this?

This matrix shows how similar the protein rankings are between every pair of prior specifications. All-blue means results are completely insensitive to prior choice.

Each cell is the Spearman rank correlation between two prior specifications. Values close to 1.0 indicate that the protein ranking is preserved. Off-diagonal entries below 0.95 flag settings where the ranking changes meaningfully.

Decision-Boundary Stability
What is this?

Each whisker shows the full range of posterior probability for one protein across all tested priors. Proteins whose whiskers cross the red dashed line (P=0.5) change classification depending on prior choice -- these are flagged in red.

Proteins are sorted left-to-right by decreasing sensitivity range. The dashed red line marks the P=0.5 decision boundary. Red markers indicate proteins that cross this boundary across the prior grid. Narrow whiskers indicate robust classification.

Posterior Overlay (Top Sensitive Proteins)
What is this?

For the most sensitive proteins, this plot shows the exact posterior probability under each prior setting as individual points. Tight clusters mean the result is stable; spread-out points reveal sensitivity to prior choice.

Each column is one protein (the top 15 most sensitive). Each dot within a column represents the posterior probability under one prior setting. The red dashed line marks P=0.5. Proteins with points on both sides of this line change classification across priors.

BMA Stacking Weights Across Prior Grid
What is this?

This chart shows how the relative importance of the two statistical models (Copula and 3c-EM) changes across different prior specifications. Stable weights indicate that model selection is robust to prior choice.

Each bar shows the LOO (leave-one-out) stacking weight for the 3c-EM and Copula models at each BMA prior grid point. Weights sum to 1.0. Large shifts across the grid indicate sensitivity of model selection to prior specification.

Component Assignment Scatter
Reset Selection

Proteins colored by MAP component assignment (H0=blue, Agnostic=gray, H1=red).
What is this?

This plot shows how the statistical model has classified each protein into one of three groups: background (non-interactors), ambiguous, or genuine interactors. Clear separation between groups indicates the model can reliably distinguish interactors from noise.

Each point is a protein, positioned by its enrichment Bayes factor (x) and correlation Bayes factor (y) on log10 scale. Colour indicates component assignment from the 3c-EM model.

Marginal Density Overlays

Histogram of combined log-BF values with fitted Gaussian components overlaid.
What is this?

These plots show how well the statistical model captures the distribution of evidence scores in your experiment. Good fits mean the model assumptions match your data.

Each panel shows the observed distribution of a Bayes factor type (histogram) overlaid with the fitted mixture components. The background (H0) component captures the null distribution; the H1 component captures genuine interactors.

EM Convergence Trace 
What is this?

This trace shows whether the model fitting algorithm reached a stable solution. A plateau at the end indicates the model has converged.

EM convergence is assessed by monitoring the log-likelihood across iterations. A monotonically increasing trace that levels off confirms stable parameter estimates.

Component Mixing Weights 
What is this?

This bar chart shows the estimated proportion of proteins in each category: background, ambiguous, and interactors. It gives an overview of how many genuine interactions the model detected in your experiment.

The three bars show the mixture weights from the 3c-EM model: H0 (background), Agnostic (ambiguous), and H1 (interactors). These proportions are estimated from the data and reflect the overall interaction prevalence.

Quality Gate Matrix

Goodness-of-fit per (marginal x component). KS test for enrichment/correlation, χ² test for detection. Green=pass, Yellow=warn, Red=fail.
What is this?

This matrix provides a quick pass/fail overview of key quality checks on the statistical model. Green cells indicate the model is behaving as expected; yellow or red cells flag potential issues.

Each cell tests a specific aspect of model fit: marginal goodness-of-fit (KS test for enrichment/correlation, chi-squared test for detection), component separation, contamination between H0 and H1, and within-class independence. Click individual cells for details.

|  | H0 | Agnostic | H1 |
| --- | --- | --- | --- |
|  |  |  |  |
| --- | --- | --- | --- |

⚠

**High disorder rate detected.**
A large fraction of docked candidates are predicted to be significantly disordered
(fraction\_disordered > 0.5). For these proteins, docking scores are unreliable and
BF\_dock has been set to 1.0 (no update). Disordered proteins are flagged with
status "disordered" in the table below.

ⓘ

**Interpreting docking scores.**
A low C2Qscore, ipTM, or BF_dock_ does *not* mean a protein is a false positive.
AlphaFold 3 was trained on stable co-crystal complexes (PDB) and performs
near-randomly on transient interactions — the very class
that AP-MS excels at detecting. Low or negative docking scores more likely indicate an
**indirect** or **transient** interaction, not the absence
of interaction. Always check the `docking_status` column before drawing
conclusions.
Scoring details

**C2Qscore** (Genz et al. 2025) is a linear combination of four
AlphaFold 3 metrics (ipTM, pTM, iPAE, ipLDDT) calibrated for AF3 docking quality
prediction. Negative values indicate low predicted structural accuracy —
this is expected for weak or transient complexes. The score is converted to a
Bayes factor via logistic regression.

**Scoring tiers:**
*C2Qscore* (preferred, when full\_data available) →
*pDockQ (legacy)* →
*ipTM (fallback)*. The tier used per pair is shown in
the `Tier` column.

**Benchmark performance** (Genz et al. 2025, 1265 AF3 models, DockQ ≥ 0.23 = correct):

| Metric | AUC-ROC | AUC-PR | MCC |
| --- | --- | --- | --- |
| **C2Qscore (4-metric)** | **0.929** | **0.978** | **0.737** |
| ipTM alone | 0.924 | 0.974 | 0.712 |
| pDockQ2 | 0.825 | 0.953 | 0.559 |

AUC-ROC = area under the receiver operating characteristic curve;
AUC-PR = area under the precision-recall curve;
MCC = Matthews correlation coefficient (best threshold).
Higher is better for all three metrics.

ipTM vs -log₁₀(PEP)
Points above ipTM 0.7 = high-confidence direct binders. Points below 0.4 = likely indirect or disordered.
What is this?

This plot compares the mass spectrometry evidence for each interaction with the structural prediction confidence from AlphaFold. Proteins in the upper-right have both strong MS evidence and plausible predicted structures.

The x-axis shows -log₁₀(PEP) from the MS-only analysis (higher values = more confident); the y-axis shows the ipTM score from AlphaFold structure prediction. An ipTM above 0.6 suggests a plausible physical interaction; below 0.2 provides no structural support.

PEP Update: Before vs After Docking
Points above the diagonal were boosted by docking evidence (higher -log₁₀(PEP) = more confident).
What is this?

This plot shows how structural evidence from AlphaFold changes interaction error rates. Proteins that shift downward gained support from structural predictions (lower PEP = more confident). The log₁₀ scale reveals changes among high-confidence interactions that would be compressed near zero on a linear scale.

Both axes show -log₁₀(PEP) (higher = more confident). The x-axis is the MS-only evidence; the y-axis is the updated value after incorporating structural Bayes factor evidence. Points above the diagonal gained confidence from docking; below lost confidence.

Docking Results
What is this?

This table lists all proteins for which structural predictions were obtained, with their structural quality scores and the resulting update to interaction probabilities.

Key columns: ipTM (interface quality), pLDDT (local structure confidence), pDockQ (docking quality), and the structural Bayes factor used to update the MS-based posterior. Values are from AlphaFold Server predictions.

Status

All
Success
Disordered
Too large
Pending

ipTM ≥

Export Docking CSV
Clear filters

| Protein | P(MS) | P(Combined) | BF Dock | ipTM Best | ipTM Std | Ranking | Disordered | PAE Min | Status | BFDR (combined) | PEP (combined) | pDockQ | C2Qscore | Tier | Tokens | 3D |
| --- | --- | --- | --- | --- | --- | --- | --- | --- | --- | --- | --- | --- | --- | --- | --- | --- |

×

Scale Detection
What is this?

BayesInteractomics expects intensity values on a log2 scale. If your data is on the original linear scale (raw intensities), the statistical models may produce unreliable results due to the highly skewed distribution of untransformed protein abundances.

Scale detection checks the maximum observed intensity value. Log2-transformed AP-MS data typically falls in the range of 10-35. Maximum values above 1000 strongly suggest linear-scale (untransformed) data. If flagged, apply log2 transformation before re-running the analysis.

Replicate Correlation
What is this?

Replicate correlation measures how consistently each replicate reproduces the same protein abundance pattern. In AP-MS experiments, Spearman correlations above 0.80 between replicates indicate good technical reproducibility.

Each cell shows the pairwise Spearman rank correlation between two replicates within the same group (sample or control). Green cells indicate strong agreement (r >= 0.80), yellow indicates moderate agreement (0.60 <= r < 0.80), and red indicates poor agreement (r < 0.60). The number of shared non-missing proteins is shown on hover.

Missingness Asymmetry
What is this?

Missing values are common in AP-MS data because low-abundance proteins may not be consistently detected across replicates. However, if one replicate has substantially more missing values than others, it may indicate a failed injection or degraded sample.

Each bar shows the fraction of proteins with missing intensity values for one replicate. Bars are grouped by experimental group (sample/control). A replicate is flagged if its missing fraction is more than 2x the group median. Green indicates normal missingness, yellow indicates elevated missingness (2-3x median), and red indicates extreme missingness (>3x median).

Intensity Distribution Shape
What is this?

The shape of the intensity distribution reveals technical artifacts. Well-behaved AP-MS data on a log2 scale typically shows an approximately normal distribution. Bimodal distributions, heavy tails, or value spikes suggest instrument issues or data processing artifacts.

This section summarizes the distribution shape checks across all replicates. High excess kurtosis (>7) indicates heavy tails; negative kurtosis (<-1.2) suggests bimodality; spike fraction >10% indicates value pile-up at a boundary. These checks do not require corrective action but help contextualize downstream results.

PCA Sample-Control Separation
What is this?

Principal Component Analysis (PCA) reduces the full protein abundance matrix to its most important axes of variation. If the experiment worked well, the biggest source of variation (PC1) should separate your bait samples from your controls.

Each point represents one replicate. The x-axis (PC1) and y-axis (PC2) show the two largest axes of variation, with the percentage of total variance they explain. Sample points are colored separately from control points. If samples and controls overlap heavily on PC1, the experimental condition may not be the dominant effect, which could reduce the sensitivity of the Bayesian analysis.

Overview

Protein interactions were analyzed using **BayesInteractomics** v
(Julia v). A total of  proteins were evaluated
for interaction with the  bait protein using
 control and  sample experiment(s).

Evidence for each candidate interaction was assessed by three independent Bayesian models:
a detection probability model, an enrichment model, and a dose-response correlation model.
The evidence from all three models was then combined using
 to produce a single posterior probability for each protein.
At a Bayesian false discovery rate (FDR) threshold of 5%, 
proteins were identified as significant interactors ( at FDR ≤ 1%).

Copy to clipboard

Prior Specification Table

Extended Details

This table lists every prior distribution used in the analysis.
All priors are weakly informative unless stated otherwise. Prior sensitivity was assessed by
varying key parameters; see the Sensitivity tab for results.
Following BARG guidelines (Kruschke, 2021), each prior is justified in the rightmost column.

| Model | Parameter | Prior Distribution | Values | Justification |
| --- | --- | --- | --- | --- |

Detection Model

Extended Details

The detection model assesses whether a protein is observed more frequently in sample
experiments than expected by chance. It uses a Beta-Bernoulli framework: each protein's
detection rate is modeled with a Beta(, )
prior, and a Bayes factor quantifies the evidence for differential detection between
sample and control conditions.

The detection Bayes factor BF_10_ compares two hypotheses:
H_1_ (detection rate differs between sample and control) versus
H_0_ (detection rates are equal). The Beta(, )
prior is symmetric and weakly informative, centred at 0.5, allowing the data to drive inference.
A BF_10_ > 3 indicates moderate evidence for differential detection
(Kass & Raftery, 1995).

Enrichment Model

Extended Details

The enrichment model evaluates whether a protein is more abundant in sample than control
experiments. It uses a Hierarchical Bayesian Model (HBM) that estimates the log_2_
fold-change between conditions. The model accounts for variability both within and across
experiments through hierarchical priors on intensity means and precisions.

Intensity means are assigned Normal(mu=,
sigma=) priors, and precisions receive Gamma(shape=,
scale=) priors. The enrichment BF_10_ tests
H_1_ (log_2_FC ≠ 0) against H_0_ (log_2_FC = 0).
Posterior medians with 95% highest-density intervals (HDI) are reported for effect sizes.

Correlation Model

Extended Details

The correlation model tests for a dose-response relationship between bait protein
abundance and each candidate interactor across experiments. It fits a Bayesian linear
regression with a  likelihood and evaluates whether
the regression slope deviates meaningfully from zero (threshold: ).

The slope prior is
.
The correlation BF_10_ compares H_1_ (|slope| > )
against H_0_ (|slope| ≤ ).

Evidence Combination

Extended Details

Bayes factors from the three models are combined into a single posterior probability
using . This approach integrates complementary evidence
(detection frequency, enrichment magnitude, dose-response correlation) while accounting
for potential model misspecification.

Calibration

Extended Details

Posterior probabilities were recalibrated using Platt scaling fitted on
 synthetic datasets generated by a parametric simulation engine.
Calibration is applied only when it improves the Expected Calibration Error (ECE),
ensuring that the recalibrated probabilities are at least as reliable as the raw estimates.

Platt scaling fits a logistic transformation:
P_calibrated_ = logistic(a · logit(P_raw_) + b),
where a and b are estimated from simulation ground truth. The ECE safety guard
prevents calibration from degrading probability estimates. Synthetic data are generated
using fitted model parameters from the real data, preserving realistic correlation structures.

Analysis Parameters
What is this?

This table lists all the settings used to run your analysis, ensuring full reproducibility. If you share these parameters, another researcher can replicate your exact analysis.

Parameters include the number of control and sample replicates, the reference protein index, prior sensitivity ranges, evidence combination method (BMA, copula, or 3c-EM), and all model-specific settings.

| Parameter | Value |
| --- | --- |

Reproducibility
What is this?

This code block provides the exact Julia commands needed to reproduce this analysis, including all parameter settings and package versions.

Copy this code block into a Julia session with BayesInteractomics installed to regenerate this report. The package version and all configuration values are captured for exact reproducibility.
